# Fast, scalable, and statistically robust cell extraction from large-scale neural calcium imaging datasets

**DOI:** 10.1101/2021.03.24.436279

**Authors:** Fatih Dinç, Hakan Inan, Oscar Hernandez, Claudia Schmuckermair, Omer Hazon, Tugce Tasci, Biafra O. Ahanonu, Yanping Zhang, Jérôme Lecoq, Simon Haziza, Mark J. Wagner, Murat A. Erdogdu, Mark J. Schnitzer

## Abstract

State-of-the-art Ca^2+^ imaging studies that monitor large-scale neural dynamics can produce video datasets that tally up to ∼100 TB in size (∼10 days transfer over 1 Gbit/s ethernet). Processing such data volumes requires automated, general-purpose and fast computational methods for cell identification that are robust to a wide variety of noise sources. We present EXTRACT, an algorithm that is based on robust estimation theory and uses graphical processing units (GPUs) to extract neural dynamics from a typical Ca^2+^ video in computing times up to ∼10-times faster than imaging durations. We extensively validated EXTRACT on simulated and experimental data and processed 199 public datasets (∼12 TB) from the Allen Institute in a day. Showcasing its superiority over past cell extraction methods at removing noise contaminants, neural activity traces from EXTRACT allow more accurate decoding of animal behavior. Overall, EXTRACT is a powerful computational tool matched to the present challenges of neural Ca^2+^ imaging studies in behaving animals.

## INTRODUCTION

State-of-the-art neural Ca^2+^ imaging experiments, such as those using fluorescence macroscopes^1,2^, can generate up to ∼300 MB of data per second, or >1 TB per hour of recording.

Faced with such data volumes, neuroscientists need computational tools that can quickly process extremely large datasets without resorting to analytic shortcuts that sacrifice the quality of results. A pivotal step in the analysis of many large-scale Ca^2+^ imaging studies is the extraction of individual cells and their activity traces from the raw video data. The quality of cell extraction is critical for subsequent analyses of neural activity patterns, and, as shown below, superior analytics for cell extraction lead to superior biological results and conclusions.

Prior methods for cell extraction identified neurons as regions-of-interest (ROIs) by manual^3–7^, semi-automated^8^ or automated image segmentation^9–14^, which in turn allowed Ca^2+^ activity in each ROI to be determined using either the identified spatial masks or multivariate regression. Other cell extraction methods, including independent components analysis (ICA), non-negative matrix factorization (NMF), and constrained non-negative matrix factorization (CNMF), simultaneously infer cells’ shapes and dynamics using a matrix factorization^15–20^. In these now widely used methods, the Ca^2+^ movie is treated as a matrix that can be approximated as the product of lower dimensional spatial and temporal matrices. The detailed assumptions about this factorization differ between the approaches and influence their relative strengths and limitations. Together, extant cell extraction methods have enabled Ca^2+^ imaging studies with a wide variety of microscopy modalities and model species, but they have generally neither been created for nor validated on TB-scale datasets.

Notwithstanding the many past successes of Ca^2+^ imaging, neuroscientists face important computational challenges as Ca^2+^ imaging technology continues to progress rapidly. Many datasets contain noise that is not Gaussian-distributed, including background Ca^2+^ signal contaminants from neuropil or neural processes, weakly labeled or out-of-focus cell bodies, and neurons that occupy overlapping sets of pixels^21^. For simplicity, prior algorithms have typically used signal estimators to infer cellular Ca^2+^ traces by assuming Gaussian-distributed contamination^11,16,18,22–25^. Thus, these prior methods poorly handle the non-Gaussian contaminants found in real experimental situations, impeding the detection of cells and inference of their Ca^2+^ activity patterns. To mitigate the resulting estimation errors, past research has proposed video processing methods to be applied prior to^23,24^ or during cell extraction^18,19^. Other mitigation strategies involve post-processing of the estimated Ca^2+^ activity traces after cell extraction^26–28^. However, a reliance on specific image processing routines can restrict a cell extraction algorithm’s utility to the particular imaging conditions or modalities for which these routines were designed, and post-processing of traces generally adds greatly to the total processing time, which is not a practical option with 10–100 TB datasets. To date, no cell extraction algorithm (i) addresses the challenges of Ca^2+^ imaging within a single, generally applicable conceptual framework and (ii) is demonstrably capable of handling the field’s burgeoning data volumes.

An important recent direction in data analytics for neuroscience has been to use deep learning approaches^28–30^, notwithstanding that they are usually much slower than traditional statistical methods. However, if trained on data from a range of imaging modalities, deep learning can provide accurate cell extraction results for in-distribution datasets, *i*.*e*., cell-types and imaging conditions that were represented in the training set. However, the mammalian brain purportedly has >5000 neuron-types^31^, and neural imaging conditions are variable across different instruments, modalities, fluorescence labeling conditions, noise distributions, surgical preparations, and animals—even within the same lab. Unfortunately, the degree to which extant deep learning based approaches to cell extraction need retraining for experimental conditions not represented in the training set has not been formally examined. This is a crucial issue, as optimization and retraining are unlikely to be feasible for every neuron-type and set of experimental conditions. Moreover, deep learning approaches are prone to hallucinations^32^, which, in the context of Ca^2+^ imaging, can lead to inferences of Ca^2+^ activity that do not exist^33^. By comparison, statistical models do not require training data that are representative of the signal distributions and instead are usually based on easily explainable assumptions about the noise distributions. Further, due to their relative simplicity, cell extraction methods based on statistical models typically have much faster run speeds^30^. Overall, while deep learning approaches to cell extraction represent a very promising frontier, they do not yet fulfill the goal of having a cell extraction algorithm that is simultaneously robust, verifiably applicable to many different imaging conditions, and sufficiently fast to process the largest Ca^2+^ imaging datasets.

Here we present a broadly applicable, fast cell extraction method that addresses the experimental limitations of real Ca^2+^ imaging datasets and efficiently processes the largest available such datasets, while avoiding assumptions specific to particular imaging modalities or fluorescence labeling patterns. Using the theoretical framework of robust estimation, we describe a minimally restrictive model of data generation and derive a statistically robust method to identify neurons and their fluorescence activity traces. Robust estimation is widely used in statistics, as it provides a potent means of analyzing data that suffers from contamination, such as outlier data points, whose statistical properties differ from those of an assumed noise model (typically Gaussian). Instead of modeling the contamination statistics, robust estimation provides statistical estimates that have quality guarantees even in the case of the worst possible contamination.

One obtains these guarantees by constructing a statistical estimator that selectively downgrades the importance of contaminated, outlier observations, to which non-robust estimators generally assign undue weight^34,35^. In the presence of Gaussian-distributed noise plus non-Gaussian outliers, non-robust estimators can suffer enormous errors, whereas a suitable robust estimator can have negligible error^34^. In cell extraction, robust estimation can formally account for spatiotemporally varying, non-Gaussian contaminants and infer neural activity with high fidelity, without having to explicitly model the contaminants in Ca^2+^ imaging experiments. The result is a modality-agnostic approach that makes minimal assumptions about the data. We term the algorithm **EXTRACT** (for **<u>EX</u>**TRACT is a **<u>t</u>**ractable and **<u>r</u>**obust **<u>a</u>**utomated **<u>c</u>**ell extraction **<u>t</u>**echnique).

EXTRACT performs quickly and accurately due to its native support for graphical processing units (GPUs) and custom-designed fast convex solver. The resulting efficiency allows EXTRACT to handle individual Ca^2+^ movies up to ∼12 TB in size, up to 2 orders-of-magnitude larger than the validated range of prior cell-extraction algorithms^20^. For a typical imaging study, processing times with EXTRACT are an order-of-magnitude briefer than the imaging session. Even with Ca^2+^ videos from recent fluorescence mesoscopes, EXTRACT runtimes on a standard personal computer are comparable to imaging durations.

We first extensively validated EXTRACT on simulated data and benchmarked it against other algorithms in 33 computational experiments spanning a wide range of challenging imaging conditions. We then analyzed experimental data from conventional, multi-plane, and mesoscopic two-photon imaging studies in head-fixed behaving mice, one-photon miniaturized microscopy studies in freely behaving mice, and the Allen Brain Observatory two-photon Ca^2+^ imaging datasets^36^. When studying data from behaving animals, we focused on how EXTRACT led to superior biological results, due to the improved quality of the Ca^2+^ activity traces as compared to those from prior algorithms. Specifically, we show enhanced identification of anxiety-encoding cell populations in the ventral hippocampus, improved identification of anatomically clustered neural activity in the striatum, and more accurate predictions of mouse location via decoding of hippocampal neural ensemble activity, all using Ca^2+^ activity traces from EXTRACT.

## RESULTS

### A defect of conventional cell extraction: *L*_2_ loss is suboptimal for realistic datasets

We first illustrate the substantial shortcomings of conventional cell sorting algorithms by using a toy model in which the Ca^2+^ movie, ***M***, contains a single neuron, has a field-of-view *h* × *w* pixels in size, and is *n*_*times*_ frames in duration (**Fig 1A**; **Fig. S1A–D**). Without loss of spatial information, we refer to the two spatial dimensions using a single scalar variable whose values have a 1:1 correspondence to points in the *x-y* plane. In this notation we can describe ***M*** as an *n*_*pixels*_ × *n*_*times*_ array, where *n*_*pixels*_ equals the total number of pixels, *hw*. Within this description, the column vector ***s*** (of size *n*_*pixels*_) denotes the cell’s spatial profile, and the row vector ***t*** (of size *n*_*times*_) denotes its Ca^2+^ activity trace. Initially, ***t*** is unknown. Our goal is to obtain an optimal estimate of the cell’s activity trace, 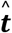, such that the square of the residual deviation, 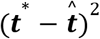, from its directly measured activity, ***t*** ^*^, is minimized. Unfortunately, unlike with *e*.*g*., intracellular electrical recordings of neural activity, we do not have a single and direct measurement, ***t*** ^*^, of the cell’s activity trace. Instead, we have a set of experimental readouts, in the form of fluorescence traces from the pixels of a Ca^2+^ movie, from which we must infer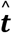. Therefore, since we cannot directly optimize 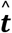 to minimize 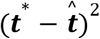, it has been conventional in the field to solve a substitute problem that both approximates the original goal and is mathematically tractable^15,18,19,23,26^.

**Figure 1.**
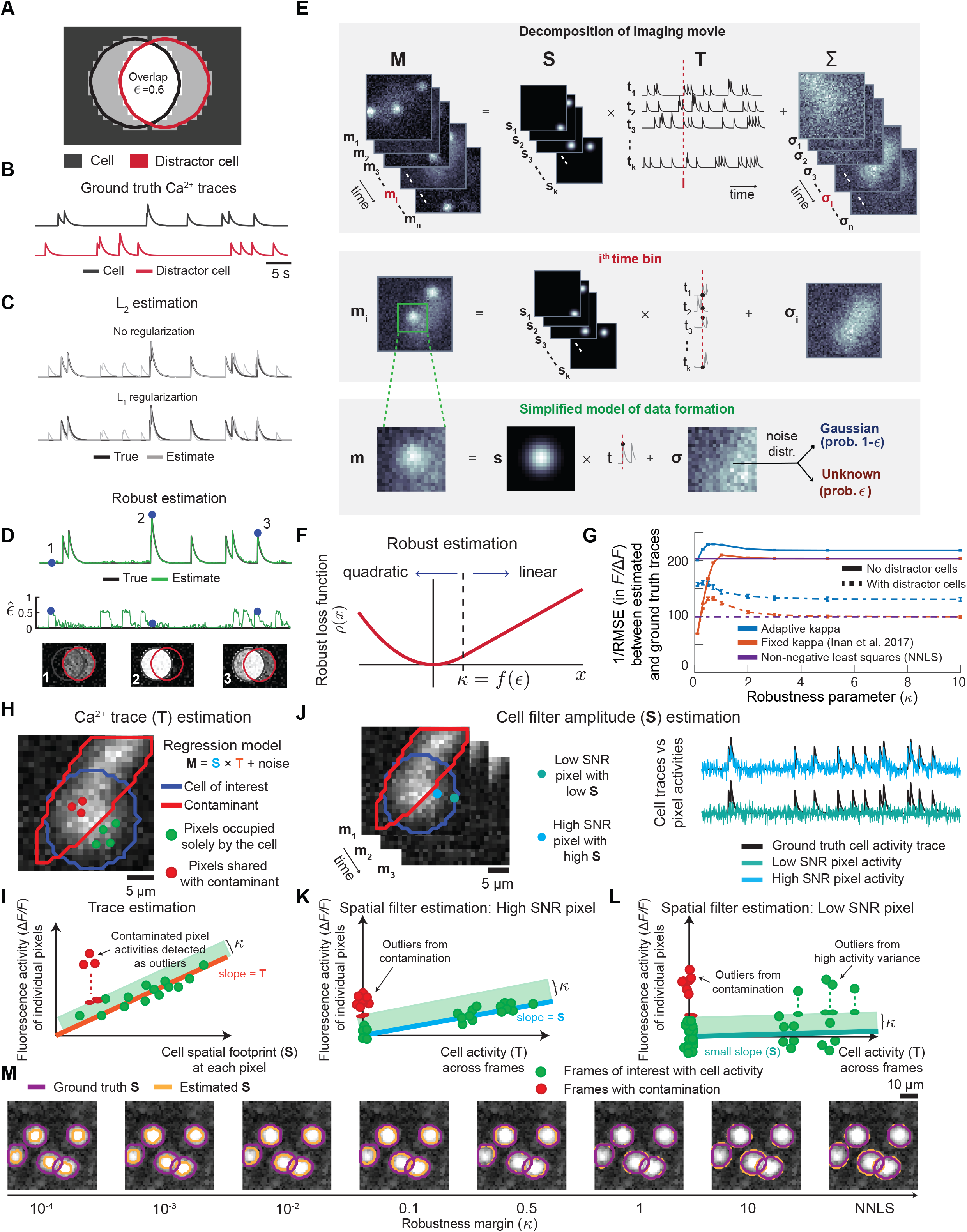
A robust estimation framework for extracting cells from Ca^2+^ video datasets. **(A)** *To showcase a basic limitation of using standard L*_2_ estimation to infer Ca^2+^ activity, we simulated an example movie with one cell of interest plus one distractor cell, which is not explicitly accounted for during the estimation process, *i*.*e*., with a corresponding spatial filter. Both cells had binary-valued images. The value of ϵ characterizes the time-dependent severity with which fluorescence photons from the distractor cell are detected in movie pixels that overlap with the image of the cell of interest. We analyzed the inference of Ca^2+^ activity in the presence of the distractor cell. **(B)** Examples of actual, ground truth Ca^2+^ traces for both cells in **(A)**. **(C)** Example results showing that *L*_2_ estimation leads to inferred Ca^2+^ activity for the cell of interest that is contaminated by the activity of the distractor cell. The addition of an *L*_1_ regularization penalty shrinks the inferred activity toward zero uniformly but does not mitigate the crosstalk. **(D)** With robust estimation, the activity trace of the cell of interest is accurately reconstructed without explicit knowledge of either the existence or the spatiotemporal characteristics of the distractor. Further, robust estimation infers the time-dependent level of contamination, ϵ, from the distractor cell onto the image of the cell of interest. **(E)** Our computational model of a Ca^2+^ video dataset represents the detected photons as originating from spatially localized cellular sources with time-dependent emission intensities, plus noise. ***Top***, A movie dataset is treated as a three-dimensional matrix, ***M***, that can be decomposed into cells’ fluorescence emissions plus contributions from noise. The first component is the product of the set of the cells’ spatial images, represented as a three-dimensional matrix, ***S***, and their activity traces, represented as a two-dimensional matrix, ***T***. The noise component, ***∑***, is additive, can vary over space and time, and can have non-Gaussian components. ***Middle***, A schematic representation of the problem of inferring the cells’ time dependent fluorescence emission amplitudes for a single time bin, *i*.*e*., one column of ***T***, given ***M*** and ***S***. Each frame of the movie, ***M***, is represented as a sum of the individual cell images, with each one weighted by the cell’s scalar valued Ca^2+^ activity at the selected time bin, plus noise. ***Bottom***, A simple application of the above model to a movie frame with one cell. Unlike conventional statistical approaches in which noise is assumed to be Gaussian-distributed, here the noise can have an unknown, non-negative contamination component that is subject to no other assumptions. **(F)** Using a framework for robust statistical estimation, we identify a loss function, ρ, that achieves optimal estimates with the least possible mean squared error (MSE) given the worst possible form for the unknown noise distribution with support on [κ,∞). ρ has a quadratic dependence for negative arguments and for positive arguments below a threshold value, κ. For arguments greater than κ, ρ rises linearly; this is what renders robust estimation relatively impervious to occasional but large non-negative noise contaminants, which typically skew conventional estimation procedures. Given the movie, ***M***, and an estimate of either its spatial or temporal components, ***S*** or ***T***, estimating the other component simply involves minimizing the residuals between the estimated and actual movie data (see **Methods**). **(G)** We compared 3 different approaches for estimating Ca^2+^ traces. For each method, we allowed the extraction algorithm to use the cells’ ground truth spatial profiles and computed the root mean square error (RMSE) between the estimated and actual Ca^2+^ traces. We assessed results using fixed (red curves) or adaptively varying (blue curves) values of κ. We also examined non-negative least squares (NNLS) regression (purple lines). The datasets comprised simulated Ca^2+^ movies with 600 cells each. The graph has plots of RMSE^−1^ as a function of either the fixed value of κ or the initial κ value when adaptive κ(*x, t*) estimation was used. The movie contained only Gaussian noise, a small portion (5%) of which was correlated across nearby cells. Solid lines: results for trace estimation when the movie reconstruction was performed using all cells’ spatial profiles. Dashed lines: results when 20% of the cells were designated as distractors, whose spatial profiles were not provided to the regression algorithm. Error bars: s.d. across 20 simulated movies. **(H–L)** To illustrate in more detail how robust estimation successfully handles contaminants, these panels depict analyses of a simulated movie with one target cell and one contaminant (representing *e*.*g*. neuropil activation). Unlike in **(A)**, ground truth images of both the cell and the contaminant are continuously valued. **(H, I)** To estimate the cell’s activity trace, each frame is examined independently. For pixels at which the cell’s spatial filter, ***S***, is non-zero, the pixel values from the filter are regressed against those from the movie frame. Some of these pixels, **(H)**, may overlap the footprint of the unknown contaminant (3 example pixels of this kind are marked with red dots), but others are occupied solely by the cell of interest (4 example pixels marked with green dots). The cell’s activity level in each movie frame is determined as the slope of the regression. Each datum in the regression plot, **(I)**, corresponds to one pixel in the cell’s spatial filter; the *x*-axis value denotes the pixel’s value in the spatial filter, ***S***; the *y*-axis value denotes the pixel’s brightness in the movie frame under analysis. The slope of the regression gives the cell’s estimated activity in this frame. Unlike conventional linear regression, our implementation of robust regression uses an iterative approach that handles outlier data points (*i*.*e*., from contaminant pixels) through two geometrically interpretable steps. First, outlier data points (depicted in red; defined as those points whose orthogonal distance to the regression line determined in the previous iteration is more than κ) are projected to the edge of the non-outlier zone (green shading; red ellipses at the edge of the green shading mark the projected values of the 3 red outlier points). Second, a conventional least-squares regression is performed, but using the projected values determined in step one for the outlier data points. This two-step process is iteratively repeated until the regression results converge. Crucially, by projecting outlier data points to the edge of the non-outliner zone, robust regression is ‘robust’ or impervious to the exact values of the outlier data. **(J–L)** To estimate the cell’s spatial filter, the fluorescence activity within each pixel of the movie is considered separately. Some pixels report the cell’s dynamics at a high SNR, whereas other pixels report the dynamics at low SNR values (see example pixels in **(J)** marked in navy blue and cyan, respectively). To determine each pixel’s contribution to the cell’s spatial filter, the cell’s estimated activity trace is regressed against the trace of the pixel’s fluorescence dynamics **(K)**. The cell’s activity level in each movie frame is determined as the slope of the regression. Each datum in the regression plot, **(K)**, corresponds to one time point in the movie; the *x*-axis value denotes the cell’s estimated activity level at that time point, ***T***; the *y*-axis value denotes the pixel’s brightness at that same time point. The slope of the regression gives the pixel’s weight in the cell’s spatial filter. Thus, pixels with high SNR reports of the cell’s activity yield regressions with substantial values of the slope, **(K)**, whereas low SNR pixels yield lower values of the slope, **(L)**. As for trace estimation, **(H, I)**, estimation of the spatial filter relies on a robust regression implemented in an iterative manner with two main steps, projection of the outlier points to the edge of the non-outlier zone (green shading in **(K, L)**) followed by a conventional linear regression using the projected values (red and green ellipses) of the outlier data points. There are two main classes of outlier points. Time bins at which the pixel reports contaminants from Ca^2+^ activity sources outside the cell fall near the *y*-axis (red data points in **(K, L)**). Additionally, for low SNR pixels, **(L)**, the variance between the pixel’s activity level and the estimate based on the cell’s activity trace can sometimes exceed κ (green data points above the green shading). **(M)** A demonstration, using the same movie simulation protocol as in **(G)**, that varying the robustness margin, κ, changes the estimated boundaries of the cells. Cell boundaries (orange lines) are shown for analyses of identical Ca^2+^ video data with different κ values. For small κ values, robust regression rejects from a cell’s spatial filter the pixels with low SNR versions of the cell’s activity trace, as evidenced by the estimated cell boundaries lying within the actual ground truth boundaries (purple lines; defined as contours 1.5 σ from the centroid of each Gaussian-shaped cell). As κ increases, the estimated boundaries expand and can even lie outside the actual cell boundaries. NNLS is equivalent to robust regression with an infinite value of κ.

This substitute problem is that of ‘movie reconstruction’. Namely, one estimates the trace, 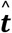, so the outer product, 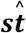, well approximates the Ca^2+^ movie, ***M***, and minimizes the magnitude of the residual,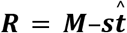. This is usually done by minimizing the sum of the squared elements of ***R*** with respect to 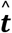. In other words, one minimizes an *L*_2_ (*i*.*e*., quadratic) loss function on the residual; the resulting estimator is referred to as an ‘*L*_2_ estimator’. This widespread method of estimating Ca^2+^ activity rests on an implicit assumption that ***R*** is Gaussian-distributed. If ***M*** contains the cell’s activity plus additive Gaussian noise, then the *L*_2_ estimator is optimal in that it reconstructs an estimated movie,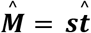, which differs from the real movie only by irreducible noise^37^.

However, in reality, Ca^2+^ imaging data are corrupted not just by Gaussian noise but also other contaminants, such as from neuropil Ca^2+^ activity, out-of-focus neurons, or cells with overlapping pixels. In the realm of optimization problems, this situation is unusual in that signals from one cell may also constitute noise contaminants that degrade the estimated activity trace of an overlapping cell (See **Supplementary Note 1**). For instance, if one adds to our toy model with one cell a partially overlapping ‘distractor’ cell, whose spatial profile was not identified by the cell extraction algorithm, this simple addition greatly impedes the estimation of Ca^2+^ signals from the first cell. Specifically, using an *L*_2_ loss function can lead to crosstalk from the distractor cell in the estimated trace, 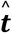, for the first cell—even when regularization enforcing sparsity is used (**Fig. 1B–D; Fig. S1A–D**). Thus, traditional *L*_2_ estimators of neural signals^18–20,23^ can be inaccurate in realistic scenarios.

### Robust statistical estimation of neural Ca^2+^ dynamics

We start our presentation of robust estimation by first relaxing the common assumption that noise is Gaussian-distributed. Signal contaminants may exist with spatially irregular and temporally non-stationary properties; this can occur when neighboring cells occupy overlapping sets of pixels or when Ca^2+^ signals arise from neuropil or out of focus cells, leading to signal sources that are absent from or poorly represented in the extracted set of cells’ spatial filters. Especially when the cells of interest are quiet, such signal contaminants can greatly exceed the Ca^2+^ signals we aim to extract. Second, we note that since nearly all fluorescent Ca^2+^ reporters have a rectified dynamic range, positive-going [Ca^2+^] fluctuations are reported far more strongly than negative-going fluctuations of [Ca^2+^] or fluorescence levels below baseline values. Based on these points, we model the noise distribution as having two components, each of which can vary spatiotemporally (**Fig. 1E,F**). The first noise component is assumed to be Gaussian-distributed and to affect a fraction, **1** − ϵ(*x, t*), of the pixel intensity measurements. The second component has an unknown distribution, *H*, and affects the remaining fraction, ϵ(*x, t*), of the measurements. We assume nothing about *H*, except that it yields non-negative measurement values, due to the rectification of the Ca^2+^ indicator. (More precisely, *H* has support on [κ(*x, t*), ∞), where κ(*x, t*) is a non-negative parameter that is linked to the severity of the contamination; typically, κ is on the order of 1 s.d. or less of the baseline noise fluctuations that persist after pre-processing, but, as discussed below, EXTRACT estimates spatiotemporally varying values of κ(*x, t*) in an adaptive manner; **Methods**).

With this noise model, what is a suitable loss function for estimating cells’ Ca^2+^ signals? The reliance on Ca^2+^ movie data, rather than direct measurements of neuronal dynamics, and the lack of a prescribed noise distribution for *H* prevents identification of an optimal loss function for estimating the Ca^2+^ activity trace, 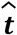. However, by using the theory of robust statistics^34,35^, we can find a loss function that is optimal in a different sense, namely that it achieves the best MSE under the worst possible probability distribution that the unknown noise could ever assume. This loss function smoothly transitions between a quadratic function and a linear function, with the transition occurring at the positive value, κ(*x, t*), that should depend on the prevalence, ϵ(*x, t*), of the unknown noise component (**Fig. 1F; Fig. S1F**). We call this the one-sided Huber loss function, in reference to the traditional Huber loss function in which non-Gaussian, outlier noise contaminants can have either sign^34^. Our loss function is thus tailor-made for the rectified signals of Ca^2+^ imaging, and it leads to mathematically provable upper bounds on the worst-case estimation errors in the limit of scalar estimation, *i*.*e*., the case of sparsely labeled neurons with uniform spatial profiles (Proposition 1 in **Methods**). A key consequence of using this loss function is that, when the first derivative of the total loss is computed across all data points for the sake of minimization, any datum at which the residual is more than κ contributes only a fixed value, 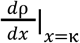, to the derivative, namely the same as if the datum had a residual of exactly κ. (This fact leads, below, to an appealing geometric interpretation for how our robust solver operates).

The simplest approach to robust regression uses fixed^35^ values of ϵ(*x, t*) = ϵ and κ(*x, t*) = κ for all pixels and frames. However, given that real Ca^2+^ movies have spatiotemporally varying levels of noise and contamination, we need a more sophisticated loss function, in which the matrices ***ϵ*:** = ϵ(*x, t*) and ***κ*:** = κ(*x, t*) vary in space and time and must be estimated from the movie (**Fig. 1D; Fig. S1C, D**). To do this, one iteratively seeks better estimates of ***ϵ*** and ***κ*** in a closed loop, while concurrently performing robust estimation with the heterogeneous ***κ*** (**Methods**). In this way, one can let the data dictate, across each frame and time bin, how the loss function should differ from its conventional *L*_2_ form to accommodate the instantaneous, non-Gaussian contaminant components. Returning to our toy model with one cell of interest and one distractor cell, our robust loss function can estimate the first cell’s Ca^2+^ trace accurately, while ignoring signals from the distractor (**Fig. 1C,D; Fig. S1B,D**).

To illustrate the utility of robust regression with adaptive estimation of ***κ***, we simulated a set of 20 Ca^2+^ movies, each containing 600 cells with non-negative Ca^2+^ signals (**Fig. 1G**; **Fig. S1E**). The movies contained Gaussian noise, but a small portion of it (5%) was correlated across nearby cells (**Methods**). Using the cells’ ground truth spatial profiles, we obtained estimates of the cells’ Ca^2+^ activity traces via robust regression using either a fixed κ or adaptively varying values of ***κ***. For comparison, we also used an *L*_2_ estimator to compute estimated Ca^2+^ traces that constitute the non-negative least squares (NNLS) solution to the Ca^2+^ movie reconstruction. In our comparisons, we considered two cases. In the first case, the spatial profiles of all the cells were input to the cell extraction algorithms. In the second case, the spatial profiles of only 80% of the cells were input to the extraction algorithm, and 20% were deemed distractor cells whose spatial profiles were not input. These distractor sources of Ca^2+^ activity lead to non-Gaussian noise distributions. Notably, in both cases, NNLS yielded the best reconstruction of the Ca^2+^ movie, whereas robust regression with adaptive estimation of ***k*** provided the smallest error on the estimated Ca^2+^ traces (**Fig. 1G**). This underscores the differences between the two estimation problems—that of reconstructing the movie *vs*. estimating the traces—and that robust regression outperforms NNLS even when the noise distribution deviates only slightly from independent Gaussian noise at each pixel. Robust regression with fixed κ values yielded modest gains over NNLS; however, these gains were only obtainable over a limited range of κ values, which a user would have to know *a priori* to choose κ appropriately.

To complement these results, we performed additional studies of simulated movies in which the pixels occupied by target cells included non-Gaussian noise contaminants arising from varying percentages of target and distractor cells. In these cases, robust estimation with adaptive estimation of ***κ*** was needed to extract neural signals most accurately for the target cells (**Fig. S1E**).

### Geometric interpretation of robust regression

Our robust loss function, the one-sided Huber loss, is designed to minimize the influence of outlier, non-Gaussian contaminants on the estimated solution to the problem of Ca^2+^ movie reconstruction. The custom robust solver that we created to minimize this loss function has an intuitive geometric interpretation (**Fig. 1H–M; Supplementary Note 2**). The minimization is achieved by iterating the following pair of steps, comprising an adjustment to the dataset followed by a regression. In the first step, any outlier data points for which the residual from the prior round of regression exceeds κ are moved to the robustness boundary, defined as the set of data points with a positive residual equal to κ (**Fig. 1I–L**). This step is a geometric implementation of the statement that outlier points should all have the same impact, 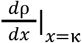, on the loss minimization. Second, a least-squares regression is computed for the adjusted dataset. These two steps are iterated until the process converges. In this way, the robust regression detects and weights outlier data points, which can arise, *e*.*g*., from the activity of distractors, so as to reduce their effects on signal estimates (**Fig. 1H, I)**.

The geometric interpretation of our custom solver for performing regressions with a one-sided Huber loss function also has implications for the estimation of cells’ spatial profiles. To this point, we have discussed the determination of cells’ estimated activity traces, 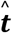, given cells’ known spatial profiles. However, for the analysis of real Ca^2+^ movies, the cells’ spatial profiles must also be estimated. Our extraction algorithm outputs both 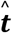 and 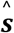, and it finds these estimates through iterative, alternating estimates of 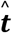 using fixed 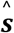, and then *vice versa*. The estimation of 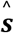 given a fixed 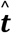 is also done using regression with a one-sided Huber loss, which minimizes the impact of movie pixels at which a target cell’s activity is represented but with low signal-to-noise ratio (SNR), and of outlier data points reflecting times when there was Ca^2+^ activity in distractor cells overlapping the target cell (**Fig. 1J–L**). Notably, for 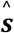 estimation, unlike for 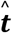 estimation, we use a fixed value for κ, for which lower κ values lead robust regression to minimize the weights in 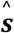 assigned to low SNR pixels, which often lie at the cell periphery (**Fig. 1M**). By comparison, cell extraction by NNLS led to estimates of 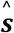 that included pixels that were nearby but outside the cell perimeter and had no clear Ca^2+^ signal, whereas robust regression using κ ∼**1** s.d. of the movie noise led to estimated spatial profiles that were comparable to the cells’ actual boundaries (**Fig 1M**). Thus, the use of a robust loss function in the filter estimation assigns higher weights in 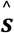 to pixels from the movie that are highly informative about the cell’s activity.

### Introducing EXTRACT: A pipeline for cell extraction using robust regression

Using our loss function and robust estimation, we now treat real data by going beyond our toy model and introduce the EXTRACT pipeline. We consider a Ca^2+^ movie, ***M***, that is a linear combination of background signal contaminants plus Ca^2+^ signals from an unknown number of cells, each of which contributes to the movie an activity trace given by the product of its spatial and temporal weights (**Fig. 1E**). We accomplish cell extraction by first performing a simple (and optional) pre-processing of the movie frames, followed by 3 main computational stages (**Fig. 2A**). A conference paper^35^ outlined our initial ideas for EXTRACT, but the EXTRACT pipeline described here and the implementations of each stage are new, use a new customized optimization solver (**Methods**; **Supplementary Note 2**), and are now rigorously validated against other cell extraction methods (below). Crucially, here we use adaptive estimation of ***κ*** = κ(*x, t*), unlike our proof-of-concept study^35^.

**Figure 2.**
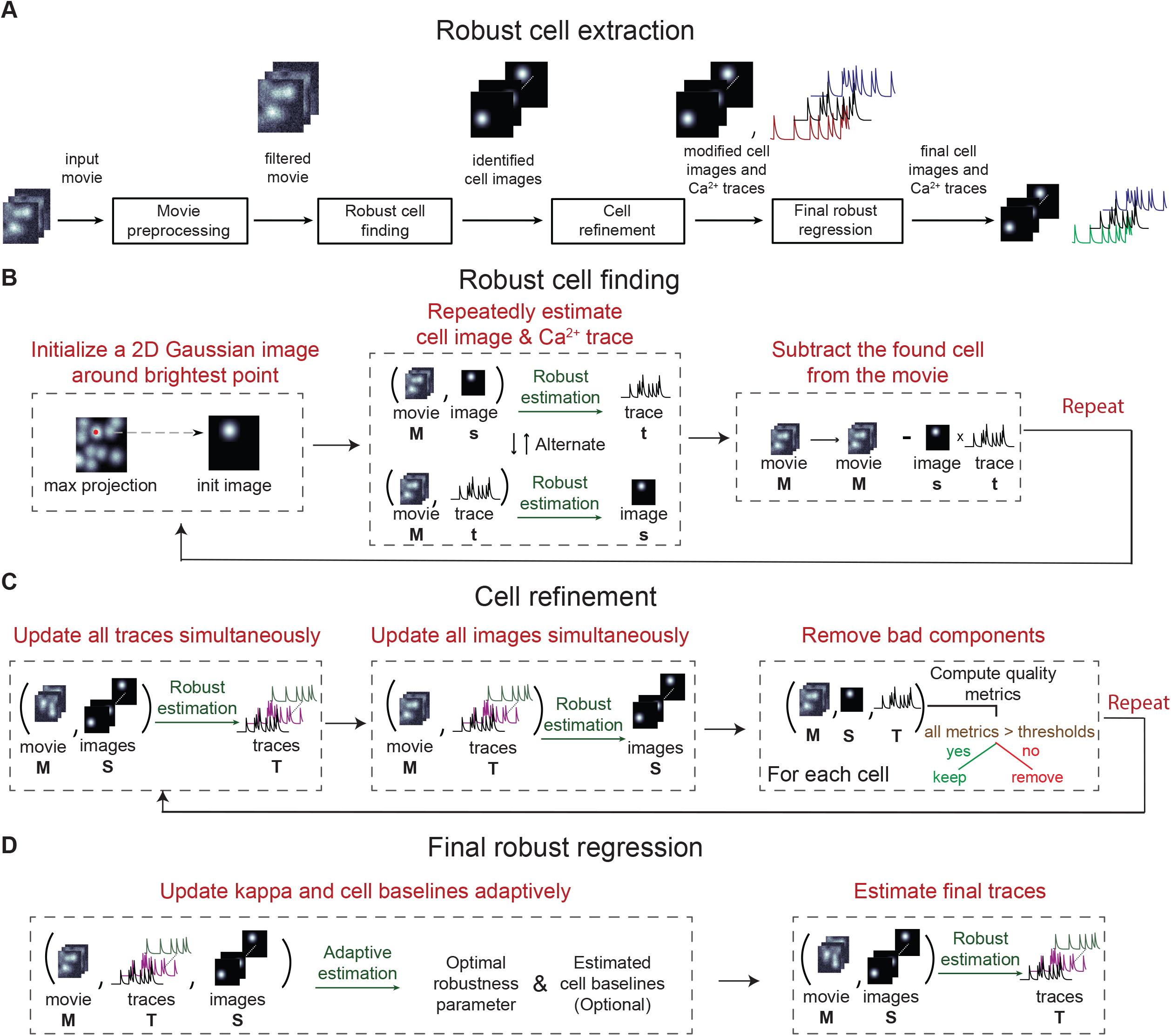
Automated identification of neurons and their Ca^2+^ activity traces with EXTRACT. **(A)** The EXTRACT algorithm comprises an optional stage for preprocessing of the raw Ca^2+^ videos, followed by 3 primary stages of cell extraction. The preprocessing stage filters fluorescence fluctuations at coarse spatial scales that arise from neuropil Ca^2+^ activity. The robust cell finding stage identifies and then extracts individual cells in an iterative manner, using robust estimation to infer each cell’s spatial and temporal weights. In the cell refinement stage, these weights are updated through iterative, alternating refinement of first the spatial and then the temporal weights, again using robust estimation. To attain the final estimated set of Ca^2+^ activity traces, the final robust regression stage employs adaptive estimations of each cell’s baseline activity level and time-dependent κ value. **(B)** Robust cell finding is an iterative procedure that identifies individual cells in a successive manner. At each iteration, a seed pixel is chosen that attains the maximum fluorescence values among all the pixels, and a cell image is initialized around it. Next, this image and the cell’s activity trace are iteratively updated via alternating applications of robust estimation. When this process converges, the cell’s estimated fluorescence contributions to the Ca^2+^ video are subtracted from the movie, and then the entire process repeats for another cell. This procedure continues until the identified seed pixel has a maximum instantaneous SNR that falls below a minimum threshold, as determined by examining the s.d. of the pixel’s intensity fluctuations across the entire movie. **(C)** Cell refinement is also an iterative procedure. At each iteration, the set of estimated Ca^2+^ traces is updated via robust estimation while holding fixed the cell images; the cell images are then updated using robust estimation while holding fixed the activity traces; a set of quality metrics is computed for each putative cell, and putative cells for which one or more metric values lies below a minimum threshold are eliminated from EXTRACT’s final output. **(D)** The final robust regression is a trace regression (see **Figure 1I**), performed using the spatial filters obtained from the cell refinement stage.

The pre-processing step applies a high-pass spatial filter to ***M*** to reduce background fluorescence and then subtracts from each pixel value its baseline fluorescence level (**Methods**). The first main stage of computation, ‘Robust cell finding’, identifies cells in the movie. The second main stage, ‘Cell refinement’, hones the estimates of cells’ spatial profiles and activity traces. The third stage, ‘Final robust regression’, performs one final regression to reconstruct ***M*** and obtain a final set of estimated Ca^2+^ traces using the last set of spatial profiles provided by the Cell refinement stage. As with the toy model above, for which an *L*_2_ loss function led to crosstalk from a distractor cell, robust estimation allows the proper isolation of individual neurons from real data, even when there is substantial spatial overlap in cells’ profiles and temporal overlap in their activity patterns.

The cell-finding stage uses a simple, iterative procedure to find cells and applies robust estimation to determine each cell’s spatial profile and activity trace (**Fig. 2B**). At each iteration, the algorithm finds a seed pixel that attains the movie’s maximum fluorescence intensity, and it initializes a candidate cell image at the seed pixel (**Methods**). The algorithm then alternatively improves its determinations of the cell’s spatial profile and activity trace via robust estimation (**Fig. 2B**). After the estimates of the spatial profile and activity trace stabilize, the cell’s inferred activity trace is subtracted from the movie, and in the next iteration the steps above repeat for another cell. The cell-finding procedure ends when the peak value for the activity trace of the seed pixel fails to reach a threshold value, which is set as a fixed multiple of the standard deviation of the background noise. Notably, the use of robust estimation for cell finding is important for providing the subsequent cell refinement stage a set of good initialization values (**Supplementary Note 1**).

After cell finding, the ‘Cell-refinement’ stage improves the estimates of cells’ spatial and temporal contributions to the movie data, by accounting concurrently for all the identified cells using multivariate robust estimation (**Fig. 2C**; **Methods**). This stage is also an iterative procedure, and each iteration has 3 steps. First, all Ca^2+^ traces are simultaneously updated using robust estimation, while holding fixed the cells’ spatial profiles. Second, all spatial profiles are concurrently updated via robust estimation, while holding fixed the activity traces. Third, a validation procedure checks a set of predetermined metrics for every putative cell and removes any cell with metrics that fail to meet user-set criteria. This 3-step procedure repeats for a fixed number of iterations, and the algorithm outputs the final estimates of cells’ spatial profiles and activity traces. Compared to *L*_2_ estimation, the use of robust estimation in the Cell refinement stage allows faster convergence and better discarding of spurious or duplicate cells (**Supplementary Note 3**). The resulting spatial profiles are then used in the ‘Final robust regression’ module to estimate the final Ca^2+^ activity traces (**Fig. 2D)**. This regression is performed with additional adaptive estimation procedures for cells’ baselines to ensure high quality output (**Methods)**.

Crucially, to perform these computations efficiently, we developed a custom fast solver for robust estimation that combines the computational cost of a first-order optimization algorithm with a convergence behavior approaching that of second-order optimization algorithm, such as Newton’s method (**Proposition 2** in **Methods; Supplementary Note 2**). Our solver is expressly adapted for and benefits greatly from computational acceleration provided by multiple graphical processing units (GPUs) and parallel computation. Internally, the solver uses the alternating direction method of multipliers^38^ (ADMM), which enforces mathematical constraints by separating optimization and constraint enforcement into two distinct problems, each of which can be solved via matrix multiplications and projections (**Supplementary Note 2**).

### Performance evaluations using large-scale simulated Ca^2+^ activity datasets

To validate EXTRACT and benchmark it against other cell extraction methods, we created a set of large-scale, simulated Ca^2+^ video datasets (∼4 TB total), allowing us to evaluate cell extraction performance quantitatively and in greater depth than prior studies (see **Table S1** for a list of 33 different computational experiments and benchmarks). We simulated Ca^2+^ videos with varying densities of cells and levels of cell overlap (**Fig. 3A–E**), across a density range slightly beyond that seen in real datasets (*e*.*g*., compare the insets of **Figs. 3E** and **4A**). The simulated activity traces had discrete non-binary Ca^2+^ events with exponentially decaying waveforms (**Fig. 3A**; **Methods**). The movies also contained additive Gaussian-distributed noise that was uncorrelated between pixels, mimicking photon shot noise, plus varying levels of spatiotemporally correlated noise contamination, mimicking blood vessel contamination and/or neuropil Ca^2+^ activation.

**Figure 3.**
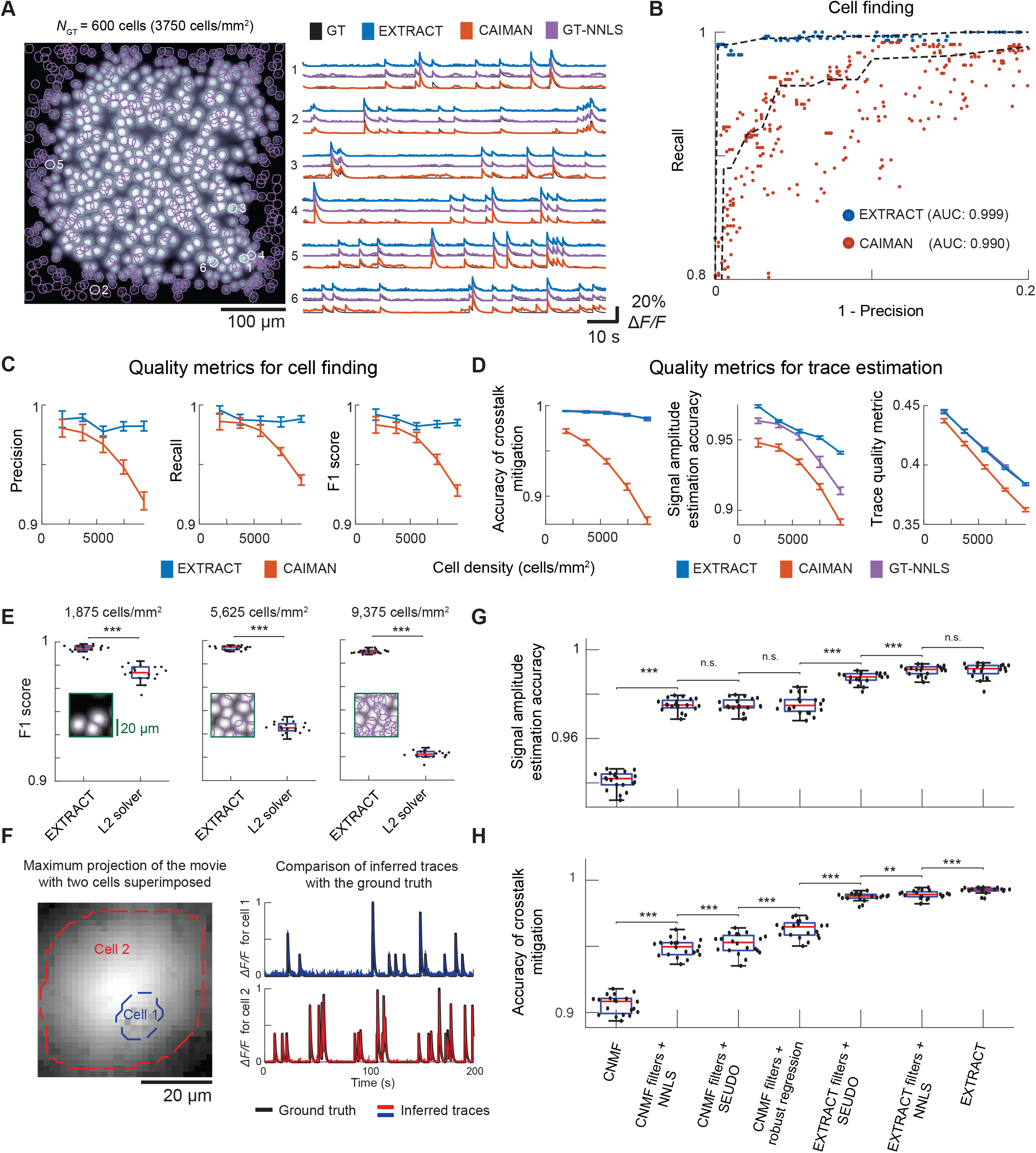
Benchmarking EXTRACT *vs*. state-of-the-art methods on simulated Ca^2+^ movies. **(A–H)** To evaluate EXTRACT against other widely used cell extraction methods, we created datasets of simulated one- and two-photon Ca^2+^ videos (∼4 TB in total), over a range of imaging conditions. **Table S1** describes 33 different numerical experiments we did. Illustrative results from experiments 12, 7, 2, 28 and 17 are respectively shown in panels **(B), (C, D), (E), (F)**, and **(G, H)**. **(A)** *Left*, An example simulated one-photon movie with 600 cells (*left*) is shown, with the cells’ footprints encircled in purple. 6 example cells whose activity patterns are shown in the right panel are encircled in white. *Right*, Ground truth (GT; black traces) and estimated Ca^2+^ activity traces obtained from EXTRACT (blue traces), CAIMAN (red traces), and via a non-negative least-squares regression of the cells’ ground truth spatial filters against the Ca^2+^ movie (GT-NNLS; purple traces), for the 6 numbered labels in the left panel. **(B)** To compare the cell finding capabilities of EXTRACT and CAIMAN, we simulated a two-photon Ca^2+^ movie with 600 cells. We then performed a precision-recall (PR) curve analysis by varying each algorithm’s hyperparameters in a nearly exhaustive manner (via grid search) and plotting the cell identification metrics, *i*.*e*., 1–precision and recall, for each method and set of hyperparameter values (**Methods**). EXTRACT was nearly impervious to the choice of hyperparameters and yielded a higher value of the area under the PR curve (AUC). Notably, EXTRACT achieved a 100% rate of identifying true positive cells with a 97% precision; CAIMAN reached the same mark only with a 70% precision. Dashed lines: median values of the true positive rate for each method, across the range of false positive rates and using a variable *x-*axis bin size to account for variations in the density of data points. **(C, D)** We used 6 different quality metrics to compare the cell finding **(C)** and trace estimation **(D)** capabilities of EXTRACT and CAIMAN on simulated one-photon Ca^2+^ videos across a wide range of cell densities. Error bars: s.d. over 20 simulated movies for each cell density value. For all 6 quality metrics, EXTRACT outperformed CAIMAN, especially for movies with high cell density. **(C)** The cell-finding precision is defined as the number of correctly identified cells divided by the total number of candidate cells returned by the algorithm. Recall is defined as the number of correctly identified cells divided by the total number of actual cells in the movie. F1 score is a summary metric equal to the harmonic mean of the precision and recall. **(D)** Under realistic, non-ideal conditions, the detection threshold needed to capture 100% of the real Ca^2+^ events also leads to a non-zero rate of false positive detection, which in turn depends on the severity of crosstalk between overlapping sources of Ca^2+^ signals. To assess the accuracy of crosstalk mitigation, we computed the area under the PR curve describing the tradeoff between the detection of true and false positive Ca^2+^ events as the Ca^2+^ event detection threshold is varied **(Methods)**. To compute the accuracy of signal amplitude estimation, for each cell identified in the movie we computed the Pearson’s correlation coefficient between the estimated and actual Ca^2+^ traces at the (actual) occurrence times of the cell’s Ca^2+^ events. We then determined the mean correlation coefficient, averaged across all identified cells. Finally, we computed a trace quality metric (TQM; validated in **Figure S3**) that assesses both signal amplitude estimation and crosstalk mitigation but does not require ground truth data and so can also be applied to real experimental data (**Figures 5H, 6D,E** and **7G, H**). The TQM is determined as the mean Pearson’s correlation coefficient, averaged across all detected cells, between a template of each cell’s image and the individual Ca^2+^ movie frames at the times of the cell’s detected Ca^2+^ events **(Methods)**. **(E)** To assess whether EXTRACT’s superior cell finding arises primarily from its use of a robust loss function, or, alternatively, from other aspects of its computational implementation, we compared EXTRACT to the use of a traditional *L*_2_ loss function, which we implemented in the EXTRACT pipeline by using a fixed robustness margin, κ, of a very high value (κ = 100). Thus, this comparison selectively isolates the performance impact of using the robust loss function. Across simulated one-photon Ca^2+^ videos with 3 different cell densities, EXTRACT consistently achieved more accurate cell finding, showing the virtue of using robust regression. *Insets*: Cropped maximum projection images showing typical levels of cell overlap at the 3 density levels. **(F)** We tested whether EXTRACT can facilitate cell extraction in even the extreme scenario in which the image footprint of one cell is fully encapsulated by that of another cell, we simulated a Ca^2+^ video with such two cells. EXTRACT correctly identified both cells and accurately estimated their Ca^2+^ traces without crosstalk. **(G, H)** To evaluate the Ca^2+^ activity traces from EXTRACT to those produced by a state-of-the-art post-processing algorithm intended to improve cell activity traces that may suffer from cross-talk, we simulated two-photon Ca^2+^ movies and performed cell extraction using CNMF and EXTRACT. Next, we ran the SEUDO post-processing algorithm to obtain putatively denoised traces, initializing SEUDO with the spatial filters provided by EXTRACT and CNMF. For comparison, we also tested the quality of Ca^2+^ traces found by using the same spatial filters together with either a robust or a conventional NNLS regression. For the resulting 7 different scenarios, we determined the accuracy of signal estimation **(G)** and crosstalk mitigation **(H)** for the estimated Ca^2+^ traces. EXTRACT achieved the highest values of both quality metrics, without any post-processing. The application of SEUDO after EXTRACT lowered both quality metrics. Box-and-whisker plots in **(E), (G)**, and **(H)** show results from 20 different conditions, denoted by individual data points; red lines denote median values; boxes span the 25th to 75th percentiles; whiskers extend to 1.5 times the interquartile range. Statistical comparisons between results of different algorithms were two-sided Wilcoxon signed-rank tests (*p < 0.05, **p < 10^−2^, ***p < 10^−3^).

When picking cell extraction algorithms to compare with EXTRACT, there were at least 15 options from the peer-reviewed published literature with openly available code (see **Supplementary Note 1**), making it impractical to run them all through a broad range of systematic comparisons. Therefore, from among the most widely used cell extraction routines described in peer-reviewed publications, we categorized algorithms into 3 categories: i) early-stage algorithms based on independent component analysis; ii) more recent, state-of-the-art algorithms that optimize reconstruction of the Ca^2+^ movie as a product of cells’ spatial filters and activity traces; iii) post-processing tools for improving estimated Ca^2+^ traces given a set of spatial filters. Based on its current and past use by hundreds of neuroscience labs, we picked PCA/ICA^16^ to represent the first category. In the second category, we chose CAIMAN^20^, which is widely used and has options for analyzing one- or two-photon Ca^2+^ movies, unlike a similar algorithm with a different implementation^19^. For the third category, we chose SEUDO, a state-of-the-art approach for improving Ca^2+^ trace estimation. To assess algorithmic performance, we supplemented standard measures of cell finding accuracy, such as precision and recall, with metrics that quantify various aspects of the cells’ estimated activity traces, such as the level of crosstalk from neighboring cells and the estimation accuracy of signal amplitudes (**Methods, Fig. 3**).

To start, we simulated large-scale one- and two-photon Ca^2+^ movies across varying cell densities, levels of neuropil contamination and correlated neural Ca^2+^ activity (**Figs. 3A–D; S2, S3, S4A)**. Notably, whereas EXTRACT makes no assumptions about Ca^2+^ event distributions or waveforms, CAIMAN is based on an assumption that Ca^2+^ transients have exponentially decaying waveforms. Thus, our simulated movies gave CAIMAN an inherent advantage, in that its formulation explicitly models the exponential Ca^2+^ event waveform. Notwithstanding this advantage to CAIMAN, EXTRACT outperformed CAIMAN in all scenarios tested.

Prior to detailed benchmarking, we ensured that each algorithm was run with a set of near optimal hyperparameters for the simulated datasets. To this end, we processed a representative movie with CAIMAN and EXTRACT using >1000 different hyperparameter configurations for each method, chosen via an extensive grid-search (**Methods**). Then, we computed the precision and recall of cell finding for each algorithm with each hyperparameter set (**Figs. 3B**; **S4A**). Whereas EXTRACT’s cell-finding accuracy was almost impervious to the choice of hyperparameters, with CAIMAN the cell-finding accuracy depended sensitively on the hyperparameter values. Even when fully optimized, CAIMAN fell short of EXTRACT’s cell-finding capabilities **(Fig. 3B; S4A)**.

In subsequent experiments, we used the optimized hyperparameter sets for CAIMAN and EXTRACT to benchmark cell finding and Ca^2+^ trace estimation accuracies on simulated one- and two-photon Ca^2+^ videos with a wide range of different conditions (**Figs. 3C,D; S3**). When analyzing one-photon movies with CAIMAN, we ran it with the option implementing the CNMF-E algorithm^23^, its preferred option for one-photon datasets. Especially with higher densities of cells—potentially representative of cutting-edge, real datasets—cell finding and trace estimation accuracies for CAIMAN but not EXTRACT declined substantially. Sometimes, EXTRACT’s trace estimation accuracy even surpassed that of a non-negative least squares (NNLS) regression using the cells’ ground truth spatial filters (**Fig. 3A**). Notably, estimated traces from CAIMAN commonly included false-negative and false-positive Ca^2+^ events (**Fig. S2**). The latter had exponentially decaying waveforms, making it hard to recognize these as false-positives based solely on their time courses.

Next, we tested whether the accuracy of EXTRACT arose from its use of robust regression, or perhaps merely from a favorable implementation of a software pipeline for video preprocessing cell extraction. To examine this point, we implemented an *L*_2_ solver in the EXTRACT software simply by using a fixed, effectively infinite value for ***κ***, which converts the robust solver into a NNLS solver. When then compared the performance of EXTRACT, run as normal with adaptive ***κ*** estimation, to that of the NNLS solver run in EXTRACT software. With simulated one- and two-photon Ca^2+^ movies with varying cell densities, EXTRACT outperformed the *L*_2_ solver at both cell-finding and trace estimation **(Figs 3E, S4B–E)**. The use of robust estimation also allowed the cell-refinement stage to converge more quickly (**Fig. S4B**; **Supplementary Note 3**).

Past work reported that *L*_2_ solvers can identify spatial filters and demix Ca^2+^ signals from cells that partially overlap^18^. To examine an extreme case of this scenario, we simulated a small, one-photon Ca^2+^ movie depicting an unusual scenario in which the image of an out-of-focus cell has an enlarged footprint that fully covers that of an in-focus cell (**Fig 3F**). EXTRACT correctly estimated both cells’ spatial filters and Ca^2+^ traces. Although at first blush this scenario might seem unrealistic, multi-plane one-photon movies can contain out-of-focus cells with enlarged footprints. To our knowledge, EXTRACT is the first extraction method to be tested in such a scenario.

Next, we compared EXTRACT to SEUDO, a state-of-the-art post-processing approach to trace estimation^26^. While a post-processing method cannot correct any inaccuracies in cell-finding, it might be able to improve the accuracy of Ca^2+^ trace estimation. To benchmark SEUDO, we simulated two-photon Ca^2+^ movies with varying levels of neuropil contamination, cell density, and Ca^2+^ trace signal-to-noise ratio (SNR). We processed these movies with EXTRACT and CNMF^18^, a MATLAB implementation of CAIMAN, to find cell’s spatial filters. Then, we used either SEUDO, robust regression, or NNLS to estimate the Ca^2+^ activity traces using the spatial filters from EXTRACT or CNMF. Notably, SEUDO improved the Ca^2+^ traces estimated by CNMF by removing some residual crosstalk, but the results fell short of robust regression (**Figs. 3G,H; S5**). SEUDO was also very slow and scaled poorly to movies with many cells **(Fig. S5D**). Further, when we attempted to use SEUDO to improve the estimated traces from EXTRACT, the results were inferior to those of either NNLS or EXTRACT (**Fig. 3G,H, S5**). Consequently, we concluded that the spatial filters from robust estimation led to more accurate Ca^2+^ trace estimation and that post-processing with SEUDO is not a suitable substitute.

Finally, we tested how EXTRACT would perform under challenging experimental conditions, such as those with slow imaging speeds, highly synchronized neural firing, and residual brain motion that had not been removed by image registration preprocessing (**Fig. S6**). We found that EXTRACT could resolve cells even when neural activity was nearly perfectly synchronized, providing that the imaging duration was sufficiently long to observe each cell fire individually at least once (**Fig. S6A,B)**. EXTRACT also tolerated downsampling in time to frame rates as slow as 2–3 Hz (**Fig. S6C**). Finally, EXTRACT could tolerate image displacements up to about the radius of an individual cell (**Fig S6D**). (Notwithstanding, we still recommend image registration prior to the application of EXTRACT).

The full list of our benchmarking studies can be found in **Table S1**, whereas additional experiments and methodological details are discussed in **Supplementary Notes 1, 3**, and **4**.

### A native implementation on (multiple) GPUs enables fast runtimes

EXTRACT’s main components are novel estimation algorithms that are customized for the challenge of cell extraction and that rely heavily on elementary matrix algebra. Thanks to several widely used software packages, such as the Intel Math Kernel Library, modern computers can perform matrix algebra operations in a highly optimized manner, which allows EXTRACT to achieve fast, scalable, and computationally efficient cell extraction. Our software implementation of EXTRACT also has native support for computation on graphical processing units (GPUs), enabling even greater efficiency for matrix operations.

When designing EXTRACT, we aimed to create a pipeline that can not only process the large neural recordings widely being acquired today with fluorescence mesoscopes^21^ but also scale well to even larger datasets, such as those with a million neurons^39^. To achieve this, we endowed EXTRACT with automated RAM and GPU memory management, such that mesoscope movies with large fields-of-view are spatially partitioned in way that both minimizes the loss of accuracy due to stitching effects and takes advantage of fast GPU matrix operations by allowing the partitions to contain thousands of cells (**Supplementary Note 2**). In contrast, although CAIMAN has been run in a parallelized way across 100 CPUs, the number of cells per partition is often 1-2 orders of magnitude smaller^20^. Overall, unlike past methods, EXTRACT is expressly designed to leverage the distinctive computational capabilities of GPUs while minimizing the processing time and errors that can arise from stitching together the results from many spatial partitions that slightly overlap.

To benchmark performance speed, we evaluated runtimes on datasets of varying sizes. For this purpose, we started with a Ca^2+^ video (4 mm^2^ field-of-view; 1024 × 1024 pixels; 30 Hz frame rate; 200,000 time bins; ∼800 GB) acquired from the neocortex of a live mouse on a laser-scanning two-photon mesoscope^1^ expressing the jGCaMP8s Ca^2+^ indicator in pyramidal neurons (**Fig. 4A**; **Methods**). To test extraction runtimes on raw datasets of different sizes, we used portions of this video, which covered subsets of the movie’s field-of-view or full duration.

**Figure 4.**
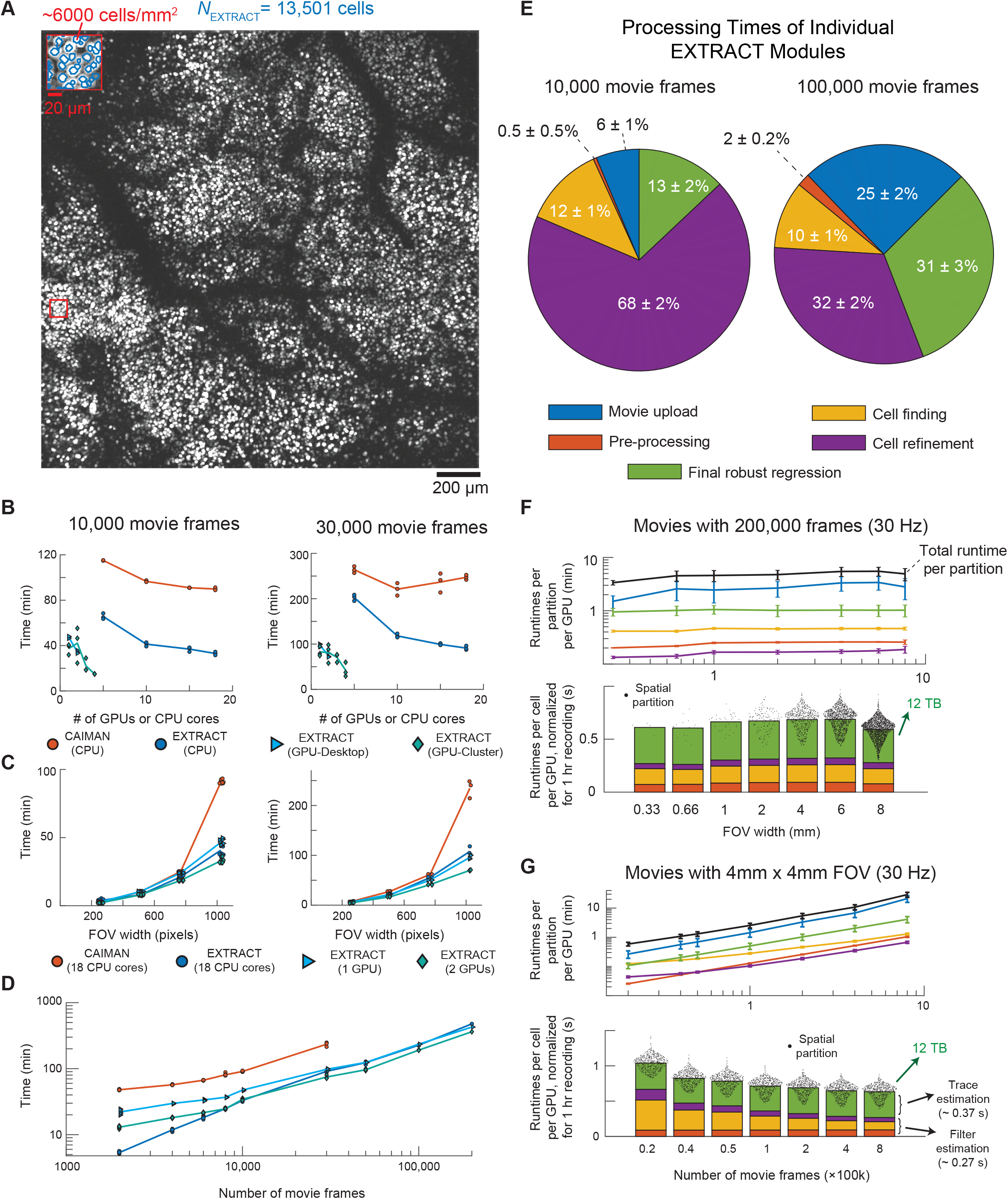
Fast and scalable processing of large-scale imaging datasets with EXTRACT. **(A–D)** To benchmark EXTRACT’s speed and scalability relative to the Python implementation of the CNMF algorithm (CAIMAN), we studied large-scale Ca^2+^ videos of densely labeled cortical pyramidal cells expressing soma-targeted jGCaMP8s that were acquired at 30 Hz using a laser-scanning two-photon mesoscope^1^ with a 2 mm × 2 mm field-of-view (FOV). **(A)** Maximum projection image of a Ca^2+^ video (∼2 hr duration; 200,000 frames) from the mesoscope. *Inset*: Magnified view of the tissue area boxed in red, showing a region with densely labeled neocortical pyramidal neurons (∼6000 cells per mm^2^). The blue solid lines mark the boundaries of cells found by EXTRACT. **(B, C)** *Left*, we first determined EXTRACT and CAIMAN runtimes on movies of short duration (10,000 frames), using variable numbers of GPUs and CPU computing cores with a fixed FOV (2 mm × 2 mm), **(B)**, or across a range of FOV sizes but with fixed computing resources (specified in the legend), **(C)**. Runtimes for both algorithms declined with greater use of distributed computing **(B)** and increased with larger FOV **(C)**. *Right*, we performed the same tests on movies with 30,000 frames. CAIMAN, but not EXTRACT, showed longer runtimes with more CPU cores, **(B)**, indicative of a deleterious phenomenon called memory thrashing. For both short and long movies, the speed differences between two algorithms became more apparent as the FOV got larger, **(C)**. **(C)** To evaluate runtime scaling on large movies, we processed movies with 2,000–200,000 frames. On a desktop computer with 256 GB RAM memory, CAIMAN failed to process movies bigger than 120 GB (∼30,000 frames). Notably, for movies with >10,000 frames, EXTRACT was faster when run on 2 GPUs than on 18 CPU cores. **(E)** To determine how the EXTRACT runtime reflects the times consumed by its internal substeps (**Fig. 2A**), we processed movies with a 2 mm × 2 mm FOV and either 10,000 or 100,000 frames using 1 GPU. The pie charts show the relative time consumed by each EXTRACT module. Notably, for movies with 100,000 frames, the movie upload times increased the most dramatically. Error bars: s.d. over 36 spatial partitions of the full 2 mm × 2 mm video. **(F, G)** To further test the scalability of EXTRACT, we created synthetic datasets by dividing or concatenating video data from the 2 mm × 2 mm, 200,000 frame video from **(A)**, in both space and time. We processed the resulting movies with EXTRACT using 2 GPUs and analyzed the times spent on each step of EXTRACT. In line with results in **(E)**, movie upload times dominated those of the other steps. **(F)** *Top*, as the FOV became larger, EXTRACT used more spatial partitions, and each partition consumed a similar runtime. *Bottom*, after uploading the movies, EXTRACT required ∼0.7 s per cell per 1 hr recording per GPU, *i*.*e*., 0.35 s with 2 GPUs. **(G)** *Top*, as the number of frames increased, EXTRACT took longer to process each partition. *Bottom*, with increasing number of frames, EXTRACT runtimes per cell per hour of recording per GPU decreased to ∼0.65 s, implying increased efficiency with longer movies. Black points in **(F, G)** show results from individual spatial partitions of the movie.

To evaluate runtimes for CAIMAN and EXTRACT (run on either CPUs or GPUs), we created two datasets with 10,000 (40 GB) and 30,000 (120 GB) frames, respectively. (As context, prior work^20^ on cell extraction termed movies with similar sizes ‘medium’ and ‘large’, although these sizes are relatively small given present experimental capabilities and far smaller than the movie of **Fig. 4A**). For these datasets, we determined the runtimes of EXTRACT and CAIMAN as a function of the numbers of CPUs and GPUs used (**Fig. 4B**) and of the spatial field-of-view (**Fig. 4C**). CAIMAN was slower than EXTRACT when both algorithms were run on CPUs, and, when EXTRACT was run on multiple GPUs, it was nearly an order-of-magnitude faster than CAIMAN.

Next, we checked runtime performance as a function of movie duration. For movies covering the full 4 mm^2^ field-of-view, EXTRACT, but not CAIMAN, was able to process the entire original movie of 200,000 time bins (800 GB) (**Fig. 4D**). By comparison, for movies of durations >30,000 frames, CAIMAN returned a memory error and was unable to process the data. To examine how the individual components of EXTRACT contributed to its overall runtime, we determined runtimes for each EXTRACT module, using 2 GPUs and movies with either 10,000 or 100,000 frames (**Fig. 4E**). Notably, the time needed to upload the ∼400 GB movie from a hard drive to the computer’s random access memory (RAM) took a much larger proportion of the total runtime than for the ∼40 GB movie. This led us to hypothesize that the scalability of EXTRACT to even larger, future datasets might be limited by upload and hard drive readout speeds, rather than algorithmic constraints.

To test this idea, we emulated futuristic, enormous datasets (∼45 TB), ∼50 times larger than present mesoscope videos, by concatenating more than one copy of the movie from **Fig 4A** in the spatial (**Fig. 4F**) or temporal (**Fig. 4G**) dimensions. Algorithmic runtimes scaled as expected, *i*.*e*., quadratically with the width of the field-of-view and linearly with movie duration (**Fig. 4F, G**). The calculated runtime per cell stayed constant as the field-of-view increased, and decreased as the movie duration grew, approaching ∼0.6–0.7 s per cell per GPU for a 1 hr recording. These results show that, unlike CAIMAN, EXTRACT can process movies that are far beyond the very largest Ca^2+^ videos presently available and is thus suited to handle the rapidly approaching deluge of neuroscience data.

Finally, to highlight that EXTRACT’s custom solver for robust regression plays a key role in its scalability to massive datasets, we did an experiment using a set of simulated Ca^2+^ videos with varying fields-of-view and numbers of cells (**Fig. S6E**). Across this set of videos, we determined the runtimes for EXTRACT’s custom solver (run as a non-negative least-squares estimator, *i*.*e*., in the limit κ → ∞) and compared these to the runtimes for MATLAB’s built-in non-negative least squares solvers (**Methods**). EXTRACT’s solver consistently outperformed and was nearly ∼100 times faster than MATLAB’s non-negative least squares solver, for movies of just 0.4 mm in width (**Fig. S6E**).

Thus, EXTRACT’s superior speed and scalability to large Ca^2+^ videos rely significantly on the speed of its internal, custom designed solver (**Supplementary Note 2**).

### Fast, comprehensive cell extraction from the Allen Brain Observatory data repository

After validating EXTRACT on both artificial and real data taken by two-photon imaging, we tested how well EXTRACT could process an entire repository of Ca^2+^ imaging data. To perform this test at a large scale, we applied EXTRACT to the publicly available Ca^2+^ imaging data repository from the Allen Institute Brain Observatory^36^ (**Fig. 5**). We downloaded 199 sessions of *in vivo* two-photon Ca^2+^ imaging data from GCaMP6-expressing cells across different visual cortical areas of behaving mice.

**Figure 5.**
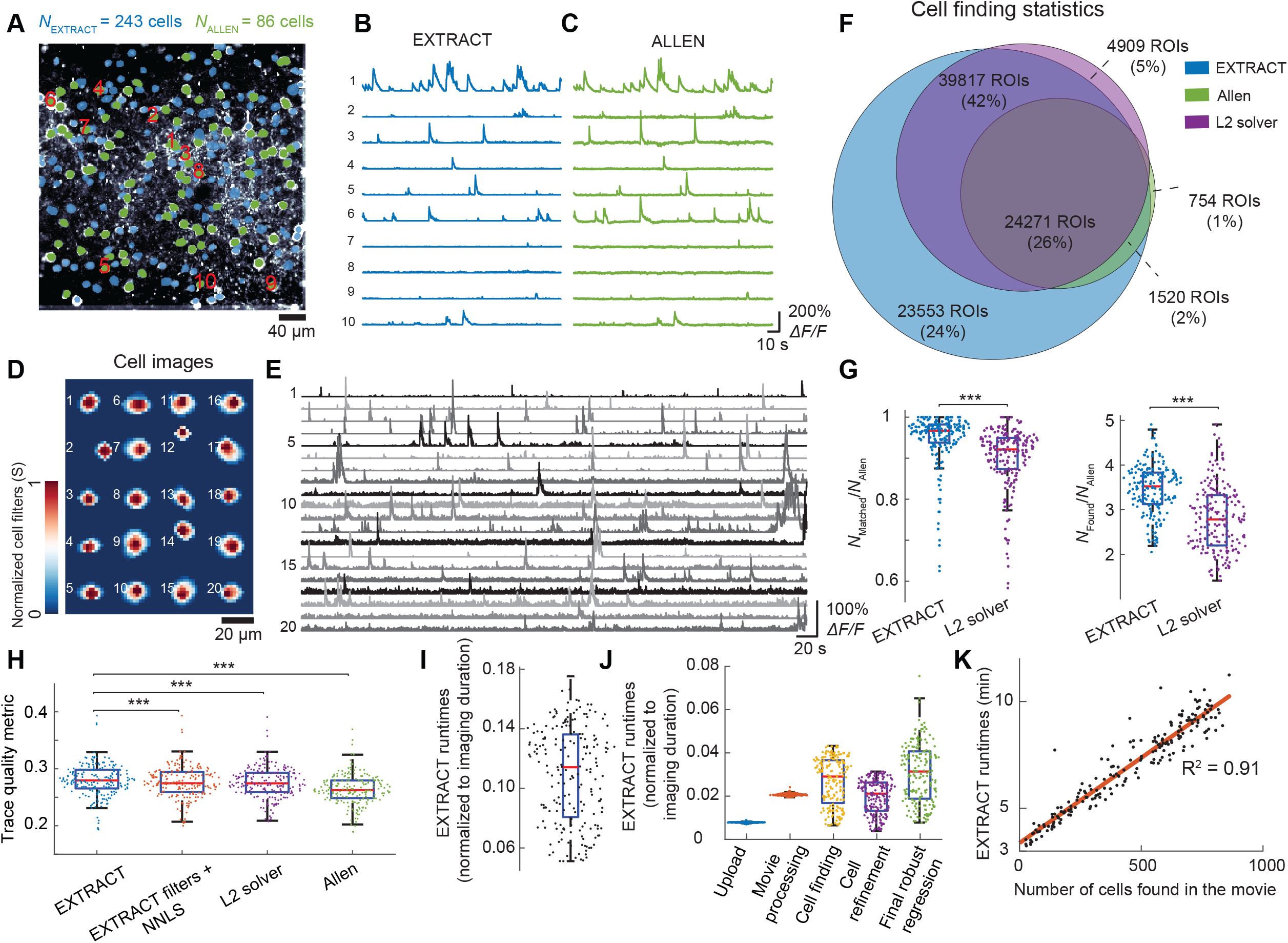
EXTRACT analyses of the Allen Brain Observatory Ca^2+^ imaging datasets. **(A–C)** Cell maps obtained by applying EXTRACT or the Allen Software Development Kit (SDK) to data from an example imaging session from the Allen Institute Brain Observatory. 86 cells colored green are those found by the SDK; EXTRACT also found all of these cells. Additional cells found by EXTRACT are colored blue; in total, EXTRACT identified 243 neurons. Estimated Ca^2+^ activity traces for the 10 neurons marked with numerals in **(A)** are shown in **(B, C)**. **(D, E)** Spatial images, **(D)**, and estimated Ca^2+^ activity traces, **(E)**, for 20 example cells that were identified by EXTRACT but not the Allen SDK. **(F–K)** To benchmark EXTRACT’s speed and cell extraction accuracy across diverse conditions, we ran EXTRACT on 199 movies from the Allen SDK in 23.1 hrs. To assess the unique contribution of the robust loss to the cell extraction quality, as in **Fig. 3**, we also processed the same datasets using EXTRACT but with the robust regression replaced with a conventional *L*_2_ loss function. **(F)** Venn diagram showing the net numbers of regions of interest output by EXTRACT, the Allen SDK, and the *L*_2_ solver determined over 199 imaging datasets. (See **Methods** for how cells were matched across the different output sets). **(G)** Box-and-whisker plots characterizing, for each of the 199 imaging sessions, the (*left*) fraction of cells found by the Allen SDK that were also found by EXTRACT or the *L*_2_ solver, and (*right*) the numbers of cells found by these two methods divided by the number found by the SDK. We used hyperparameters that were optimized for a single imaging session and then applied uniformly to the other 198 sessions. EXTRACT consistently surpassed the *L*_2_ solver, detected nearly all cells found by the Allen SDK, and identified ∼3.5 times the numbers of Ca^2+^ sources as the SDK. In panels **(G)** and **(H)**, *** denotes p < 10^−3^ using a two-sided Wilcoxon signed-rank test, **(H)** Box-and-whisker plots of the trace quality metrics (**Fig. 3**; **Methods**) for the estimated Ca^2+^ activity traces, averaged across all cells identified by the EXTRACT, the *L*_2_ solver, and the Allen SDK. Statistically, EXTRACT yielded higher quality traces than the Allen SDK, the *L*_2_ solver, and from the application of a non-negative least squares (NNLS) regression performed using the spatial filters from EXTRACT. Mean and s.e.m. values for EXTRACT: 0. 280 ± 0. 002, EXTRACT + NNLS: 0. 277 ± 0. 002, *L*_2_ solver: 0. 276 ± 0. 002, and Allen SDK: 0. 2640 ± 0. 002. **(I)** Box-and-whisker plot of the EXTRACT runtime (including the movie upload time) using a single GPU, divided by the movie duration across the 199 imaging sessions. Runtimes were typically ∼10 times faster than the duration of each Ca^2+^ movie. **(J)** Box-and-whisker plots showing the runtimes of each of EXTRACT’s modules relative to the Ca^2+^ movie duration for the 199 movies. **(K)** Scatter plot of EXTRACT runtimes using 1 GPU versus the number of cells found by EXTRACT. Each datum shows results from one of the 199 movies. Red line: Linear regression to the data (*t*_*runtime*_ = 3. 4 mins + 0. 008 * *N*_*cells*_ ; *R*^*2*^ = 0.91), implying that it took ∼11.4 mins to find 1000 cells, or ∼0.65 s per cell per movie hour per GPU, in accord with results of **Fig. 4**. Box-and-whisker plots in **(G–J)** show results from 199 movies; red lines denote median values; boxes span the 25th to 75th percentiles; whiskers extend to 1.5 times the interquartile range; data points denote results from individual movies.

The repository’s software development kit (SDK) has estimated spatial profiles for cells from each movie. The spatial profiles are regions-of-interest (ROI) estimates for each cell based on its morphology. Each cell’s Ca^2+^ trace comes from a linear regression of the Ca^2+^ movie onto the cell’s ROI, after subtracting an estimate of background Ca^2+^ activity in the neuropil. Owing to the lack of a non-negativity constraint, these traces will exhibit negative-going noise fluctuations and potentially lower SNR values than traces estimated when non-negativity is enforced. Therefore, to make even-handed comparisons with EXTRACT, we took cells’ spatial profiles from the Allen SDK and performed a non-negative least squares regression against the same pre-processed version of the Ca^2+^ movie as used in EXTRACT. With this approach, we benchmarked the Allen SDK and three other cell extraction methods using 199 movies from the repository with diverse attributes. The latter three methods were EXTRACT, a conventional *L*_2_ solver, and non-negative least squares regression applied using cells’ spatial profiles as obtained from EXTRACT.

Using a common set of hyperparameters, EXTRACT consistently identified nearly all cells found by the Allen SDK and many more that the SDK missed (**Fig. 5A–G**). For a typical movie, EXTRACT identified >97% of cells found by the SDK while also finding 3-4 times more cells (and some dendrites) than the Allen SDK (**Fig. 5G**). Visual curation of the cells identified by EXTRACT led to an estimated cell-finding precision of 96% (**Methods**). EXTRACT all found more cells than the *L*_2_ solver (**Fig. 5F,G**) and led to better trace quality than all the other methods (**Fig. 5H**). An in-depth study of 5 randomly chosen movies showed that hyperparameter optimization allowed EXTRACT to find all the cells found by the SDK.

Next, we quantified runtimes of EXTRACT on the 199 Allen movies. For movies that were ∼64 min long, EXTRACT had runtimes that were 0.11 ± 0.03 (s.d.) of each movie’s acquisition time (**Fig. 5I,J**). By comparison, CAIMAN reportedly had runtimes comparable to image acquisition times for movies of a similar scale, even when parallelized across 112 CPUs in a high-performance cluster (see Fig. 8 in Ref. ^20^). This observation suggests EXTRACT provides an order-of-magnitude or more speed improvement using only a single GPU on a standard desktop computer.

As the Allen SDK movies were all ∼1 hr in duration and covered an equivalent spatial area, the main factor leading to runtime differences was the number of cells in each movie. This led to a linear relationship (R^2^ = 0.91) between runtime and the number of cells in a movie, namely 3.4 min plus 8 min per thousand cells (**Fig. 5K**). This equals 0.64 s per cell per GPU for a 1 hr movie, consistent with the results of **Fig. 4** and confirming EXTRACT’s speed on a widely used database.

### Spatiotemporally clustered Ca^2+^ activity in striatal spiny projection neurons of active mice

As a first test of whether EXTRACT can yield superior biological results, we studied Ca^2+^ imaging data that we previously acquired in the dorsomedial striatum of freely behaving mice with a head-mounted, miniature epi-fluorescence microscope^40^. Each video is a recording of Ca^2+^ activity, as reported using the GCaMP6m Ca^2+^ indicator, in spiny projection neurons of either the direct or indirect pathway of the basal ganglia (dSPNs and iSPNs, respectively). We compared results from EXTRACT to those from PCA/ICA, the original MATLAB implementation of CNMF-E, and the Python version of the CNMF-E algorithm as implemented in CAIMAN^20,23^ (**Fig. 6**).

**Figure 6.**
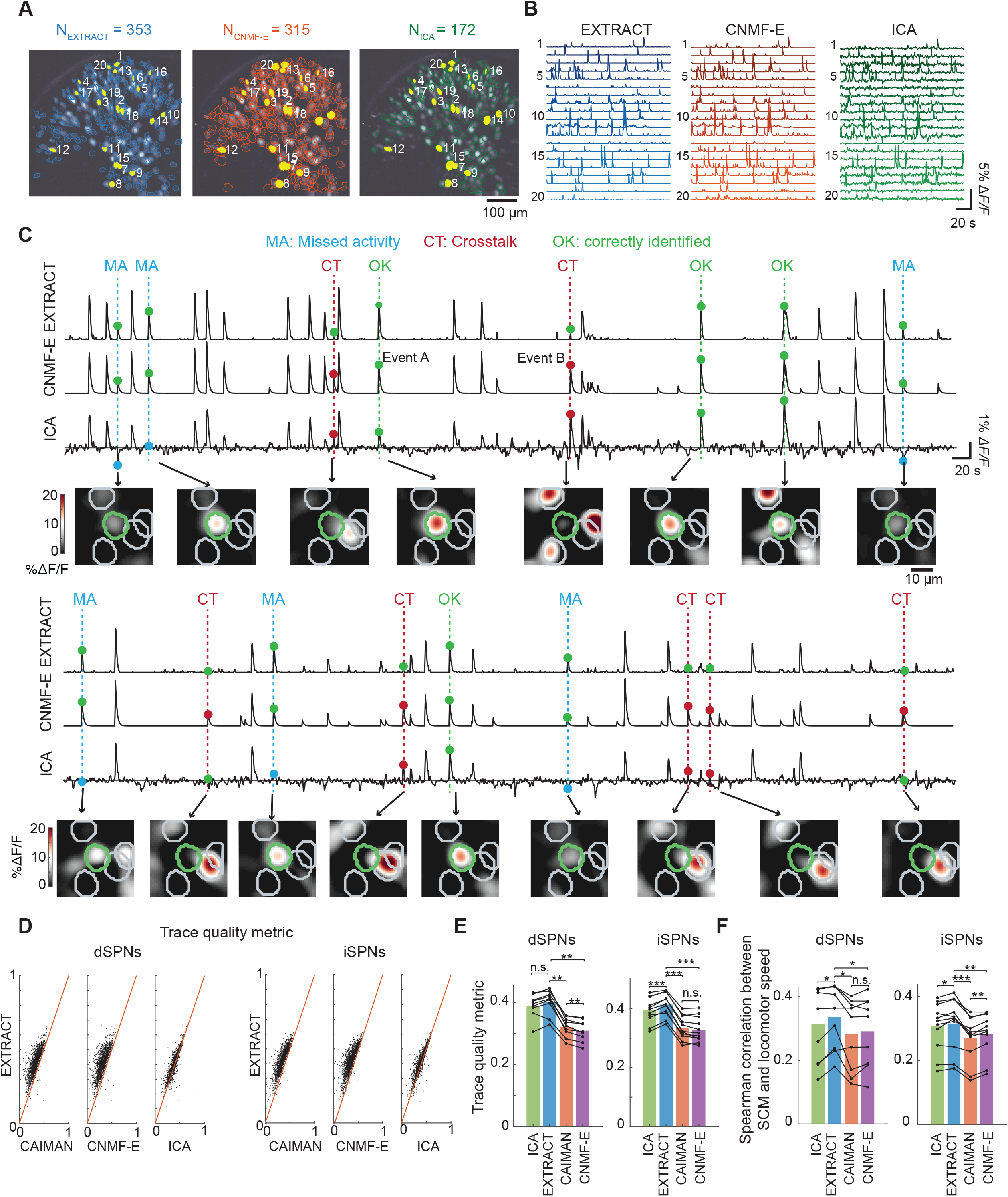
Evaluations of EXTRACT, CNMF-E, and ICA for the analysis of Ca^2+^ imaging data from direct and indirect pathway striatal spiny projection neurons (dSPNs and iSPNs). **(A)** Example maps of direct pathway spiny projection neurons (dSPNs) virally expressing GCaMP6m, as identified by EXTRACT, CNMF-E, and ICA from a representative, freely behaving mouse. 20 example cells found by each of the algorithms are marked with numerals. Data are from Ref.^40^. Note the greater size of the cells’ spatial footprints as estimated by CNMF-E, in accord with illustrations in **Fig. 1M**. **(B)** Estimated Ca^2+^ activity traces for the 20 example cells marked in panel **(A)**. **(C)** Magnified views of the Ca^2+^ activity traces for an individual example cell, a dSPN, as determined by the 3 different algorithms, allowing detailed qualitative assessments of the outputs at specific time points. Instances of missed Ca^2+^ activity (MA), crosstalk from a nearby cell (CT) and correctly identified Ca^2+^ transients (OK) are highlighted on the traces; alongside are image frames from the Ca^2+^ videos at the relevant time points, with the cell of interest shown in green and its immediate neighbors shown in gray. A spatial bandpass filter was applied to the images frames to enhance visualization (**Methods**). Colored dots mark mistakes in the estimated Ca^2+^ activity trace; green dots mark instances in which the Ca^2+^ activity trace is correct. Traces from ICA commonly exhibited missed Ca^2+^ transients. Traces from ICA and CNMF-E both had visible crosstalk from nearby cells, which led to incorrect estimation of Ca^2+^ event amplitudes and identification of false-positive events. (Compare Events A and B, for which CNMF and ICA wrongly estimated the Ca^2+^ event amplitudes). **(D, E)** To test the quality of dSPN and iSPN Ca^2+^ traces, we calculated trace quality metrics (**Fig. 3, Methods**) for cells matched across 4 algorithms (EXTRACT, ICA, CNMF-E, and CAIMAN). **(D)** Scatter plots show values of the trace quality metric for individual cells (black data points) for EXTRACT and the other 3 methods. EXTRACT consistently yielded higher quality traces across nearly all of the matched cells. **(E)** Plots (black lines and points) of the mean trace quality metric for individual mice, averaged over all cells extracted in each mouse, for the 4 different extraction algorithms. Colored bars show the mean values, averaged across mice. **(F)** As in Ref.^40^, for each time point of the Ca^2+^ videos we computed a spatial coordination metric (SCM) that characterized the extent to which neurons exhibited spatially clustered activity (**Methods**). Ca^2+^ activity traces from EXTRACT yielded significantly greater Spearman correlation coefficients between the SCM values and locomotor speeds. Data in **(D–F)** were from an identical set of 9 mice (for dSPNs) or or 11 mice (iSPNs). *p < 0.05, **p < 10^−2^, ***p < 10^−3^; n.s. = not significant; two-sided Wilcoxon signed-rank tests.

When we inspected the estimated Ca^2+^ activity traces, our observations fit well with those from simulation experiments (**Figs. 3, S2**). Notably, traces from PCA/ICA sometimes omitted Ca^2+^ transients that were plainly visible by simple inspection of the movie data (**Fig. 6C**, *cyan dots*). Further, activity traces from both PCA/ICA and CNMF-E exhibited crosstalk between neighboring cells (**Fig. 6C**, *red dots*). We quantitatively confirmed these observations by computing values of the trace quality metrics for all cells and mice (**Fig. 6D, E**). The MATLAB and CAIMAN implementations of CNMF-E yielded very similar but not identical results **(Fig. 6D–F)**, both of which tend to overestimate cells’ spatial footprints (**Fig. 6A**, *middle panel*), in line with simulation results using NNLS (**Fig. 1M**), the core algorithm in CAIMAN and CMNF-E.

We next examined whether differences arising in cell extraction quality might impact neurophysiological assessments. Our own prior study of striatal SPNs found that mouse locomotion led to activation of SPNs in a spatiotemporally clustered manner^40^. However, assessments of clustered activity are likely to be influenced by missing Ca^2+^ transients or crosstalk between spatially adjacent cells. For instance, crosstalk could elevate estimates of cells’ co-activation. Omitted Ca^2+^ transients might lead to underestimates of spatiotemporal clustering. To investigate, we used a spatial coordination metric (SCM), defined similarly to as in Ref. 40, to quantify the extent of spatially clustered activity in the striatum at each time frame (**Methods**). We compared results obtained by analyzing the activity traces from EXTRACT, PCA/ICA, and both versions of CNMF-E for a common set of cells. SCM values for the trace outputs of EXTRACT had significantly higher correlation coefficients with the mouse’s locomotor speed then the traces from other methods (**Fig. 6F**). We also verified that EXTRACT works well with two-photon Ca^2+^ videos of dSPN and iSPN activity (**Fig. S7**). Overall, our results show that superior cell extraction can lead to neurophysiological signatures that relate more precisely to animal behavior.

### EXTRACT detects dendrites and their Ca^2+^ activity

Some past cell extraction algorithms often do not provide sensible results when applied to Ca^2+^ videos of dendritic activity. Thus, we tested and validated EXTRACT on videos of dendritic Ca^2+^ activity in cerebellar Purkinje cells and neocortical pyramidal neurons in live mice (**Fig. S8**). Although the default mode of EXTRACT discards candidate cells whose spatial areas or eccentricities are uncharacteristic of cell bodies (**Fig. S9**), the user can opt to retain candidate sources of Ca^2+^ activity without regard for their morphologies, thereby allowing EXTRACT to identify active dendrites. For example, in large-scale movies of Purkinje neuron dendritic Ca^2+^ spiking activity acquired with a two-photon mesoscope^1^, EXTRACT identified the dendritic trees of >500 cells per mouse, and the extracted spatial forms had the anisotropic shapes that are characteristic of these cells’ dendritic trees, which are highly elongated in the rostral-caudal dimension^16^ (**Fig. S8A, B**). We also used EXTRACT to analyze Ca^2+^ videos acquired by conventional two-photon microscopy in apical dendrites of layer 2/3 or layer 5 cortical pyramidal cells in live mice (**Fig. S8C, D**). EXTRACT identified ∼850–900 dendritic segments per mouse, and, as expected, they had a wide variety of shapes and temporally sparse Ca^2+^ transients. For both cerebellar and neocortical neurons, we found no limitations to the dendrite shapes that EXTRACT could identify, and it readily identified large numbers of dendritic segments without making assumptions about dendrite morphology or spatial continuity on the imaging plane as discussed elsewhere^13^.

### EXTRACT improves identification of anxiety-encoding cells in ventral hippocampus

As another test of whether EXTRACT can improve biological findings, we examined the Ca^2+^ activity of pyramidal neurons in the CA area of the ventral hippocampus (**Fig. 7A**). We tracked the dynamics of these cells in freely behaving mice navigating a 4-arm elevated plus maze (EPM, **Fig. 7B**). The EPM had 2 enclosed and 2 open arms, arranged conventionally on the perpendicular linear paths of the maze. The EPM assay is based on rodents’ innate aversion to open, brightly lit spaces and is widely used to investigate anxiety-related behavior^41^. A subset of ventral CA1 neurons, termed ‘anxiety cells’, show enhanced activity when the mouse is in anxiogenic regions of the EPM, namely the open arms^42–44^. We used EXTRACT and CNMF-E to obtain Ca^2+^ activity traces of ventral CA1 cells and compared their encoding of the open and closed arms (**Fig. 7C**).

**Figure 7.**
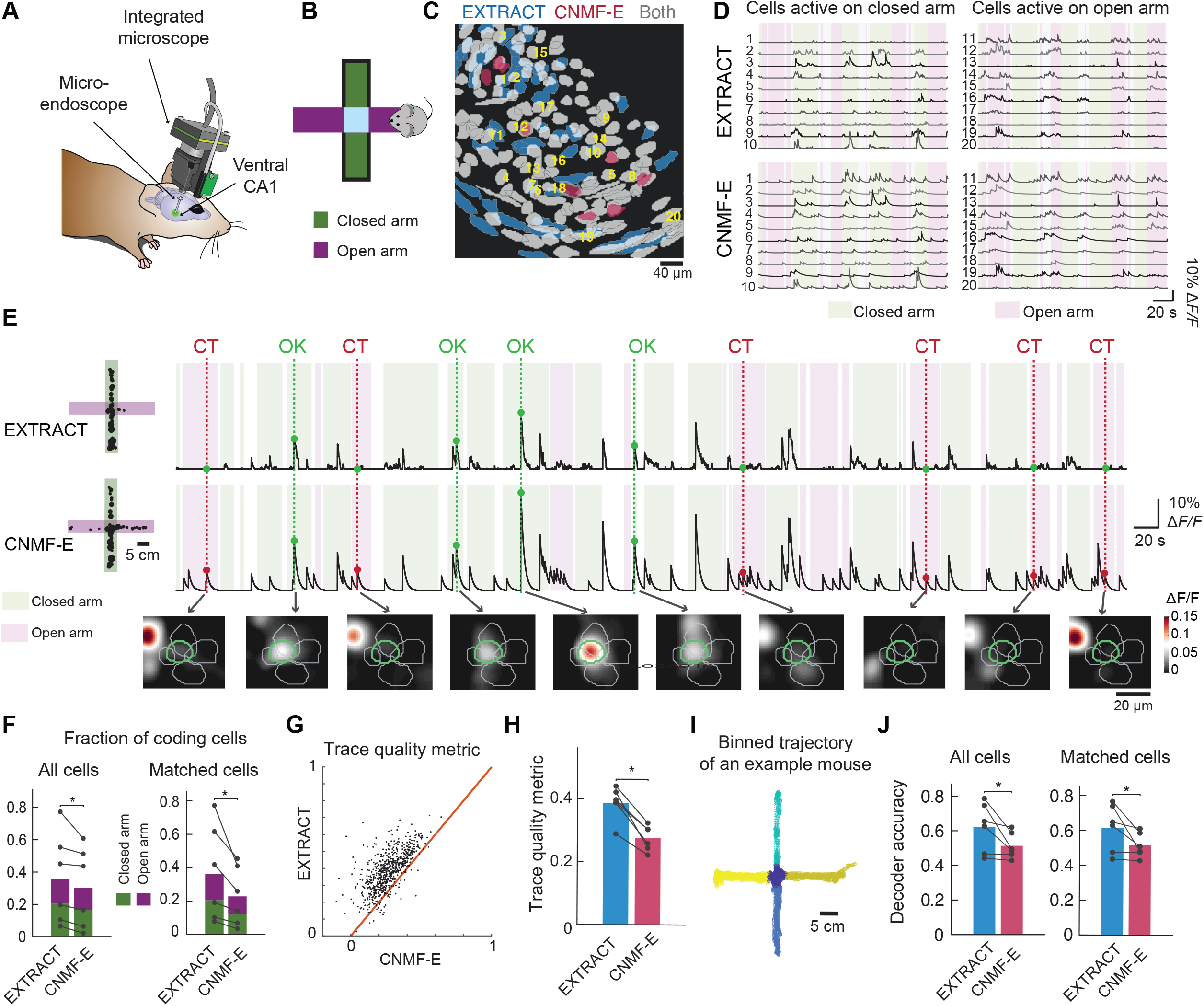
EXTRACT enables superior identification of anxiety-coding cells in ventral CA1. **(A)** We used the integrated, miniature fluorescence microscope and an implanted microendoscope to image the somatic Ca^2+^ activity of pyramidal neurons expressing the Ca^2+^ indicator GCaMP6s in the ventral portion of the CA1 hippocampal subfield in freely behaving mice. **(B)** Mice navigated an elevated plus maze (EPM) consisting of two open arms and two arms enclosed with walls. **(C, D)** We analyzed the Ca^2+^ videos using EXTRACT or CNMF-E, a variant of CNMF that is better suited for one-photon Ca^2+^ imaging. Panel **(C)** shows a map of pyramidal cells found by one or both of the two algorithms. A majority of detected cells were found by both algorithms, although EXTRACT identified a greater number of cells. 20 example cells found by both methods are marked with yellow numerals; panel **(D)** shows Ca^2+^ activity traces for these cells, as estimated by EXTRACT and CNMF-E. 10 of the neurons (*left*) were preferentially active when the mouse was in one of the closed arms (periods marked in light green). The other 10 cells (*right*) were preferentially active when the mouse was in one of the open arms (periods marked in pink). **(E)** Many pyramidal cells in ventral CA1 were preferentially active when the mouse was in either the closed or open arms of the maze, as illustrated across this panel for an example cell that was more active when the mouse was in a closed arm. *Left*, Maps of the EPM showing the mouse’s locations (black dots) at which the cell exhibited a Ca^2+^ transient, as detected using EXTRACT and CNMF-E. The area of each dot is proportional to the peak magnitude of the corresponding Ca^2+^ transient. *Right*, The cell’s traces of Ca^2+^ activity, as determined by the two extraction algorithms. The abbreviation ‘OK’ above the traces marks Ca^2+^ transients (green dots) that were correctly found by both methods. ‘CT’ marks instances of crosstalk in the trace from CNMF-(pink dots). The pink and light green shading respectively indicate periods when the mouse was in the open and closed arms of the maze. Images below the traces are from individual image frames and show the activity of the example cell (outlined in green) and its immediate neighbors (outlined in gray). Note the false transients reported by CNMF-E when the mouse is in the open arm, yielding the incorrect impression that the cell is active on both the closed and open arms. **(F)** We identified cells that encoded the arm-type of the EPM using the Ca^2+^ traces output by each algorithm for the recordings from 6 different imaging sessions (*N =* 3 mice) (**Methods**). EXTRACT yielded a greater proportion of cells that encoded the arm-type than CNMF-E, across all extracted cells and for cells found by both methods (Wilcoxon signed-rank test; \**p* < 0.05). Black data points and lines denote data from individual imaging sessions. **(G, H)** Trace quality metrics (**Fig. 3, Methods**) computed for cells matched across the outputs of EXTRACT and CNMF-E from 6 different imaging sessions. **(G)** Scatter plot of the trace quality metric for individual cells (black data points) for EXTRACT and CNMF-E. EXTRACT consistently yielded higher quality traces for nearly all cells. **(H)** Values of the mean trace quality metric for individual mice (black lines and points), averaged over all cells found in each mouse, for the 2 different algorithms. Colored bars show the mean values, averaged across 6 imaging sessions. **(I, J)** We divided the EPM into 5 spatial bins. We used support vector machine classifiers to predict the mouse’s spatial bin based on the Ca^2+^ events detected in traces from either EXTRACT or CNMF-E. Panel **I** shows the locomotor trajectory of an example mouse, color-coded so each of the 5 bins is shown in a distinct color. Panel **(J)** shows mean decoder accuracies (colored bars), averaged over 6 imaging sessions with 3 different mice and 100 random splits of the data into decoder training and testing portions. EXTRACT led to superior classification of mouse location. Black data points and lines denote results from individual imaging sessions. In **(F), (H), (J)**, * indicates *p*<0.05, two-sided Wilcoxon signed-rank test, n *=* 6 imaging sessions.

In Ca^2+^ traces from both EXTRACT and CNMF-E, a subset of cells responded differentially when the mouse was in the open versus the closed arms (**Fig. 7D**). Namely, distinct subsets of ventral CA1 cells were active when the mouse occupied the two different arm-types, in accord with past reports of their anxiety-related coding^42,43^. However, Ca^2+^ traces from EXTRACT generally exhibited a purer form of coding, in that the traces were typically silent when the mouse was in one arm-type but had high activity levels in the other arm-type. Traces from CNMF-E tended not to distinguish the two arm-types as clearly (**Fig. 7E**). Traces from EXTRACT also corresponded more precisely to Ca^2+^ activation events that were plainly apparent in the movies (**Fig. 7E**, *lower panel*).

To quantify these observations, we compared the arm-coding cells identified using the traces from the two different extraction algorithms. Notably, EXTRACT yielded significantly more arm-coding cells than CNMF-E, even when the analysis was restricted to cells identified by both algorithms (**Fig. 7F;** Wilcoxon signed-rank test, p < 0.05). To assess how well the activity traces from the two algorithms reflected events in the Ca^2+^ video data, we computed mean values of the trace quality metric for all cells and mice. Activity traces from EXTRACT had superior trace quality metric values than those from CNMF-E (**Fig. 7G,H**; Wilcoxon signed-rank test, p<10^−5^ for n=665 matched cells, p = 0.03 for n=6 mice), showing that EXTRACT more accurately captured the Ca^2+^ dynamics present in the movie data.

Finally, we evaluated how well the sets of activity traces from the two algorithms allowed one to estimate the mouse’s behavior using decoders of neural ensemble activity. We divided the EPM into 5 spatial bins (**Fig. 7I**) and trained support vector machine (SVM) classifiers to predict the spatial bin occupied by the mouse, based on the neural ensemble activity pattern at each time step (**Methods**). We compared the decoder accuracies using a distinct subset of the data than that used for decoder training. Strikingly, for every mouse, decoders based on traces from EXTRACT outperformed the decoders based on traces from CNMF-E (for both common and all cells, **Fig. 7J**).

## DISCUSSION

Here we have introduced the first major data analytic pipeline in systems neuroscience that is based on the mathematical framework of robust statistics^45^. Our contributions to this framework include: the one-sided Huber loss function (**Fig. 1F**), which we expressly developed to handle the rectified nature of Ca^2+^ activity; adaptive estimation of kappa, the parameter describing the levels and spatiotemporal variations of non-Gaussian noise contaminants in the Ca^2+^ movie (**Fig. 1F, G**); an implementation of multivariate robust regression using ADMM (**Supplementary Note 2**); a geometric interpretation of robust regression as applied to the estimation of neural activity traces and spatial profiles (**Fig. 1I–L**); and proofs providing guaranteed upper bounds on worst-case estimation errors (**Methods**). Based on these elements, we created an open-source computational pipeline, EXTRACT, in which cell finding, cell refinement, and the estimation of Ca^2+^ activity traces are all implemented using multivariate robust regression with a one-sided Huber loss (**Fig. 2**).

Although it is commonplace to use the word ‘robust’ to describe results or analyses that are resilient to the presence of noise or perturbations, such colloquial usage does not refer to the formal theory of robust statistics^34,46^ that was first developed in the 1950s. Prior work on cell extraction algorithms in neuroscience used the term ‘robust’ colloquially^26,29^, but, traditionally, robust estimators should exhibit certain formal properties^45^, including being able to handle a large percentage of outlier data points, providing correct results in the absence of outliers, generalizing well to diverse datasets, resisting the influence of extreme outliers, and computational and operational simplicity. EXTRACT exhibits all of these properties.

### EXTRACT is a versatile method for analyzing a broad range of Ca^2+^ imaging datasets

EXTRACT provides a superior means of analyzing somatic or dendritic Ca^2+^ data acquired with conventional, multi-plane or large-scale two-photon microscopes, or with head-mounted epi-fluorescence microscopes (**Figs. 4–7; S7, S8**). This broad applicability stems from one major factor, namely that the theoretical framework on which EXTRACT is based makes minimal assumptions about the nature of the data.

The estimation framework on which EXTRACT is based does not model noise sources; instead it aims to isolate cellular Ca^2+^ signals from contamination sources while staying agnostic to the latter’s exact form. This approach leads to formal robustness and great flexibility. Here, the minimization of a robust loss function, the new one-sided Huber loss, together with adaptive estimation of ***κ***, the parameter describing the extent of non-Gaussian noise contaminants, enables automated detection of outlier data points in a way that is adaptively updated from the data itself. This allows EXTRACT to handle a high-percentage of outlier data points. However, when the noise contaminants approximate statistically independent, Gaussian-distributed noise at each image pixel, the loss function used in EXTRACT adapts itself to behave like a linear regression loss and thereby achieves the optimal statistical efficiency of a standard maximum likelihood estimator^37^. In an opposite extreme case, when the data suffer from large contaminants due to Ca^2+^ activity in overlapping cells or neuropil, the EXTRACT loss function modifies its robustness parameter so as to reject these contaminants. Further, EXTRACT makes no assumptions about cell morphology and/or the temporal waveforms of Ca^2+^ activity, which allows EXTRACT to generalize well and to detect activity in somata or dendrites of many different neuron classes. Its fast and scalable solver is geometrically interpretable (**Fig. 1H–L**) and provides computational simplicity. Thus, EXTRACT satisfies the formal hallmarks of robust statistical analyses.

Several prior methods for cell extraction took a different approach and used explicit models of neural attributes to separate cellular Ca^2+^ activity from strong background contaminants. For instance, CNMF-E seeks to infer neural Ca^2+^ activity while modeling background activity as a linear combination of the residual activity within nearby pixels^23^. Minian is also based on the CNMF method and, like CNMF-E, is mainly intended for analyses of one-photon Ca^2+^ imaging datasets^24^. It applies several image processing steps to the movie data, carefully initializes cell locations, and then applies the CNMF method. Other authors have applied *post hoc* denoising of Ca^2+^ activity traces, by taking a set of previously identified neurons and re-estimating the Ca^2+^ activity traces in a way that seeks to minimize crosstalk and contamination^26–28^. Common to all these prior approaches are efforts to either model the noise sources or to remove them, based on certain assumptions about the data. Unfortunately, as we found throughout our benchmark studies using simulated data, explicit modeling of the background noise distribution fails—even when the model assumptions are perfectly met (**Figs. 3A–D**; **S2, S3**), *i*.*e*., with Gaussian noise, exponentially decaying Ca^2+^ signals, and Gaussian-shaped cell filters. This failure is due to (a) the non-Gaussian contaminants introduced by overlapping cells, and (b) the spatially correlated, non-homogenous levels of Gaussian noise that stem from cells’ nonbinary, continuously valued spatial profiles.

By comparison, EXTRACT makes few assumptions about the data and little use of image processing. Thus, while our robust estimation framework has not been fine-tuned to work optimally under specific statistical conditions, it is designed to yield high fidelity results across a wide spectrum of data statistics, as is expected from a robust estimator. Moreover, traditional robust estimation approaches, including our prior work^35^, lacked adaptive ***κ*** estimation and do not automatically adapt to different levels of non-Gaussian contamination. This capability greatly enhances EXTRACT’s ability to achieve excellent analytic performance on datasets from a variety of brain areas and imaging modalities (**Fig. 1G**).

It is worth noting that EXTRACT’s robustness and its ability to generalize to multiple imaging modalities stands in direct contrast to recent deep network-based approaches for cell extraction^30^, which have very slow runtimes and seem to require network re-training for new datasets, neuron-types or imaging modalities. Still lacking are comprehensive studies that prove the ability of deep network approaches to provide trustworthy cell extraction results for datasets with statistics that differ from those of the data used to train the network. It will be crucial to perform careful studies of the hallucinations that will inevitably arise and infect the analytic results of deep network-based cell extraction^32,33^. Rigorous studies of this kind will also need to evaluate Ca^2+^ activity traces with metrics that are more sophisticated than correlation coefficients between estimated and ground truth fluorescence activity traces, which can be highly dominated by periods of cellular inactivity, do not isolate hallucinated or crosstalk activity, and overly reflect the level of concordance between the estimated and actual decay time-constant of Ca^2+^ transients. While the future of cell extraction may eventually lie in this direction, current deep network approaches to cell extraction remain early-stage. A potentially attractive future possibility is that perhaps deep network cell extraction methods might be trained on EXTRACT’s robust traces.

Notwithstanding, neuroscientists should recognize that robust estimators do have certain limitations. Under conditions with very low optical SNR, the estimator trades robustness for fidelity, causing it to behave more like an *L*_2_ estimator (**Methods**). Although EXTRACT applies spatial filtering during pre-processing and cell finding steps to enhance the input SNR, for movies with extremely low SNR, robust estimators do not boost performance of trace estimation much beyond that of least-squares. This stems from the fact that, in a low SNR movie, fluorescence signals in individual pixels have large variances that typically are larger than the non-Gaussian contaminants and thereby dominate the errors in the estimated Ca^2+^ traces. Nonetheless, the outputs from EXTRACT should still be sensible due to its model-agnostic nature; in the low SNR scenario, ***κ*** adaptively takes on a large value that allows the regression to approximate least-squares regression, selectively using pixels with the highest SNR for filter estimation (**Fig. 1K–M**).

### An efficient implementation for fast cell extraction that scales well to large datasets

Owing to recent advances in optical technologies, such as fluorescence mesoscopes and multi-arm microscopes that can monitor multiple brain areas concurrently, Ca^2+^ imaging data is now routinely collected at a scale of several terabytes per publication^1,2,36,40,47^. Notably, time-lapse studies with multiple imaging sessions for each animal can readily produce datasets of this magnitude ^40,47,48^. Such datasets are so large that the raw data from a single original research study are typically not shared on the most commonly used public data repositories. Aside from issues of data sharing, the sheer volume of leading-edge datasets necessitates faster processing algorithms to avoid a major bottleneck in the pace of systems neuroscience research.

To handle the most massive datasets, we developed EXTRACT and showed that it can process typical Ca^2+^ movies in times that are as much as ∼10-fold briefer than the movie durations (**Fig. 5I-K**). EXTRACT’s built-in support for multiple GPUs substantially accelerates processing, allowing cell extraction from a 1 hr movie in ∼0.6–0.7 s per cell per GPU. For a Ca^2+^ movie with a thousand cells, analyzed with 2 GPUs, this translates to ∼6 min of processing, or ∼10% of the movie duration. This represents an order-of-magnitude speedup over past methods (**Fig. 4**).

On large datasets, EXTRACT performed quickly in all regimes, and runtimes scaled gracefully as dataset sizes (**Fig. 4**). On the Allen Institute Brain Observatory data, EXTRACT ran in ∼11% of the time of a typical recording session; this enabled batch processing of ∼212 hours of recordings in less than a day (**Fig. 5I–K**). On two-photon mesoscope recordings, EXTRACT ran much faster than the widely used CAIMAN method and successfully analyzed datasets ∼10 times bigger than the maximum sizes that CAIMAN could handle (**Fig. 4B–D**). With these recordings, EXTRACT was not limited by RAM and/or GPU memory, did not require a high-performance cluster, and needed only a typical desktop PC to handle futuristic, 12 TB datasets (**Fig. 4F, G**).

The accelerated computation from EXTRACT’s use of GPUs does not require any special handling, such as explicit parallelization or algorithmic variations. EXTRACT runs the same code on CPUs and GPUs, if the latter are available to the user. With any suitable MATLAB compatible GPU installed on the analysis computer, one can readily use EXTRACT with GPU processing to achieve major speed-ups over the CPU runtime. GPUs typically cost a fraction of the analysis computer, and nowadays most pre-configured computers include GPUs that have computing capability. In addition to faster runtimes, EXTRACT’s built-in GPU support implies that, since its computationally intensive tasks are run on the GPU, the user can run other CPU-demanding software at the same time. We have tested EXTRACT on a variety of computing platforms, including laptops, desktops, and high-performance clusters (**Methods**), which gives researchers the freedom to choose their computing resources based on their experimental needs, unconstrained by EXTRACT software. Further, EXTRACT does not require pre-installation of other computing packages than MATLAB and thus is readily compatible with high-performance computing platforms.

### EXTRACT enables improved scientific results

The identification of neurons from movie data is a crucial step in neuroscience experiments that rely on Ca^2+^ imaging techniques for large-scale recording of neural dynamics. Extraction of individual cells and their activity traces reduces the raw data to a set of time series, the accuracy of which is crucial for the success of all subsequent analyses. Thus, EXTRACT aims to achieve high-fidelity results by avoiding extraneous image processing as much as possible (and the existing preprocessing steps are fully interpretable in terms of their effects to the movie, **Supplementary Note 1**) while also improving the inference of cellular activity and removing non-Gaussian contaminants through robust statistical estimation. Unlike some past approaches to cell detection, we found that EXTRACT works well with Ca^2+^ videos of dendritic activity, which often do not provide as many fluorescence photons as videos of somatic activity. Further, our results from two separate biological experiments highlight that the use of EXTRACT can lead to improved scientific results (**Figs. 6**,**7**).

First, we evaluated EXTRACT, CNMF-E (Python and MATLAB implementations), and PCA/ICA using Ca^2+^ imaging data taken from striatal spiny projection neurons (SPNs) (**Fig. 6**), which exhibit spatially clustered activity patterns during animal locomotion^40^. When we analyzed these activity patterns, EXTRACT provided higher quality traces (**Fig 6D,E**) that led to higher correlations between the spatially clustered activity and the animal’s locomotor speed (**Fig 6F**). This fits with our observations that EXTRACT made the fewest mistakes during the cell extraction process, as seen by comparing the traces from all 3 algorithms to the raw data (**Fig. 6C**).

Second, we characterized anxiety-related representations in the ventral hippocampus of mice behaving within an elevated plus-maze for studies of anxiety (**Fig. 7**). Using the neuronal Ca^2+^ traces from EXTRACT, we identified significantly more cells with anxiety-related coding than when we used the outputs of CNMF-E (**Fig. 7E,F**). The use of EXTRACT also led to superior decoding analyses (**Fig. 7I–J**), in that the traces from EXTRACT enabled better estimates than CNMF-E of the animals’ locomotor trajectories (**Fig. 7J**). These results confirm that accurate biological findings require accurate reconstructions of neuronal activity and show that EXTRACT improves the results from downstream computational analyses, especially when the raw data may have substantial noise or fluorescence contaminants.

### OUTLOOK

Ca^2+^ imaging technology continues to progress rapidly, with new tools emerging for multi-color Ca^2+^ imaging of multiple cell-types and volumetric Ca^2+^ imaging^21,39^. Techniques for high-speed optical voltage imaging are also making rapid strides and provide direct access to neural membrane voltage dynamics. Because EXTRACT makes so few assumptions about data statistics, future versions of the algorithm should be applicable to the data from these emerging imaging modalities with only straightforward modifications^49^.

To increase the numbers of neurons that can be tracked simultaneously, new imaging approaches are arising in which cells from multiple planes in tissue are deliberately superposed in the raw video data^50–53^; the cells and their activity traces must then be disentangled in offline analyses. EXTRACT’s capability for high-fidelity isolation of individual cells, even when cells greatly overlap one another in the raw images (**Fig. 3C–F**), should facilitate multi-plane imaging by allowing a greater number of planes to be sampled concurrently while still being able to computationally extract the individual neurons from dense sets of overlapping cells.

Another ongoing set of advancements concerns interventional experiments in which neural activity or animal behavior is manipulated in real-time, based on recorded patterns of neural activity. Irrespective of whether such online interventions are implemented in a closed- or open-loop form, they generally require specialized versions^17,54^ of cell extraction algorithms that have data processing delays corresponding to only one or only a few image frames in duration, but typically at the cost of notably diminished estimation accuracy. Robust regression-based cell extraction, such as with a future real-time version of EXTRACT, seems especially well suited for such applications, as it does not need acausal information from the movie to estimate Ca^2+^ traces, unlike methods that apply a decaying exponential kernel to Ca^2+^ traces^17^.

Beyond cell extraction from Ca^2+^ movies, we expect that the general framework of robust regression will have broad applications across neuroscience for analyses of many types of recording data, both optical and electrophysiological. Moreover, the scalability of EXTRACT (and algorithms inspired by it) to datasets approaching the petabyte scale will likely be a crucial feature of future analytic pipelines.

## ACKNOWLEDGEMENTS

We gratefully acknowledge research support from HHMI (M.J.S.), the Stanford *CNC* Program (M.J.S.), DARPA (M.J.S.), the NIH BRAIN Initiative (M.J.S.), an NSF NeuroNex grant (M.J.S.) and the Stanford Mind Brain Computation & Technology Program (F.D.). We thank R. Chrapkiewicz, A. Christensen, M.S. Ebrahimi, H. Kim, A. Shai, M. White, S. Boyd, P. Nobel for helpful conversations, C. Gillon and J. Zylberberg for videos of dendritic Ca^2+^ activity, and O. Akengin for manual annotation of the Allen SDK datasets. Some of the computing for this project was performed on the Sherlock cluster. We would like to thank Stanford University and the Stanford Research Computing Center for providing computational resources and support that contributed to these research results.

## METHODS

### MICE

All procedures were approved by the Stanford University Administrative Panel on Laboratory Animal Care (APLAC). Ca^2+^ imaging studies in ventral hippocampus used male double-transgenic CaMKII-GCaMP6s mice (tetO-GCaMP6s-2Niell/J: Camk2a-tTA-1Mmay/DboJ, Jackson Laboratory, stock #007004 and #024742 respectively) aged 12-16 weeks at the start of experimentation^55^.

For Ca^2+^ imaging studies of cerebellar Purkinje cell dendritic trees, we used mice that were a cross of PCP2-Cre driver mice with a Bl6-129 genetic background and Ai148 transgenic mice^56^; the resulting double transgenic mice (*PCP2-cre*/*TIGRE-loxP-stop-loxP-CAG-tTA2-TRE-GCaMP6f [Ai148]*) expressed the GCaMP6f Ca^2+^ indicator selectively in Purkinje cells. We discuss the mouse preparation for each experiment below in the corresponding section.

### NOTATION FOR MATHEMATICAL VARIABLES

Throughout the methods section, we denote the size of the imaging field-of-view as *h* × *w*, in units of pixels. We refer to the scalar product *hw* as *n*_*pixels*_. We use boldface characters for arrays and non-boldface characters for scalars. As in the main text, we denote the movie matrix as ***M*** (flattened in space, so that ***M*** is a two-dimensional matrix), the matrix of spatial weights (cell images) as ***S***, and the matrix of temporal weights (Ca^2+^ traces) as ***T***.

### THEORY BEHIND EXTRACT’S ROBUST SOLVER

Starting with the model described in the main text (**Figure 1E**), the common goal of cell extraction routines is to reconstruct an estimated movie, 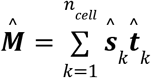, where *n*_*cell*_ corresponds to the number of estimated cells,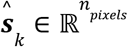 and 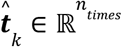 denote the estimated spatial profile and Ca^2+^ activity trace for the *k*’th cell, *n*_*pixels*_ is the number of pixels in the movie, and *n*_*times*_ is the number of movie frames. This problem differs from the toy example in **Figure 1A–D** in two key details. First, both the cells’ spatial profiles and Ca^2+^ activity traces need to be jointly estimated. Second, the number of cells in the movie is unknown and should be estimated from the data.

To solve this estimation problem, most existing approaches^19,20^ (**Supplementary Note 1**) aim to minimize the *L*_2_ loss on reconstruction of the movie,

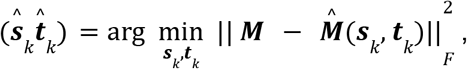

where ∥. ∥_*F*_ denotes the Frobenius norm. To detect cells that are spatially localized and have connected spatial profiles, the cell extraction process often starts with an initialization stage^57^, termed ‘cell finding’. This is followed by refinement stages, in which one of 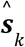 or 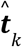 is held fixed and the other is updated by minimizing the movie reconstruction loss^19,20^, a subproblem that constitutes a constrained linear regression.

Unlike traditional regression methods, EXTRACT uses the one-sided robust Huber loss function, which defines a robust version of the regression problem (**Fig. 1F, H–L**). In this section, we introduce and motivate the robust regression framework (**Algorithm 1** below), which is used by EXTRACT as a subroutine to update its estimates of cells’ spatial profiles and Ca^2+^ activity traces.

#### Theory of robust estimation in the presence of large non-negative contaminants

Here, we introduce our signal estimation approach, based on the theory of robust M-estimation. This theory is well-developed for symmetric and certain asymmetric contamination regimes^34,35,45,46^. However, prior theoretical work does not readily suggest an optimal estimator that is suitable for estimating Ca^2+^ signals, which are largely rectified (*i*.*e*., positive-going), under conditions with spatiotemporally varying levels of noise contamination that is also positively rectified. Thus, extending on the theory of robust M-estimation developed in our prior work^35^, which uses a one-sided Huber loss function suited to conditions of asymmetric contamination, we first introduce a simple mathematical abstraction for treating contaminants that vary across time bins and cells. Then, we propose a fast solver that minimizes the robust loss using matrix multiplications that are amenable to parallel processing, *e*.*g*., over multiple GPUs. This solver adaptively estimates cells’ baseline levels of Ca^2+^ activity from the movie data and also adaptively updates the estimated level of non-Gaussian contamination. Finally, we provide a convergence proof for the generalized solver and discuss our adaptive method for estimating contamination levels. To start, we consider univariate estimation using the robust one-sided Huber loss function. We then discuss the generalization to multivariate regression.

Given the nature of signal contaminants in Ca^2+^ imaging datasets, we created a noise model based on the observation that most fluctuations in the fluorescence background are well modeled as being Gaussian-distributed. This type of noise stems from the stochastic emission, propagation and detection of photons, which are all Poisson processes, implying that the numbers of detected photons are Gaussian-distributed when there are large numbers of photons. However, the fluorescence background also contains other sources of noise or contamination, such as from neuropil Ca^2+^ activity, out-of-focus cells, and residual activity of overlapping cells that are not detected and well accounted for by the cell extraction method. This latter category of contamination is very distinct from normally distributed noise; namely, it is non-negative (or above the signal baseline), its characteristics can be highly irregular, and it may take on large values. Hence, we model the data generation process as having an additive noise source that is normally distributed a fraction **1** − ϵ of the time, but which is free to be any positive value greater than a threshold otherwise:

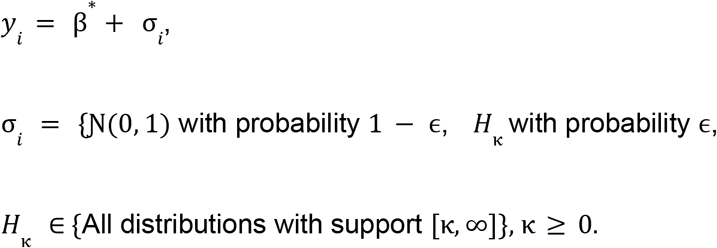

Here, *y*_*i*_ denotes an experimental observation, which deviates from β^*^, the true value of the measured quantity, due to corruption with an additive noise term, σ_*i*_. This noise term, σ_*i*_, is normally distributed with **1** − ϵ probability and according to an unknown distribution, *H*_κ_, with probability ϵ. For the sake of generality, we allow *H*_κ_ to be any probability distribution with support over the range [κ, ∞), for a value κ ≥ 0. When κ is estimated adaptively, it is determined for each cell and time bin directly from the movie data, as part of the loss minimization process (see below). Hence, ϵ can be interpreted as setting the extent or severity of ‘gross contamination’. If ϵ is small, the noise will be close to Gaussian-distributed. On the other hand, as ϵ nears one, the noise distribution deviates from a normal distribution to an arbitrary extent. The parameter κ can be interpreted as a threshold for automated detection of pixels with outlier levels of fluorescence activity, as detailed in **Fig. 1F, H–L** and **Supplementary Note 2**. We denote the full distribution of the noise as 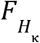, subscripted by *H*_κ_.

Given a set of experimental observations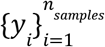, we form an estimate, 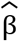, of the true parameter, β^*^, by considering an equivariant M-estimator:

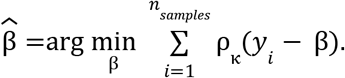

Typically, M-estimators are characterized by estimator functions, Ψ_κ_, that are defined as the derivative of 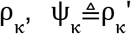. Here we consider Ψ_κ_ with specific properties that enable efficient optimization and allow general theoretical guarantees.

We define a set, Ψ = {Ψ_κ_ ∣Ψ_κ_ *is a monotonically increasing function*}, which implies that each member of Ψ is a derivative of a convex function^58^. If we choose a set of estimator functions, Ψ_κ_ ∈Ψ, finding a point estimate, 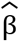, is equivalent to solving the following first-order condition for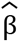:

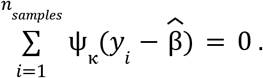

We seek an M-estimator for our noise model that is robust to variations in the noise distribution (*H*_κ_ in particular), in the sense of minimizing the worst-case deviation from the true parameter, β^*^, as measured by the mean squared error. Following our prior work^35^, we define an estimator function, Ψ_0_, as follows:

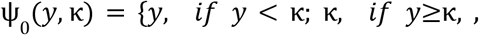

where κ is defined in terms of the contamination level, ϵ, according to

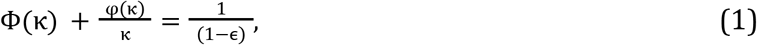

in which Φ(⋅) and φ(⋅) denote the cumulative distribution and density functions for a standard normal variable. Ψ_0_ (*y*, κ) is the estimator function for the ***one-sided Huber loss***, ρ_κ_ (.). Clearly, Ψ_0_∈ Ψ, and therefore the loss function, ρ_κ_(.), is convex. When the value of κ is the same for all movie pixels, Ref.^35^ provides an asymptotic minimax result for Ψ_0_ :

##### Proposition 1.

*The one-sided Huber function*, Ψ_0_, *yields an asymptotically unbiased M-estimator, i*.*e*., *an estimator that is unbiased in the limit of an infinite number of data samples, for the family of noise distributions, F* = {(**1** − ϵ)Φ + ϵ*H*_κ_ }, *that denotes a weighted mixture of Gaussian and non-Gaussian noise contaminants with weights* (**1** − ϵ) *and* ϵ, *respectively. Further*, Ψ_0_ *minimizes the worst-case asymptotic variance in F, i*.*e*.,

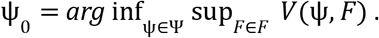

***Proof:*** A proof is provided in our prior work^35^. Here, we reproduce it with more elaboration. First, to obtain an M-estimator, we use the law of large numbers to obtain the first-order condition, *E*_*F*_ [Ψ_0_ (σ − *c*)] = 0, with σ denoting the underlying noise variable from a noise distribution, *F* = (**1** − ϵ)Φ + ϵ*H*_κ_. In the limit of an infinite number of samples, *c* approaches the bias of the estimator and is zero for an unbiased estimator. The condition for unbiasedness is:

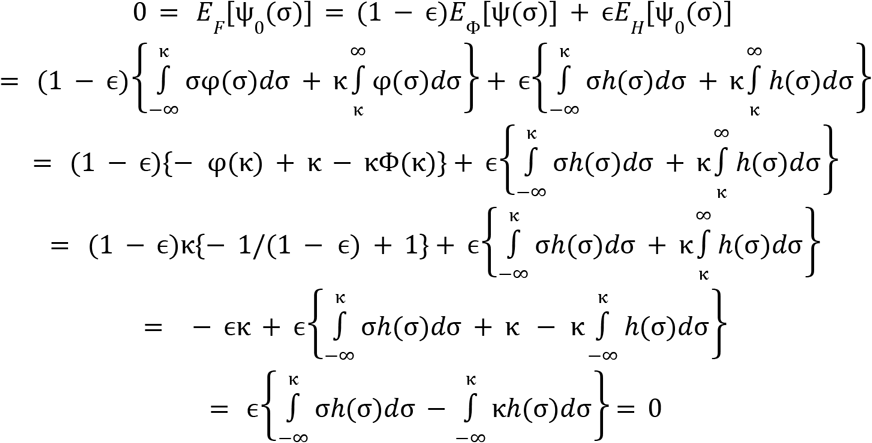

Here, we used the identity Φ(κ) + φ(κ)/κ = **1**/(**1** − ϵ) during the transition from the third to fourth row, and the fact that support of *H* is [κ, ∞) in the final row.

Next, we calculate the variance of the unbiased estimator, with the formula 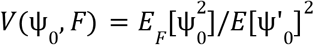. This formula can be calculated similarly to as in the original work^34^ by Huber, *i*.*e*., by considering the Taylor expansion around the first-order optimality condition. A straightforward calculation^35^ leads to *V*(Ψ_0_, *F*) = [(**1** − ϵ)Φ(κ)] ^−**1**^. Notably, this is a constant over the contamination class, *H*_κ_, meaning that it does not depend on the specific form of the distribution, only the contamination level κ.

Now, we define the following distribution *F*_0_ by its density function, *f*_0_ :

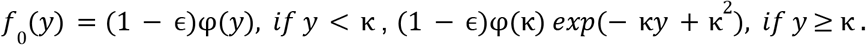

*F*_0_ is indeed Gaussian for *y* < κ and integrates to one, thus *F*_0_ ∈ *F*. Since the variance of estimators within family *F* is constant, and since *F*_0_ is an estimator within this class, we arrive at:

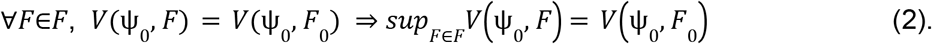

This relates the worst-case variance of Ψ_0_ across all possible noise distributions within the *F* family.

The next step is to relate the variances of all estimators given a common noise distribution, *F*_0_.

Specifically, the application of the Cauchy-Schwartz inequality to the variance formula yields

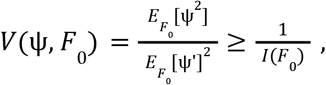

where *I*(*F*_0_) = (**1** − ϵ)Φ(κ) = *V*(Ψ_0_, *F*_0_) is the Fisher information for estimation of β^*^, *i*.*e*., the minimum asymptotic variance of our estimate, 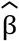. Then, we obtain

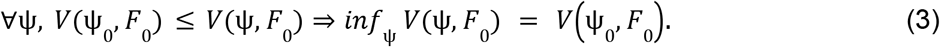

Hence, for the noise distribution *F*_0_, we have related the variance of all possible estimators, Ψ, with that of the optimal estimator, Ψ_0_. Now, we are ready to combine the results of Eqs. (2) and (3):

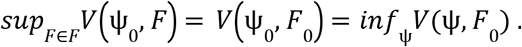

To connect back to the original statement, we note the following:

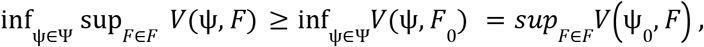

which implies the equality:

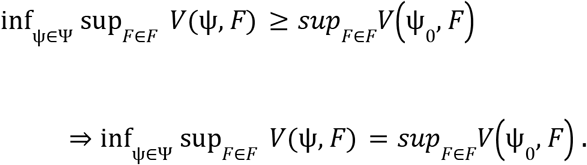

This says that Ψ_0_ minimizes the worst-case asymptotic variance in *F*, concluding the proof.

We now discuss the multivariate regression that we use to estimate the spatial and temporal weight matrices, ***S*** and ***T***. We illustrate the simple case of solving for one row of 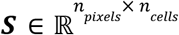 or one column of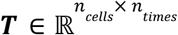. The experimental observations comprise the set of data samples, 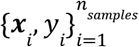, where *n*_*samples*_ refers to either *n*_*pixels*_ or *n*_*frames*_ depending on the subproblem of interest, 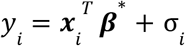, where 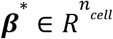 is the true value of the parameter, 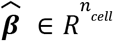, to be estimated, and σ_*i*_ are noise components. We estimate ***β***^*^ as

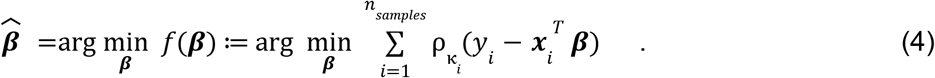

Classical M-estimation theory establishes, under certain regularity conditions, that the minimax optimality in the univariate case carries over to multivariate regression; we refer the reader to Ref.^46^ for details. Here, because these regularity conditions do not always apply to neural recordings, we opted to validate multivariate robust regression with a one-sided Huber loss function by performing extensive computational experiments (**Table S1**). We discuss below and in **Supplementary Note 2** how κ_*i*_ are estimated from the movies in an adaptive manner.

### Solving the robust regression problem with a fast, custom solver

We seek to solve the robust regression problem of Eq. (4) in a large-scale setting, namely that of Ca^2+^ videos with wide fields-of-view and extended durations. Hence, the solver for our problem should ideally be tractable for a large value of *n*_*samples*_ and yield results that are as accurate as possible. To this end, we propose a fast optimization method that has a step cost equal to that of gradient descent, while making use of second-order information and exhibiting convergence behavior similar to that of Newton’s method:

#### Algorithm 1

**Fast robust solver**

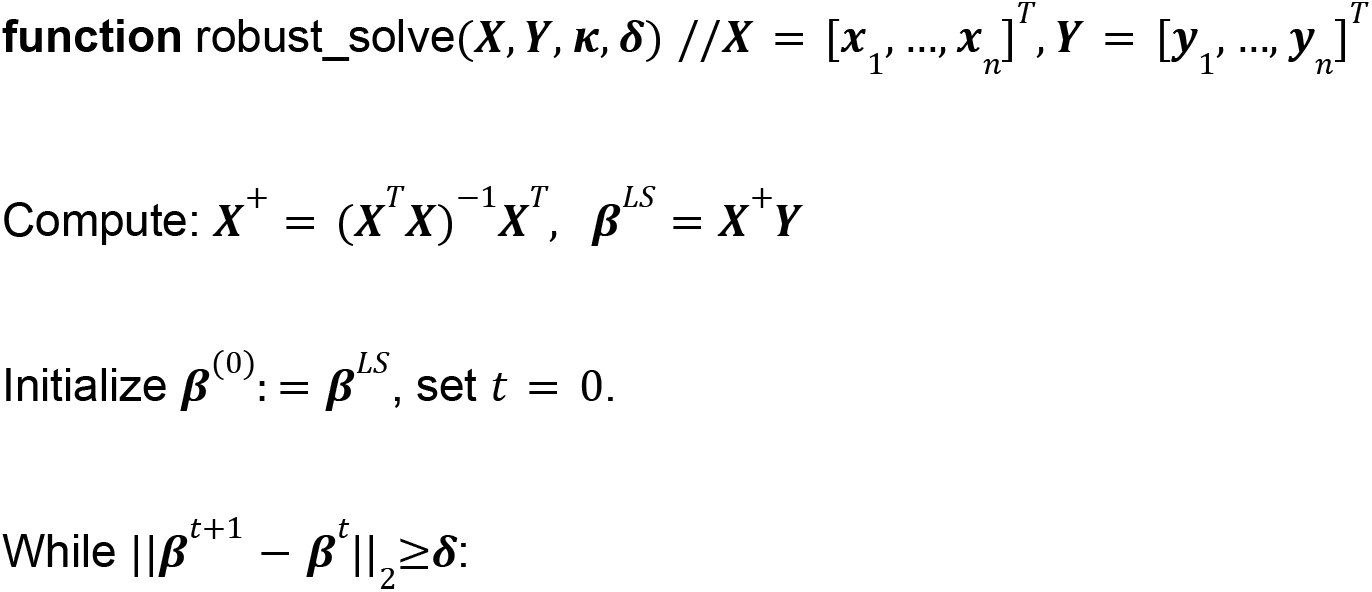
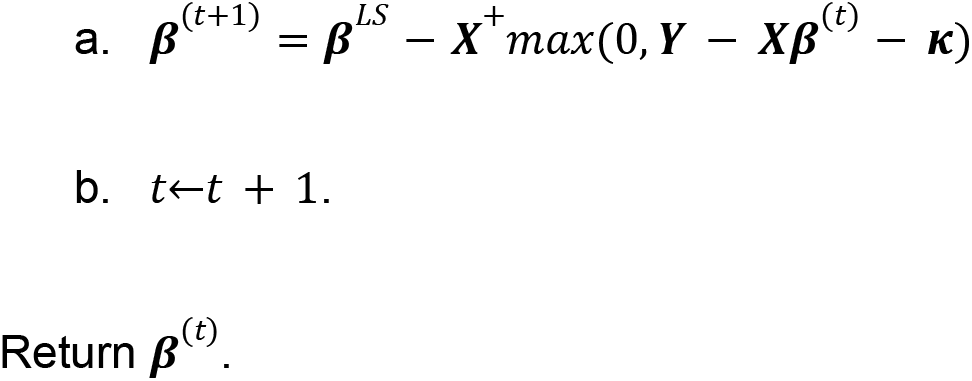

Unlike in Ref. ^35^, here we allow ***κ*** to be a vector with potentially varying entries, and we initialize ***β***^(0)^ : = ***β*** ^*LS*^, which in practice significantly speeds convergence.

Algorithm 1 is used to solve the two subproblems of cell extraction, *i*.*e*., estimation of cells’ spatial profiles (while Ca^2+^ traces are held fixed) and the estimation of Ca^2+^ activity traces (while cells’ spatial profiles are held fixed.) Owing to the matrix operations in Algorithm 1, these subproblems can be quickly solved on GPUs by vectorizing them across the entire Ca^2+^ movie (**Supplementary Note 2**). Below we present a proposition and then its proof regarding the convergence of the solver described in Algorithm 1.

#### Proposition 2.

*Let* 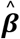 *be the solution of Algorithm 1 for the problem in Eq*. (4), *and let* λ_*max*_ *and* λ_*min*_ > 0 *denote the extreme eigenvalues of* 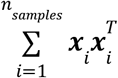, *and let max*_*i*_ ‖ ***x***_*i*_ ‖_2_ ≤*k. Assume that for a subset of indices S*⊂ {**1**, 2, …, *n*}, ∃Δ_*s*_ > 0 *such that* 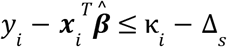, *and denote the extreme eigenvalues of* 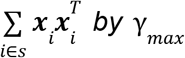 *and* γ_*min*_ > 0 *satisfying*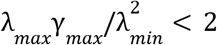. *If the initial point* ***β***_0_ *is close to the true minimizer, i*.*e*., 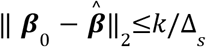, *then Algorithm 1 converges linearly*,

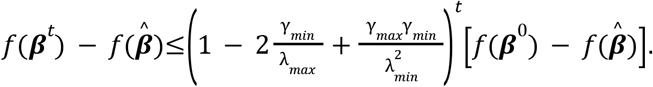

### Proof

This proof is similar to the one we provided earlier for the solver in our prior work^35^ using a homogenous ***κ***. We consider the following objective function,

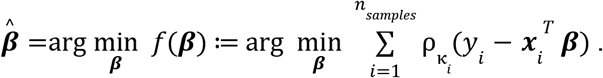

We start by assuming 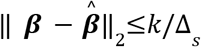 for some ***β***, the variable over which we are minimizing.

Then, we have for ∀*i*∈*S*,

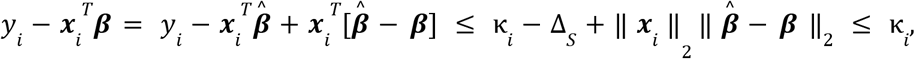

where we used the fact that the dot product between two vectors can be upper-bounded by the product of their norms and the assumptions 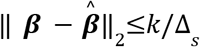 and *max*_*i*_ ‖ ***x***_*i*_ ‖_2_ ≤*k*. Notably, the inequality that 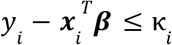 is also the condition that a particular sample *i* is in the quadratic regime of the loss function, *i*.*e*., contributes to the Hessian calculations. However, though we have shown that for all ∀*i*∈*S*, the *i* th sample contributes to the Hessian calculation, the inverse statement is not necessarily true. In other words, there may be other samples contributing to the Hessian, leading to the following property of the Hessian:

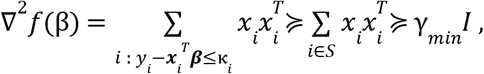

which says that in the ball 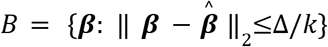, the objective function *f* is γ_*min*_ -strongly convex. Yet, strong convexity implies smoothness, i.e., ∇^2^*f*≼γ_*max*_ *I* for ∀***β***∈*B*.

Assuming that the current iterate is ***β***^(*t*)^, our approach takes a step of the following form:

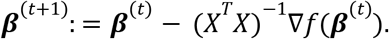

By γ_*max*_ -smoothness, we can write

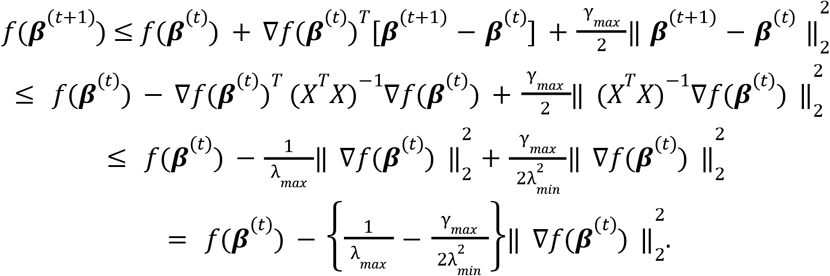

By γ_*min*_ -strong convexity, for any ***β*** and ***β***’:

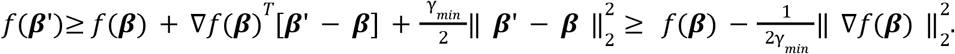

The second inequality follows from setting ***β***’ = ***β*** − **1**/γ_*min*_ ∇*f*(***β***), which is the minimizer of the right-hand side. Choosing 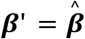 above yields

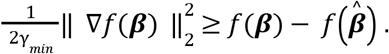

Using this and the smoothness inequality above, we write

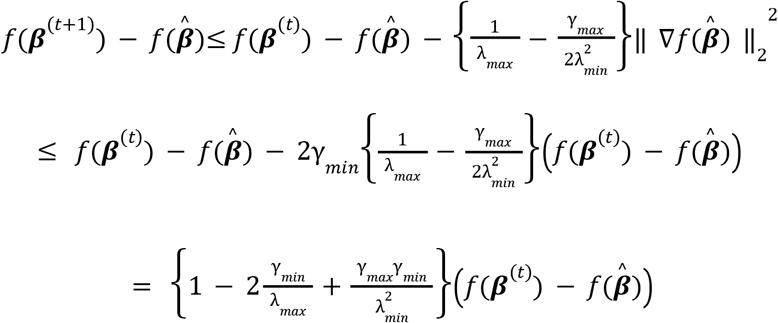

This is linear convergence with coefficient 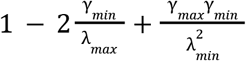, as long as the condition,

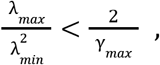

is satisfied. This concludes the proof.

Empirically, we found that the fast solver retains fast convergence even outside the realm covered by this proof and is compatible with the ADMM framework (**Supplementary Note 2**).

#### Relation between our fast solver and Newton’s method for the robust estimation problem

For a convex function 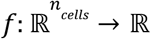, unconstrained Newton update on the parameter 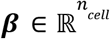 entails

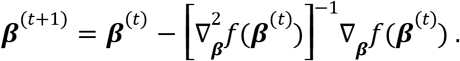

In our algorithm, within the update step above we approximate the Hessian as 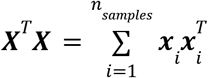. instead of 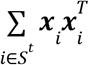 **(Supplementary Note 2**). This implies that our solver is second-order and so its convergence behavior should be similar to that of Newton’s method. However, there is one caveat: the second-derivative of the one-sided Huber loss is not continuous. Hence, one cannot expect a quadratic rate of convergence; this issue of a non-continuous second derivative is commonly encountered in robust estimation^34^. Nevertheless, Algorithm 1 converges very quickly in practice.

#### Setting κ Adaptively in Robust Estimation

To estimate ***κ*** adaptively, we first estimate the contamination level, ϵ, for individual cells and each time bin. For a given time bin, assume that 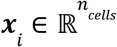 is the weight of the cell’s spatial footprint at pixel *i, yi* is the fluorescence in pixel *i*, and 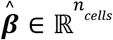 is the set of estimated Ca^2+^ activity traces for the cells occupying pixel *i*. To compute the contamination level for each cell, we first examine the residuals, 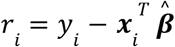, at each pixel. Then, we compute the fraction of positive residuals, averaged across all the pixels occupied by a particular cell of interest, *j*:

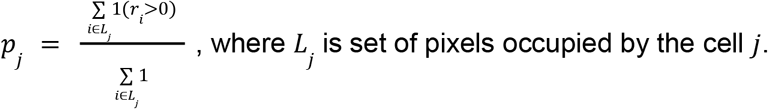

In practice, we do not update ***κ*** at every iteration of the solver in Algorithm 1; hence, we will use a distinct variable, *q*, to denote the iterations of the adaptive updates of ***κ***. Specifically, we update the estimated probability of non-Gaussian contamination for cell *j* using:

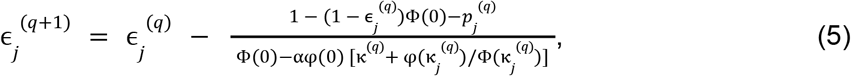

where α = 0. 05 sets the update rate and κ_*j*_ ^(*q*)^ and ϵ_*j*_ ^(*q*)^are initial values or prior estimates (**Supplementary Note 2**). The estimates, κ^(*q*+**1**)^ and ϵ^(*q*+**1**)^, replace the prior ones, κ_*j*_ ^(*q*)^ and ϵ_*j*_ ^(*q*)^. In practice, we start with a value of κ_*j*_ ^(0)^ that is initialized to a user-provided value (by default, 0.7 sd of median pixel activity). Then, we compute the corresponding contamination level ϵ_*j*_ ^(0)^. Next, we first update the contamination level using Eq. (5) and obtain the corresponding ***κ*** using Eq. (1).

Overall, we summarize the procedure to estimate ***κ*** below:

##### Algorithm 2

Estimation of ***k***:

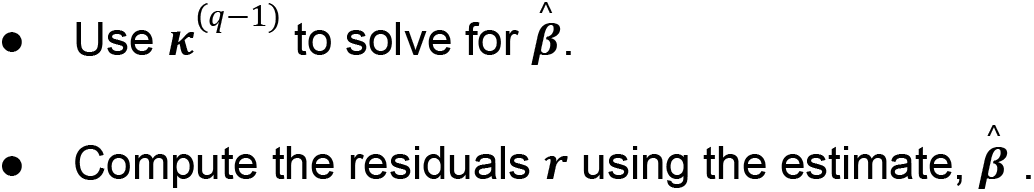
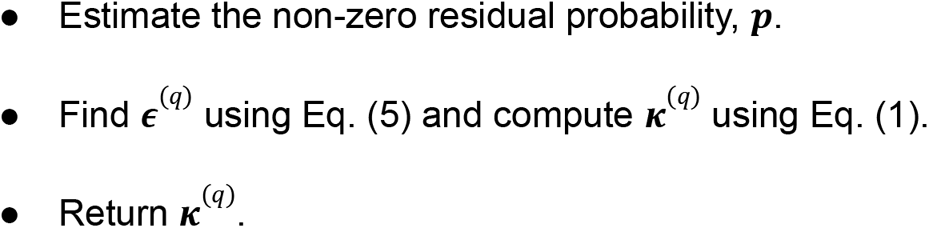

#### Interpretation of the robust loss function as an automated detector of outlier data points

The specific form of the fixed-point solver in Algorithm 1 allows us to define a variable that functions as a detector of outlier data points (**Supplementary Note 2**):

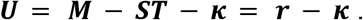

If it were the case that ∀_*i,j*_ : *U*_*ij*_ < 0 for all data points at pixel coordinates (*i, j*), then a minimization of the one-sided Huber loss function would be equivalent to a least-squares minimization. However, when there are outlier data points that make the least-squares solution suboptimal, the optimization of robust loss function involves an automated subtraction of outliers. To elaborate, for outlier data points with *U*_*ij*_ > 0, the values of the pixel fluorescence are projected to the outlier detection margin, ***r*** = ***κ*** (**Fig. 1H–L**). Then, any residual activity beyond the margin, *i*.*e*., ***r*** > ***κ***, which cannot be reasonably explained as reflecting either neural activity or a Gaussian noise component, is subtracted from the movie to obtain:

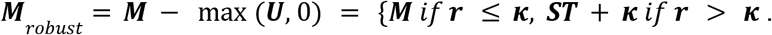

Once the outlier contributions are subtracted in this way, the least-squares estimates of cells’ Ca^2+^ activity traces or spatial profiles are obtained from ***M***_*robust*_. This procedure has a simple geometric interpretation; namely, minimization of the robust loss function involves subtraction from the movie of any unexplainable (*i*.*e*., with support on [κ(*x, t*), ∞)) residual fluorescence activity (**Figs. 1I–L**).

Regardless of the value of the non-Gaussian positive contaminant, if its support is within [κ(*x, t*), ∞), then the contaminant noise will be deemed an outlier and projected to the outlier detection margin, ***r*** = ***κ***. This geometrical interpretation also applies to the original, symmetric Huber loss function^34^, but to our knowledge has not been previously discussed in the published literature. This principled treatment of outlier data points is what provides EXTRACT with the key properties of robustness, as described above in the introduction to this work, as opposed to prior colloquial usages of the word ‘robustness’ in the neuroscience literature^26,29^. Here, we go beyond prior uses of robust regression approaches in the published statistics literature, in that our solvers can mitigate large contaminants that vary in both space and time. In other words, the adaptive estimation of ***κ*** used here is important for our application of robust regression to analyses of large-scale neural recordings, as the use of a homogenous ***κ*** leads to suboptimal estimation of Ca^2+^ activity traces (**Fig. 1G**).

### EXTRACT ALGORITHM

The EXTRACT pipeline applies our one-sided Huber loss function to the cell extraction problem. The pipeline has a modular structure, which affords the user operational flexibility and comprises four base modules:

1. Preprocessing module.
2. Cell finding module.
3. Cell refinement module.
4. Final robust regression module.

The first module transforms the Ca^2+^ video into a standardized format for the downstream modules; it also performs an optional spatial filtering of the movies, with the goal of removing noise and making the videos more suitable for cell extraction. The main goal of the second and third stages is to accurately estimate cells’ spatial filters; the user has the option of running these two stages on a temporally downsampled movie. For faster processing speed, cell finding and refinement are, by default, run without adaptive estimation of ***κ*** and cells’ baseline activity values, which take on fixed, user-defined values (*e*.*g*., the default, baseline activity value is Δ*F*/*F*_0_ = 0, below which cell activity is truncated to zero). This default is motivated by our observation that use of adaptive estimation within these intermediate modules generally does not lead to substantially different results, although the user can select to adaptively estimate these quantities if desired. By comparison, the goal of the final robust regression is to accurately estimate cells’ activity traces, which should thus be performed without temporal downsampling and with adaptive estimation of ***κ*** and cells’ baseline activity values. When running the EXTRACT pipeline, the user can choose to bypass one or more of the above modules, *e*.*g*., to perform their own pre-processing, provide their own initialization parameters for cells’ spatial footprints that are later refined, or perform the final robust regression with a set of spatial filters, ***S***, provided by the user.

#### Definition of signal-to-noise ratio (SNR)

Before discussing the details of the EXTRACT pipeline, we first define the signal-to-noise ratio (SNR) for a given signal as the signal’s maximum value divided by the s.d. of the noise. We estimate the noise s.d by obtaining the power spectral density of the signal with a Fourier transform, integrating the spectral power across the upper half of the frequency range (where most of the Ca^2+^ trace comprises noise fluctuations), and extrapolating the power found there to the rest of the spectrum. We compute the SNR at an individual image pixel by considering the time-varying fluorescence from that pixel as the signal.

#### EXTRACT preprocessing module

This module takes the input Ca^2+^ movie, which is presumed to have undergone motion-correction but no other preprocessing, and puts it into a standardized format for efficient and effective processing. The movie is first re-expressed in terms of the relative fluorescence changes that occur at each pixel (Δ*F*). (A normalization by baseline fluorescence levels, *F*, is applied to each cell’s activity trace only after the final robust regression, right before EXTRACT delivers its final outputs, so that the traces provided are expressed in units of Δ*F*/*F*_0_). This is followed by an optional spatial high-pass filtering, which can be beneficial in some cases for removing contaminants that may exist at low spatial frequencies, such as from neuropil Ca^2+^ activation.

The spatial high-pass filter used in this second step is a fourth-order high-pass Butterworth filter that is designed in the frequency domain, with a cutoff set by the user-provided average cell radius. To design this filter, we first model cells’ spatial footprints as 2D Gaussian functions, with the cell radius equal to twice the s.d. of the Gaussian function. We set the corner frequency, *w*_*c*_, of the high-pass filter to the spatial frequency corresponding to twice the s.d., *i*.*e*.,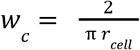, where *r*_*cell*_ is the user-defined cell radius. Then, the cutoff frequency of the Butterworth filter is determined by dividing the corner frequency by a dimensionless factor that is set by the user (the default value is 5). The user has the option to set the amount of spatial filtering to be distinct in the *x* and *y* spatial directions, *i*.*e*., for cases in which the neuron-type has an anisotropic morphology, such as with Ca^2+^ spikes in Purkinje neuron dendritic trees (**Fig. S8**). The resulting high-pass filter is applied to each frame of the Ca^2+^ movie in the spatial frequency domain and then transformed back to real space. The EXTRACT preprocessing module can also perform an optional spatial low-pass filtering of the raw movie data, for the sake of smoothing and boosting the SNR of cellular signals, for use only by the cell finding module. Like the high-pass filter, this low-pass filter is also a fourth-order Butterworth, for which the cut-off frequency is set by multiplying the corner frequency (see above) by another user-set, dimensionless constant (the default value is 2).

#### EXTRACT Cell Finding Module

In the cell finding module, we first compute a ‘smoothed’ maximum projection image of the whole movie, which we obtain as follows. For each movie pixel in ***M***, we first identify the time point at which the Ca^2+^ activity of the pixel reaches its maximum value. We record this information in an array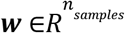, here *n*_*samples*_ stands for the total number of pixels in the movie. We compute the smoothed maximum projection image, 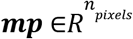, as

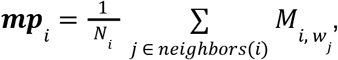

where *N*_*i*_ is the number of pixels that neighbor the *i*’th pixel. In other words, for each pixel *i*, we average its fluorescence values over the time points at which the neighboring pixels reached their maximum fluorescence amplitudes. The function, *neighbors*(·), selects the neighboring pixels of a given pixel; this is done in practice by creating a binary circular mask around the query pixel with a radius of 2 pixels and returning the indices that are nonzero. This procedure for determining the maximum projection image has the property that, for pixels that lie within cells, it reports values close to the maximum value attained at each pixel (owing to the co-activation of neighboring pixels that lie within the same cell); whereas, for pixels that lie outside cells, the smoothing procedure typically yields much lower values.

At every iteration of the cell finding module, a seed pixel is chosen as the brightest pixel in the smoothed maximum projection array, ***mp***, and then a cell image centered at the seed pixel is initialized. This initialization is done either by generating a Gaussian shape with a radius equal to a user-defined estimate of the cell radius. Alternatively, the initialization is done by first computing the Pearson’s correlation coefficient between the seed pixel’s fluorescence trace and the traces of other pixels, with the movie, and then truncating to zero the spatial map of correlation coefficients for all pixels for which the correlation coefficient with the seed pixel was below 0.5 the maximum correlation coefficient. This latter method to initialize the cell image is the default option.

With the resulting estimate of the cell image, the cell’s activity trace is obtained by performing a univariate robust regression of the cell image against the movie data. Next, the cell image is re-estimated using a univariate robust regression, by regressing the cell’s estimated activity trace against the movie. This alternating estimation scheme is repeated either 10 times, or until the relative changes in the cell image and trace estimates are <1% between iterations, as measured by the *L*_2_ norm. For these univariate robust regressions, we optimize the one-sided Huber loss using our custom solver (see Algorithm 1 above), with a non-negativity constraint on both the cell image and the activity trace. Since each regression is univariate in this module, the non-negativity constraint is easily enforced by solving the optimization problem first without it, and then applying the non-negativity constraint at the end, which yields the optimum non-negative solution. After obtaining the cell image, ***s***, and the trace, ***t***, for the identified cell, we subtract the contribution of this cell (*i*.*e*., by setting ***M*** ← ***M*** − ***st***). We then re-compute the smoothed maximum projection, ***mp***, for only the pixels that were affected by the activity subtraction.

At the end of each iteration, we apply a quality check to both the image and the activity trace of the identified cell, to decide whether to include it in the set of identified cells. We discard cells that occupy an abnormal number of pixels given the expected area of a typical cell (as computed from the user-provided estimate of a cell’s radius). We also compute the trace SNR for each cell, and discard it if the trace SNR is less than the user-provided threshold.

We terminate cell finding if any of the following conditions are met: (1) The maximum allowed number of iterations set by the user has been exceeded; (2) The pixel-wise SNR in the current seed pixel is lower than the user-provided SNR threshold; (3) The running yield, defined as the fraction of good cells over the last 10 iterations, is less than 1 in 10. The cell finding module outputs the spatial and temporal weights of the identified components in two matrices: the spatial weights matrix ***S***, whose columns contain the (flattened) cell images, and ***T***, the temporal weights matrix, whose rows contain the corresponding Ca^2+^ traces.

#### EXTRACT Refinement Module

In the refinement module, we alternatively update the entire matrix of spatial weights and the entire matrix of temporal weights, by performing multivariate regressions of each matrix against the movie data, with the constraints that the estimates of both ***S*** and ***T*** must be non-negative, as in the cell finding module. We perform these regressions using the fast custom solver of Algorithm 1, together with a consensus optimization method based on dual ascent, termed the ‘alternating direction method of multipliers’ (ADMM^38^). We use ADMM because it allows us to add the non-negativity constraints in a straightforward manner, with our fast solver, *robust*-*solve*(⋅), as a subroutine within ADMM. **Supplementary Note 2** has further details.

When solving for ***S***, we compute a binary mask ***B*** that is obtained by convolving each cell image with a disk filter of a radius equal to the average cell radius, followed by binary thresholding. We then add the following constraint:

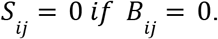

This constraint ensures that the estimation of each component is restricted to a local neighborhood, preventing artifacts that might otherwise arise due to strongly correlated Ca^2+^ activity between spatially separated portions of the movie. This locality constraint defines a convex set, hence it can be added to the estimation problem without violating convexity.

Overall, given ***M*** and ***T***, the ***S***-estimation step solves the following problem:

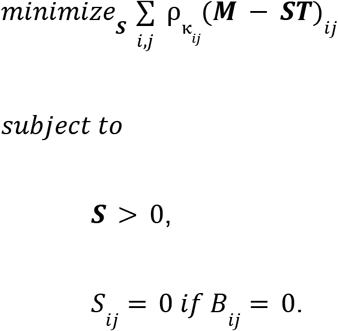

Given ***M*** and ***S***, the ***T***-estimation step solves the following problem:

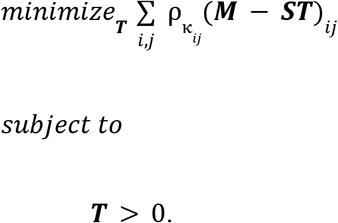

After each alternating estimation step, which involves first solving for ***T*** given ***S***, and then for ***S*** given ***T***, we compute several quality metrics and discard the subset of cells for which any of the computed metrics are worse than certain user-set thresholds. In particular, we compute the following quality metrics:

##### Trace SNR

We compute the trace SNR for each component given its Ca^2+^ trace. We eliminate cells whose trace SNR is below the trace SNR threshold.

##### Area of the cell image

We compute the area of each cell image by summing the number of pixels with spatial weight >0.1 times the maximum weight. If the calculated area is smaller than a lower threshold or higher than an upper threshold, then the cell is discarded.

##### Duplicate cells

We check whether cells are duplicates by separately examining (a) the similarities of cell images, and (b) the overall similarities of cells’ spatiotemporal profiles. For the former check, we first smooth the cell images by convolving them with a two-dimensional Gaussian kernel with σ equal to half the average cell radius. After this, we compute Pearson’s correlation coefficients between pairs of smoothed cell images and then apply a binary threshold at 0.8. We then treat this thresholded correlation matrix as a graph adjacency matrix, and we find the connected components using MATLAB’s *graphconncomp*() function. For each set of connected components, we identify the component with the most edges in the set, and we mark it as a duplicated cell. Although this procedure identifies only one cell per iteration within a highly similar set of cells, we have empirically found it to be effective in eliminating duplicates across iterations of cell refinement. For identification of duplicates based on spatiotemporal similarity, we follow the same procedure, but we fuse the spatial and temporal similarity through the following two steps: (1) We obtain a temporal correlation matrix by first pre-conditioning the temporal matrix, ***T***, with the matrix of correlations between smoothed cell images and then computing the Pearson’s correlation coefficients between pairs of components in the pre-conditioned ***T***. This allows us to enforce spatial proximity within the computations of trace similarity. (2) We obtain a spatiotemporal similarity matrix via an element-wise multiplication of the temporal correlation matrix with the spatial correlation matrix computed above. A binary thresholding is applied to the resulting correlation matrix at 0.95 to obtain the graph adjacency matrix, and the above steps are repeated for this procedure to identify duplicates.

##### Spatial corruption metric

We compute a spatial corruption metric that measures the lack of local smoothness within a cell’s spatial profile. We do this using a heuristic that compares the variance of spatial weights for each cell to a “local variance” for the same cell. We compute the variance as the empirical variance of the spatial weights that are larger than 10^−3^. We compute the local variance as the squared *L*_2_ distance between a pixel’s spatial weight and its weight after applying a 2D low-pass filter based using a square kernel with uniform weights over 4 × 4 image neighborhood. The spatial corruption metric is the ratio of the local variance to the variance of spatial weights. Intuitively, better looking cell spatial weights have negligible local variance when compared to the variance, so the spatial corruption metric will be small for these cells. In the algorithm, the default threshold value for spatial corruption is set at 1.5, based on the typical distribution of spatial corruption metric values across many different datasets.

##### Spatiotemporal match metrics

We use a quality metric that is intended to assess the spatiotemporal contribution of the cell, relative to its surrounding, with respect to the power of the cell signal. This metric looks at the mean gap (averaged over movie frames) between a cell’s fluorescence trace and nearby fluorescence activity in its spatial vicinity, by removing the portion of the estimated activity belonging to a cell that is also represented in its surroundings. Our implementation for this metric can be found in our codebase inside the function *find_spurious_cells*(), which can be referred to for full details on how the various fluorescence activity traces are computed. This metric must be > 0.01 for EXTRACT to accept the identified cell in the output, though we keep this metric off by default for most movies to prevent prolonged runtimes.

#### EXTRACT Final Robust Regression Module

As noted above, the cell finding and refinement steps are usually performed on a temporally downsampled version of the Ca^2+^ movie and without adaptive estimation of ***κ*** and/or cells’ baseline activity values. The final set of Ca^2+^ activity traces, however, should generally be obtained from the original movie, without downsampling, and with adaptive estimation. In other words, the final robust regression module solves the following optimization problem:

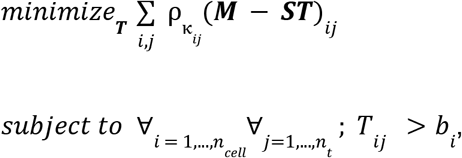

where the cell baselines, *b*_*i*_, and robustness parameters, κ_*ij*_, are adaptively computed from the movie. As in the cell refinement module, the final robust regression is performed using the fast custom solver of Algorithm 1, along with ADMM (See **Supplementary Notes 1, 2**).

#### Algorithmic Output

EXTRACT provides 3 main options for the final set of estimated Ca^2+^ activity traces, termed ‘non-negative’, ‘baseline_adjusted’, or ‘no_constraint’ in the software Github. In all cases, the robust solver operates under the constraint that Ca^2+^ signals must be non-negative until the end of the cell refinement process. The motivation for this constraint is that EXTRACT considers activity below each cell’s baseline level to be noise, where the baseline is determined by the cell’s time-averaged, mean fluorescence level. The algorithm truncates all activity below this baseline, leading to intermediate non-negative activity traces, which are then used to estimate cells’ spatial profiles accurately using mainly the Ca^2+^ events. Then, based on the option selected, EXTRACT performs a final robust estimation to solve for the activity traces using the final set of cells’ spatial profiles, which may include the noise baseline depending on the solver type chosen.

#### Getting Started with EXTRACT

To help users get started with EXTRACT, we designed six tutorials (**Figure S9)** to familiarize the users with the hyperparameters of different modules in EXTRACT. These tutorials also include additional functions to pre-process Ca^2+^ imaging movies, including a wrapper code for motion correction, and to visualize the cell extraction results. The details are provided in our user manual (**Supplementary Note 5**) and the tutorial codes can be found in our Github repository^59^.

### SIMULATION BENCHMARK

#### Simulated Ca^2+^ Imaging Datasets

To benchmark the performance of different cell extraction algorithms, we created artificial datasets designed to be representative of the Ca^2+^ activity of cortical pyramidal neurons. In all cases, the simulated Ca^2+^ videos had a frame-rate of 10 fps. Here we summarize the process of creating the Ca^2+^ videos; **Supplementary Note 4** lists the default values of the simulation parameters and has additional details for how these parameters were varied across the many different computational experiments that we performed (**Table S1**). Generation of the artificial Ca^2+^ videos involved 3 steps.

In the first step, we simulated Ca^2+^ traces of neurons with exponentially decaying, optical spike waveforms. To achieve this, we first simulated Ca^2+^ event trains for each cell by assuming that the probability of a Ca^2+^ event occurrence in each time bin was governed by a Bernoulli random variable with a probability of 0.01, corresponding to a Ca^2+^ event rate of *r*_*event*_ = 0. 1 Hz. Since real Ca^2+^ events can report the occurrences of multiple action potentials or spike bursts via their amplitudes, for our simulations we next generated a set of randomly determined, discretely valued yet non-binary Ca^2+^ amplitudes for each of the Ca^2+^ events in the video. The minimum Ca^2+^ event amplitude, corresponding to the occurrence of a single action potential, was set by multiplying a user-defined value of the s.d. of the noise level within individual pixels of the movie, σ_*pixel*_, with a parameter called the ‘minimum SNR’ per cell, *SNR*_*min*_. This allowed us to set randomly the amplitudes, *T*_*amp*_, for each Ca^2+^ event using the model:

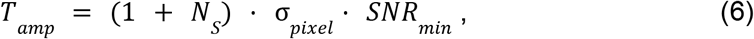

where *N*_*s*_ = *Poisson* (*A*_*spike*_) is a Poisson random variable with *A*_*spike*_ ≥ 0 setting the characteristic variations in amplitude. In this model, if *A*_*spike*_ = 0, the Ca^2+^ events all take on equal amplitudes, whereas if *A*_*spike*_ > 0, the Ca^2+^ events have discretely valued but randomly varying amplitudes. Then, we built in refractory periods by deleting spikes that occurred exactly one frame after a prior spike. Since the simulated firing rates were low, such deletions occurred with extremely low probability and barely altered the net rates of Ca^2+^ events. Finally, having set the amplitudes of all Ca^2+^ events, we convolved the resulting event trains with an exponentially decaying temporal kernel of the form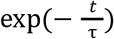, with τ = 10 time bins. This corresponds to a decay time constant of 1 s, roughly comparable to that of *e*.*g*., GCaMP6m^4^ or jGCaMP7s^60^. For simulated movies of independent spiking across the different cells in the movie, the resulting Ca^2+^ traces were then incorporated into the artificial movie (see below).

However, to simulate movies with correlated neural spiking, instead of generating the activity trace of each cell independently, we synchronized the instantaneous firing probabilities within groups of cells, such that each cell fires *p*_*synch*_ of its Ca^2+^ events jointly with the group. Each cell can participate in many different groups. To simulate high levels of correlated activity, for each time point we clustered randomly chosen cells (*i*.*e*., without regard to their spatial locations) into groups of a randomly chosen size, uniformly distributed between 50–100 cells. Then, for each time point, with an adjusted firing probability *p*_*synch*_ × *r*_*event*_, we assigned a new Ca^2+^ event to each neuron such that every cell in the group had a synchronized Ca^2+^ event at that time point. The amplitudes of the synchronized Ca^2+^ events were set independently for the different cells in the group using Eq. (6) above. We independently adjusted the baseline event rate of each cell to keep its mean firing rate constant at 0.1 Hz. We enforced a refractory period by deleting all spikes that initiated exactly one frame after a previous spike. Finally, as above we convolved the Ca^2+^ event trains with an exponentially decaying temporal kernel, 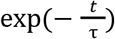, with τ = 10 time bins.

In the second step of movie generation, we created the spatial profiles of individual cells. We used a square field-of-view, with each pixel having a width of 1.6 μm in the specimen plane. In nearly all experiments, with the exception of Experiment 21 (**Table S1**), we created each cell’s fluorescence image by randomly sampling a two-dimensional Gaussian distribution that was oriented in a random direction relative to the *x-y* coordinate axes of the movie. For each cell, we approximated its effective diameter as the width of this Gaussian, for which the s.d. was selected randomly from a uniform distribution ranging between 3.5–4.5 pixels. We then truncated to zero all pixels of the Gaussian spatial profiles that had values <5% of the peak value. We then randomly set the centroid of each cell from a uniform spatial distribution across the field-of-view. However, we required that no pair of cells could have their centroids <4 pixels (6.4 μm) of each other.

For one experiment, Experiment 21, we compared the performance of several cell extraction algorithms using Ca^2+^ movies with donut-shaped neurons. In this case, we used the functional form in Ref. ^18^ to simulate the spatial profile for donut shaped neurons:

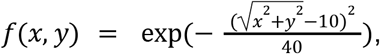

where *x* and *y* are in units of pixels. Donut-shaped spatial profiles were truncated and distributed across the field-of-view as described above for all other computational experiments (**Table S1**).

In the third and the final step, we generated the noise components of the synthetic Ca^2+^ movie by randomly sampling values from a Gaussian distribution that were uncorrelated for each spatial pixel and time point, with a s.d. that was assigned according to the desired mean pixel-wise SNR for the simulated movie. This uncorrelated Gaussian noise models the stochastic emission, propagation and detection of photons, which are governed by Poisson processes at the level of individual photons but which approximate Gaussian processes in large photon numbers. In addition to this spatiotemporally uncorrelated noise, we also added correlated noise to the movie, representing non-negative neuropil contamination, by applying a spatial bandpass filter (4th-order Butterworth filter with low and high cutoff frequencies of 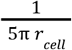 and 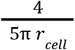, respectively, where *r*_*cell*_ is the mean cell radius used in the simulations) to i.i.d. samples of Gaussian noise at each movie pixel and then convolving the resultant with a temporal kernel identical to that used for the Ca^2+^ transients in the neural activity traces. The total Gaussian noise at each movie pixel was determined as a weighted sum of the uncorrelated (95% weight) and correlated (5%) noise traces. Finally, for simulations of one-photon fluorescence Ca^2+^ movies, we added non-Gaussian background Ca^2+^ activity by simulating out-of-focus cells with varying diameters that were uniformly distributed between 50–180 μm. (Please see **Supplementary Note 4** for more detailed information about each computational experiment and how background Ca^2+^ activity traces were generated). In all cases, we generated the final synthetic Ca^2+^ movie by taking the product of the matrix of all cells’ spatial weights and that of their Ca^2+^ traces (including background cells when appropriate, *e*.*g*., for simulating one-photon Ca^2+^ imaging movies; **Supplementary Note 4**), and then adding to the resulting noiseless movie the matrix holding the total time-dependent noise values for each pixel.

#### Systematic evaluations of cell extraction quality

For the different cell extraction algorithms evaluated in this paper, we benchmarked several aspects of cell extraction quality, such as the ability to accurately estimate cells’ spatial footprints and Ca^2+^ activity traces. To quantify the results, we defined several metrics that used the known ground truth underlying the simulated Ca^2+^ videos. In addition, we defined one metric of trace quality that focuses on the similarity of a cell’s estimated Ca^2+^ trace to the Ca^2+^ activity seen in the movie for the same cell.

##### Evaluating the cell-finding accuracy in simulations

Substantial prior research tested the quality of cell-finding and focused on an algorithm’s ability to find human-annotated cells within a set of Ca^2+^ videos^11,19,20,61^. Owing to this body of research, several quality metrics that quantify cell-finding accuracy have become relatively standardized and are summarized here (**Figure 3**). The cell-finding ‘Precision’ is defined as the number of correctly identified cells, divided by the total number of cell candidates that the algorithm outputs. The cell-finding ‘Recall’ is defined as the number of correctly identified neurons, divided by the total number of true cells in the movie. The ‘F1 score’ for cell-finding is a joint summary metric that is defined as the harmonic mean of the Recall and the Precision:

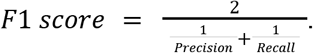

All three of these metrics range from 0–1, with 1 being the best score. We emphasize that Precision and Recall must be considered as a pair, for it is easy for an algorithm to achieve a perfect score in either one of the two individual metrics, but there is nearly always a crucial trade-off between the two. Thus, assessments of which algorithms achieve superior cell-finding performance should be made by examining the Precision-Recall (PR) curve (**Figure 3B, S4A**), which quantitatively describes the trade-off between the two metrics. Thus, use of the F1 score helps to find the optimal point in the PR curve that maximizes the harmonic mean of the two constituent metrics.

##### Trace quality metrics for use with simulated datasets

We used two different metrics to quantify the quality of trace estimation from simulated Ca^2+^ videos, namely the accuracy of crosstalk mitigation and that of signal amplitude estimation. The former metric operates at the level of Ca^2+^ events and quantifies how well individual Ca^2+^ events are identified, which requires both the correct identification of a cell’s Ca^2+^ signals and suppression of crosstalk from neighboring cells. The latter metric quantifies how well the exact amplitudes of a cell’s Ca^2+^ transients are estimated. A key distinction between the two metrics here is that the crosstalk mitigation accuracy does not assess whether Ca^2+^ signal amplitudes are correctly estimated, whereas signal amplitude estimation accuracy does not assess how well contamination from other sources is suppressed within a cell’s estimated Ca^2+^ activity trace.

To evaluate the levels of crosstalk in a set of estimated Ca^2+^ activity traces, our metric examines the number of false positive Ca^2+^ events that must be tolerated to identify a given level of true positive Ca^2+^ events. We create a PR curve for Ca^2+^ event detection for each cell and compute the area under the curve (AUC). This AUC value is then averaged across all cells identified by the cell extraction algorithm, yielding the value of the crosstalk mitigation accuracy.

To extract Ca^2+^ events from cellular activity traces, we applied an exponential deconvolver (with an exponential decay time-constant matching the ground truth value for the simulations) to the set of estimated activity traces provided by the cell detection algorithm. We swept the detection threshold from high to low values across the range 1–0, in units of each cell’s peak Ca^2+^ signal. Then, to compute Ca^2+^ event Recall and Precision metrics, we matched ground truth Ca^2+^ events to the set of detected Ca^2+^ events by using a greedy matching scheme. Specifically, we computed a matrix of temporal intervals between the sets of ground truth and detected events, applied a matching threshold so that the time intervals of matched events had to be ≤ 3 image frames, and then matched the events greedily in an iterative manner. With the set of matched events in hand, we computed the Recall as the ratio of correctly detected Ca^2+^ events to the total number of ground truth Ca^2+^ events. We determined the Precision as the ratio of correctly matched events to the total number of detected Ca^2+^ events. As we swept the Ca^2+^ event detection threshold from high to low values, we maintained the Recall values of Ca^2+^ events that were matched at previous, higher values of the detection threshold, and greedily increased the Recall values as new Ca^2+^ events were matched for the new, lower detection threshold.

To compute the signal estimation accuracy, we computed Pearson’s correlation coefficient between the ground truth Ca^2+^ trace and the estimated Ca^2+^ trace at the time bins at which the ground truth events occurred. Unlike correlation coefficients computed across the entire movie duration, as has been previously used as a trace quality metric^20^, in our metric we focused on signal estimation by dissecting away the effects of crosstalk. Overall, we quantified two important aspects of trace quality, crosstalk and event amplitude, with separate quality metrics.

##### A trace quality metric applicable to real experiments

To assess the quality of trace estimation for real Ca^2+^ videos, we formulated a ‘trace quality metric’ that does not require the use of ground truth activity traces but remains interpretable. With simulated datasets, we found that this trace quality metric followed the general trends as the crosstalk and trace accuracy metrics described above, arguing for its validity and utility (**Figure S3**).

The trace quality metric is a relative measure that compares two or more estimated versions of the same signal, extracted using different algorithms, for a cell that is matched across the outputs of the different cell extraction algorithms. To compute this metric, we first matched cells across the outputs of the different algorithms and extracted Ca^2+^ events through exponential deconvolution as described above. To avoid penalizing algorithms that are highly sensitive to weak Ca^2+^ events, we compared the different algorithms using only the highest amplitude Ca^2+^ events that were detected by the algorithms under evaluation, with the number of Ca^2+^ events considered set equal to the smallest number of events found by any of the algorithms.

For each cell, we used a consensus spatial profile, computed as the intersection of the spatial profiles found for this cell by the different algorithms under evaluation. We then converted this consensus spatial footprint into a binary spatial mask. Notably, the use of the consensus spatial profile ensures that comparisons across algorithms are performed across the same set of movie pixels. Using this mask, and the cell’s activity trace estimated by each algorithm, we found the set of time points at which the Ca^2+^ events had amplitudes >90% of the maximum event amplitude, and across these time points we averaged the fluorescence values for each pixel lying within the mask. This yielded an estimated spatial profile characterizing the cell’s activation pattern, calculated separately for each extraction algorithm. Then, we determined the Pearson’s correlation coefficients between the masked portions of the raw or preprocessed Ca^2+^ movie and the corresponding cell spatial profiles, across the set of estimated Ca^2+^ event times. To compute the value of the trace quality metric, we computed the sum of these correlation coefficients, weighted by each the amplitude of each event Ca^2+^ across the set of matched cells.

#### Benchmarking tests with published cell extraction algorithms

Using simulated datasets, we benchmarked the performance of EXTRACT against those of several other cell extraction algorithms. For tests involving the cell extraction routine, CNMF, we used the CNMF code available at the open-source CaImAn-MATLAB Github repository^62^. To evaluate CAIMAN, a Python implementation of CNMF, we used CAIMAN version 1.9.10, downloaded from the open-source CAIMAN Github repository^63^. To evaluate the post-processing tool, SEUDO, we used the published (and currently only) version from the SEUDO Github repository^64^. In some tests, we compared the performance of robust regression to that of non-negative least squares regression, both of which we performed using our implementation of Algorithm 1 (above) in MATLAB. During these tests, we observed that the standard linear regression functions, *lsqlin()* and *lsqnonneg()*, built into MATLAB were very slow. Hence, we wrote our own regression routine in MATLAB (**Supplementary Note 2**), which was up to two orders-of-magnitude faster than MATLAB’s native regression routines. **Supplementary Note 4** has further details about each computational experiment performed to benchmark the different algorithms tested.

Making performance comparisons between different cell extraction algorithms required us to match the cells output by the different algorithms to the ground truth underlying the simulated Ca^2+^ movie. To do this, we wrote custom MATLAB code (provided in the EXTRACT Github) implementing a greedy matching method to identify matched pairs of cell spatial profiles that represent the same actual cell. This method relies on the two-dimensional matrix of Pearson’s correlation coefficients computed between the set of ground truth spatial profiles and the spatial profiles of the detected cells. We greedily determined matched cell pairs by iteratively identifying the cell pair with the highest correlation coefficient, removing the two corresponding spatial profiles from the sets of ground truth and detected cells, and then repeating the process until no there were no more cell pairs with correlation coefficients above a minimum threshold value (typically 0.7). In some cases, we also compared 2 or more different algorithms using real datasets; in these cases, we applied the greedy matching algorithm to all possible pairs of algorithms and then determined the set of cells that were consistently matched across all the algorithms under evaluation.

To tune the hyperparameters of the aforementioned algorithms, we optimized the cell-finding metrics described. Notably, the determination of the PR curve for cell-finding is computationally extensive. Even for a single Ca^2+^ movie, it took about a week to find the PR curve for cell finding for either CAIMAN or EXTRACT, as we tested >1000 hyperparameter combinations for each algorithm (**Figure 3B**). Given this protracted testing duration, we initiated our studies of CAIMAN by starting with hyperparameter values taken from the demos in the CAIMAN Github repository^20^ and then performing a grid-search across 1000 different sets of crucial hyperparameters.

In subsequent computational experiments with EXTRACT, CAIMAN or CNMF (**Table S1**), we started with a near optimal set of hyperparameters as determined via the experiments of **Figure 3B** and then made modest adjustments to the algorithm’s hyperparameter values so that the number of cells identified was roughly the same as the number of cells in the movie. This involved setting the ‘minimum SNR’ hyperparameter values of these algorithms, the total number of cells to be initialized, and hyperparameters characterizing levels of spatial corruption or thresholds of cell size. (For computational experiments with simulated one-photon Ca^2+^ videos, the minimum SNR parameter in CAIMAN was replaced with min_pnr and min_corr, hyperparameters from CNMF-E^23^ to be used with one-photon datasets, and we used CAIMAN’s default value of 1.4 for the ‘ring radius’ hyperparameter.) We also initialized CAIMAN with the movie frame rate and the ground truth decay times of the cells’ optical spike waveforms. CAIMAN’s autoregressive system parameter was set to be p=1, consistent with the data generation process for exponential traces. In principle, these choices gave CAIMAN an unfair advantage over EXTRACT, because CAIMAN was able to model the correct shape of the simulated optical spike waveforms, which EXTRACT does not do. Hence, the superior performance of EXTRACT over that of CAIMAN (**Figures 3, S2, S3**) showcases the power and utility of the robust estimation framework.

For studies of the SEUDO post-processing routine, we first performed a hyperparameter grid search on a set of Ca^2+^ movies; this process took more than a week but identified several options for the hyperparameter set that provided reasonable results, whereas about 60% of the hyperparameter configurations that we tried for SEUDO led to estimated Ca^2+^ traces that were actually worse than those that we input to SEUDO. We therefore identified a hyperparameter configuration that generalized well between different Ca^2+^ movies and levels of signal contamination and then used it for subsequent computational experiments.

### CELL EXTRACTION WITH REAL IMAGING DATA

#### Implementations of CAIMAN, CNMF-E, ICA, and EXTRACT

To run CNMF-E^23^, we used the original author’s implementation, taken from a Github repository called CNMF_E^65^. We based our implementation of CNMF-E on the demo script given in the Github for running it on large data, taking most settings from this script. For both analyses of both striatal and ventral CA1 imaging data, we used gSig = 3, gSiz = 2*gSig, min_pnr=2.5, and min_corr = 0.7. To run CAIMAN, we used the optimal hyperparameter set found via the studies of **Figure 3B**, but with the exception that we used p=2, min_corr = 0.7, min_pnr = 8, ring_size_factor = 1 after performing a local optimization around the default hyperparameter values.

To run PCA/ICA^16^, we used the authors’ published version, which is available on MATLAB’s FileExchange forums. This method first performs a principal components analysis (PCA) to reduce the dimensions of the data and then runs independent components analysis (ICA) to unmix the components spatiotemporally. In all our studies, we ran ICA with µ = 0.1 (which sets the contribution of temporal information in the ICA step), its recommended value in the original paper^16^. We used a maximum of 750 fixed-point iterations for the ICA step. In our studies with simulated data, we set both the number of principal components and the number of independent components to 1.5 times the number of ground truth cells.

We ran EXTRACT mainly using the default values of the hyperparameters, as provided in the EXTRACT Github^59^. However, we optimized a small subset of the hyperparameters that are worth tuning for the analysis of specific datasets, including cell_min_snr, thresholds.T_min_snr, avg_cell_radius, cellfind_max_steps, etc. To test and showcase the utility of robust regression across a variety of analyses, we ran the solver and implementation of robust regression within EXTRACT using a variety of configurations. For the studies of **Figures 4** and **5**, we used adaptive estimation of ***κ*** during the final robust regression (see above). However, using a uniform fixed value of ***κ*** speeds up cell extraction. For the studies of **Figure 6**, we used a fixed κ = **1**. For **Figure 7**, we used adaptive estimation of ***κ*** in the cell finding, refinement and final robust regression stages. To perform the speed benchmarking of **Figure 4**, we allowed both CAIMAN and EXTRACT to temporally downsample the Ca^2+^ movies by a factor of 8, to 3.75 Hz. To ensure fair comparisons, we arranged the numbers of initialized cells to achieve 4500 initialized cells/mm^2^.

#### Detection of Ca^2+^ Transients

To analyze the quality of Ca^2+^ traces estimated for real datasets, we first detected the times of Ca^2+^ transient events within the estimated activity traces. CAIMAN and CNMF-E provide Ca^2+^ event times and amplitudes within their outputs. To find Ca^2+^ events within the traces output by either PCA/ICA or EXTRACT, we used the same exponential deconvolver as used for analyses of simulated datasets as described above. The exponential decay time of the deconvolver was set to approximately match the decay time of the Ca^2+^ indicator used in each real experiment. The thresholds for detection of Ca^2+^ events were 5σ for studies of striatum (**Figure 6**) and 3σ for studies of hippocampus (**Figure 7**), where σ denotes the s.d. of the baseline noise in the deconvolved traces. Below, we provide further details about the analysis of each experimental dataset examined.

### TWO-PHOTON MESOSCOPE IMAGING

#### Movie Acquisition

We recorded movies of Ca^2+^ activity in the V1 cortical area (and adjacent regions) of C57BL/6J mice (Jax: 000664)^66^, genetically modified to express the soma-targeted Ca^2+^ indicator jGCamp8s-Ribo via the CaMk2a promoter in cortical pyramidal neurons (**Figure 4**), using an upgraded version of our two-photon fluorescence mesoscope^1^. This mesoscope, now equipped with faster electronics and more advanced array detectors, employs multiple time-multiplexed illumination beams to scan a 4 mm^2^ area of brain tissue, generating high-resolution two-photon images at 30 frames per second. For this study, the largest Ca^2+^ movie acquired was 800 GB, covering a 4 mm^2^ field of view and consisting of 200,000 frames.

#### Viral expression of Ca^2+^ indicator and surgical preparation

To express jGCamp8s-Ribo in pyramidal cells, we made a viral construct based on the p-AAV-syn-jGCaMP8s-WPRE construct (Addgene plasmid #162374) and the pyc126m ribosome-targeting sequence (Addgene plasmid #158777). The HHMI Janelia Viral Tools facility produced the virus, AAV2/PHP.eB-CamkII-jGCamP8s-Ribo.

To deliver the virus, we injected adult C57BL/6J mice (12–16 week old, male and female) with AAV2/PHP.eB-CamkII-jGCamP8s-Ribo (500 nL of 7E12 GC/mL virus injected at each of 3 sites around the right cortical area V1; injection site coordinates of –3.4 AP, 2.6 ML; –2.7 AP, 2.2 ML; –2.7 AP, 2.9 ML). To do this, we anesthetized mice with isoflurane (4–5% induction, 1–2% maintenance, both in O_2_) and held them in a stereotactic frame for the entire surgery. Body temperature was maintained using a heating pad. We used a SU-P2000 micropipette puller (World Precision Instruments, WPI) to pull a glass capillary (1B100F-4, WPI) and preloaded the virus into the micropipette. We performed a craniotomy centered on the injection coordinates using a round carbide bur (0.7 mm in diameter; Fine Science tools # 19007-07). We infused 500nL of the viral solution at each injection site at a rate of 150 nL/min. We waited for 5 min after each injection and then slowly removed the glass capillary.

Next, we created a cranial window by removing a 5-mm-diameter skull flap (centered at AP –2.5 AP, 2.7 ML) that was positioned over the right visual cortical area V1 and surrounding cortical tissue. We covered the exposed cortical surface with a 5-mm-diameter glass cover slip (#1 thickness, 64-0700, CS-5R, Warner Instruments) that was attached to a circular steel annulus (1 mm thick, 5 mm outer diameter, 4.5 mm inner diameter; #50415K22 McMaster). We secured the annulus to the cranium with ultraviolet-light curable cyanoacrylate glue (Loctite 4305). To enable head-fixation during imaging sessions, we also cemented a metal head plate to the cranium using dental acrylic. Imaging sessions began at 4 weeks after viral infusion.

#### Speed Benchmarking

We processed the mesoscope movies of **Figure 4** with CAIMAN and EXTRACT. With both algorithms, we divided the field-of-view area into spatial patches and performed an 8-fold temporal downsampling for the cell-finding and refinement stages. Both algorithms process the individual patches separately and have internal processes to stitch the results together prior to outputting their final results. When running CAIMAN, we set the width of the spatial patches to be either 80 or 120 (**Figure 4B–D**) pixels wide, with 10 pixels overlap. With EXTRACT (**Figure 4B–E**), we processed most 2 mm × 2 mm movies on a single GPU using either 16 spatial patches (with each patch slightly wider than 256 pixels, accounting for the overlap), or on multiple GPUs using 36 patches (each about 180 pixels in width). To allow fair comparisons with CAIMAN, in **Figure 4C,D** we ran both algorithms on 18 CPU cores; with EXTRACT, we used up to 108 image patches, each about 100 pixels in width. In other cases, the size of the spatial patches used with EXTRACT depended on the RAM requirements. To emulate even larger datasets for the analyses of **Figure 4F,G**, we concatenated two or more portions of this movie in space and time.

### PROCESSING ALLEN DATASET

Ca^2+^ videos from the Allen Brain Observatory (**Figure 5**) were originally 512 × 512 pixels in size and about ∼1 h in duration^67^, but before running EXTRACT we downsampled them to 256 × 256 pixels. We processed 199 movies as a batch, for which the key hyperparameters were tuned on a typical movie. For the cell-finding and cell-refinement stages of EXTRACT, we temporally downsampled the movies by a factor of 6, down to 5Hz, for but not for the final robust regression. We capped the total number of cells EXTRACT that could initialize at either 1000 or 5 times the number of cells found by the Allen SDK^67^, whichever value was smaller.

Given that the Allen SDK uses an unconstrained *L*_2_ regression in its final estimation of Ca^2+^ activity traces, for the sake of making even-handed comparisons we first sought an improved set of traces based on the cellular spatial profiles provided by the SDK. Specifically, we took these spatial profiles and performed a non-negative least-squares (NNLS) regression to obtain a set of estimated Ca^2+^ activity traces. We then compared these traces to those provided by EXTRACT, using robust regression and adaptive ***κ*** estimation. We also took the spatial profiles provided by EXTRACT and similarly performed a NNLS regression. In this way, we were able to isolate the distinct benefits of using robust regression for determinations of cells’ spatial profiles and activity traces (**Figure 5H**). We then calculated trace quality metric values as explained above.

### ANALYSES OF STRIATAL SPINY PROJECTION NEURAL ACTIVITY

For analyses of striatal neural activity, we used published datasets of Ca^2+^ activity in spiny projection neurons, and to compute the spatial coordination index of neural activity we followed closely the approach published in the original paper^40^.

We first computed a matrix of centroid distances between each pair of cells in a movie and detected Ca^2+^ events from the output traces. To make even-handed comparisons between different cell extraction algorithms, we examined an equal number of Ca^2+^ events from the traces output by the different cell extraction algorithms under evaluation. To do this, for each cell we first identified the algorithm that output the smallest number of Ca^2+^ events. We then selected the same number of Ca^2+^ events from the traces provided by the other algorithms for the same cell, with the events chosen to be those with the largest amplitudes. Finally, we created a binarized event trace in which we marked the as ‘active’ the 1-s-period surrounding each Ca^2+^ event. The motivation for this temporal expansion is that it improves the identification of spatially clustered neural activity, which may not be perfectly synchronous, as described previously for these striatal recordings^40^. For each time point, using the centroid distance matrix, we obtained a histogram of pairwise centroid distances for all pairs of active cells at each time point. We also performed the same computations using shuffled versions of the same data in which the cells’ identification numbers were randomly permuted. From these shuffled datasets, we obtained a null distribution by aggregating the histograms of pairwise distances over 100 different permutations. For each time point, we then compared the histogram of pairwise distances for the real data to the null distribution using a one-sample Kolmogorov-Smirnov test with one tail, performed using MATLAB’s *kstest()* function. This allowed us to test statistically whether the pairwise centroid distances in the real data were less than expected by chance. We took the negative base-10 logarithm of the resulting p-value as the spatial coordination metric (SCM). We compared the correlations between SCM values and the mouse’s locomotor speed using the sets of cells that were matched across the outputs of CNMF-E, CAIMAN, ICA, and EXTRACT (**Figure 6F**).

### DETECTION OF DENDRITIC Ca^2+^ ACTIVITY

For the analyses of **Figure S8**, involving dendritic activity in cerebellar Purkinje and neocortical pyramidal neurons^68^, we set the ‘dendritic awareness’ parameter in EXTRACT to 1. The default setting for this parameter is 0; however, when this parameter is set to 1, EXTRACT no longer discards candidate sources of Ca^2+^ activity whose spatial areas or eccentricity values are uncharacteristic of cell bodies. This setting allows EXTRACT to detect Ca^2+^ activity sources, such as dendritic segments, with a wide range of shapes.

The Ca^2+^ imaging data used for **Figure S8C**,**D** were acquired in studies of dendritic excitation in neocortical pyramidal neurons^68^, for which processed data are publicly available^69^. To process the raw Ca^2+^ movies, we first temporally downsampled the Ca^2+^ videos from 31 fps to 7.75 fps and ran EXTRACT on the downsampled movies. For studies of Purkinje neuron dendrites, we first sought to initialize EXTRACT with a reasonable set of candidate dendrites. To determine this set, we first denoised the movie by performing a factor analysis, through a singular value decomposition of the movie. We discarded the noise components of the movie, as determined through the factor analysis, and spatiotemporally smoothed the resultant by convolving the movie with a filter that was 3 time bins in duration and 3 pixels wide in both spatial dimensions. We ran EXTRACT on the denoised, low-pass filtered movie version and used the resulting set of dendritic spatial profiles as the starting point for another iteration of EXTRACT, as performed on a denoised version of the movie that was spatially filtered as before but not temporally smoothed. Within both iterations of EXTRACT, we used the algorithm’s internal Butterworth spatial filtering in the pre-processing module, but with greater filtering along the rostral-caudal dimension then the medial-lateral dimension, to account for the rostral-caudal elongation of the Purkinje cell dendritic trees. After the second iteration of EXTRACT, we visually inspected the results and retained the larger dendritic segments with substantial Ca^2+^ activity.

### ANALYSES OF VENTRAL HIPPOCAMPUS NEURAL ACTIVITY

#### Surgical Procedures

For microendoscopic Ca^2+^ imaging studies of mouse ventral hippocampus, we conducted all surgeries under aseptic conditions using a digital stereotaxic frame (David Kopf Instruments). We used double-transgenic mice (tetO-GCaMP6s-2Niell/J (JAX: 024742)^55^ crossed with Camk2a-tTA-1Mmay/DboJ (JAX: 007004)^70^) that expressed GcaMP6s in hippocampal pyramidal neurons (and in other neural populations). We anesthetized the mice with isoflurane (5% induction, 1–2% maintenance, both in O_2_) in the stereotactic frame for the entire surgery. Body temperature was maintained using a heating pad. A craniotomy centered on the injection coordinates was performed using a trephine drill (1.0 mm in diameter). To prevent increased intracranial pressure due to the insertion of the microendoscope, we aspirated brain tissue until the white fibers of the corpus callosum became visible. Next, we slowly lowered a custom 0.6-mm-diameter microendoscope probe (Grintech GmBH) to the stereotaxic coordinates –3.40mm AP, –3.75mm ML, –3.75mm DV. We fixed the implanted microendoscope to the skull using ultraviolet-light-curable glue (Loctite 4305). To ensure stable attachment of the implant, we inserted two small screws into the skull above the contralateral cerebellum and contralateral sensory cortex (18-8 S/S, Component Supply). We applied Metabond (Parkell) around both screws, the implant and the surrounding cranium. Lastly, we applied dental acrylic cement (Coltene, Whaledent) on top of the Metabond, for the joint purpose of attaching a metal head bar to the cranium and to further stabilize the implant. After surgery, we maintained the animal’s body temperature using a heating pad until it fully recovered from anesthesia.

Mice recovered for 3–6 weeks, at which point we checked the brightness of GCaMP6s expression using a miniature microscope (nVista HD, Inscopix, Inc.). If the expression was sufficiently bright, a baseplate for repetitive mounting of the miniature microscope was fixed onto the skull using blue-light curable composite (Pentron, Flow-It N11VI).

#### Ca^2+^ Imaging Sessions

During imaging studies of ventral CA1 pyramidal neurons, 6 individual mice explored a standard elevated plus maze, comprising an elevated platform (72 cm above the floor) with two opposing open (35 cm × 8 cm), and two opposing closed arms [35 cm × 8cm, wall height: 23 cm] for a duration of 10 min for 4 mice and 20 min for 2 other mice. To start the assay in a uniform manner, we placed each mouse in the center of the platform (8 × 8 cm) facing a closed arm. Ambient illumination in the open arms was 350-400 Lux.

#### Classification of Arm-coding Cells in the Ventral Hippocampus

We wrote custom MATLAB software to determine mouse trajectories within the elevated plus maze, and we manually verified the accuracy of the estimated locations. For each mouse and arm-type of the maze (*viz*., closed *vs*. open), we computed the mean value of each cell’s Ca^2+^ activity trace across all time bins in which the mouse occupied a given arm. For each cell, we determined the difference, *d*, between the mean Ca^2+^ activity levels for when the mouse occupied closed *vs*. open arms of the maze. To perform shuffle controls, we performed the same computations after circularly shifting the Ca^2+^ trace of each cell by a random number of time bins. We computed the activity differences, *d*, between open and closed arms of the maze for 1000 different circular permutations, yielding a null distribution of *d* values for the shuffled datasets. We classified a cell as closed- or open-arm coding, respectively, if its *d* value computed for the real data was at the 95^th^ or higher percentile, or at the 5th or lower percentile, of the *d* values in the null distribution. Cells with *d* values (determined from real data) that fell within the 5%–95% percentile range were classified as non-coding.

#### Decoding of Mouse Locations from the Ca^2+^ Traces of Ventral CA1 Pyramidal Neurons

To quantify the mouse’s position in the elevated plus maze, we divided it into 5 spatial bins: left arm, right arm, upper arm, lower arm and the stem. For our decoding analyses of the neural activity, we first obtained the times and amplitudes of the Ca^2+^ events by applying the deconvolution and event detection routines described above to the traces output by the cell extraction algorithms. We then temporally smoothed the deconvolved activity traces with a 0.5 s running average, to avoid having traces in which many time bins exhibited no Ca^2+^ activity. Ca^2+^ event thresholds were set as noted above.

To create decoders, we trained support vector machines (SVM) to predict the spatial bins of the mouse’s location based on the smoothed Ca^2+^ event traces. We used the *templateLinear()* function in MATLAB with SVM learners, using ridge regularization with the regularization level set through cross-validation studies. To prevent information leakage between the train and test sets due to the smoothing of Ca^2+^ event rates, we selected random time intervals (each 50 time bins = 2.5 s in duration), which in total comprised ∼30% of the full datasets, to be used for decoder testing. We used the remainder of the data for decoder training. In this way, we created 100 different divisions of the data from each imaging session into testing and training subsets, and we averaged the decoding performance across these 100 instances.

## COMPUTER HARDWARE

To perform the benchmarking studies of **Figures 3, S1–S6**, and figures in **Supplementary Note 1** and **3**, we used a combination of several computers as detailed in **Supplementary Note 4**. For the studies of **Figure 4A–D**, we used a desktop computer with two NVIDIA GeForce RTX 3090 GPUs and an Intel Core(TM) i9-10980XE processor with 18 CPU cores. For **Figure 4B**, we ran EXTRACT on Stanford’s High Performance Computing cluster (Sherlock) with 4-8 V100S_PCIE GPUs. For **Figure 4E,H**, we used a desktop computer with an Nvidia Geforce RTX 3090 Ti GPU and an Intel Core i9-9900X Skylake X 10-Core processor. For **Figure 4F,G** and **Figure 5**, we used a desktop computer with two NVIDIA GeForce RTX 3080 Ti GPUs and an Intel Core(TM) i9-10980XE processor with 18 CPU cores. For **Figures 6,7**, and **S7**, we used a desktop computer with an Intel® Xeon(R) CPU E5-2637 v4 @ 3.50GHz × 16 processor and a single NVIDIA GTX 1080 GPU to process EXTRACT and CNMF-E outputs. We processed the CAIMAN outputs of **Figure 6** with an Intel Core i9-9900X Skylake X 10-Core processor. For **Figure S8**, we used a desktop computer with Intel Core i9-9900X Skylake X 10-Core processor and a Nvidia Geforce RTX 2070 GPU.

## DATA AND STATISTICAL ANALYSIS

We performed all data analyses, statistical analyses, and simulations using MATLAB (Mathworks; version R2017b for processing Figures 6 and 7, version R2019a for the rest). All statistical tests were two-sided.

## DATA AND CODE AVAILABILITY

The EXTRACT code is available in an open Github repository^59^. The data are available from the corresponding authors upon request. We used open source software for extracting individual cells and their activity traces from Ca^2+^ videos using principal component and then independent component analysis^71^, constrained non-negative matrix factorization^62,63^, and extended CNMF-E^65^.

## FIGURE CAPTIONS

**Table S1.** List of 33 computational experiments performed to benchmark EXTRACT. To benchmark EXTRACT, we ran 33 separate computational experiments, in which we evaluated the performance of EXTRACT against those of two other widely used cell extraction algorithms, CAIMAN^20^ (both the MATLAB and Python implementations) and PCA-ICA^16^, and a state-of-the-art algorithm for denoising neural Ca^2+^ activity traces, SEUDO^26^. The table contains a description of each experiment, the algorithms compared, and the figure panels. **Appendix Figs. 1-4** are in **Supplementary Note 1**, and **Appendix Figs. 5-10** are in **Supplementary Note 3**.

| No | Experiment description | Figure Panels | Algorithms |
| --- | --- | --- | --- |
| **I. Benchmarks testing the need for the robust estimation framework** |  |  |  |
| 1 | Need for the robust estimation of Ca<sup>2+</sup> activity traces | Fig 1G, Fig S1E | EXTRACT, Inan et al., 2017 |
| 2 | Need for the robust loss function in cell extraction | Fig 3E | EXTRACT |
| 3 | Validation of cell refinement procedures with a robust loss function | Fig S4B | EXTRACT |
| 4 | Faster cell refinement with robust loss function under iid Gaussian noise | Fig A5 | EXTRACT |
| 5 | Need for the robust estimation of cells' spatial profiles and activity baselines | Fig S4C,D | EXTRACT |
| 6 | Changing the robustness parameter during cell finding | Fig S4E | EXTRACT |
| **II. Benchmarking EXTRACT against the state-of-the-art cell extraction algorithms** |  |  |  |
| 7 | Benchmarking EXTRACT against CAIMAN on movies with independently firing neurons | Fig 3A,C,D, Fig S2, S3A | EXTRACT, CAIMAN |
| 8 | Benchmarking EXTRACT against CAIMAN on low SNR movies with independently firing neurons | Fig S3B | EXTRACT, CAIMAN |
| 9 | Benchmarking EXTRACT against CAIMAN on movies with correlated neurons | Fig S3C | EXTRACT, CAIMAN |
| 10 | Benchmarking EXTRACT against CAIMAN on movies with high neuropil contamination and correlated neurons | Fig S3D | EXTRACT, CAIMAN |
| 11 | Benchmarking EXTRACT against CAIMAN on movies with short duration | Fig A8 | EXTRACT, CAIMAN |
| 12 | Hyperparameter search and precision-recall curves | Fig 3B, S4A | EXTRACT, CAIMAN |
| 13 | Suboptimality of averaging pixel activities inside ROIs | Fig A2 | EXTRACT |
| 14 | Benchmarking SEUDO on ground truth cell profiles | Fig A10 | EXTRACT, SEUDO |
| 15 | Benchmarking SEUDO on mismatched cell profiles and varying levels of neuropil contamination | Fig A10 | EXTRACT, SEUDO |
| 16 | Benchmarking SEUDO on mismatched cell profiles with varying levels of profile qualities | Fig A10 | EXTRACT, SEUDO |
| 17 | Comparison of EXTRACT against SEUDO as post-processing tools on movies with varying cell SNRs | Fig 3G,H, S5A,B,C,E,F | EXTRACT, CNMF, SEUDO |
| 18 | Comparison of EXTRACT against SEUDO as post-processing tools on movies with varying cell densities | Fig S5E,F | EXTRACT, CNMF, SEUDO |
| 19 | Comparison of EXTRACT against SEUDO as post-processing tools on movies without a substantial neuropil contamination | Fig S5E,F | EXTRACT, CNMF, SEUDO |
| 20 | Comparisons of MATLAB based cell extraction algorithms on movies with varying SNRs | Fig A9 | EXTRACT, CNMF, ICA |
| 21 | Comparisons of MATLAB based cell extraction algorithms on movies with varying SNRs and donut shaped neurons | Fig A9 | EXTRACT, CNMF, ICA |
| **III. Validation experiments for EXTRACT's constraints and algorithmic choices** |  |  |  |
| 22 | Empirical validation of the theoretical Ca<sup>2+</sup> activity trace estimation errors | Fig A1 | EXTRACT |
| 23 | Empirical validation of the non-negativity constraint | Fig A3 | EXTRACT |
| 24 | Empirical validation of the spatial high-pass filtering | Fig A4 | EXTRACT |
| 25 | Empirical validation of the temporal downsampling during cell finding and cell refinement | Fig S6C | EXTRACT |
| **IV. Benchmarks testing the limits of EXTRACT** |  |  |  |
| 26 | Comparisons with varying cell radii | Fig A6 | EXTRACT, CNMF, ICA |
| 27 | Robustness to increased spatiotemporal correlations | Fig S6A,B | EXTRACT, CAIMAN |
| 28 | Demixing of activities in the extreme limit: cells encapsulated by other cells | Fig 3F | EXTRACT |
| 29 | Testing robust trace estimation under rigid motion with ground truth spatial profiles | Fig S6D | EXTRACT |
| 30 | Testing EXTRACT's cell finding and trace estimation capabilities under rigid motion | Fig A7 | EXTRACT, CNMF |
| **V. Speed benchmarks for the fast solver** |  |  |  |
| 31 | EXTRACT as a fast NNLS solver | Fig S6E | EXTRACT |
| 32 | Scalability of EXTRACT's solver with increasing FOV | Fig S6E | EXTRACT |
| 33 | Benchmarking EXTRACT's speed against SEUDO as a post-processing tool | Fig S5D | EXTRACT, SEUDO |

**Figure S1.**
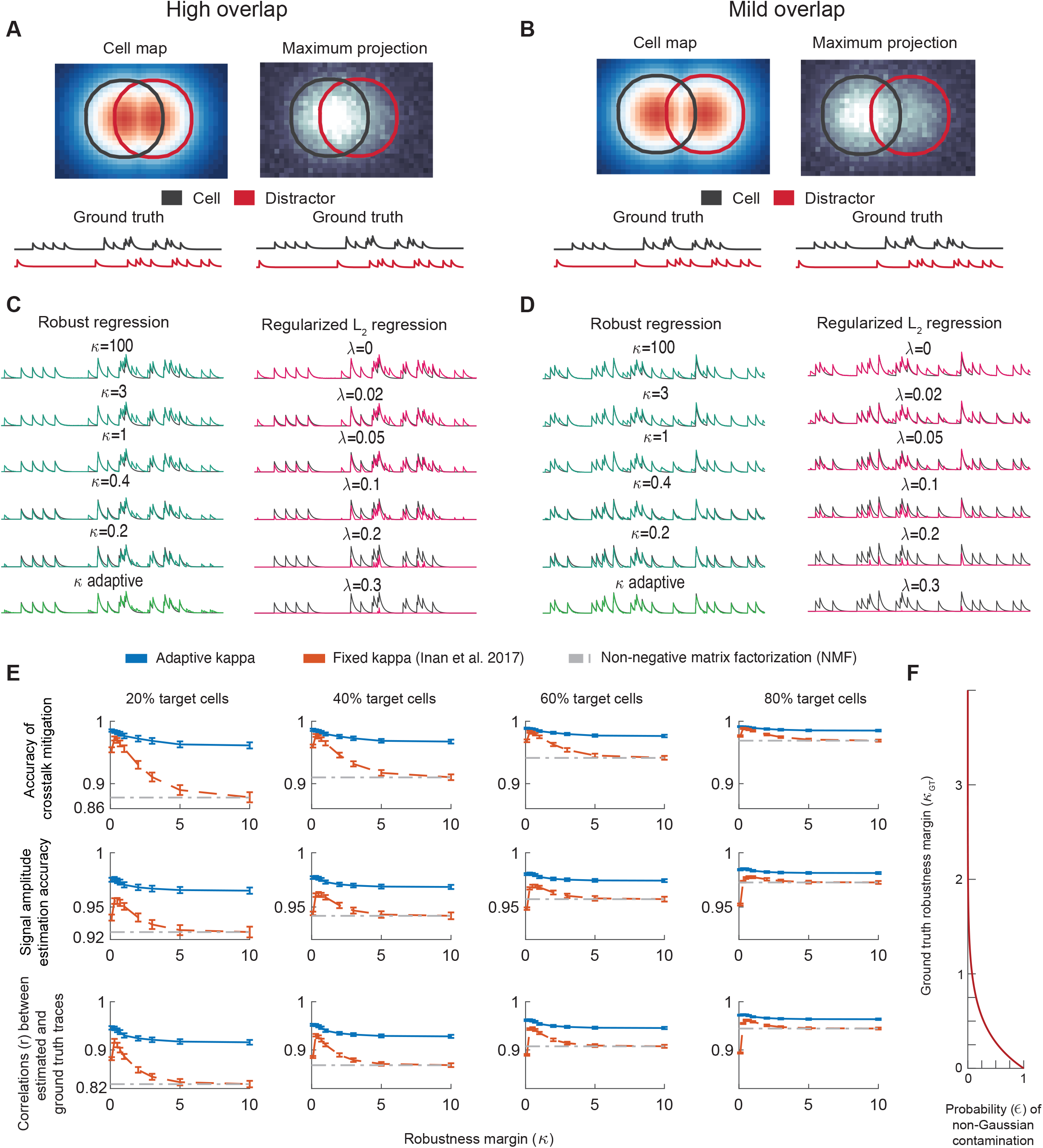
Evaluations of activity trace estimation, using either robust regression with adaptive κ estimation or conventional regression methods. **(A, B)** We simulated the same scenario as in **Fig. 1**, with one neuron of interest (outlined in black) and another overlapping ‘distractor’ neuron (outlined in red). We simulated ground truth Ca^2+^ activity traces for both cells, shown in black and red, respectively, under the assumption that their dynamics were independent and the traces had an SNR value of 5 (**Methods**). Panels **(A)** and **(B)** differ only with regard to the degree of spatial overlap in the two cells’ footprints. **(C, D)** For the two scenarios shown in **(A)** and **(B)**, in panels **(C)** and **(D)**, respectively, we inferred the Ca^2+^ activity trace for the neuron of interest using either robust regression (*green traces*) or *L*_2_-regression with *L*_1_-regularization (*pink traces*). For the robust regression, we systematically varied the robustness margin, κ, which characterizes the expected level of non-Gaussian noise contamination, between 0.2–100 (*top 5 traces*). We also performed a version of robust regression in which the value of κ was estimated in an adaptive manner (*bottom trace*). Smaller values of κ led to reduced Ca^2+^ transient amplitudes in the inferred activity trace; this effect was substantially more pronounced at times when the distractor cell was active. Consequently, reduced values of κ suppressed crosstalk from the distractor more than it suppressed the estimated activity of the cell of interest. Varying κ in an adaptive manner across time yielded the best result, in accord with the fact that the level of crosstalk from the distractor cell also varied across time and was maximal when the distractor cell was most active. For *L*_1_-regularized *L*_2_-regression, we varied the regularization penalty, λ, between 0–0.3. Larger values of λ indiscriminately suppressed the contributions of both the distractor cell and the cell of interest to the inferred activity trace. **(E)** We repeated the analysis of **Fig. 1G**, but with four different relative proportions of target and distractor cells (20%, 40%, 60% and 80% target cells, respectively, for the results shown left to right in the four columns of the figure panel; 600 cells per movie). More distractor cells leads to more (non-Gaussian) crosstalk in the activity traces of the target cells. We evaluated results from robust regression using either fixed^35^ (red curves) or adaptively varying values of ***κ*** (blue curves), and from a traditional non-negative matrix factorization (NMF) of the movie data under an assumption of Gaussian noise (dashed gray curves). For each data point, the *x*-axis value of all plots denotes either the fixed value of κ used (red points) or the κ value used to initialize the adaptive ***κ*** estimation (blue points). We evaluated the algorithms using 3 different performance metrics, the correlation coefficient between the estimated and ground truth activity traces (bottom row), the accuracy of signal amplitude estimation (middle row), and the accuracy of crosstalk mitigation (top row). (The caption to **Fig. 3D** has the definitions of the latter two performance metrics). Because NMF does not involve κ, for each graph the mean performance of NMF is marked with a horizontal line. Robust regression with adaptive ***κ*** estimation led to the most accurate estimation of Ca^2+^ traces, which was nearly insensitive to the initial value of κ in all cases. Error bars: s.d. across 20 simulated movies. **(F)** Plot of the mathematical relationship between the level of non-Gaussian noise contamination, ϵ, and the ground truth value of the robustness margin, κ_GT_, in our robust estimation framework (see **Supplementary Note 2**). When the level of non-Gaussian contamination is near zero, the robustness margin becomes large, implying that regression with our robust loss function behaves similarly to an *L*_2_-estimator. However, when ϵ is high, indicating high levels of non-Gaussian noise contamination, the robustness margin attains low values, skewing the loss function to reject positively valued contaminants.

**Figure S2.**
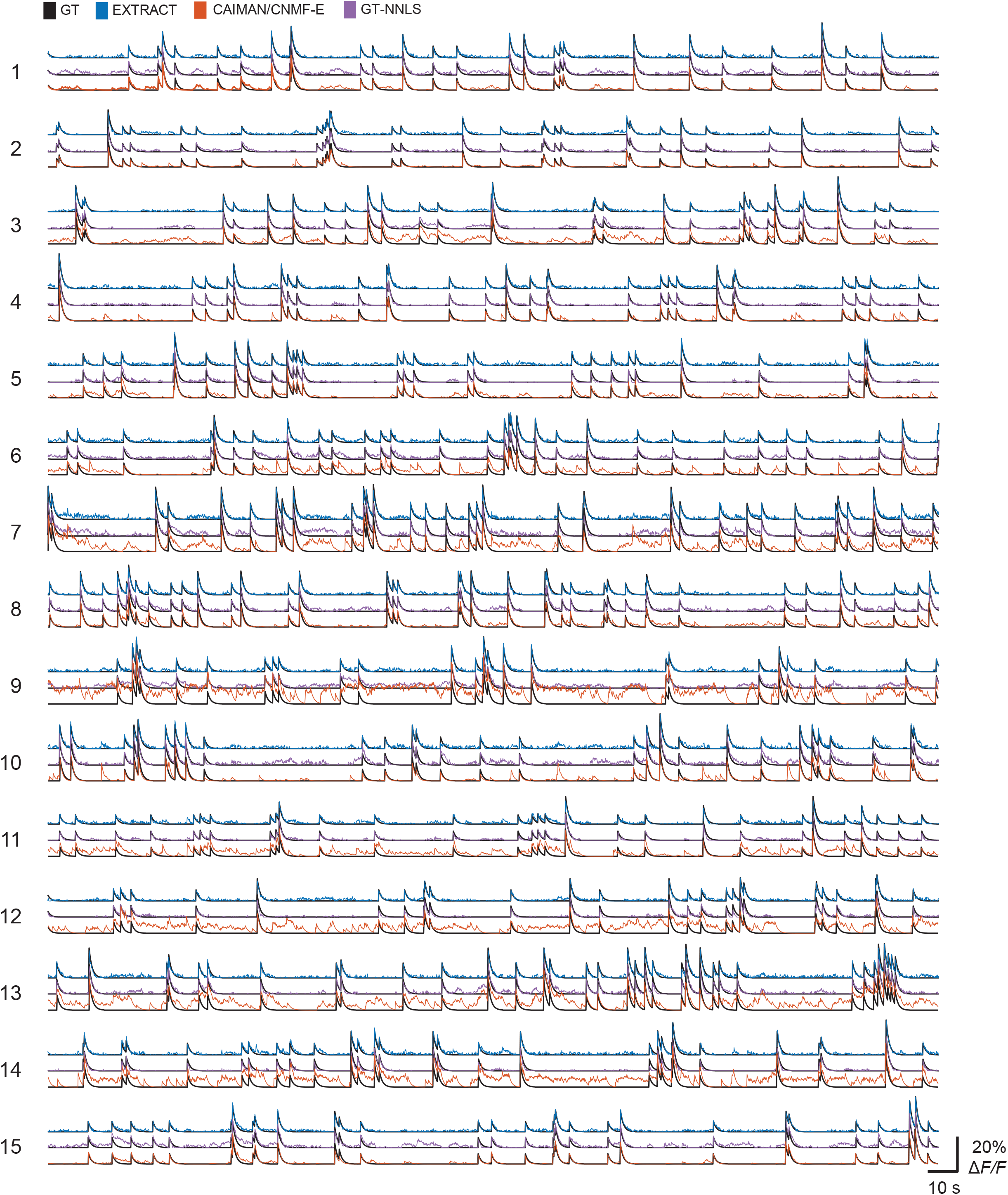
Example sets of estimated Ca^2+^ traces for a simulated one-photon Ca^2+^ video. Using the one-photon Ca^2+^ video of **Fig. 3A** with 600 neurons, we extracted neurons and their activity traces using either EXTRACT, CNMF-E (as implemented in CAIMAN), or a non-negative least squares regression using the cells’ ground truth spatial filters (GT-NNLS). For 15 example neurons, this figure shows the actual, ground truth Ca^2+^ activity traces (black traces) and the 3 different estimated traces (colored traces) for each of the neurons. Cells 1–6 are the same as those in **Fig. 3A**, but plotted over an extended duration. Estimated traces from CNMF-E exhibited a lot of false positive activity, as well as missed Ca^2+^ transients, *i*.*e*., false negative Ca^2+^ activity. For visualization purposes only, we normalized all traces by their peak values to account for differences between the algorithms in how they calculate the amplitudes of Δ*F*/*F* activity. The magnitude of the vertical scale bar refers to the Δ*F*/*F* activity levels in the ground truth traces.

**Figure S3.**
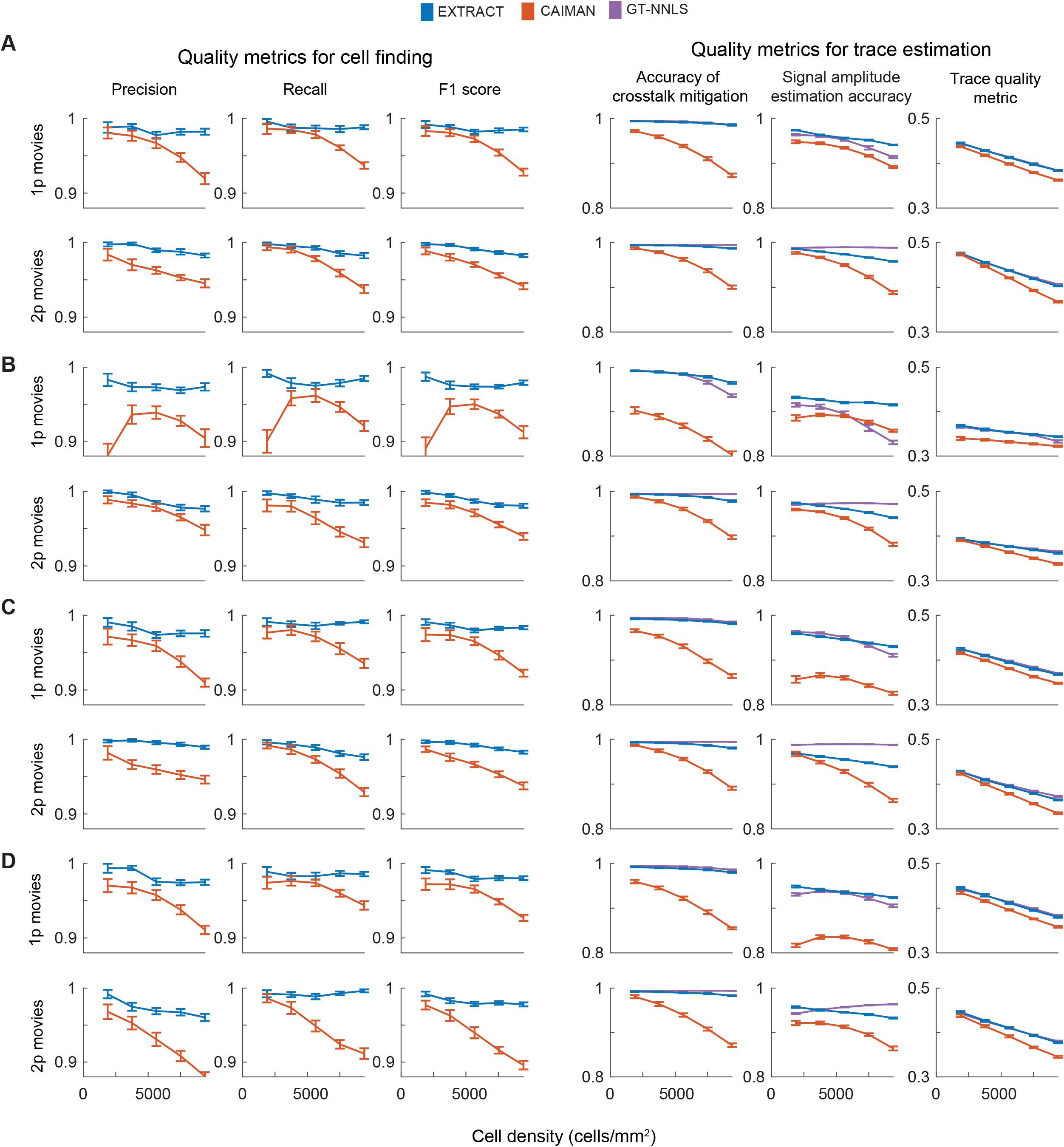
Performance evaluations of cell extraction and trace estimation by EXTRACT and CAIMAN. To compare the performances of EXTRACT (blue curves), CAIMAN (red curves) and non-negative least squares regression using the cells’ actual (ground truth) spatial footprints (GT-NNLS; purple curves), we simulated one- and two-photon Ca^2+^ imaging datasets over a wide range of different conditions. The field-of-view was fixed at 400 μm × 400 μm, but the simulated datasets had varying cell densities, levels of spatially correlated spiking, neuropil contamination levels, and SNR values for the Ca^2+^ activity traces (see **Table S1**). The left three columns of plots show performance metrics for cell finding; since the NNLS approach used the cells’ actual spatial footprints, there were no cell-finding metrics for this method. The right three columns show metrics for the estimation of Ca^2+^ activity traces. In all metrics, EXTRACT outperformed CAIMAN, especially for datasets with high cell densities. See **Fig. 3** and **Methods** for definitions of performance metrics. **(A, B)** Performance metrics for simulated one- and two-photon Ca^2+^ videos with independently spiking cells and 5% correlated noise (see Section *Simulated Ca*^*2+*^ *Imaging Datasets* in **Methods**). The simulated Ca^2+^ transients had a minimum SNR value of 4 in the videos of **(A)** and 2.5 in those of **(B)**. **(C, D)** Performance metrics for simulated one- and two-photon Ca^2+^ videos of neurons with correlated spiking and Ca^2+^ transients with a minimum SNR value of 4. In **(C)**, the level of spatiotemporally correlated noise was 5%, and in **(D)** it was 30%. Error bars: s.d. across 20 simulated movies.

**Figure S4.**
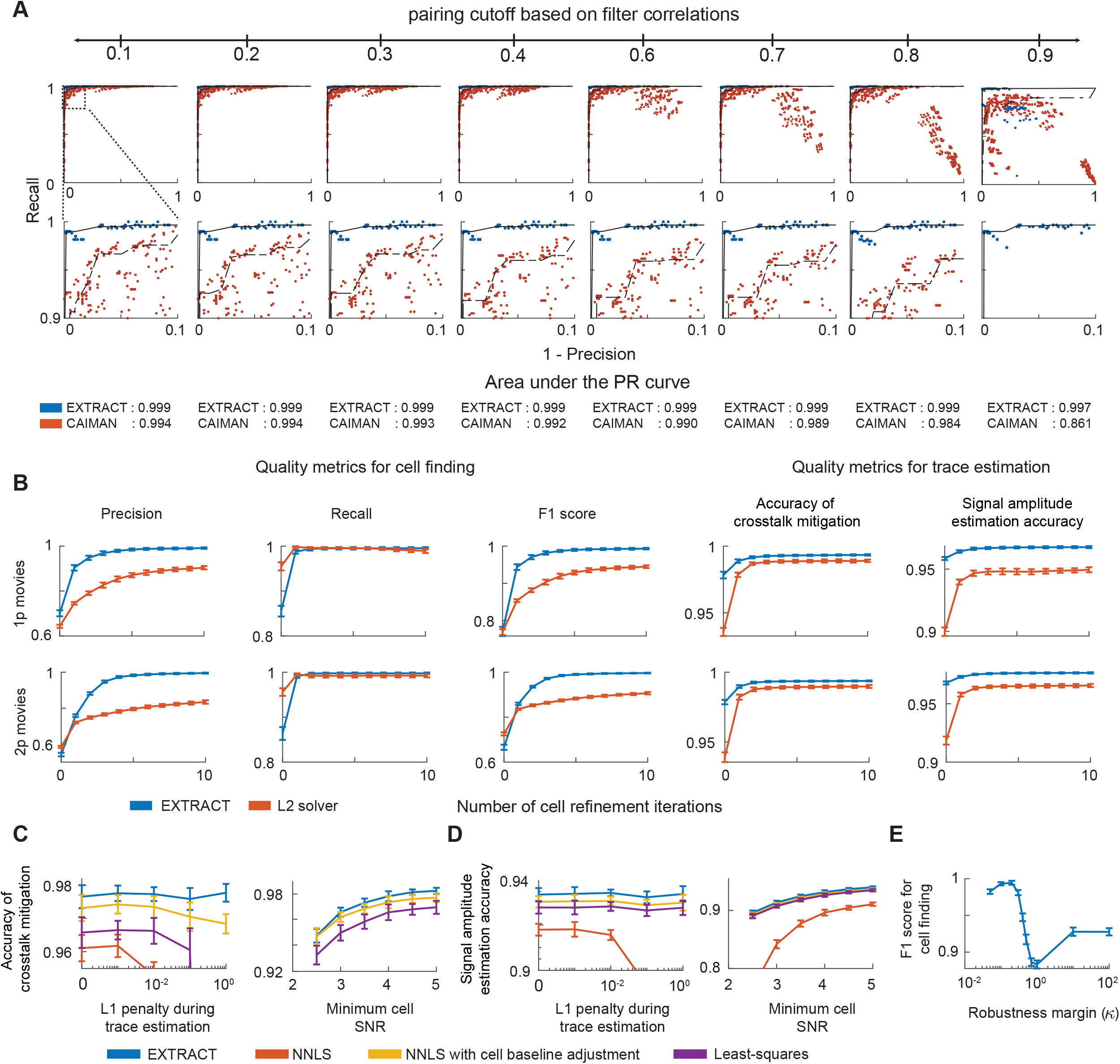
The superior performance of EXTRACT arises from its reliance on robust regression methods, not just a superior software implementation of cell extraction routines. **(A)** Additional results for the computational experiment of **Fig. 3B**, which compares the cell finding capabilities of EXTRACT (blue data) and CAIMAN (red data) via a precision-recall (PR) curve analysis for cell identification while varying each algorithm’s hyperparameters in a nearly exhaustive manner (**Methods**). The top row of plots shows the PR curves across different values of the threshold used to determine when an algorithm has successfully found a cell, as applied to the Pearson’s correlation coefficient between the cell’s estimated and actual spatial footprints. (In **Fig. 3B**, we used a threshold value of 0.5). The bottom row of plots shows a magnified view of the leftmost side of each plot in the top row, showcasing that EXTRACT was nearly impervious to the choice of hyperparameters and consistently yielded higher values of the area under the PR curve (listed for each algorithm beneath each column of plots). **(B)** To test whether the superiority of EXTRACT might arise from a better implementation of the cell extraction pipeline, rather than its use of robust statistics, we compared EXTRACT (blue data) against a conventional *L*_2_ solver (red data) that was implemented identically as EXTRACT but without the use of robust regression (realized in practice by setting κ = 100; **Methods**). We simulated one- and two-photon Ca^2+^ movies with 600 cells and 5% correlated noise, and we examined the performance metric values of EXTRACT and the *L*_2_ solver across iterations of cell refinement, for both cell finding (left three columns) and Ca^2+^ trace estimation (right two columns). Crucially, the use of robust regression consistently led to higher values of all performance metrics and generally reduced the number of iterations needed for convergence. See **Fig. 3** and **Methods** for definitions of the performance metrics. **(C, D)** Plots of crosstalk mitigation accuracy, **(C)**, and signal amplitude estimation accuracy, **(D)**, for 4 different approaches to cell extraction. In all 4 approaches, we first ran EXTRACT to obtain the cells’ spatial filters using its robust regression methods. To find the final set of Ca^2+^ traces, we then performed the final regression using either: EXTRACT with adaptive ***κ*** and baseline estimation (blue data); conventional least squares (*L*_2_) regression (purple data); non-negative least squares (NNLS) regression (implemented in EXTRACT by setting κ = 100; red data); NNLS regression with adaptive baseline estimation (orange data). With all 4 approaches, we used varying levels of *L*_1_ regularization during determination of the cell’s Ca^2+^ activity traces. All raw movie data were pre-processed identically, with identical spatial highpass filtering, prior to the application of the 4 different cell extraction methods. In the left plots of each panel, we varied the value of the *L*_1_ regularization parameter. In the right plots, we varied the minimum SNR value of the cells’ Ca^2+^ transient amplitudes (**Methods**). Higher values of the *L*_1_ regularization parameter lead to sparse assignment of Ca^2+^ events between neighboring cells, which is beneficial for discarding duplicates. NNLS performed the poorest, and its results do not even appear in one of the graphs because they are so low. Whereas, NNLS with adaptive baseline estimation outperformed least-squares, showing the utility of estimating the cells’ baseline activity values separately from the estimation of Ca^2+^ transients. EXTRACT performed best. **(E)** Plot of the F1 score for cell finding, for the simulated one-photon movie datasets of **Fig. 3A**, across varying κ values, without adaptive κ estimation. The graph shows that cellfinding benefits from the use of relatively small κ values, which promotes the discarding of false-positive candidate cells. Error bars: s.d. across 20 simulated movies.

**Figure S5.**
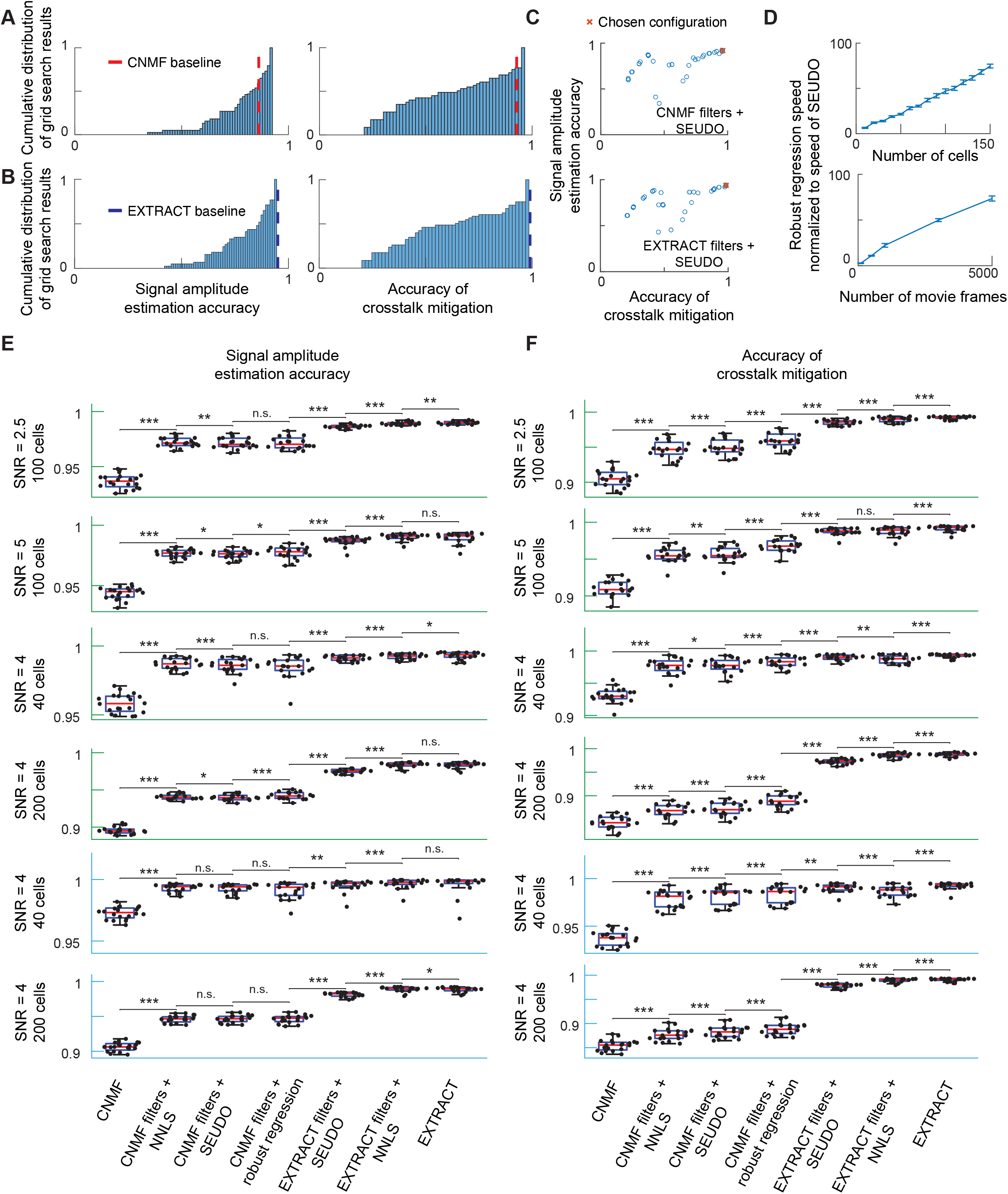
Post-processing Ca^2+^ activity traces with SEUDO did not improve trace estimation quality when cell filters were accurately estimated with robust regression. We evaluated a state-of-the-art approach to post-processing of Ca^2+^ activity traces, SEUDO^26^, by comparing metrics of activity trace quality for Ca^2+^ traces that were estimated in one of seven different ways, by using: (i) EXTRACT; (ii) CNMF; SEUDO using as inputs the cells’ spatial filters provided by (iii) EXTRACT or (iv) CNMF; non-negative least squares regression (NNLS) using cells’ spatial filters provided by (v) EXTRACT or (vi) CNMF; (vii) robust regression using spatial filters provided by CNMF. **(A–C)** To optimize SEUDO, we first simulated a two-photon Ca^2+^ movie with high levels of neuropil contamination (**Supplementary Note 4**) and then extracted cells with CNMF or EXTRACT. Then, we initialized SEUDO with the cell filters from CNMF, **(A)**, and EXTRACT, **(B)**, and performed a grid search across the space of hyperparameters used by SEUDO. The graphs show the cumulative distributions of signal amplitude estimation accuracies (*left plots*) and crosstalk mitigation accuracies (*right plots*) across all sets of SEUDO hyperparameters, starting with spatial filters from CNMF (*top plots*) or EXTRACT (*bottom plots*). The vertical dashed lines in each plot mark the accuracy metric values of CNMF or EXTRACT without using SEUDO. When used with CNMF, SEUDO required fine-tuning of its hyperparameters but was able to improve the estimated Ca^2+^ activity traces. When used with EXTRACT, SEUDO degraded the estimated activity traces in all cases. Panel **C** has scatter plots showing values of the two trace quality metrics for each set of SEUDO hyperparameters, when SEUDO was applied to spatial filters from CNMF (*top*) or EXTRACT (*bottom*). Each datum corresponds to an individual set of hyperparameter values. The data points labeled with a red × mark the SEUDO hyperparameter sets used in **(E), (F)** and in **Fig**. **3G, H** and were picked to maximize the accuracy of crosstalk mitigation while also providing a signal estimation accuracy within 0.5% of this metric’s highest value. **(D)** Plots of the speed of robust regression relative to that of SEUDO, across two-photon Ca^2+^ movies (160 µm × 160 µm field-of-view) of varying densities of target cells (*top*; 1000 movie frames) or duration (*bottom*; 50 target cells). In all cases, the number of distractor cells plus target cells tallied to 150. SEUDO took 1 to 2 orders of magnitude longer to run and scaled far more poorly with large datasets than the fast robust solver we introduced for EXTRACT^35^. For instance, with 150 cells, EXTRACT’s robust solver, as run on 1 GPU, was ∼75 times faster than SEUDO, as run on 18 CPU cores. Error bars: s.d. over 10 movies. **(E, F)** To benchmark the 7 approaches to trace estimation listed above, we simulated two-photon Ca^2+^ movies with varying levels of the activity trace SNR, numbers of cells, and levels of neuropil contamination (**Methods**). Numbers of cells and SNR levels are given in each graph; the graphs with green and blue axes are for movies, respectively, with and without neuropil contamination. Box-and-whisker plots show the values of the trace quality metrics obtained for 20 different movies, denoted by individual data points; red lines denote median values; boxes span the 25th to 75th percentiles; whiskers extend to 1.5 times the interquartile range. Statistical comparisons between algorithms were two-sided Wilcoxon signed-rank tests (*p < 0.05, **p < 10^−2^, ***p < 10^−3^). EXTRACT performed best under all conditions tested when both metrics were evaluated.

**Figure S6.**
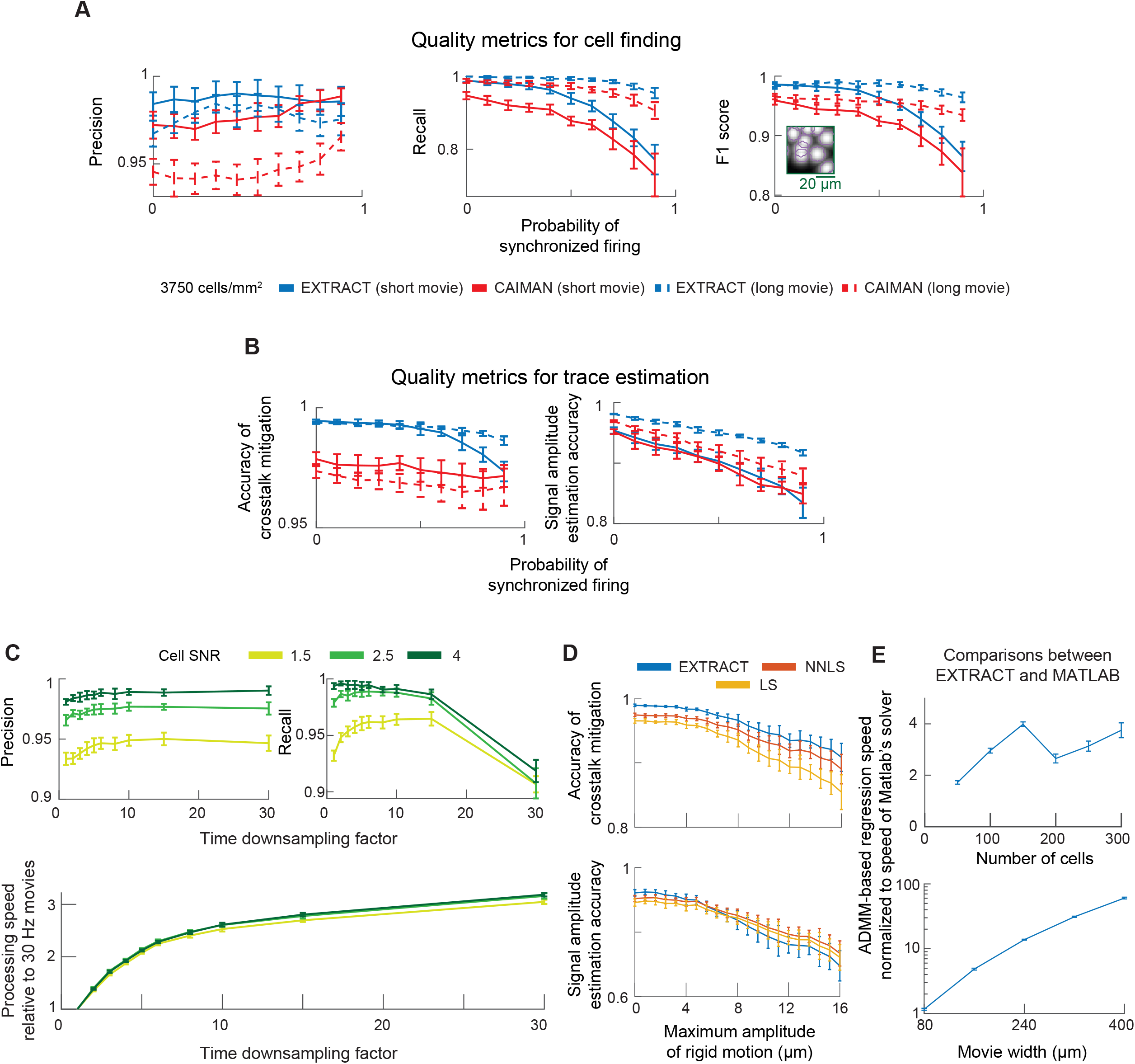
EXTRACT retains high performance across diverse experimental conditions. **(A, B)** We evaluated the capability of EXTRACT to find cells, **A**, and estimate Ca^2+^ traces, **B**, when cells’ activity patterns were spatiotemporally correlated. For this test, we simulated two-photon Ca^2+^ movies (400 μm × 400 μm; 600 cells) with varying levels of synchronized spiking across all cells in the field-of-view (**Methods**). Movies had either 1000 or 5000 image frames (termed short and long movies, respectively). With higher levels of synchronized spiking, the recall metric for cell finding declined, as did the signal amplitude estimation accuracy, although these effects were mitigated for movies of the longer duration. Error bars: s.d. over 20 movies. See **Fig. 3** and **Methods** for definitions of the performance metrics. *Inset*: Cropped maximum projection image showing a typical level of cell overlap at the density level used for these simulation studies (3750 cells/mm^2^) **(C)** We examined the extent to which cell finding by EXTRACT was impervious to a use of temporally downsampled Ca^2+^ movies as inputs. We simulated two-photon Ca^2+^ movies with a 30 Hz frame rate and Ca^2+^ transients with an exponential decay time-constant of 1 s, at 3 different SNR values for the Ca^2+^ traces. We ran EXTRACT on different versions of these movies with varying degrees of temporal downsampling. The top row of graphs shows the precision (*left*) and recall (*right*) of cell finding for downsampling factors ranging from one (no downsampling) to 30 (one sample every second). EXTRACT found cells well with sampling frequencies as low as 2 Hz, which is about the Nyquist frequency for sampling at the Ca^2+^ transient decay rate. The bottom graph shows the processing speed of EXTRACT, normalized to its processing speed for the 30-Hz-movie, as a function of the downsampling factor. Error bars: s.d. over 10 movies. **(D)** To evaluate the extent to which Ca^2+^ trace estimation by EXTRACT was resilient to the presence of brain motion in the Ca^2+^ movies, as manifested by rigid lateral displacements of the field-of-view, we simulated two-photon Ca^2+^ movies with varying degrees of rigid motion (400 µm × 400 µm; 600 cells). Lateral displacements of the specimen comprised a sudden lateral jump in a randomly chosen direction, followed by an exponential relaxation in time back to the original position. Jumps were introduced at a randomly chosen 5% of the movie frames (**Methods**). For each movie, we initialized trace estimators for 80% of the cells in the movie and obtained the estimated Ca^2+^ traces via robust regression implemented in EXTRACT. The graphs show performance metrics for Ca^2+^ trace estimation as a function of the maximum amplitude for the sudden jumps. EXTRACT was resilient to the presence of jumps of ∼6.5 µm, about half the width of a cell, but, when larger jumps were introduced into the Ca^2+^ movies, crosstalk mitigation declined and true Ca^2+^ signals were incorrectly rejected as outliers. Thus, while EXTRACT can accommodate moderate levels of brain motion artifact, we recommend correcting for brain motion by performing image registration across all frames of a Ca^2+^ movie as a pre-processing step before running EXTRACT. Error bars: s.d. over 20 movies. **(E)** To test whether our custom fast ADMM-based solver (**Supplemental Note 2**) provides accurate results even when the density of cells is high (which increases the complexity of the regression due to the presence of highly correlated predictors), we compared the non-negative traces provided by our solver (in the limit κ→∞, practically κ = 100) with those provided by the lsqlin solver of MATLAB (Mathworks). The plots show the speed of our solver relative to that of the MATLAB solver, across Ca^2+^ movies with varying densities of cells (*top*; 160 µm × 160 µm field-of-view) and movie widths (*bottom*; cell density fixed at 3,906 cells/mm^2^). EXTRACT was both faster and scaled better with increasing dataset size. Error bars are s.d. over 20 and 10 movies for the top and bottom graphs, respectively.

**Figure S7.**
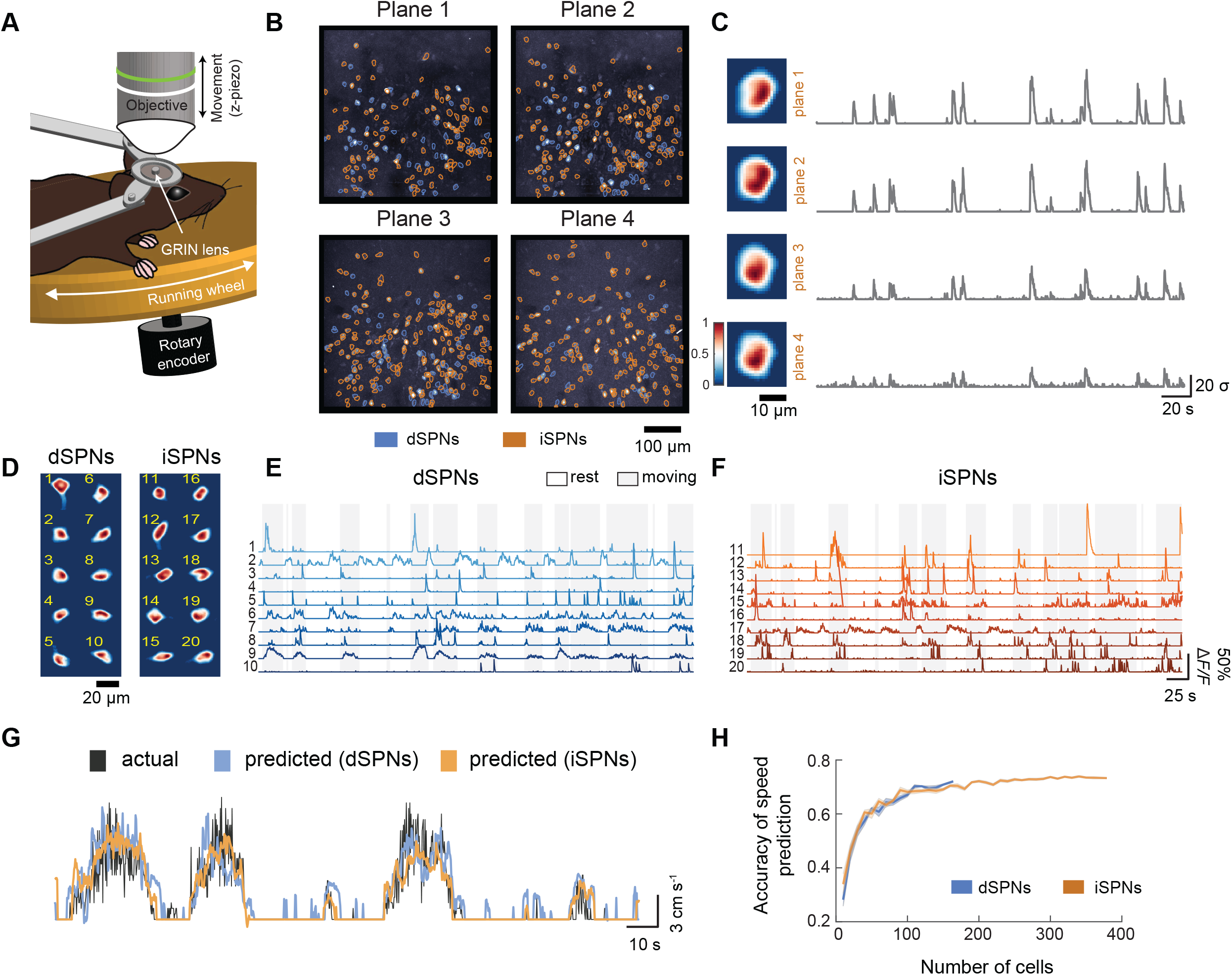
EXTRACT detects spiny projection neurons in multi-plane two-photon imaging data acquired in dorsal striatum. **(A)** We re-analyzed previously published datasets^40^ in which we had used dual-color, multi-plane two-photon imaging to track the Ca^2+^ dynamics of spiny projection neurons of the basal ganglia’s direct and indirect pathways (dSPNs and iSPNs) in the dorsomedial striatum of head-fixed mice at liberty to walk or run on a wheel. Both neuron-types expressed GCaMP6m, but only dSPNs expressed an additional red fluorophore, tdTomato. Within each mouse we sampled SPN Ca^2+^ dynamics within four different optical focal planes spaced 15 μm apart in the axial dimension. **(B)** We ran EXTRACT on the Ca^2+^ imaging data acquired from each of the four different planes. Following cell extraction, we identified each neuron as either an iSPN or a dSPN according to whether the cell expressed tdTomato or not, in addition to GcaMP6m. **(C)** An example cell identified in all four planes. After running EXTRACT, we merged multiple instances of single cells on different planes based on correlations among spatial and temporal components across planes. **(D–F)** The identified components of a representative set of 10 dSPNs and 10 iSPNs. **(D)** Cell images for iSPNs (left) and dSPNs (right). **(E)** Ca^2+^ traces of the dSPNs shown in **(A)** with matching cell IDs. **(F)** Ca^2+^ traces of the iSPNs shown in **(A)** with matching cell IDs. **(G, H)** We used support vector classifiers in conjunction with regularized linear regression to detect movement and predict the locomotor speed simultaneously, using the detected events from the Δ*F/F* traces of the algorithm output. **(G)** When we deployed this method for iSPNs and dSPNs separately, we observed that the estimated locomotor speed tracked very closely the actual speed on a held-out test portion of the data with both populations. **(H)** We quantitatively measured the prediction performance by computing the Pearson’s correlation coefficient between the predicted and the actual locomotor speed on randomly held-out test data over repeated runs. By using either iSPN or dSPN population activity, we could reach reasonably high correlation values, consistent with Ref.^40^.

**Figure S8.**
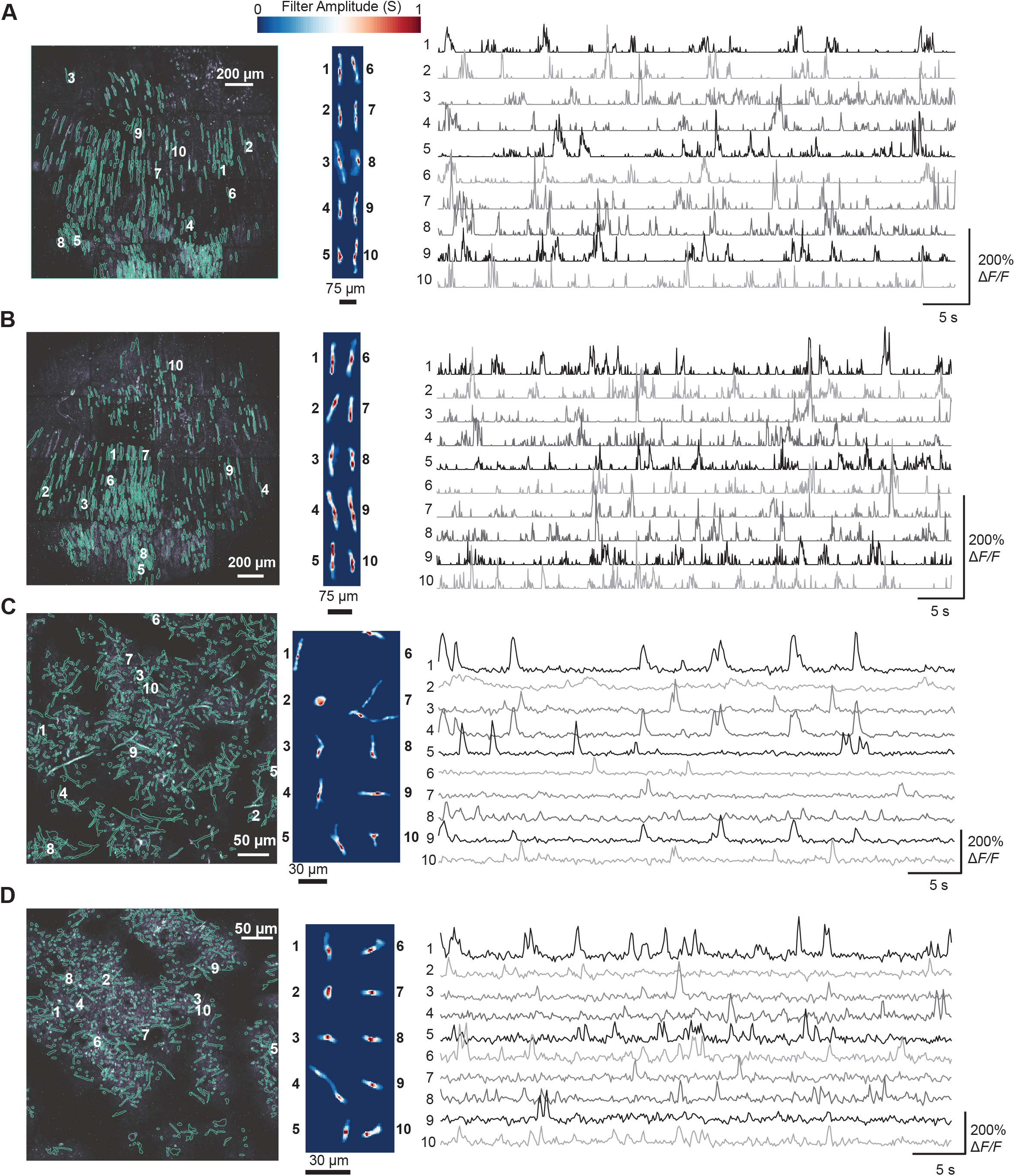
EXTRACT identifies dendritic Ca^2+^ activity in cerebellar Purkinje and neocortical pyramidal neurons in live mice. **(A, B)**. Cell maps of cerebellar Purkinje neuron dendritic trees (*left panels*), along with the extracted spatial forms (*middle panels*) and corresponding Ca^2+^ traces (*right panels*) for 10 example cells in each of two mice, as obtained by applying EXTRACT to Ca^2+^ activity datasets acquired in live mice with a two-photon mesoscope^1^. EXTRACT found the dendritic trees of 507, **(A)**, and 646, **(B)**, Purkinje neurons in the two mice. (**C, D**) Analogous panels to those in **(A), (B)** but for Ca^2+^ videos acquired with a conventional two-photon microscope in the layer 1, apical dendrites of neocortical pyramidal neurons in live mice. EXTRACT found 860 dendritic segments, **(C)**, and 905, **(D)**, dendritic segments for layer 2/3 and layer 5 pyramidal neurons, respectively, in the two mice.

**Figure S9.**
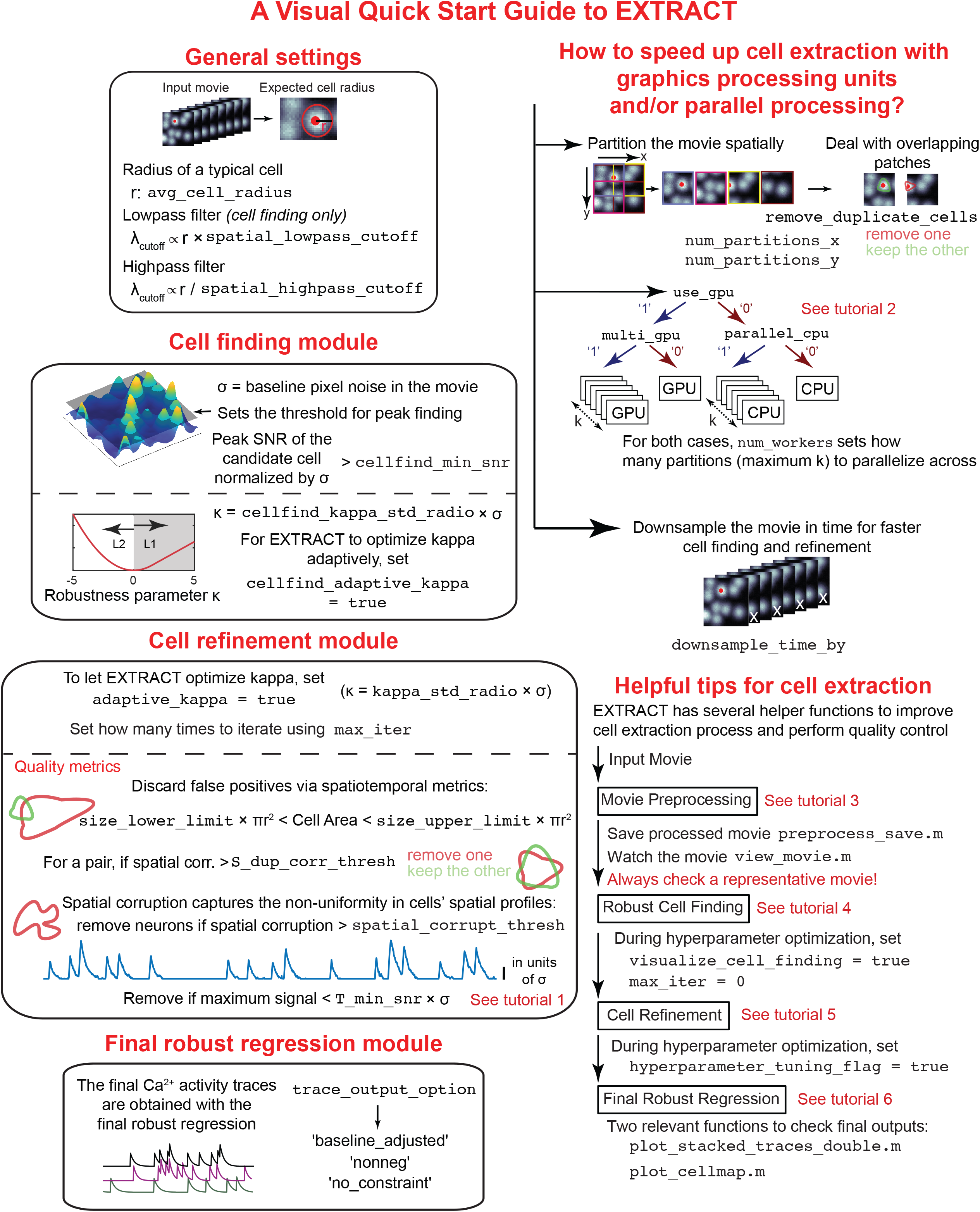
A visual guide to EXTRACT’s internal workings and hyperparameters. We designed six tutorials to help new users adopt EXTRACT, each of which provides a deep look into a distinct aspect of EXTRACT. Besides the cell extraction routine, EXTRACT’s Github repository^59^ includes helper functions to facilitate pre-processing of Ca^2+^ imaging movies and visually inspect the cell extraction results. **Supplementary Note 5**, a detailed user manual, utilizes the six tutorials and discusses the codebase.

