## Supplemental Data 1 for "Fast, scalable, and statistically robust cell extraction from large-scale neural calcium imaging datasets"

This file includes five supplementary notes that provide additional details on various aspects of EXTRACT discussed in the main text.

**Supplementary Note 1: "Cell extraction in estimation theory framework"**

### §1. Overview

Cell extraction is the process of identifying and estimating the spatial profiles ($\boldsymbol{S}$) and activity traces (referred to as $Ca^{2+}$ traces, $\boldsymbol{T}$) of cellular sources in $Ca^{2+}$ imaging movies. The quality of any subsequent data analysis is inextricably linked to the quality of cell extraction, making it a crucial step in research studies utilizing $Ca^{2+}$ imaging movies. Yet, despite the importance of cell extraction, the cell algorithms developed in the field have generally not been placed within the mathematical framework of estimation theory.

To fill this gap, in this supplementary note we first introduce the general mathematical problem of $Ca^{2+}$ movie reconstruction using concepts from estimation theory (**§2**). We then provide some background material regarding the use of non-negative matrix factorization for extracting individual cells and their activity traces from $Ca^{2+}$ imaging movies (**§3**). In **§4**, we describe some of the common $Ca^{2+}$ activity trace estimators used by researchers in the field, and in **§5** we discuss several non-idealities that are regularly encountered in experimental applications and that necessitate the use of robust regression models. In **§6**, we evaluate estimators of neurons' $Ca^{2+}$ activity traces and show that preprocessing the $Ca^{2+}$ movies with a spatial high-pass filter can benefit cell extraction. In **§7**, we review existing approaches to cell extraction using the estimation theory framework introduced earlier. In **§8,** we summarize the key results of this supplementary note, with an eye toward future research on cell extraction algorithms.

### §2. Introduction

To establish a mathematical framework for cell extraction algorithms, we start by discussing three fundamental ingredients within estimation theory[1]:

- A statistical model of the collected data;

- An estimation procedure;
- A loss function that quantifies the performance tradeoffs between different candidate solutions to the estimation problem.

### 2.1. Statistical model

Throughout this note, we will consider $Ca^{2+}$ movies in a flattened (2D) format, $\boldsymbol{M} \in \mathbb{R}^{n_{pixels} \times n_{times}}$, such that $n_{pixels}$ is the number of pixels in the movie and $n_{times}$ is the total number of time bins in the $Ca^{2+}$ movie. We assume that the movie is generated by a general linear model,

$$\boldsymbol{M} = \boldsymbol{S}_{cell}\boldsymbol{T}_{cell} + \boldsymbol{S}_B\boldsymbol{T}_B + \boldsymbol{\sigma} \quad , \tag{1}$$

which includes a noise component $\boldsymbol{\sigma} \in \mathbb{R}^{n_{pixels} \times n_{times}}$ and $n_{roi}$ regions of interest (ROIs), which can include both neurons and background sources of fluorescence modulation, with fluorescence traces $\boldsymbol{T} \in \mathbb{R}^{n_{roi} \times n_{times}}$ and spatial profiles, $\boldsymbol{S} \in \mathbb{R}^{n_{pixels} \times n_{roi}}$. The ROIs comprise two distinct subsets:

i) $n_{cells}$ neuronal cells, $(\boldsymbol{S}_{cell}, \boldsymbol{T}_{cell})$, whose fluorescence $Ca^{2+}$ activity traces are driven by neural action potentials;

ii) background activity sources, $(\boldsymbol{S}_B, \boldsymbol{T}_B)$, such as neuropil $Ca^{2+}$ activity or blood vessels whose pulsations modulate fluorescence $Ca^{2+}$ signals.

Traditionally, the primary objective of the cell extraction process is to accurately estimate the $Ca^{2+}$ activity traces of the cells, $\boldsymbol{T}_{cell}$, whereas background sources may need to be estimated, depending on the algorithm, to decrease the estimation errors on $\boldsymbol{T}_{cell}$.

Assumptions of the model: The linear model described in Eq. (1) is well-established in the existing literature[2–4] and incorporates a key assumption: The $Ca^{2+}$ activity sources are stationary, *i.e.*, any brain motion in the movies should be computationally corrected before the cell extraction process begins. Additionally, cell extraction algorithms often make additional assumptions about the noise component, $\boldsymbol{\sigma}$, which is usually assumed to come from a specific family of statistical distributions. Most commonly[4,5], $\boldsymbol{\sigma}$ is assumed to represent independent and identically distributed (i.i.d.) Gaussian noise with zero mean and unknown variance. But, how realistic are these assumptions?

*Motion correction:* The correction of brain motion across diverse $Ca^{2+}$ movies remains a problem for which there is not yet a fully general solution. However, for $Ca^{2+}$ movies acquired under typical experimental conditions, several algorithms are able to correct the brain's motion to sub-pixel precision[6–8], motivating the choice of stationary $Ca^{2+}$ activity sources in Eq. (1). An alternative approach would be to assume a more complicated model for movie generation, one that involves temporally varying locations for the ROIs, $\boldsymbol{S} = \boldsymbol{S}(t)$. However, this choice would likely complicate the optimization problem by requiring additional assumptions and novel solvers. Thus, in this note, we assume that brain motion has been removed from the $Ca^{2+}$ movies prior to cell extraction.

*Noise distribution:* The desire to model the inference of $\boldsymbol{S}$ and $\boldsymbol{T}$ as a set of regression problems necessitates an assumption of i.i.d. noise within the fluorescence traces of individual movie pixels. Yet, any assumption that the noise is Gaussian is not physiologically justified and may thus be detrimental to accuracy of cell extraction. Fortunately, the mathematical framework of robust estimation allows the assumption of Gaussian noise to be relaxed[9,10]; this separates EXTRACT from the previous literature on cell extraction.

Throughout the remainder of this note, for simplicity of presentation and unless explicitly stated otherwise, we group the background fluorescence sources into the unknown noise contaminant, $\boldsymbol{\sigma} := \boldsymbol{\sigma} + \boldsymbol{S}_B \boldsymbol{T}_B$, such that the ROIs in Eq. (1) explicitly refer to the neurons, *i.e.*, $\boldsymbol{T} := \boldsymbol{T}_{cell}$.

#### 2.2. Estimation procedure

A estimation procedure or estimator is a map, δ, that assigns to the data, *e.g.,* the $Ca^{2+}$ movie, the quantities of interests, *e.g.*, the estimates $\hat{\boldsymbol{T}}$ and $\hat{\boldsymbol{S}}$:

$$\delta : \boldsymbol{\mathcal{X}} \rightarrow \mathbb{D} \quad \Rightarrow \quad \delta(\boldsymbol{M}) = (\hat{\boldsymbol{S}}, \hat{\boldsymbol{T}})$$

Here, $\boldsymbol{\mathcal{X}}$ and $\mathbb{D}$ are data and estimation subspaces, respectively. δ is a map that takes the movie matrix as an input and outputs two matrices, the cells' $Ca^{2+}$ activities and spatial profiles. For example, a simple estimator would involve creating ROIs in $Ca^{2+}$ imaging movies[11,12], which thus serve as estimates, $\hat{\boldsymbol{S}}$, and then computing traces of the spatially averaged fluorescence activity within these ROIs, providing estimates $\hat{\boldsymbol{T}}$. We discuss more sophisticated estimators below.

#### 2.3. Loss function

The loss function, $\mathcal{L}$, quantifies the performance of candidate estimators. It penalizes bad estimations and incentivizes good ones. A (scalar) loss function traditionally has the property:

$$\mathcal{L}(\delta, d) \geq 0, \quad \text{and} \quad \mathcal{L}(\delta, d) = 0 \Leftrightarrow \delta = d,$$

where $d$ is the target with respect to which the estimator, δ, is scored. A simple and widely used example is the $L_2$ loss function, $\mathcal{L}(\delta, d) = (\delta - d)^2$, which panelizes the squared difference between the estimator's prediction and the target. In a more traditional setting, one would define the risk associated with the estimator, $\delta$, as the mean loss averaged across the entire sample space[1]. Here, for simplicity of notation, we do not distinguish between risk and loss.

Given the definition of the loss function and the movie data, $\boldsymbol{M}$, we can place the estimator of the cells' $Ca^{2+}$ activity traces and spatial profiles in a rigorous context by defining:

$$\delta^{*}(\boldsymbol{M}) := (\boldsymbol{S}^{*}, \boldsymbol{T}^{*}) \in \arg\min_{\hat{\boldsymbol{T}}, \hat{\boldsymbol{S}}} \mathcal{L}(\hat{\boldsymbol{T}}(\boldsymbol{M}), \hat{\boldsymbol{S}}(\boldsymbol{M}), \boldsymbol{T}, \boldsymbol{S}) \ .$$

Here, the optimal estimator, $\delta^{*}$, is chosen to minimize the objective function, $\mathcal{L}$, which quantifies the distance between the estimates ($\hat{\boldsymbol{S}}(\boldsymbol{M})$ or $\hat{\boldsymbol{T}}(\boldsymbol{M})$) and their ground truth values ($\boldsymbol{S}$ or $\boldsymbol{T}$ from Eq. (1)). If a minimum for the loss function exists, the returned pair, $(\boldsymbol{S}^{*}, \boldsymbol{T}^{*})$, is considered to be the optimal solution to the estimation problem.

#### 2.4. Unique challenges of cell extraction

Framing cell extraction as an optimization problem involves two unique challenges, which necessitate incorporation of domain knowledge into the solution of the problem: i) The existence of multiple outputs or objectives, $(\boldsymbol{S}^{*}, \boldsymbol{T}^{*})$, that should be accurately estimated, and ii) the number of movie pixels or 'samples' that can be used to estimate each cell's $Ca^{2+}$ activity trace is the number of pixels occupied by the cell, which is typically a relatively small number. Therefore, in the context of real experiments, for cell extraction algorithms to succeed they must generally perform well in a regime in which it is infeasible to estimate well both $\boldsymbol{S}^{*}$ and $\boldsymbol{T}^{*}$ and in which there are relatively few data samples available to constrain the optimization; these two challenges typically make data-hungry approaches ill-suited for cell extraction.

*Multiple Objectives:* In general, when the optimization problem has multiple objectives, it is often not possible to find a single solution that constitutes a global minimum for the multi-objective loss function. Instead, one usually searches for a particular Pareto optimum, which generalizes beyond the set of samples used for the optimization. A common example arises in the case of regularization in neural networks, in which a secondary objective function, *e.g.*, the norm of the parameters, should be minimized in addition to the primary objective, *e.g.*, the loss function of the task. In this scenario, a regularization parameter, usually denoted by $\lambda$, defines how much value to assign to the secondary objective relative to the primary one. In the estimation theory framework, picking a specific $\lambda$ corresponds to picking the Pareto optimum that one desires the network to find, which is often defined via a cross-validation procedure. However, unlike with neural network training, for the estimation of $Ca^{2+}$ activity traces we lack access to the ground truth, and, moreover, there are no training samples from which we expect the algorithm to generalize. Instead, the optimization algorithm has only one chance to estimate accurate $Ca^{2+}$ activity traces, making cross-validation infeasible for deciding which objectives are more important than others. Instead, one needs to rely on theoretical and experimental domain knowledge, as well as simulations with reproducible and known conditions.

One argument in the existing literature assigns all the emphasis of the loss function to the accurate estimation of $\boldsymbol{T}$. These post-processing approaches[13–15] often utilize cells spatial profiles, $\hat{\boldsymbol{S}}$, extracted by other algorithms and design optimization approaches that only aim to estimate $\boldsymbol{T}$ accurately. However, in practice, incorrect estimation of $\hat{\boldsymbol{S}}$ often leads to overfitting for the estimation of $Ca^{2+}$ activity traces, as picking the correct pixels, *i.e.*, estimating $\hat{\boldsymbol{S}}$, from which we aim to extract the cells' $Ca^{2+}$ activity traces is crucial to the extraction of cells' $Ca^{2+}$ activity traces. In other words, the quality of the data used for the estimation is at least as important as the estimation algorithm, if not more.

In our extensive benchmarks, discussed in the main text (**Fig. 3E,F**), we confirmed this theoretical prediction. SEUDO[13], a post-processing algorithm applied after $\hat{\boldsymbol{S}}$ are estimated by CAIMAN[5], led to inferior traces compared to the least-squares estimates obtained from cells' spatial profiles found by the robust solvers of EXTRACT. Therefore, the solution to the complex challenges put forth by the existence of multiple objectives may not be as simple as ignoring half of the problem, *i.e.*, the estimation of cells’ spatial profiles. Thus, cell extraction algorithms aiming to provide an optimal solution to the cell extraction should likely refrain from considering the estimation of both $\hat{\boldsymbol{S}}$ and $\hat{\boldsymbol{T}}$ as separate problems. Instead, as we discuss in Section 2.5 below, experimental domain knowledge should guide the design process of the multiobjective optimization.

*Sparse data regime:* In a classical estimation problem, one often defines a class for the estimators of interests and proves desirable quantities, such as unbiasedness, that hold in the limit of large sample size. However, in $Ca^{2+}$ imaging movies, especially those with a large field of view[16], each cell occupies roughly tens of pixels. Thus, estimating cells’ $Ca^{2+}$ activity traces is a non-traditional problem, one with a very sparse number of samples.

To appreciate how having a low number of samples can pose a problem, consider fitting a line using only a few noisy data points. Even when the data points were to be sampled from a linear model, the resulting line would likely not be a faithful representation of the ground truth. Yet, if the number of data points were to increase, the estimation error would decrease. Owing to the experimental conditions with which the $Ca^{2+}$ imaging movies are sampled, the trace estimation problem inevitably lies in the sparse data regime, in which only few data points constrain the model training. Consequently, what cannot be constrained by the data should be guided by the inductive biases, which follow from years of experimental domain knowledge. Fortunately, as we discuss next, systems neuroscience has been building such domain knowledge over several decades, culminating towards a simple optimization paradigm.

#### 2.5. Movie reconstruction paradigm

As we discussed in the previous section, overfitting can easily become a severe issue when the number of samples is low, which in turn requires the application of inductive biases in every step of the cell extraction. To achieve this, the statistical model defined in Eq. (1) presents a straightforward path: Reconstruct a denoised movie of the form $\hat{\boldsymbol{M}} = \hat{\boldsymbol{S}}\hat{\boldsymbol{T}}$, which contains only the cells' $Ca^{2+}$ fluorescence, not the noise components, and minimize the reconstruction error:

$$\delta^{*}(\boldsymbol{M}) := (\boldsymbol{T}^{*}, \boldsymbol{S}^{*}) = \arg\min_{\hat{\boldsymbol{S}},\hat{\boldsymbol{T}}} \mathcal{L}(\boldsymbol{M}, \hat{\boldsymbol{M}}), \tag{2}$$

where $\mathcal{L}(\boldsymbol{M}, \hat{\boldsymbol{M}})$ is a scalar loss, constrained to consider estimators of the form $\hat{\boldsymbol{M}} = \hat{\boldsymbol{S}}\,\hat{\boldsymbol{T}}$. We define this surrogate problem as the '**movie reconstruction**' paradigm, which constitutes the starting point of the several established cell extraction algorithms[2–4], as well as EXTRACT.

It is important to note that the movie reconstruction paradigm is, although a clever trick, a surrogate problem, not the primary objective. Since we do not have direct measurements of the target quantities, *i.e.*, $\boldsymbol{S}$ and $\boldsymbol{T}$, movie reconstruction presents itself as the next best approach that is analytically and numerically tractable and has shown empirical success. However, minimizing the surrogate loss function associated with the movie reconstruction at all costs and exactly, as is traditionally done in the literature[2–5,13], is often not the optimal strategy. A simplest example is the assumption of non-negativity for the $Ca^{2+}$ activity traces, which is regularly assumed in the previous literature. This assumption would inevitably increase the error on the movie reconstruction (See Section **6.2.** and **Appendix Figure 1** below), but is still desired to accurately estimate the $Ca^{2+}$ activity traces[5].

For the rest of this note, we aim to unify the existing cell extraction heuristics via the estimation theory framework and provide supporting evidence for (or lack thereof) the common knowledge methods developed in the field.

### §3. Past approaches to the cell extraction problem

In this section, we summarize the history of cell extraction within the estimation theory framework. To do so, we start by defining the most common loss function, the $L_2$ loss function, used in Eq. (2) to quantify the error in the movie reconstruction:

$$\mathcal{L}_2(\boldsymbol{M}, \hat{\boldsymbol{M}}) = \sum_{i=1}^{n_{pixels}} \sum_{j=1}^{n_{times}} (\boldsymbol{M}_{ij} - \hat{\boldsymbol{S}}_i \hat{\boldsymbol{T}}_j)^2 = ||\boldsymbol{M} - \hat{\boldsymbol{S}}\hat{\boldsymbol{T}}||_F^2,$$

where $||\,.\,||_F$ stands for the Frobenius norm of a matrix.

Without any further assumption, minimizing $\mathcal{L}_2(\boldsymbol{M}, \hat{\boldsymbol{M}})$ has infinitely many solutions for $\hat{\boldsymbol{S}}$ and $\hat{\boldsymbol{T}}$, all of which equivalently lead to $\hat{\boldsymbol{M}} = \boldsymbol{M}$, a tautology. Consequently, further biologically relevant assumptions are required to constrain this optimization problem into a useful format. To this end, one relevant assumption is to impose a maximum rank on the reconstruction, which corresponds to the number of neurons in the movie. This leads to the modified optimization problem:

$$\begin{aligned} &\text{Minimize} && \mathcal{L}_2(\boldsymbol{M}, \hat{\boldsymbol{M}}) \\ &\text{subject to} && \text{rank}(\hat{\boldsymbol{M}}) \le n_{cell} \end{aligned} \tag{3}$$

The solution to this problem is provided by the Eckart–Young–Mirsky theorem[17], in which one takes the singular value decomposition (SVD) of the matrix $\boldsymbol{M}$ and utilizes the first $n_{cell}$ modes with highest singular values to obtain $\hat{\boldsymbol{M}}$. In this problem, the low-rank matrices reconstructing $\hat{\boldsymbol{M}}$ would represent the cells' spatial profiles and $Ca^{2+}$ activity traces.

There are several short-comings with the approach in Eq. (3). First, plain SVD can assign negative weights for the spatial profiles, which are assumed to be non-negative by design. Second, there are no theoretical or empirical guarantees that using modes with highest singular values would preferentially pick cell activities over global modes of spatiotemporally correlated noise, *e.g.*, neuropil. Due to these reasons, while SVD constitutes a first step, it is not suitable for extracting neuronal signals from $Ca^{2+}$ imaging movies. Yet, the idea of constraining the rank of the movie plays a central role in the cell extraction algorithms.

#### 3.1. Non-negative matrix factorization

A more refined version of the matrix factorization approach in Eq. (3) includes enforcing non-negativity constraints on both low-rank reconstruction matrices, which was formulated in the literature as follows[18]:

$$\begin{aligned} &\text{Minimize} \quad \mathcal{L}_2(\boldsymbol{M}, \hat{\boldsymbol{M}}) \\ &\text{subject to} \quad \text{rank}(\hat{\boldsymbol{M}}) \le n_{cell}, \quad \hat{\boldsymbol{T}} \ge 0, \quad \hat{\boldsymbol{S}} \ge 0. \end{aligned} \tag{4}$$

Due to the existence of non-negativity constraints, this formulation of the cell extraction can no longer be solved through an SVD approach. Instead, the observation that the loss function is bi-convex in $\hat{\boldsymbol{S}}$ and $\hat{\boldsymbol{T}}$ allows one to take an alternating minimization approach. Specifically, non-negative least-squares estimation is applied to estimate one while holding the other fixed, and vice versa. The process alternates until convergence. This algorithm, called the non-negative matrix factorization[18] (NMF), addresses the first issue raised above, *i.e.*, the non-negativity of the spatial profiles (and $Ca^{2+}$ activity traces), but not the second one. Particularly, there is still no constraint that incentivizes the extraction of cells, rather than neuropil or other globally correlated signals.

#### 3.2. Initialization for the non-negative matrix factorization

NMF has been the basis for the current state-of-the-art cell extraction algorithms. As noted in the previous section, however, one aspect of the cell extraction problem remains: How to pick cells preferentially compared to other spatiotemporally correlated components?

The key insight lies in the realization that the optimization problem in Eq. (4) is non-convex. Therefore, the initialization plays a crucial role in whether the algorithm converges to a ``good" or ``bad" minimum of the loss function, which puts further emphasis on the initialization techniques. For example, one can develop an initialization routine for the spatial profiles, $\hat{\boldsymbol{S}}$, such that they primarily encapsulate cells, not undesired components such as neuropil, which has been one of the major contributions of the follow-up work[2] to NMF[18]. Supplied with further assumptions that discourage the training to ``jump out" of the good local minima, a good initialization can then lead to desirable cell extraction results. But, how do we enforce that the problem remains within this 'good' local minimum?

One approach[2,4] has been to enforce the locality and sparsity for $\hat{\boldsymbol{S}}$, which prevents the spatial profiles from indefinitely growing due to correlated global background activities. Then, the (modified) NMF problem becomes:

$$\begin{aligned} &\text{Minimize} \quad \mathcal{L}_2(\boldsymbol{M}, \hat{\boldsymbol{M}}), \quad \text{with an initialization routine for } \hat{\boldsymbol{S}} \\ &\text{subject to} \quad \text{rank}(\hat{\boldsymbol{M}}) \leq n_{cell}, \quad \hat{\boldsymbol{T}} \geq 0, \quad \hat{\boldsymbol{S}} \geq 0, \quad \forall_{j \in cells} \forall_{i \notin O(j)} \hat{\boldsymbol{S}}_{ij} = 0, \end{aligned} \tag{5}$$

where $O(j)$ stands for the region of support for the cell $j$, whose definition depends on the specific algorithm. In general, the region of support is chosen to be local and sparse, in line with our discussion above.

The solution of the Eq. (5) follows a two step process, one following the other:

1) Cell finding: This is an initialization procedure, often performed in a greedy manner[2], for finding the approximate locations of the cells. It ensures that the problem is initialized to be within the vicinity of a ``good" local minimum.

2) Matrix decomposition: This is solved via the alternating non-negative least squares, similar to NMF above. The initialization is provided by the previous step, and additional constraints on the locality and sparsity of $\hat{\boldsymbol{S}}$ are enforced.

Due to potential imperfections in the cell finding procedures, which may lead to non-neuronal and/or duplicate components, the problem described in Eq. (5) is often empirically suboptimal. Specifically, it misses a step that discards garbage components that might arise during the cell extraction. That being said, one can introduce an interesting theoretical concept via Eq. (5). In this minimization problem, replacing $\hat{\boldsymbol{S}}$ with the ground truth filters $\boldsymbol{S}$ for the estimation of traces, one can define an ``idealized $L_2$ solver". This can be considered as an upper bound on the $L_2$ movie reconstruction paradigm without additional background modeling, which is what we used in the main text (**Figs. 3** and **S3**) to benchmark EXTRACT.

#### 3.3. Constrained non-negative matrix factorization

Modified NMF and idealized $L_2$ solver present educational starting points for creating a practical pipeline for cell extraction. Particularly, after incorporating cell refinement procedures that discard spurious components that do not belong to cells, we arrive at a practical solver for an arbitrary loss function $\mathcal{L}(\boldsymbol{M}, \hat{\boldsymbol{M}})$:

For a fixed number of iterations

Minimize $\mathcal{L}(\boldsymbol{M}, \hat{\boldsymbol{M}})$, with an initialization routine for $\hat{\boldsymbol{S}}$

subject to $\text{rank}(\hat{\boldsymbol{M}}) \le n_{cell}, \quad \hat{\boldsymbol{T}} \ge 0, \quad \hat{\boldsymbol{S}} \ge 0, \quad \forall_{j \in cells} \forall_{i \notin O(j)} \hat{\boldsymbol{S}}_{ij} = 0$ (6)

Delete spurious cells and repeat

Run one final regression with resulting $\hat{\boldsymbol{S}}$ to obtain the $Ca^{2+}$ activity traces $\hat{\boldsymbol{T}}$

We refer to the solution of the optimization problem in equation (6) under the $L_2$ loss function, $\mathcal{L}_2$, as the ``$L_2$ solver," which consists of three steps:

- Cell finding: Similar to the modified NMF, this step provides good initial guesses for where cells are located and initializes $\hat{\boldsymbol{S}}$.

- Cell refinement: Through iterative application of alternating optimization, followed by the deletion of spurious cells, cell filters in $\hat{\boldsymbol{S}}$ are refined and spurious cells in $\hat{\boldsymbol{S}}$ are removed.
- Final regression: Once the cell filters $\hat{\boldsymbol{S}}$ are refined, one final regression is applied to obtain cells' $Ca^{2+}$ traces, $\hat{\boldsymbol{T}}$.

The $L_2$ solver introduced in Eq. (6) does not contain any assumptions on the trace dynamics and performs the cell refinement process iteratively, in which garbage removal is performed gradually instead of all at once. A version of this approach, with an exponential decay assumption on the calcium trace dynamics $\hat{\boldsymbol{T}}$ and a single garbage removal step, is first introduced in Ref.[2] as the ``constrained non-negative matrix factorization" (CNMF) and later applied in several other pipelines[3–5]. Unlike the NMF[18] in Eq. (4) we started with, the $L_2$ solver (and relatedly CNMF) framework in Eq. (6) has three main advantages:

- The initialization procedure via the cell finding module ensures that the problem starts out close to a ``good" local minimum.
- The locality constraint, on top of the good initialization, preferentially picks and retains $Ca^{2+}$ activity sources that putatively resemble cells over other forms of global spatiotemporal $Ca^{2+}$ activity sources.
- The fact that spurious cells are deleted through the refinement procedures desensitizes the dependence on specific parameters, such as the initial rank, $n_{cells}$, and increases the accuracy of the cell extraction outputs.

It has been empirically observed over decades of work in the field of calcium imaging that (C)NMF framework provides a good starting point for cell extraction. Even trace post-processing approaches such as SEUDO[13] and FISSA[14] use a version of this framework, with additional background modeling, to accurately estimate cells' $Ca^{2+}$ activity traces.

Before we conclude our step-by-step historical account, leading up to the prior state-of-the-art cell extraction routines, *i.e.,* CNMF, we discuss one final aspect of cell extraction that has perhaps not been given due attention before. As we initially argued, one should not forget that CNMF framework aims to solve not the primary objective of cell extraction, but a surrogate problem, *i.e.*, the movie reconstruction, and with several simplifying assumptions. Therefore, as is often the case with surrogate problems, exact minimization of the loss function can lead to overfitting errors. As we show in our simulation benchmarks (See **Table S1** and **Figs. 3, S3-6**), some of the assumptions made by CNMF, *e.g.*, the exponential decay of the calcium signals and background modeling, may not be well justified due to the overfitting issues, even for movies that perfectly match these assumptions. Specifically, performing non-negative least-squares estimates to obtain the $Ca^{2+}$ activity traces provided higher quality $Ca^{2+}$ activity traces compared to the raw CNMF outputs (**Figs. 3E, 3F, S5E,** and **S5F**), in which the latter incorporated both the background modeling and exponentially decaying kernels.

Inspired by this observation, to develop further algorithms of cell extraction, we opted to use the $L_2$ solver (also introduced in this work, as an improvement over existing approaches) as an algorithmic basis, rather than the CNMF framework, when developing the EXTRACT software introduced in this work.

#### 3.4. Cell extraction with EXTRACT

The majority of cell extraction algorithms, if not all, focused on minimizing the $L_2$ loss on the movie reconstruction. However, as we discussed above, solving a surrogate objective exactly may lead to overfitting problems, even when the data generation process of the $Ca^{2+}$ imaging movies match the underlying assumptions. Instead, minimizing loss functions that have theoretically proven robustness guarantees has yielded empirical success over the last century[9]. When designing EXTRACT, we have taken this lesson by heart and replaced the loss function in the $L_2$ solver with the one-sided Huber loss function (See **Methods** in the main text for details).

Yet, how does this affect the full cell extraction pipeline in Eq. (6), which has been developed primarily for the $L_2$ loss minimization?

Fortunately, the theoretical motivations  we provided for each of the design choices in Eq. (6) made no assumption on the nature of the loss, and can be seamlessly incorporated into a robust algorithm that minimizes a different loss function. Fortunately, as shown through our extensive simulation benchmarks (**Table S1**), the replacement of the loss function in Eq. (6) endows EXTRACT with additional unique advantages compared to the CNMF framework:

- Similar to CNMF, EXTRACT performs cell finding greedly, but with each greedy subroutine minimizing the robust loss. Due to the fact that increased robustness prevents the cells' spatial profiles from occupying low-information pixels (**Fig. 1J-L**), the use of the robust loss helps increase the efficiency and effectiveness of cell finding, especially by mitigating the initialization of duplicate filters.
- Unlike CNMF, EXTRACT performs the alternating estimation and removal of garbage cells in an iterative manner. Thanks to this gradual nature of the process, garbage and/or duplicate cells that were not picked up in earlier iterations by the quality metrics can be discarded in later iterations, which allows milder quality cutoffs to prevent discarding real cells to achieve high precision (**Figs. 3B** and **S4A**).
- The use of the robust loss for the cell-refinement leads to faster convergence even when the movie has only Gaussian noise, as shown in our simulation benchmarks (See **App. Fig. 4** in **Supplementary Note 3**).
- After the removal of the garbage cells, the cells' activity traces are re-estimated with EXTRACT, which allows to capture any missing signals that might have been explained away by the existence of duplicate or spurious filters (See **Section 6.1.** below).

Though the use of the robust loss contributes to every step of the cell extraction pipeline, arguably the most important step is the final regression to estimate the $Ca^{2+}$ activity traces. This step constitutes the cell extraction outputs, and any mistake in this step can have effects on the biological conclusions drawn from the data analysis (**Figs. 6** and **7**). Therefore, next we consider the common methods used in the field for estimating $Ca^{2+}$ activity traces.

### §4. Common estimators for the $Ca^{2+}$ activity traces

The final step in the cell extraction routine introduced in Eq. (6) involves fixing the identified cells' spatial profiles, $\hat{\boldsymbol{S}}$, and performing one final regression on the movie, $\boldsymbol{M}$, to estimate the temporal activities of the cells, $\hat{\boldsymbol{T}}$. The goal is to obtain an accurate estimate of $\hat{\boldsymbol{T}}$ under the unknown noise distribution, $\boldsymbol{\sigma}$, which might have spatiotemporal correlations due to unobserved cell activities, neuropil, and/or imaging related noise, all of which can arise in an experimental setting. However, which optimization method is the best fit for this problem?

To answer this question, we consider an idealized scenario with the ground truth $\boldsymbol{S}$, and provide a theoretical account of errors in statistical estimators regularly utilized in the field to estimate $\hat{\boldsymbol{T}}$, whereas Section **§5** will discuss the effects of non-idealities introduced by the background components and the estimation errors in $\hat{\boldsymbol{S}}$. For this section, without loss of generality and for notational simplicity, we make two assumptions: i) $\boldsymbol{S}^T\boldsymbol{S}$ has been normalized to have unity diagonal (See **Section 6.4**), corresponding to normalized areas for cells, and ii) $\boldsymbol{T} \in \mathbb{R}^{n_{cells}}$, $\boldsymbol{M} \in \mathbb{R}^{n_{pixels}}$ are vectors, *i.e.*, the movie has a single frame.

#### 4.1. Quantifying the errors in the $Ca^{2+}$ activity trace estimators

To quantify the errors of the $Ca^{2+}$ activity trace estimators, $\hat{\boldsymbol{T}}$, and provide theoretical guarantees on their use, we now turn to the point estimation theory and conduct a theoretical study by first defining a metric of error. For simplicity, and interpretability, we choose the mean squared error:

$$\mathcal{L}(\boldsymbol{T}, \hat{\boldsymbol{T}}) = \frac{1}{n_{cell}} E[||\boldsymbol{T} - \hat{\boldsymbol{T}}||_2^2] = \frac{1}{n_{cell}} E[Tr[(\boldsymbol{T} - \hat{\boldsymbol{T}})^T (\boldsymbol{T} - \hat{\boldsymbol{T}})]] \,,$$

which can be written in terms of bias and variance terms:

$$\mathcal{L}(\boldsymbol{T}, \hat{\boldsymbol{T}}) = \frac{1}{n_{cell}} E[||\boldsymbol{T} - \hat{\boldsymbol{T}}||_2^2] = \frac{1}{n_{cell}} ||\boldsymbol{T} - E[\hat{\boldsymbol{T}}]||_2^2 + \frac{1}{n_{cell}} E[||\hat{\boldsymbol{T}} - E[\hat{\boldsymbol{T}}]||_2^2] \,,$$

in which the first term corresponds to the bias squared and the second term to the variance.

#### 4.2. Least-squares estimation

If the noise distribution, $\boldsymbol{\sigma}$, follows a set of iid Gaussian distributions with the variances $\sigma_{iid}^2$, ignoring spatiotemporal correlations, the $Ca^{2+}$ activity traces can be estimated by minimizing the $L_2$ movie reconstruction loss $||\boldsymbol{M} - \boldsymbol{S}\hat{\boldsymbol{T}}||_F^2$ , which leads to the least-squares estimate:

$$\hat{\boldsymbol{T}}_{ls} = (\boldsymbol{S}^T\boldsymbol{S})^{-1}\boldsymbol{S}^T\boldsymbol{M} = \boldsymbol{T} + (\boldsymbol{S}^T\boldsymbol{S})^{-1}\boldsymbol{S}^T\boldsymbol{\sigma}, \quad (7)$$

where we used the statistical model of the movie in Eq. (1) to obtain a statistical distribution for the estimate $\hat{\boldsymbol{T}}_{ls}$, which follows a multivariate Gaussian distribution with mean $\boldsymbol{T}$ and variance $Var[(\boldsymbol{S}^T\boldsymbol{S})^{-1}\boldsymbol{S}^T\boldsymbol{\sigma}]$. The current state-of-the-art pipelines primarily use this principle, supported by assumptions regarding calcium trace shapes, non-negativity, and background modeling, to estimate the $Ca^{2+}$ activity traces.

*__Bias:__* If the cells are not spatially overlapping, no two rows of $\boldsymbol{S}^T\boldsymbol{\sigma}$ would contain a shared component of $\boldsymbol{\sigma}$ and $\hat{\boldsymbol{T}}_{ls}$ would have uncorrelated noise. On the other hand, if cells are overlapping, the random noise in the movie pixels would mix and match, leading to $\hat{\boldsymbol{T}}_{ls}$ having Gaussian, but correlated noise. Consequently, even if there are no correlations in the ground truth cell activities, the least-squares estimate of the $Ca^{2+}$ activity traces for overlapping cells could have spurious, non-vanishing correlations. Fortunately, since the least-square estimation has zero bias, $E[\hat{\boldsymbol{T}}] = \boldsymbol{T}$, the crosstalk between neighbors can be mitigated, at least, at the time-averaged activities.

<u>*Variance:*</u> Unfortunately, the least-squares estimate can potentially have a high variance:

$$\frac{1}{n_{cell}}E[||\hat{\boldsymbol{T}}_{ls} - E[\hat{\boldsymbol{T}}_{ls}]||_2^2] = \frac{1}{n_{cell}}E[Tr[\boldsymbol{\sigma}^T\boldsymbol{S}\,(\boldsymbol{S}^T\boldsymbol{S})^{-2}\boldsymbol{S}^T\boldsymbol{\sigma}]] = \frac{\sigma_{iid}^2}{n_{cell}}Tr[(\boldsymbol{S}^T\boldsymbol{S})^{-1}] \geq \sigma_{iid}^2. \quad (8)$$

The lower bound stems from the identity $Tr[(\boldsymbol{S}^T\boldsymbol{S})^{-1}] \geq n_{cells}^{\;2}Tr[\boldsymbol{S}^T\boldsymbol{S}]^{-1}$ and the fact that the diagonals of $\boldsymbol{S}^T\boldsymbol{S}$ are all normalized to one. If the cells are non-overlapping, $\boldsymbol{S}^T\boldsymbol{S} = \boldsymbol{I}$, then the variance achieves its lower bound. However, if two cells are highly overlapping, a scenario we discuss in **Section 6.1**, the variance can grow indefinitely. For now, we consider the case of mild/negligible overlaps.

<u>*Mild overlap between cells:*</u> Next, we consider how least-squares estimates handle mild overlaps between cells. To understand this, we model the symmetric overlap matrix as $\boldsymbol{S}^T\boldsymbol{S} \approx \boldsymbol{I} + \delta\boldsymbol{\Sigma}$, where $\boldsymbol{\Sigma}$ is a symmetric matrix and has zero diagonal and $\delta$ is a small number. Moreover, $\boldsymbol{\Sigma}$ is assumed to be sparse (mild overlap) and whenever it is not zero, $\Sigma_{ij} \sim O(1)$ such that the order of magnitude value is absorbed into the small $\delta$. Performing a Taylor approximation for $(\boldsymbol{S}^T\boldsymbol{S})^{-1}$, we obtain:

$$(\boldsymbol{I} + \delta\boldsymbol{\Sigma})^{-1} = \boldsymbol{I} - \delta\boldsymbol{\Sigma} + \delta^2\boldsymbol{\Sigma}^2 + O(\delta^3),$$

which leads to the approximate trace estimation error:

$$\frac{1}{n_{cell}}E[||\boldsymbol{T} - \hat{\boldsymbol{T}}_{ls}||_2^2] = \frac{\sigma_{iid}^2}{n_{cell}}Tr[(\boldsymbol{S}^T\boldsymbol{S})^{-1}] \approx \frac{\sigma_{iid}^2}{n_{cell}}Tr[(\boldsymbol{I} + \delta\boldsymbol{\Sigma})^{-1}] \approx \sigma_{iid}^2(1 + C\delta^2),$$

for some $C = O(1)$ that is proportional to the average number of neighbors.

This theoretical calculation provides a potential explanation for the success of the least-squares estimation for the case when cells are mildly overlapping, *e.g.*, the two-photon imaging conditions, as the linear contribution of the overlap corrections to the trace estimation errors vanish. On the other hand, this is no longer the case when the overlap between cells becomes significant, as we will discuss in **Section 6.1,** which makes least-squares not a viable option for a diverse set of imaging and experimental conditions.

#### 4.3. Averaging inside regions of interests

The second approach to the estimation of $Ca^{2+}$ activity traces, which is also the fastest method, is to average pixel activities inside $\boldsymbol{S}$, referred to as 'ROI averaging':

$$\hat{\boldsymbol{T}}_{roi} = \boldsymbol{S}^T\boldsymbol{M} = \boldsymbol{S}^T\boldsymbol{S}\,\boldsymbol{T} + \boldsymbol{S}^T\boldsymbol{\sigma}.$$

This trace estimation approach is frequently used in prior work[11,12], especially when there are strict processing time constraints. Similar to the least-squares estimates, $\hat{\boldsymbol{T}}_{roi}$ follow a multivariate Gaussian distribution with mean $\boldsymbol{S}^T\boldsymbol{S}\,\boldsymbol{T}$ and variance $Var[\boldsymbol{S}^T\boldsymbol{\sigma}]$.

*Bias:* At a first glance, $\hat{\boldsymbol{T}}_{roi}$ has a bias of $(\boldsymbol{S}^T\boldsymbol{S} - \boldsymbol{I})\boldsymbol{T}$. This finite bias, which can lead to cross-talk between neighboring cells, is the primary reason that ROI averaging is seen as inferior to the least-squares estimate in Eq. (7), though the two are equivalent when $\boldsymbol{S}^T\boldsymbol{S} = \boldsymbol{I}$, *e.g.*, when there is no overlap between cells.

*Variance:* If the noise distribution, $\boldsymbol{\sigma}$, follows a set of iid Gaussian distributions with the variances $\sigma^2_{iid}$, the variance of the ROI estimator is constant regardless of $\boldsymbol{S}^T\boldsymbol{S}$:

$$\frac{1}{n_{cell}}E[||\hat{\boldsymbol{T}}_{roi} - E[\hat{\boldsymbol{T}}_{roi}]||_2^2] = \frac{1}{n_{cell}}Tr[E[\boldsymbol{\sigma}^T\boldsymbol{S}\,\boldsymbol{S}^T\boldsymbol{\sigma}]] = \frac{1}{n_{cell}}Tr[\boldsymbol{S}\,\boldsymbol{S}^T E[\boldsymbol{\sigma}\boldsymbol{\sigma}^T]] = \sigma^2_{iid}.$$

In words, $\hat{\boldsymbol{T}}_{roi}$ achieves the lower bound on the variance of $\hat{\boldsymbol{T}}_{ls}$ in all cases, *i.e.*, $\hat{\boldsymbol{T}}_{roi}$ can have significantly lower errors compared to $\hat{\boldsymbol{T}}_{ls}$ when the overlap between cells is high.

<u>*Mild overlap between cells:*</u> To understand how averaging the pixel activities inside an ROI performs when there is mild overlap between cells, we once again turn to our model with $\boldsymbol{S}^T\boldsymbol{S} \approx \boldsymbol{I} + \delta\,\boldsymbol{\Sigma}$, which leads to the approximate trace estimation error:

$$\frac{1}{n_{cell}}E[||\hat{\boldsymbol{T}}_{roi} - \boldsymbol{T}||_2^2] \approx \sigma_{iid}^2 + \frac{\delta^2}{n_{cell}}Tr[\boldsymbol{T}^T\Sigma^2\boldsymbol{T}] \approx \sigma_{iid}^2(1 + C(\boldsymbol{T})\delta^2)$$

for some $C(\boldsymbol{T}) = O(1)$ that is proportional to the average number of neighbors and the activity levels of the neurons.

Similar to the case with least-squares estimates, the linear term of the overlap has a vanishing contribution to the trace estimation errors for $\hat{\boldsymbol{T}}_{roi}$. However, ROI averaging is distinct in the sense that the proportionality constant of the second order correction depends on the overall activity levels of the neurons in the movie. Specifically, when cells are silent, *i.e.*, $\boldsymbol{T} = 0$, ROI averaging can outperform least-squares estimation. In contrast, when neighboring cells are active, which contributes to the error term via $Tr[\boldsymbol{T}^T\Sigma^2\boldsymbol{T}]$, ROI averaging can lead to significant cross contamination. Therefore, similar to the least-squares, ROI averaging is not a viable option for a diverse set of imaging and experimental conditions.

#### 4.4. Least-squares estimation with model-inspired constraints

As we discussed above, least-squares estimation can have high variance, whereas ROI averaging, despite its low variance, can introduce undesirable bias, *e.g.*, the crosstalk contamination between neighbors. Using the fact that cells' $Ca^{2+}$ traces are assumed to be non-negative in the movie generation model (Eq. (1), $\boldsymbol{T} \geq 0$), the non-negative least-squares estimate can provide a desirable common ground. Specifically, it can account for the crosstalk between observed cells thanks to the least-squares regression, and has reduced, bounded variance thanks to the non-negativity constraint, as we show next.

*Bounded variance of non-negative least-squares:* The non-negative least-squares estimator is defined as the solution of the problem:

$$\begin{aligned} &\text{Minimize} \quad ||\boldsymbol{M} - \boldsymbol{S}\hat{\boldsymbol{T}}_{nnls}||_2^2 \\ &\text{subject to} \quad \hat{\boldsymbol{T}}_{nnls} \geq 0 \end{aligned}$$

Although there is no closed-form solution for $\hat{\boldsymbol{T}}_{nnls}$, a simple argument proves that the variance of $\hat{\boldsymbol{T}}_{nnls}$ is bounded from above. This is because, since $\hat{\boldsymbol{T}}_{nnls} \geq 0$ and $\boldsymbol{S} \geq 0$ by design, one can bound $\hat{\boldsymbol{T}}_{nnls}$ by using the extreme values of $\boldsymbol{S}$ and $M$ via $\hat{\boldsymbol{T}}_{nnls} \leq \max_{i \in pixels} (\boldsymbol{M}_i) / \min_{\boldsymbol{S}_i \neq 0} (\boldsymbol{S}_i) = \tilde{\boldsymbol{T}}_{nnls}$.

For any $\hat{\boldsymbol{T}}_{nnls} \geq \tilde{\boldsymbol{T}}_{nnls}$, the resulting movie reconstruction error is higher following the inequalities:

$$\text{For } \hat{\boldsymbol{T}}_{nnls} \geq \tilde{\boldsymbol{T}}_{nnls}: \quad (\boldsymbol{S}\hat{\boldsymbol{T}}_{nnls} - \boldsymbol{M})_i \geq (\boldsymbol{S}\tilde{\boldsymbol{T}}_{nnls} - \boldsymbol{M})_i \geq 0 \Rightarrow ||\boldsymbol{M} - \boldsymbol{S}\hat{\boldsymbol{T}}_{nnls}||_2^2 \geq ||\boldsymbol{M} - \boldsymbol{S}\tilde{\boldsymbol{T}}_{nnls}||_2^2.$$

*Additional constraints for trace estimation:* In addition to non-negativity, previous research has enforced exponential kernels during the estimation of cells' $Ca^{2+}$ activity traces[5]. This approach aims to reduce the variance significantly by incorporating a prior regarding the temporal profiles of $Ca^{2+}$ activities. An example approach is the CAIMAN pipeline[5], which performs simultaneous fitting of an exponential kernel and its deconvolution to model the $Ca^{2+}$ signals. However, this adds an additional complexity to the estimation of $Ca^{2+}$ activities, and is not necessarily needed as per our extensive benchmarks (**Table S1, Figs 3,** and **S3**).

#### 4.5. An experimentally appropriate unit for traces

When we considered the variance of the $Ca^{2+}$ activity traces, *e.g.*, for the least-squares estimator, see Eq. (8), we observed an independence on the number of pixels encapsulated by the cell. At first, this seems to be at odds with the empirical observation that having higher resolution decreases the variance of the estimated $Ca^{2+}$ activity traces. Yet, as we show in this section, it is not. To understand why, we need to revisit the normalization we assumed at the beginning of our discussions for analytical convenience, *i.e.*, $\boldsymbol{S}^T\boldsymbol{S} = I$. This normalization ignores the spatial scale and simply assigns unit area to each cell, and are not in the units we typically use in experiments.

Consider a simplified scenario with two non-overlapping cells with approximately equal brightness, but very different cell areas (call them $A_1$ and $A_2$ with $A_2 \gg A_1$). In the normalization $\boldsymbol{S}^T\boldsymbol{S} = I$, the two cells would end up having different values for their $Ca^{2+}$ activity traces, $T_2 = T_1\sqrt{A_2/A_1}$. In other words, even though both cells had approximately equal brightness, in this normalization, the absolute value of the $Ca^{2+}$ activity trace is larger for the cell with higher area. This is experimentally counter-intuitive, and is the reason why $\boldsymbol{S}^T\boldsymbol{S} = I$ normalization is only a theoretical tool, not a practical one.

Practically, it is common to use the $\Delta F/F$ normalization, in which the maximum value of cells' spatial profile, $\boldsymbol{S}$, not the cells' area, is normalized such that $\forall_{j=1,\dots,n_{cells}}: \max_i \boldsymbol{S}_{ij} = 1$. Unlike before, this normalization leads to the equal values for cells as long as they have the same brightness level, even if they may have different areas. In this normalization, noting that $\boldsymbol{S}^T\boldsymbol{S} \sim O(N_{pixel\,per\,cell})$, the trace estimation errors follow $\sim \sigma_{iid}^2/O(N_{pixel\,per\,cell})$, not $\sigma_{iid}^2$ as in Eq. (8). In this normalization, it is clear that having higher cell resolution leads to lower trace estimation errors.

Returning to our original concern, we conclude that $\boldsymbol{S}^T\boldsymbol{S} = I$ normalization masks the effect of adding more pixels, *i.e.,* more samples, for the trace estimation problem. In this case, while the variance of the estimated $Ca^{2+}$ activity trace stays the same with added pixels, the activity levels in $\boldsymbol{T}$ increase with $\sim O(\sqrt{N_{pixel\ per\ cell}})$, leading to higher signal to noise ratio. This recapitulates the results we obtained above with the $\Delta F/F$ normalization, though with additional steps required to put things in context. Therefore, instead of the counter-intuitive $\boldsymbol{S}^T\boldsymbol{S} = I$ units, in the outputs of EXTRACT, we use the $\Delta F/F$ normalization.

#### §5. Estimation of $Ca^{2+}$ activity in movies with background components

Until now, we assumed that the noise distributions in the movies were identical and independent. In reality, movies (Eq. (1)) contain correlated noise components, which may be non-Gaussian and are usually modeled as a linear background ($\boldsymbol{S}_B, \boldsymbol{T}_B$) as in Eq. (1). To handle the potential contamination of estimated $Ca^{2+}$ signals by the background components, prior algorithms either modeled them explicitly as part of the regression problem[5], or estimated the background separately and subtracted from the movie [4].

In practice, modeling of the background explicitly often requires careful care and fine-tuning of hyperparameters, *e.g.*, the number of rank components to model in CAIMAN[5] or the regularization parameters in SEUDO[13] (**Fig. S5A, B**). Yet, inaccurate or incomplete identification of the background components may introduce non-Gaussian contaminants in the form of spurious activity or explain away true-positive neural signals, which we discuss next.

#### 5.1. Inaccurate estimation of cells' spatial profiles

Though, for simplicity of discussion so far, we had assumed that the estimators of cells' $Ca^{2+}$ activity traces had access to the cells' ground truth spatial profiles, $\boldsymbol{S}$, in reality, there are often inevitable errors in the estimation of $\hat{\boldsymbol{S}} \neq \boldsymbol{S}$. Consequently, instead of the ground truth $\boldsymbol{S}$, the estimation of the $Ca^{2+}$ activity traces often relies on an estimate, $\hat{\boldsymbol{S}} = \boldsymbol{S} - \Delta_S$. Thus, even if the noise components of the movie, $\boldsymbol{\sigma}$, are sampled from purely iid Gaussians and background components, $\hat{\boldsymbol{S}}_B$, are completely (but not necessarily correctly, $\hat{\boldsymbol{S}}_B \neq \boldsymbol{S}_B$) identified, the estimation of cells' $Ca^{2+}$ signals would need to account for non-Gaussian contaminants. Specifically, the statistical model of the movie generation in Eq. (1) can be rewritten as:

$$\boldsymbol{M} = \boldsymbol{S}\,\boldsymbol{T} + \boldsymbol{\sigma} = \hat{\boldsymbol{S}}\,\boldsymbol{T} + \Delta\boldsymbol{S}\,\boldsymbol{T} + \boldsymbol{\sigma},$$

in which $\Delta\boldsymbol{S}\,\boldsymbol{T}$ constitute spatiotemporally correlated non-Gaussian contaminants, *i.e.*, unexplained $Ca^{2+}$ activity from cells or background components, for the subsequent estimation of $Ca^{2+}$ activity traces.

#### 5.2. Non-Gaussian noise from inaccurate number of estimated spatial profiles

Apart from the inaccurate estimation of the spatial profiles, estimating the number of background components or cells incorrectly can also introduce non-Gaussian contaminants. The estimation can be incorrect in two distinct manners: i) missing spatial components for the $Ca^{2+}$ sources, and/or ii) estimating duplicate profiles for the same cells' spatial location.

Missed components: In the case of missed components, we can simply rewrite the spatiotemporally correlated $Ca^{2+}$ activity contributions to the movie as:

$$\boldsymbol{S}\,\boldsymbol{T} = \boldsymbol{S}_{found}\boldsymbol{T}_{found} + \boldsymbol{S}_{missed}\boldsymbol{T}_{missed},$$

where the second term adds additional terms to the non-Gaussian and correlated contaminants.

Duplicate components: For the case of duplicate identification of spatial profiles for the same cell, the situation is a bit more nuanced and potentially more dangerous. In this case, the $Ca^{2+}$ activity events of a single cell can be shared between two putative cells due to the increased multicollinearity of the regression stemming from the duplication. Not only can this introduce spurious correlations to the population, but it can mask reliable signals by dividing events into two spurious cells. Additionally, this scenario introduces yet another non-Gaussian noise component. To illustrate how, let us consider a simple scenario, in which all cells are identified, but there is also one more spurious spatial profile such that $\widetilde{\boldsymbol{S}} = [\boldsymbol{S}; \boldsymbol{s}_{extra}]$. Then, the movie model becomes:

$$\boldsymbol{M} = \boldsymbol{S}\hat{\boldsymbol{T}} - \boldsymbol{s}_{extra}\boldsymbol{t}_{extra} + \boldsymbol{\sigma}.$$

In this case, the term, $\boldsymbol{s}_{extra}\boldsymbol{t}_{extra}$, adds unnecessary complexity to the problem by contributing to the non-Gaussian noise component. Moreover, since $\boldsymbol{s}_{extra}$ has a likely overlap with another profile from $\boldsymbol{S}$, due to the duplication, the spurious cell can catch some of the real $Ca^{2+}$ activity signal with $\boldsymbol{t}_{extra}$ and subtract it from the movie.

To sum up, while missing spatial components often add crosstalk to the estimation of traces, having extra duplicates can explain signal away. Given the low cell finding accuracy of the $L_2$-estimation based algorithms, these effects can become particularly prominent in high density scenarios (**Fig. 3C-E**). Additionally, post-processing tools that denoise $Ca^{2+}$ activity traces using the spatial profiles from these pipelines cannot correct the imperfections in cells' profile estimations. However, our experiments in the main text (**Figs. 1A-D and S1**) suggest that robust estimation of $Ca^{2+}$ activity signals mitigates the non-Gaussian noise, even when 80% of all cells' spatial profiles are not identified from the movie.

#### 5.3. Spatial high-pass filtering for mitigating neuropil

Instead of explicitly modeling the background[5] or performing heuristic neuropil removal steps[4], a principled approach that can remove spatially correlated neuropil activities, while retaining the cells' $Ca^{2+}$ activity signals, is the use of spatial high-pass filtering. To understand how spatial high-pass filtering can remove the spatially correlated background by keeping the neuronal signals unaffected, we start by rewriting Eq. (1):

$$\boldsymbol{M} = \boldsymbol{S}_{cell}\boldsymbol{T}_{cell} + \boldsymbol{S}_{B}\boldsymbol{T}_{B} + \boldsymbol{\sigma},$$

in which we assume that some background components, $\boldsymbol{S}_B$, are correlated in spatial distances that are significantly larger than cells' diameters, and can induce cross-talk during the estimation of $Ca^{2+}$ activity traces. Such cases are common, especially in one-photon movies, due to the increased background noise and $Ca^{2+}$ activities from the out of focus sources[19].

To understand the effects of spatial high-pass filtering, we first consider a general linear filter, $\mathfrak{F}[.]$, applied to each movie frame independently with the goal to remove the background contaminants. This approach would affect $\boldsymbol{S}$ but not $\boldsymbol{T}$, which we show by using the fact that any linear filter, $\mathfrak{F}[.]$, can be considered as a convolution with a corresponding kernel, $\mathfrak{f}(.)$:

$$\mathfrak{F}[\boldsymbol{M}]_{ij} = \sum_{h \in pixels} \boldsymbol{M}_{hj}\mathfrak{f}(i-h) = \sum_{h}[\boldsymbol{S}_h\boldsymbol{T}_j + \boldsymbol{\sigma}_{hj}]\mathfrak{f}(i-h) = [\sum_{h}\boldsymbol{S}_h\mathfrak{f}(i-h)]\boldsymbol{T}_j + \sum_{h}\boldsymbol{\sigma}_{hj}\mathfrak{f}(i-h),$$

$$\Rightarrow \quad \boldsymbol{M} = \mathfrak{F}[\boldsymbol{S}_{cell}]\boldsymbol{T}_{cell} + \mathfrak{F}[\boldsymbol{S}_B]\boldsymbol{T}_B + \mathfrak{F}[\boldsymbol{\sigma}]. \tag{9}$$

As long as linear filtering is applied only in the spatial domain, without mixing the temporal components, the statistical model gets updated with a new set of filters $\mathfrak{F}[\boldsymbol{S}]$ and noise $\mathfrak{F}[\boldsymbol{\sigma}]$; but the temporal dynamics remain unaffected. In other words, a linear spatial filtering approach can selectively reduce the background without decreasing the signal as long as $\mathfrak{F}[\boldsymbol{S}_B] \approx 0$ and $\mathfrak{F}[\boldsymbol{S}_{cell}] \approx \boldsymbol{S}_{cell}$. Yet, spatial filtering may affect the shape of cells' spatial profiles, the spatial profiles of the background contaminants, and the noise distribution $\boldsymbol{\sigma}$.

We use a linear spatial high-pass filter, a Butterworth's filter of 4th order (**Methods**), in which we can pick the high-pass cutoff frequency such that only spatial structures that are an order of magnitude larger than the cells' typical sizes would be removed. In this case, $\mathfrak{F}[\boldsymbol{S}_B] \approx 0$

for large contaminants, whereas $\mathbb{F}[\boldsymbol{S}_{cell}] \approx \boldsymbol{S}_{cell}$ for the cells or smaller contaminants. The only caveat is the potential change in noise distribution, *i.e.*, $\mathbb{F}[\boldsymbol{\sigma}]$ may not be the same as $\boldsymbol{\sigma}$, which may throw off $Ca^{2+}$ activity trace estimators with strong assumptions.

### §6. Theoretical and empirical evaluations of common $Ca^{2+}$ activity trace estimators

To test the concepts introduced in this note so far, we perform several benchmarks. First, we consider a simplified theoretical scenario, in which there are two overlapping cells with known ground truth spatial profiles. This case is distinct, and easier than, the scenario in **Figure 1A-D**, which contains an unknown distractor cell. We use this theoretical case study to illustrate that the least-squares estimates can have infinite variance. As the second benchmarking strategy, we perform an empirical study with simulated $Ca^{2+}$ imaging movies, as part of our extensive benchmarking strategy (**Table S1**)

#### 6.1. An analytical case study with two overlapping cells

We consider a case study scenario of a hypothetical $Ca^{2+}$ imaging movie, which contains two overlapping cells with the overlap area $(S^T S)_{1,2} = (S^T S)_{2,1} = \alpha$, where $\alpha \in [0, 1)$. Based on our discussions in the previous sections, we can compute the trace estimation errors as:

$$\mathcal{L}_{ls} = \frac{\sigma_{iid}^2}{2} Tr[(\boldsymbol{S}^T\boldsymbol{S})^{-1}] = \frac{\sigma_{iid}^2}{1-\alpha^2},$$
$$\mathcal{L}_{roi} = \sigma_{iid}^2 + \frac{Tr[\boldsymbol{T}^T(\boldsymbol{S}^T\boldsymbol{S}-I)^2\boldsymbol{T}]}{2} = \sigma_{iid}^2 + \alpha^2 < t^2 >,$$

where we define $< t^2 >$ as the average activity of the cells. As hinted before, the mean squared error of the least-squares estimates can blow-up as $\alpha \to 1$, *i.e.*, when cells are highly overlapping. In contrast, the trace estimation errors for the ROI averaging remain bounded above by $\sigma_{iid}^2 + < t^2 >$. When the cells are silent, ROI averaging outperforms the least-squares, unless $\alpha = 0$, in which case both estimators are equivalent. For any non-zero $< t^2 >$, there exists a critical overlap value $\alpha_c$ beyond which traces estimated via ROI averaging outperform least-squares estimates. Thus, surprisingly, even traces estimated via crude ROI averaging can outperform those estimated via least-squares, as long as the cells are highly overlapping.

Though the variance may be high, what does this mean in a real dataset? Practically, this can lead to events being shared between highly overlapping cells, potentially leading to true-positive signals being explained away. This may become particularly concerning when duplicate cells are initialized in the same spatial location. Moreover, though cells are not necessarily highly overlapping in a traditional two-photon $Ca^{2+}$ imaging movie (though see **Figure 4** for a case with highly packed cells), the neuropil signals regularly overlap with cells' locations. Bringing all together, the theoretical example discussed in this section provides an important motivation for why least-squares is not a reasonable estimator, instead additional constraints may be needed to accurately estimate cells' $Ca^{2+}$ activity traces.

While the scenario with two overlapping cells is insightful and interesting to consider analytically, to understand how non-negative least squares and robust regression models compare to the traditional approaches to $Ca^{2+}$ activity estimation under realistic constraints, we resort to empirical benchmarking for the rest of this section.

#### 6.2. Empirical validation of the theoretical errors

To empirically validate the theoretical formulas we derived for the $Ca^{2+}$ activity trace estimators, *e.g.*, Eq. (8) for the least-squares estimates, we simulated $Ca^{2+}$ imaging movies with highly overlapping cells (**Appendix Figure 1**). These movies contained only Gaussian noise, in line with the assumptions of least-squares problem.

To estimate the cells' $Ca^{2+}$ activity traces, we used three trace estimation algorithms: least-squares, non-negative least-squares, and non-negative robust estimation with adaptive $\boldsymbol{\kappa}$ updates. We initialized these trace estimators with cells' ground truth spatial profiles, and afterwards computed the errors in the estimated $Ca^{2+}$ activity traces. The empirical errors of the least-squares estimates matched the theoretical predictions from Eq. (8) for all simulations, regardless of the resolution (**Appendix Figure 1A**). Non-negative estimators, robust or least-squares, had significantly lower errors, in line with our discussion **in Section 4.4**, indicating that the non-negativity constraint mitigated the high variance of the least-squares estimation.

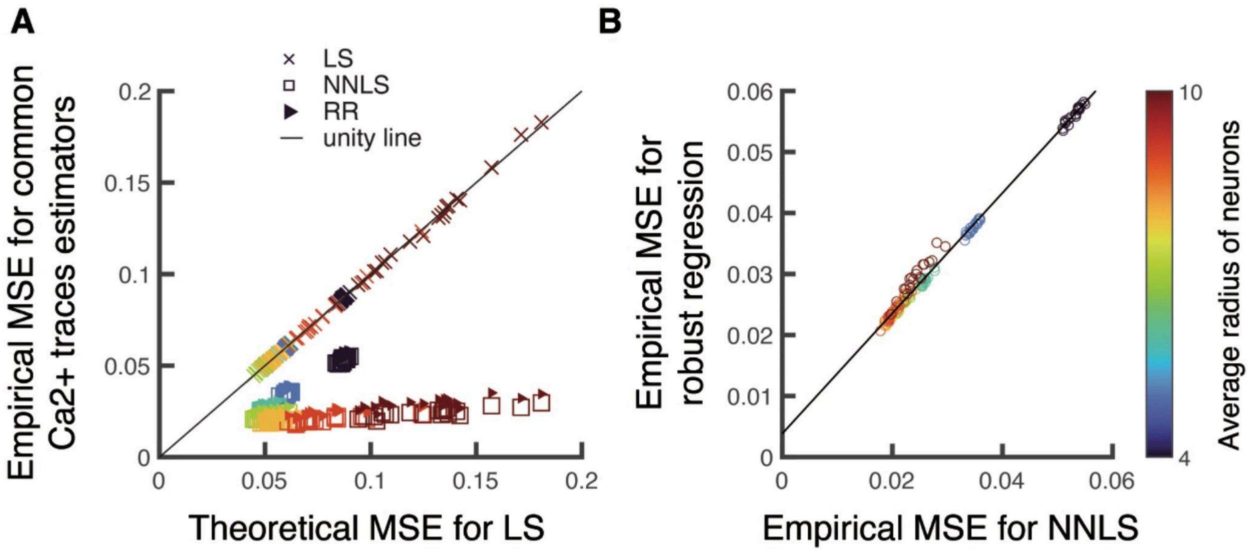

***Appendix Figure 1. Validation of the theoretical and empirical trace estimation errors of common estimators.***

To compare the trace estimation errors (in units of $(\Delta F/F)^2$, normalized by $\sigma_{iid}^{2}$), we simulated movies with $160 \times 160$ $mm^2$ FOVs, which contained only Gaussian noise and 100 cells each. Each datum in the plots corresponds to a movie, in which we averaged the mean squared error between ground truth and estimated $Ca^{2+}$ activity traces across 100 cells. **A.** The predicted theoretical mean squared errors for $\hat{\boldsymbol{T}}_{ls}$ matched the empirical values, whereas both non-negative least squares (NNLS) and (non-negative) robust regression (RR) had lower trace estimation errors. **B.** The empirical trace estimation errors between NNLS and RR followed a linear relationship with a near unity slope, $(MSE_{RR} = 0.99 MSE_{NNLS} + 0.004, R^2 = 0.99)$.

#### 6.3. Empirical validation of the adaptive $\kappa$ estimation

In **Appendix Figure 1**, we considered ideal $Ca^{2+}$ imaging movies with only Gaussian noise and trace estimators were initialized with cells' ground truth spatial profiles. Consequently, due to the absence of non-Gaussian contaminants, the trace estimator was expected to adjust $\kappa$ values to become an approximate $L_2$ solver, *i.e.*, the limit $\kappa \to \infty$. We hypothesized that EXTRACT's adaptive estimation module (**Methods)** for $\kappa$ would be able to recover this limit approximately, by identifying the absence of non-Gaussian noise contaminants and providing similar trace estimation errors to the non-negative least-squares.

To test this hypothesis, we compared the errors computed from the non-negative least-squares and robust regression estimates for the $Ca^{2+}$ activity traces, which followed a linear relationship with near unity slope (**Appendix Figure 1B**). The close match between the errors of the two estimators provided an empirical validation that the adaptive estimation of $\kappa$ values captured the Gaussian nature of the movie noise, albeit with small errors due to finite number of updates to approximate $\kappa \to \infty$.

#### 6.4. Verification of the robust regression

Having benchmarked the trace estimation algorithms on idealized $Ca^{2+}$ imaging movies, we next simulated realistic movies with 600 cells each and some correlated noise, which aimed to model background components such as neuropil (**Appendix Figure 2, Methods**). To introduce non-Gaussian contamination (See **Section §5**), we initialized the trace estimators with a portion of ground truth cells' spatial profiles, and compared the errors of $Ca^{2+}$ activity trace estimation.

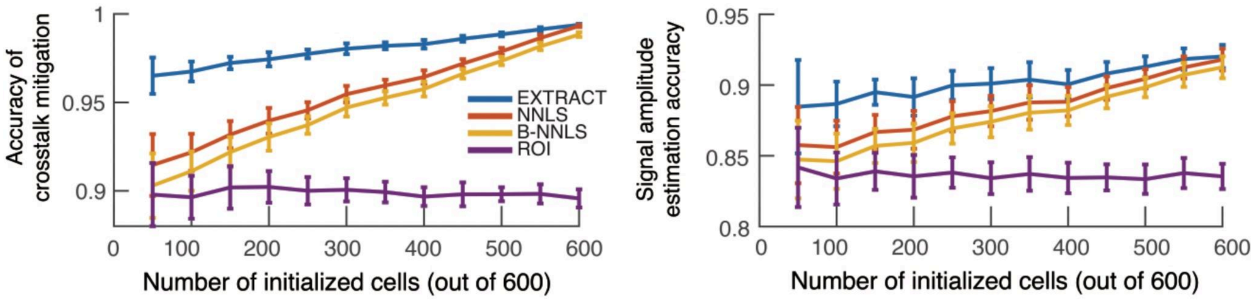

***Appendix Figure 2. Trace estimation errors under correlated and non-Gaussian noise contamination.***

To benchmark the $Ca^{2+}$ activity trace estimation quality under realistic scenarios, we simulated 20 movies with $400 \times 400$ mm$^2$ FOVs, each with 600 cells. The movies inherently contained correlated noise (5% of total noise, **Methods**), and we introduced explicit non-Gaussian noise, both of which made (non-negative) least-squares estimates non-ideal. To introduce the non-Gaussian contaminants, we initialized the $Ca^{2+}$ activity trace estimators with a portion of ground truth cells' spatial profiles (x-axis, lower number of initialized cells corresponds to larger contaminants) and compared the cross-talk mitigation and signal amplitude estimation accuracies. B-NNLS refers to non-negative least-squares estimates based on binarized spatial profiles for cells, ROI refers to the ROI averaging inside the binarized spatial profiles.

When initialized with all cells, *i.e.*, no non-Gaussian contamination, both EXTRACT and NNLS estimated the $Ca^{2+}$ activity traces with the same accuracy (**Appendix Figure 2**), which was in line with the results above (**Appendix Figure 1**). With increased background contamination, NNLS led to inferior estimation of $Ca^{2+}$ acticity traces due to high crosstalk between neighboring cells, and had similar errors as the $Ca^{2+}$ activity traces estimated via ROI averaging. In contrast, EXTRACT was able to demix neural signals even under extreme background contamination. Thus, non-negative least squares cannot account for the mitigation of crosstalk from unobserved cells and/or background sources, for which robust regression is needed. Finally, we observed that using binarized versions of $\boldsymbol{S}$, *e.g.*, those that are obtained by circling regions of activity on $Ca^{2+}$ imaging movies, led to lower quality $Ca^{2+}$ activity traces compared to their continuous valued counterparts. This observation underscores the importance of accurately estimating cells' spatial profiles for the subsequent estimation of cells' $Ca^{2+}$ activity traces.

#### 6.5. Validation of the non-negativity constraints

In **Section 4.4**, we theoretically argued for the use of the non-negativity constraint to decrease the variance in the estimated $Ca^{2+}$ activity traces. This, in turn, is particularly relevant for $Ca^{2+}$ imaging movies with low signal-to-noise (SNR) ratios, in which high levels of noise in comparison to signal can lead to increased variance. To validate this theoretical observation experimentally, we simulated additional $Ca^{2+}$ imaging movies with the same recipe as in **Appendix Figure 2**, *i.e.*, 600 cells each and with 5% correlated noise contaminants (**Appendix Figure 3A**).

To estimate cells' $Ca^{2+}$ activity traces using least-squares with and without non-negativity constraints, we initialized both algorithms with 80% percentage of ground truth cells' spatial profiles, in which 20% of the unknown cell population was intended as non-Gaussian noise contaminants. In line with the theoretical predictions, non-negative least-squares led to superior $Ca^{2+}$ activity traces, with larger benefits of non-negativity constraint observed at lower cell SNRs (**Appendix Figure 3A**). Overall, this experiment supported our observations in **Fig. S4C-D**, and provided extra evidence for the choice to use non-negativity constraints in EXTRACT's solvers.

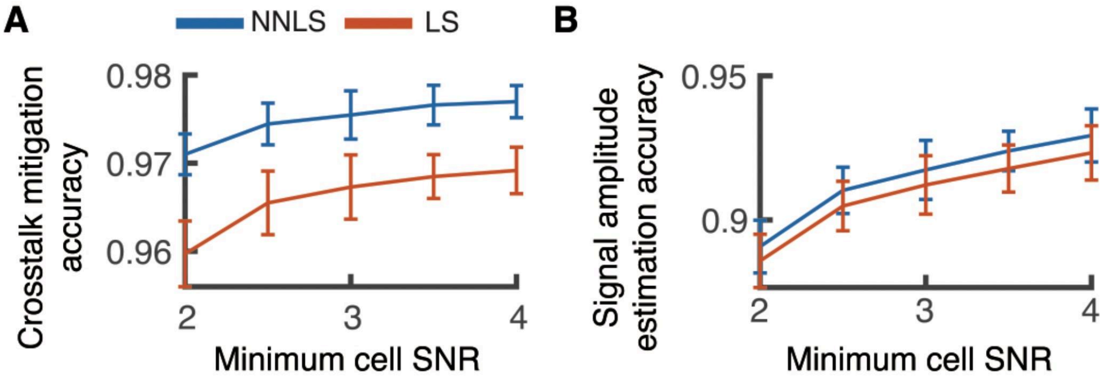

***Appendix Figure 3. Validation experiments for the non-negativity constraint.***

To test the necessity of non-negativity constraint, we simulated two-photon movies with mild (5%) correlated noise, varying levels of minimum cell SNRs, and $400 \times 400$ mm$^2$ FOVs, each containing 600 cells. To estimate cells' $Ca^{2+}$ activity traces, we initialized trace estimation algorithms with 80% percentage of ground truth cells' spatial profiles. Non-negativity constrained led to increased crosstalk mitigation (**A**) and the signal amplitude estimation (**B**) across all SNR values, but mostly for $Ca^{2+}$ imaging movies with lower quality signals.

#### 6.6. Test of spatial high-pass filtering for trace estimation

In **Section 5.3**, we theoretically argued that spatial high-pass filtering can be performed in a manner that does not affect $Ca^{2+}$ activity traces, but can filter out background components such as neuropil. To test this prediction, we simulated one- and two-photon movies with 600 cells in a $400\ \mu m\ x\ 400\ \mu m$, initialized the $Ca^{2+}$ activity trace estimators with 80% of the cells, allowed two iterations of cell refinement to best estimate $\boldsymbol{S}_{cell}$ after the spatial filtering, and (re-)extracted $Ca^{2+}$ activity traces while varying levels of the spatial high-pass cutoff (**Appendix Figure 3B**). Here, a higher cutoff corresponds to a milder filtering, *e.g.*, a cutoff of $5$ intuitively means that any movie object larger than $5$ times the radius of a typical cell would be removed via the high-pass. We observed that both for types of $Ca^{2+}$ imaging movies, mild spatial high-pass filtering led to superior $Ca^{2+}$ activity traces, particularly for the simulated one-photon $Ca^{2+}$ imaging movies with large out-of-focus cells (**Appendix Figure 3B**, *left*).

There *are* cases in which spatial high-pass filtering can be detrimental. Particularly, when the cells are firing in a highly correlated manner across the FOV, performing a high-pass filtering can end up removing neuronal $Ca^{2+}$ activities alongside the neuropil. This includes, for example, the study of dendritic activities we considered in **Fig. S8**, or the simulated $Ca^{2+}$ imaging movies with high FOV-wide neuronal activity correlations in **Fig. S6A, B.** In such cases, we either apply an extremely mild cutoff, mainly during cell finding and/or refinement, or omit it altogether.

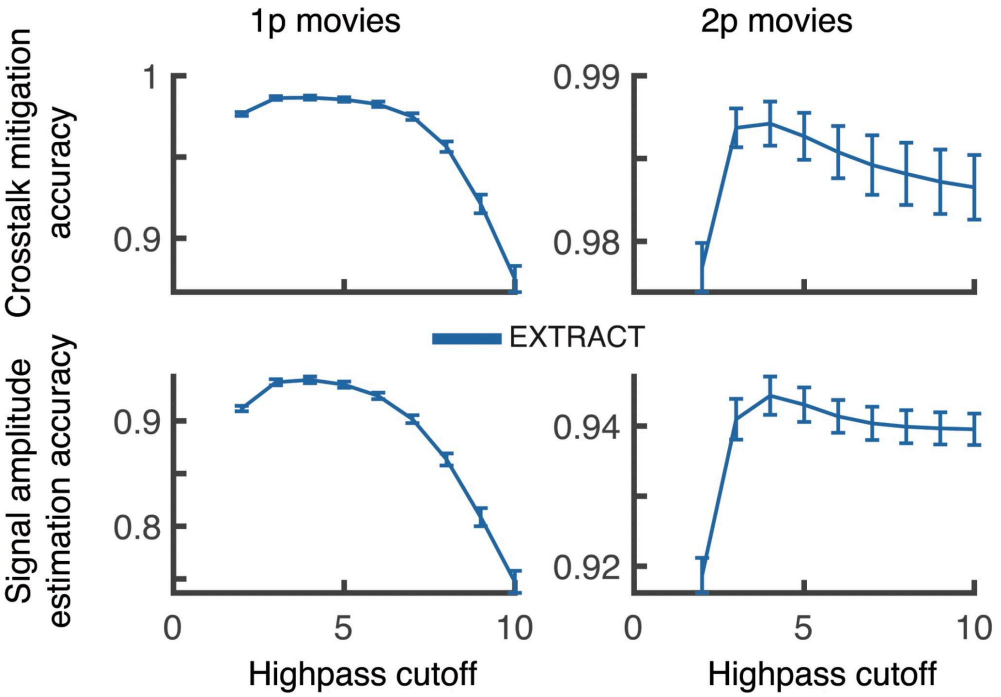

***Appendix Figure 4. Validation experiments for the high-pass filtering.***

To validate the mild high-pass filtering used by the EXTRACT pipeline, we simulated one- and two-photon movies with $400 \times 400$ mm$^2$ FOVs, each with 600 cells. Once again, to estimate cells' $Ca^{2+}$ activity traces, we initialized trace estimation algorithms with 80% percentage of ground truth cells' spatial profiles. The movies all contained moderately (30%) correlated noise, and we enforced 50% of $Ca^{2+}$ events for each cell to be jointly firing within groups of randomly chosen 50-100 neurons. Thus, we artificially created high spatiotemporal correlations not only in the background contaminants, but also in cell activity patterns. Recalling that a large high-pass cutoff value corresponds to milder spatial filtering (**Methods**), *e.g.* $\infty$ corresponds to no filtering at all, mild spatial high-pass filtering was beneficial to the cell extraction quality in both one- and two-photon $Ca^{2+}$ imaging movies.

### §7. Review of existing cell extraction algorithms

Currently, there is a long list of available cell extraction and trace estimation algorithms in the field of systems neuroscience[2–5,11–15,20–28]. Thus, to guide our benchmarking efforts, we first divided the existing algorithms into several broadly defined categories.

*Early approaches:* The simplest, and earliest, cell extraction approach is the process of hand-annotating cells by drawing ROIs on $Ca^{2+}$ imaging movies and estimating the $Ca^{2+}$ activity signals via averaging inside these (often binarized) regions. Remnants of this manual approach are still implemented in newly published pipelines[11,12]. With the advent of reasonably large movies, the early automated cell extraction approaches were developed, most notably the PCA/ICA approach[20], which we used as part of our benchmarks in this work. These early algorithms provided the first automation to the cell extraction process, but their underlying assumptions, specifically the independence assumption of $Ca^{2+}$ activity signals, were often violated (**Figures 6** and **7**).

*Deep learning approaches:* Within the last decade, cell extraction routines based on deep learning have also been applied to the cell extraction, though with limited user base and experimental success to this date[12,15,21,22,25,29]. The latest state-of-the-art approach in this direction explored deep learning based cell extraction using 3D U-nets on 1p movies, and the resulting algorithm is named DeepWonder[29]. This work reported increased cell finding accuracy, though with marginal increase in $Ca^{2+}$ activity trace estimation quality compared to CNMF-e (See Fig. 2i of Ref.[29]), which are shown in this work to be severely suboptimal (**Figures 3, S2,** and **S3**).

While a deep learning approach to cell extraction may be beneficial for specific datasets that they are trained on, ability of the deep learning algorithms to adapt to new datasets may be limited[15,29], often requiring re-training on novel datasets. Moreover, deep learning methods are known to spit out spurious information, which is carefully discussed in the traditional deep learning literature[30] but not in context with cell extraction. Given the possibility of spurious $Ca^{2+}$ activity signal predictions by deep learning models, their adoption in the field would likely benefit significantly from (or maybe require) a detailed hallucination study, which discusses not only how they perform with respect to existing algorithm based approaches, but also when they tend to fail and how. Until then, we believed this category of cell extraction methods to be newly emerging and did not test a deep learning approach in our benchmarks, spots in which were reserved for testing the state-of-the-art approaches to cell extraction.

<u>*Current state of the art methods:*</u> The current state-of-the-art approaches to cell extraction - most notably CAIMAN[5] and Suite2p[4] , but also see[24,26,31,32] - focus largely on the usability by a wide range of users. Consequently, the cell extraction pipelines in this generation include additional utilities such as motion correction[6] and across day cell registration[33]. Though these pipelines are well benchmarked in terms of their cell finding accuracies, *e.g.*, through the Neurofinder challenge[34], the quality of $Ca^{2+}$ activity trace estimation is often benchmarked with a correlation based metric. However, correlation metrics can often be misleading, *e.g.*, due to the (mis)match of exponential kernels between ground truth and the estimated $Ca^{2+}$ activity traces. Instead, in this work, we perform extensive evaluations of the trace quality via carefully designed and empirically validated quality metrics (**Methods**).

Our analysis in the main text indicated a low quality of estimated $Ca^{2+}$ activity traces obtained by a state of the art method across one- and two-photon $Ca^{2+}$ imaging movies (CAIMAN[5], **Figs. 3, S2,** and **S3)**. This observation is in line with the previous work that has shown the inferiority of $Ca^{2+}$ activity traces estimated via $L_2$ estimation methods (including Suite2p[4] and CAIMAN[5]) from two-photon $Ca^{2+}$ imaging movies. Yet, these cell extraction methods are still widely used. Therefore, from this group of algorithms, we picked CAIMAN as the representative for benchmarking, which is currently the most widely used published cell extraction method and has been validated to process one-photon $Ca^{2+}$ imaging movies[3].

<u>*Trace denoising algorithms:*</u> Following the inability of the current cell extraction approaches to provide accurate $Ca^{2+}$ activity traces, several recent works focused on developing post-processing tools to correct for errors during the trace estimation. These

algorithms often utilize the already extracted cells’ spatial footprints, $\boldsymbol{S}$ (potentially by other algorithms), and provide a set of denoised traces by accounting for neuropil contamination and/or neighboring cells’ unaccounted activity. By design, this branch of algorithms divides the cell extraction problem into two distinct parts, and focuses primarily on the latter, the trace estimation. Yet, in this approach, errors made during the estimation of cells’ spatial profiles remain uncorrected such that the trace denoising algorithms may face unnecessarily increased amounts of non-Gaussian noise contaminants, which may affect the estimation of $Ca^{2+}$ activity traces (See **Section 5.1**). This theoretical observation is in line with our findings in the main text (**Figures 3** and **S5**), in which robust estimation of cells’ spatial footprints, $\boldsymbol{S}$, led to superior $Ca^{2+}$ activity traces for all trace estimation algorithms. Most notable trace denoising algorithms include FISSA[14] for neuropil subtraction, DeepCINAC[15] and SEUDO[13] for detecting and removing false transients.

For our benchmarking experiments, we tested SEUDO, the current state-of-the-art trace denoising method. Similar to our earlier design[10] of EXTRACT, SEUDO starts with the observation that $Ca^{2+}$ imaging movies tend to have large positive outliers. Alternative to our robust regression approach, SEUDO instead trains two estimators to minimize differently regularized $L_2$ losses, in which the background contamination is modeled with a set of Gaussian shaped blobs that can absorb unexplained activity. The $Ca^{2+}$ activity traces are then computed as the minimum of the two estimated values, to prevent large positive outliers. Our benchmarks replicated the original claims in the previous work[13], namely SEUDO led to higher quality traces when applied as a post-processing tool to CAIMAN (**Figures 3** and **S5**). Yet, EXTRACT outperformed SEUDO in all our benchmarks, had less dependency on finding the right hyperparameters, and was orders of magnitude faster (**Figures 3** and **S5**).

*The use of the term robustness:* In the prior works to cell extraction, the word robust is often used colloquially, perhaps in place of the word ‘accurate’. For example, SEUDO is termed as a 'robust' approach, but our benchmarks has shown that it is hypersensitive to hyperparameters and the quality of extracted cells' spatial footprints (**Figures 3** and **S5)**. Similarly, Ref.[25] uses the term robust solely in the title, with no discussion of robustness in the rest of the work. However, there is a long list of prior work in the field of statistics for the properties of robust estimators[9]. Our use of the term 'robust' is in accord with this line of work.

*EXTRACT:* The field of statistical estimation has regularly tested ideas to mitigate unexplained non-Gaussian signals in the regression problems (similar to cell extraction literature summarized above), and robust statistics[9] has emerged as a state-of-the-art approach standing the test of time with the ability to generalize across wide range of problems. When designing EXTRACT, we based our cell extraction methodology to this literature, by introducing a new version of the Huber loss better suited for processing $Ca^{2+}$ imaging movies. This choice not only increased the generalizability and the accuracy of the resulting algorithm, but also allowed scalability thanks to the replacement of explicit background modeling with the minimization of a bi-convex loss function (**Supplementary Note 2**). Therefore, we denote EXTRACT as the flagship method of a new generation of cell extraction algorithms, which no longer requires any post-processing for denoising traces and can be implemented orders of magnitude faster compared to previous methods to allow processing large datasets (~TBs) of today and future.

**§8. Summary of key heuristics developed over the years for cell extraction**

So far, we introduced and reviewed several heuristics under the estimation theory framework, which are regularly utilized by the current state-of-the-art pipelines for extracting cells from $Ca^{2+}$ imaging videos. We first introduced the movie reconstruction paradigm, then discussed the motivations behind state-of-the-art design choices to estimate $Ca^{2+}$ activity traces in the presence of non-Gaussian noise contaminants, some of which passed our benchmarking tests and therefore are incorporated into EXTRACT, and concluded with a set of simulation experiments providing theoretical and empirical evidence for why $L_2$ approaches to cell extraction are suboptimal. In this section, we now provide a final (non-exhaustive) listed summary of the key heuristics developed in the field to this date and explain what type of inductive biases (or lack thereof) they incorporate into the cell extraction problem.

Greedy cell finding: Existing state-of-the-art approaches, including Suite2p[4] and CAIMAN[5], often use a greedy cell finding step before performing alternating estimation steps to estimate cells' $Ca^{2+}$ activity traces and spatial profiles jointly. This initial step is crucial due to the non-convex nature of the movie reconstruction problem, as the initialization routine plays an important role to arrive at a 'good' local optimum. However, the greedy cell finding process inevitably suffers from non-Gaussian contaminants, since neighboring cells' signals contribute to the noise during the greedy initialization of the current cell's spatial footprint. Thus, the use of robust loss, as discussed in **Section 3.2** above, is needed to account for non-Gaussian noise contaminants. Therefore, though we still use a greedy cell finding step, EXTRACT uses the robust loss function during the cell finding module (**Methods**).

Cell refinement iterations and quality checks: The bi-convex nature of the movie reconstruction loss (See Eq. (6)) allows finding fast solutions by performing iterative (non-negative) least-squares estimates in the case of $L_2$ estimators such as Suite2p[4] and CAIMAN[5]. In prior algorithms, the quality checks on the refined components are often performed at the end of these iterations, and spurious cells are discarded. EXTRACT improves on this process in two distinct ways. First, instead of the $L_2$ loss function, we minimized the 1-sided Huber loss function (**Methods**), which turned the least-squares estimates into robust regression solutions. Second, we added quality checks in between the iterations, discarding spurious cells gradually and allowing the spatial footprints of true cells to adjust in-between the steps. We call this new and improved process of alternating iterations as 'cell refinement.'

*Non-negativity condition:* The non-negativity constraint on the $Ca^{2+}$ activity traces is motivated by the physics of $Ca^{2+}$ imaging experiments and the biology of the $Ca^{2+}$ indicators. As discussed in **Section 4** above, when the assumptions of non-negativity are met in the statistical model of the $Ca^{2+}$ imaging movie generation, the non-negative estimates outperform their unconstrained counterparts. Motivated by this observation, for EXTRACT, we use the non-negativity constraint when performing the robust regression, but supply it with an adaptive cell baseline estimators, which ensure that the cells' baselines (below which the non-negativity is to be enforced) are correctly estimated from the data (**Methods, Supplementary Note 2**).

*Exponential kernel fitting:* As discussed in **Section 4**, least-squares estimates can have very high variance, especially when cells in the FOV are moderately overlapping. To mitigate the variance, current approaches often enforce exponentially decaying kernels for the estimated $Ca^{2+}$ activity traces[4,5]. However, this assumption may sometimes be too restrictive, especially when the cells' fire persistently in prolonged durations. When designing EXTRACT, since we do not perform least-squares estimates and non-negativity constraint significantly decreases the variance (**Appendix Figure 1**), we opted not to use this restrictive design choice.

*Background modeling and/or subtraction:* Previous work incorporated background subtraction and/or modeling routines[3–5] to remove and/or account for neuropil contamination. This takes, often, the form of subtracting a fraction of the activity in the pixels[4] surrounding the cell or in a ring of pixels with a relatively large radius[3], or the explicit modeling of background with low-rank[2] or high-rank[13] components. However, there is no theoretical guarantee that such heuristic approaches would not remove cells' signals alongside the neuropil contaminations. In fact, in our experiments, we observed that replacing the $Ca^{2+}$ activity traces obtained by CAIMAN with the non-negative least-squares estimates increased their quality significantly (**Figures 3E-F** and **S5**), which can be attributed to overfitting due to extra assumptions of CAIMAN including the background modeling. In contrast, as we have shown in **Sections 5.3** and **6.6** above, spatial high-pass filtering can remove spatially correlated neuropil signals without harming cells’ signal thanks to the spatial separation of scales between a typical cell’s radius and global background contaminations. Thus, in EXTRACT, we replaced explicit background modeling with two implicit components: a mild high-pass filtering that removes global contaminants without affecting cells’ $Ca^{2+}$ activity traces and an agnostic robust regression approach that can handle local non-Gaussian contaminants.

In summary, when designing and validating EXTRACT, we first put the existing key heuristics in the field in an estimation theory framework, which allowed us to first theoretically and next empirically test and validate them. Those that passed our validation tests were incorporated into EXTRACT, with additional advantages brought forth by the use of the robust loss function in every step. The theoretical framework we introduced and developed in this supplementary note shall be helpful when designing new and improved computational analysis pipelines supporting systems neuroscience research.

### Supplementary Note 2: “Mathematical derivations for EXTRACT’s custom solver”

### §1. Overview.

Modeling real-world data often requires the ability to tolerate deviations from the expected statistical model of data generation. In practice, this requires mitigating risks associated with several statistical models, not just the one that provides the best fit. Minimax estimators, *i.e.*, those that minimize the maximum risk in a broad class of potential estimators, are trained to perform well under the worst-case scenario, hence are well suited to handle real-world data[1]. Yet, despite its proven effectiveness and efficiency over the last half century, the application of the robust minimax estimation to the problem of cell extraction from $Ca^{2+}$ imaging movies has largely been missing. The sole exception is perhaps our conference paper from seven years ago[2], which outlined our initial ideas for how robust regression models can contribute to solving the cell extraction problem, albeit did not address one of the most important questions: How can one convert a proof-of-concept study using controlled simulations into a scalable, interpretable, and generalizable algorithm that can handle complex non-idealities in real-data applications?

To address this question, this note introduces the one-sided Huber loss function under conditions of inhomogeneous contamination (across movie frames and pixels) and gives a geometric picture for the resulting robust solution (**§2**). Then, **§3** provides mathematical details for the optimization procedures of EXTRACT’s fast, scalable, and robust solver. **§4** discusses the unique algorithmic tricks introduced in our implementation to speed EXTRACT and minimize its dependence on RAM and GPU memory. Finally, **§5** concludes with a summary of design features that are unique to EXTRACT, which may help guide future work developing fast and scalable software implementations for systems neuroscience.

### §2. Robust statistics for extracting cells from $Ca^{2+}$ imaging movies.

The starting point of our investigations is the intuition that many contamination signals, such as from out-of-focus cells and/or neuropil $Ca^{2+}$ dynamics, lead to positive-going fluorescence fluctuations from $Ca^{2+}$ indicators. If the movie reconstruction problem, introduced in **Supplementary Note 1**, does not preferentially allow for large positive values in the reconstruction errors, these fluctuations will appear within the cells' estimated $Ca^{2+}$ activity traces, leading to crosstalk effects as documented in our benchmarks (**Figures 3A** and **S2**) and prior literature[3,4]. Consequently, demixing of signal and noise sources in $Ca^{2+}$ movies should benefit from a loss function with an asymmetric valuation of positive and negative errors.

Our prior conference paper[2] showed that use of the one-sided Huber loss, instead of its traditional, two-sided symmetric counterpart[1], led to superior estimated traces under a family of statistical models, in which part of the noise is sampled from a Gaussian distribution and the rest from a family of non-negative distributions. The latter part of the noise model was designed to accommodate positive outlier patterns, such as neuropil activation, in the $Ca^{2+}$ imaging datasets. While Ref.2 provided a proof-of-concept study for the application of robust statistics to the cell extraction problem, theoretical (*e.g.*, how to pick the robustness parameter that defines the specific family of interest) and practical (*e.g.*, developing a fast solver suited for $Ca^{2+}$ imaging datasets) limitations prevented the development of a full pipeline.

A first step towards overcoming these issues is to understand and interpret the inner workings of the (unconstrained) robust solver. To achieve this, this section provides a detailed account of the unconstrained robust regression problem by developing a quantitative measure for outlier detection and a qualitative, geometric understanding of the resulting outlier mitigation effects. Building on the theory developed here, **§3** and **§4** will discuss our efforts to overcome the aforementioned limitations via our theoretical and algorithmic advances, respectively.

#### 2.1. Introduction of the robust movie reconstruction problem

In **Supplementary Note 1**, we introduced the movie reconstruction paradigm, in which a movie, $\boldsymbol{M}$, is assumed to consist of $Ca^{2+}$ activity sources with spatial profiles, $\boldsymbol{S}$, and $Ca^{2+}$ activity traces,

$\boldsymbol{T}$, and unknown noise components, $\boldsymbol{\sigma}$. Then, cell extraction is achieved by estimating a reconstructed movie, $\hat{\boldsymbol{M}} = \hat{\boldsymbol{S}}\hat{\boldsymbol{T}}$, consisting of the cells' $Ca^{2+}$ activity signals. The pairs, $\hat{\boldsymbol{S}}$ and $\hat{\boldsymbol{T}}$, are jointly optimized to minimize a movie reconstruction loss such that:

$$(\boldsymbol{T}^*, \boldsymbol{S}^*) \in \arg\min_{\hat{\boldsymbol{S}},\hat{\boldsymbol{T}}} \mathcal{L}(\boldsymbol{M}, \hat{\boldsymbol{M}}),$$

where $(\boldsymbol{T}^*, \boldsymbol{S}^*)$ stands for an optimal pair that minimizes the loss function $\mathcal{L}$.

In **Supplementary Note 1**, we reviewed prior cell extraction pipelines, which often relied on minimizing the $L_2$ loss function. EXTRACT, building on the prior work on robust statistics[1,2], replaces the $L_2$ loss function with the one-sided Huber loss function, denoted as $\rho_{\boldsymbol{\kappa}}$:

$$\mathcal{L}_{robust}(\boldsymbol{M}, \hat{\boldsymbol{M}}) = \sum_{i=1}^{n_{pixels}} \sum_{j=1}^{n_{times}} \rho_{\boldsymbol{\kappa}_{ij}}(\boldsymbol{M}_{ij} - \sum_{k=1}^{n_{cells}} \hat{\boldsymbol{S}}_{ik}\hat{\boldsymbol{T}}_{kj}), \quad (1)$$

where $n_{pixels}$ is the number of pixels in the movie, $n_{times}$ is the number of movie frames, and the one-sided Huber loss function, $\rho_{\kappa}(.)$, is defined as:

$$\rho_y(x) = \{(x^2 + y^2)/2 \quad if \quad x \le y; \quad xy \quad if \quad x > y\} \quad ,$$

which is differentiable, but does not have a well-defined Hessian as it is not twice differentiable.

In Eq. (1), $\boldsymbol{\kappa}_{ij}$ is a parameter that defines the severity of non-Gaussian noise in the $i$th pixel during the $j$th movie frame. Notably, this is a substantially different noise model than the one assumed in prior work[2], in which the contamination level was assumed to be constant across all pixels and movie frames.

With the introduction of the convex robust loss function, Eq. (1) has the desirable property of bi-convexity, which means the estimation of cells' $Ca^{2+}$ activity traces, $\hat{\boldsymbol{T}}$, is convex when $\hat{\boldsymbol{S}}$ is held fixed and *vice versa*. In such cases, another desirable property of Eq. (1) arises: The loss minimization can be divided into several independent problems, which can be solved in a scalable and fast manner owing to parallelization across (multiple) graphics processing units (GPUs). Taken together, these facts allow the implementation of a fast convex routine, analogous to alternating (non-negative) least-squares[5], which we detail below.

To illustrate how the loss function can be divided into independent components, consider the case when $\hat{\boldsymbol{S}}$ is held fixed and $\hat{\boldsymbol{T}}$ is sought. Then, we have:

$$\mathcal{L}_{robust}(\boldsymbol{T}, \hat{\boldsymbol{T}}) = \sum_{j=1}^{n_{frames}} \left[ \sum_{i=1}^{n_{pixels}} \rho_{\boldsymbol{\kappa}_{ij}} (\boldsymbol{M}_{ij} - \sum_{k=1}^{n_{cells}} \hat{\boldsymbol{S}}_{ik} \hat{\boldsymbol{T}}_{kj}) \right] = \sum_{j=1}^{n_{frames}} \mathcal{L}_j(\boldsymbol{T}, \hat{\boldsymbol{T}}^{(j)}), \tag{2}$$

where $\hat{\boldsymbol{T}}^{(j)}$ refers to the cells' estimated $Ca^{2+}$ activity traces during the $j$th frame of the movie and $\mathcal{L}_j(\boldsymbol{T}, \hat{\boldsymbol{T}}^{(j)})$ is a term in the loss function that contains the only dependence of $\hat{\boldsymbol{T}}^{(j)}$ and does not depend on any other $\hat{\boldsymbol{T}}^{(l)}$ for $l \neq j$. A similar discussion for the case when $\hat{\boldsymbol{T}}$ is held fixed reveals that the estimation of $\hat{\boldsymbol{S}}$ can be divided into independent components for each movie pixel.

Owing to the inherent parallelization properties of alternating robust regressions, we can, without loss of generality, consider a general regression problem of the form:

$$\mathcal{L}_{robust}(\boldsymbol{Y}, \boldsymbol{\beta}) = \sum_{i=1}^{n_{samples}} \rho_{\boldsymbol{\kappa}_i} (\boldsymbol{Y}_i - \sum_{k=1}^{n_{cells}} \boldsymbol{X}_{ik} \boldsymbol{\beta}_k). \tag{3}$$

Here, when $\hat{\boldsymbol{S}}$ is held fixed, $\boldsymbol{Y}_i$ is the activity of the $i$th pixel, $\boldsymbol{X}$ corresponds to $\hat{\boldsymbol{S}}$, $\boldsymbol{\beta}$ refers to a column of $\hat{\boldsymbol{T}}$, and $\mathcal{L}_{robust}(\boldsymbol{Y}, \boldsymbol{\beta})$ is equivalent to $\mathcal{L}_j(\boldsymbol{T}, \hat{\boldsymbol{T}}^{(j)})$.

As we discuss next, $\mathcal{L}_{robust}(\boldsymbol{Y}, \boldsymbol{\beta})$ can be solved with fixed-point updates (**Algorithm 1 in Methods**), which can be vectorized to solve jointly all loss components in Eq. (2). The same arguments hold for the case when $\hat{\boldsymbol{T}}$ is held fixed, hence without the loss of generality we will explicitly focus on Eq. (3) for the rest of this note.

### 2.2. Minimization of the one-sided Huber loss function

Though the robust loss function in Eq (3) is convex with respect to $\boldsymbol{\beta}$, due to the fact that the Hessian of ρ is ill-defined (See the discussion in **Section 2.1** and **Methods**), even the simplest form of the optimization problem, *i.e.*, without any additional constraints, requires a closer look. We do this here in this section and defer discussion on minimizing the robust loss under additional constraints, such as the non-negativity of $Ca^{2+}$ activity traces, to **Section §3**.

We start by computing the gradient of the loss function in Eq. (3), which is as follows:

$$\frac{\partial \mathcal{L}_{robust}(\boldsymbol{Y},\boldsymbol{\beta})}{\partial \boldsymbol{\beta}_k} = \sum_i [\boldsymbol{X}_{ik}\boldsymbol{U}_i \Theta(\boldsymbol{U}_i > 0)] + \sum_{i,j} [\boldsymbol{X}_{ik}\boldsymbol{X}_{ij}\boldsymbol{\beta}_j - \boldsymbol{X}_{ik}\boldsymbol{Y}_i] \ ,$$

where we define $\Theta(.)$ as the Heaviside function with $\Theta(x > 0) = 1$, and zero otherwise. Here, we define $\boldsymbol{U}_i = \boldsymbol{Y}_i - \sum_j \boldsymbol{X}_{ij}\boldsymbol{\beta}_j - \boldsymbol{\kappa}_i$ as the outlier detection variable, which we will expand on later below. As the second step, we calculate the Hessian:

$$\frac{\partial^2 \mathcal{L}_{robust}(\boldsymbol{Y},\boldsymbol{\beta})}{\partial \boldsymbol{\beta}_k \partial \boldsymbol{\beta}_l} = \sum_i \boldsymbol{X}_{ik}\boldsymbol{X}_{il}\Theta(\boldsymbol{U}_i \leq 0) = \sum_i \boldsymbol{X}_{ik}\boldsymbol{X}_{il} - \sum_{i:\, \boldsymbol{U}_i > 0} \boldsymbol{X}_{ik}\boldsymbol{X}_{il}$$

The fixed-point solver introduced in the main text (**Algorithm 1** in **Methods**) starts with the observation that $\boldsymbol{X}^T\boldsymbol{X}$ is positive semi-definite, thus $\Delta\boldsymbol{\beta}_{fixed-point} = -(\boldsymbol{X}^T\boldsymbol{X})^{-1}\nabla\mathcal{L}_{robust}(\boldsymbol{Y},\boldsymbol{\beta})$ would be a descent direction for minimization of the loss function in Eq. (3). Moreover, since the Hessian calculated above is approximately $\boldsymbol{X}^T\boldsymbol{X}$, with a correction involving only the outlier samples, this descent direction is expected to be close to the Newton's step $\Delta\boldsymbol{\beta}_{Newton}$. Fortunately, unlike the Newton's step that requires explicit calculation of the Hessian at every iteration due to the dependence of $\boldsymbol{U}$ on $\boldsymbol{\beta}$, $\boldsymbol{X}^T\boldsymbol{X}$ can be precomputed at the beginning of the optimization and used throughout, hence the name 'fixed-point'. Explicitly calculating

$\Delta\boldsymbol{\beta}_{fixed-point}$ results in the update rule:

$$\boldsymbol{\beta}^{(t+1)} := \boldsymbol{\beta}^{(t)} - (\boldsymbol{X}^T\boldsymbol{X})^{-1}\nabla\mathcal{L}_{robust}(\boldsymbol{Y},\boldsymbol{\beta}) = \boldsymbol{\beta}_{ls} - (\boldsymbol{X}^T\boldsymbol{X})^{-1}\boldsymbol{X}^T max(\boldsymbol{U}, 0) \tag{4}$$

where we define the least-squares estimate as $\boldsymbol{\beta}_{ls} = (\boldsymbol{X}^T\boldsymbol{X})^{-1}\boldsymbol{X}^T\boldsymbol{Y}$ .

Even though Eq. (4) resembles the update rule derived in our prior work[2], it is distinct in the sense that $\boldsymbol{\kappa}$ is a vector with nonhomogenous values and enters the equation through $\boldsymbol{U}$. Yet, since both update rules share the same form, our solver in Eq. (4) naturally inherits the convergence properties developed in the prior work[2], *i.e.*, is practically a second-order method in convergence but has computational complexity comparable to the gradient descent.

#### 2.3. Automated outlier detection

The update rule in Eq. (4) provides an intuitive geometrical interpretation for how the use of $\boldsymbol{X}^T\boldsymbol{X}$ leads to an automated outlier detection, which we briefly discussed in the main text (**Figure 1H-L**). We provide more details here and start by restating the outlier detection variable:

$$\boldsymbol{U} = \boldsymbol{Y} - \boldsymbol{X}\boldsymbol{\beta} - \boldsymbol{\kappa} \quad , \tag{5}$$

which affects the parameter update in Eq. (4) only for outlier samples, for which $\boldsymbol{U}_i$>0. If such points exist, the optimal estimate, $\boldsymbol{\beta}^*$, differs from $\boldsymbol{\beta}_{ls}$ by a small outlier correction.

To study the implications of this outlier correction, we first note $\Delta\boldsymbol{Y} = \boldsymbol{Y} - \boldsymbol{X}\boldsymbol{\beta}$, the residual error for the movie reconstruction. A negative residual points to a negative noise component, which can only be due to the Gaussian noise, and not the outliers, by the assumed model (**Methods**), implied by the quadratic form of the loss function for negative error values (**Figure 1F**). A positive error in the movie reconstruction loss, $\Delta\boldsymbol{Y}_i > 0$, can potentially stem from two sources: the random Gaussian noise, or the arbitrary non-Gaussian contaminant, that is non-zero only for $[\boldsymbol{\kappa}_i, \infty)$. Rewriting Eq. (5) as $\boldsymbol{U} = \Delta\boldsymbol{Y} - \boldsymbol{\kappa}$, we observe that $\boldsymbol{U}_i > 0$ if and only if $\Delta\boldsymbol{Y}_i > \boldsymbol{\kappa}_i$, *i.e.*, when the residual lies within the support of the non-Gaussian contaminant. Therefore, by identifying samples for which $\boldsymbol{U}_i > 0$, one can detect, without human intervention, *potential outlier samples,* in which the fluctuations in the movie reconstruction residual error, $\Delta\boldsymbol{Y}$, are large enough to have the possibility of originating from non-Gaussian contamination. This is similar to the winsorizing cutoffs[6], in which deviations beyond a fixed threshold are projected down to the threshold, but is distinct in the sense that the cutoff is actively learned for each sample and updated through the iterations of the robust regression.

Equation (4) provides further insights into how the detected outliers affect the optimization problem. First, we rewrite Eq. (4) assuming convergence $\boldsymbol{\beta}^{(t)} \rightarrow \boldsymbol{\beta}^*$:

$$\boldsymbol{\beta}^* := (\boldsymbol{X}^T\boldsymbol{X})^{-1}\boldsymbol{X}^T[\boldsymbol{Y} - max(\boldsymbol{U}^*, 0)] \quad (6),$$

where $\boldsymbol{U}^* = \boldsymbol{Y} - \boldsymbol{X}\boldsymbol{\beta}^* - \boldsymbol{\kappa}$. This is a least-squares solution on a projected movie $\boldsymbol{Y}^* = \boldsymbol{Y} - max(\boldsymbol{U}^*, 0)$, in which the robust loss *projects* (not removes) the outlier points to the detection margin. In the limit $\boldsymbol{\kappa} \rightarrow \infty$, *i.e.*, the movie is assumed to contain only Gaussian noise, no sample is considered to be an outlier, and Eq. (6) simplifies to the least-squares solution. This observation explains why $\boldsymbol{\beta}_{ls}$ is a good initialization for the robust solver (**Algorithm 1** in the main text) and $\boldsymbol{X}^T\boldsymbol{X}$ is a proper approximation for the Hessian of the loss function in Eq. (3).

#### 2.4. Geometric interpretation of the robust statistics

While we motivated the outlier detection procedure for the case of the converged solution in the previous section, the interpretation is in fact more general. Specifically, we can divide the algebra in Eq. (4) into three interpretable steps:

1) For a given $\boldsymbol{\beta}^{(t)}$, compute the outlier detector: $\boldsymbol{U}^{(t)} := \boldsymbol{Y} - \boldsymbol{X}\boldsymbol{\beta}^{(t)} - \boldsymbol{\kappa}$.

2) Obtain the projected movie: $\boldsymbol{Y}^{(t)} = \boldsymbol{Y} - max\,(0, \boldsymbol{U}^{(t)})$.

3) Estimate the least squares solution on the projected movie: $\boldsymbol{\beta}^{(t+1)} = (\boldsymbol{X}^T\boldsymbol{X})^{-1}\boldsymbol{X}^T\boldsymbol{Y}^{(t)}$.

These three steps can be performed iteratively until convergence, equivalently to the updates performed in Eq. (4). Thus, minimization of the robust loss through the fixed-point solver amounts to iterative application of data-driven outlier detection, subtraction of potential contamination from the original movie, and then performance of a subsequent least-squares estimate. Here, the updates are performed through iterative applications of least-squares and margin-based outlier detection, and no assumption is made about the outliers other than that they cause unwanted crosstalk that cannot be explained *by the $Ca^{2+}$ activity traces of the extracted cells*. This process is distinct from the crosstalk removal performed by the least-squares estimate. Least-squares necessitates that all background components are either extracted, subtracted, or explicitly modeled[7], whereas robust regression can remove crosstalk from unmodelled background.

The three interpretable steps introduced here provide the mathematical background for our discussion in **Figure 1H-L**, which illustrates the consequences of applying robust regression specifically to the estimation of cells' spatial footprints when $Ca^{2+}$ activity traces are held fixed and *vice versa*.

**§3. Algorithmic details for the optimization of EXTRACT's fast and scalable solver.**

Here, we introduce the algorithmic details of our custom-written fixed-point solver, discussed as **Algorithm 1** in the main text, which has been heavily optimized compared to our prior design[2]. Specifically, the new solver (i) has the ability to enforce non-negativity, locality, and L1-regularization using a novel application of the alternating direction method of multipliers

(ADMM)[8], (ii) incorporates new adaptive estimation procedures to update $\boldsymbol{\kappa}$ and cells' baseline activities, and (iii) can now adaptively account for the GPU and RAM memory requirements. In this section, we provide mathematical details for the first two aspects: ADMM methodology, which incorporates necessary constraints for cell extraction from $Ca^{2+}$ imaging movies, and how the robustness parameter, $\boldsymbol{\kappa}$, and the adaptive cell baselines are estimated from the $Ca^{2+}$ imaging movies.

#### 3.1. Alternating direction method of multipliers (ADMM)

In this section, we review the main ideas behind the ADMM framework[8], which can be utilized to minimize an optimization problem of the following form:

$$\begin{aligned} \text{minimize} \quad & \mathcal{L}_{robust}(\boldsymbol{Y}, \boldsymbol{\beta}) = \sum_{i=1}^{n_{samples}} \rho_{\boldsymbol{\kappa}_i}(\boldsymbol{Y}_i - \sum_{j \in cells} \boldsymbol{X}_{ij} \boldsymbol{\beta}_j) \\ \text{subject to} \quad & \boldsymbol{\beta} \geq 0, \ \forall j, \ \forall i \in O_j : \boldsymbol{\beta}_i = 0 \end{aligned} \tag{7}$$

Here, $\boldsymbol{\beta}$ is the variable of interest, corresponding to either $\hat{\boldsymbol{T}}$ or $\hat{\boldsymbol{S}}$, depending on the cell extraction subroutine (**Figure 2**). $\boldsymbol{X}$ is the remaining variable that is fixed for the duration of the optimization subroutine. $\boldsymbol{Y}$ either corresponds to a single pixel (when estimating $\hat{\boldsymbol{S}}$) or to a single frame (when estimating $\hat{\boldsymbol{T}}$) of the movie $\boldsymbol{M}$. $O_j$ is a set and denotes the pixels that cannot be occupied by the $j$th cell, *i.e.*, enforces locality (or other similar) constraints if desired.

The non-negativity condition, $\boldsymbol{\beta} \geq 0$, (or an alternative version with estimated cell baselines, see below) may be used in both cases. The condition $\forall j, \ \forall i \in O_j : \boldsymbol{\beta}_i = 0$ enforces locality constraints and is primarily used when estimating $\hat{\boldsymbol{S}}$. We note that one can always set $O = \emptyset$, which means Eq. (7) can be considered a general problem of interest for cell extraction, which we refer to as "the constrained non-negative robust estimation." We can minimize this problem by transforming Eq. (7) with the addition of a dummy variable:

$$\begin{aligned} \text{minimize} \quad & \mathcal{L}_{robust}(\boldsymbol{Y}, \boldsymbol{\beta}) \; + \; g(\boldsymbol{\gamma}) \quad , \\ \text{subject to} \quad & \boldsymbol{\gamma} = \boldsymbol{\beta} \end{aligned}$$

where $g(\boldsymbol{\gamma})$ is an indicator function that is zero for data points for which $\boldsymbol{\gamma} \geq 0,\ \forall j,\ \forall i \in O_j$ : $\boldsymbol{\gamma}_i = 0$ and $\infty$ otherwise. In other words, the dummy variable $\boldsymbol{\gamma}$ has to satisfy the original constraints in Eq. (7) and has to be equal to $\boldsymbol{\beta}$. The new transformed equation can be solved via the ADMM updates, which are derived in Eqs. (3.5-3.7) of Ref. 8:

$$\begin{aligned} \boldsymbol{\beta}^{(k+1)} &:= \arg\min_{\boldsymbol{\beta}} \left(\mathcal{L}_{robust}(\boldsymbol{Y}, \boldsymbol{\beta}) \; + \; (\chi/2)||\boldsymbol{\beta} - \boldsymbol{\gamma}^{(k)} + \boldsymbol{z}^{(k)}||_2^2\right), \\ \boldsymbol{\gamma}^{(k+1)} &:= \arg\min_{\boldsymbol{\gamma}} \left(g(\boldsymbol{\gamma}) \; + \; (\chi/2)||\boldsymbol{\beta}^{(k+1)} - \boldsymbol{\gamma} + \boldsymbol{z}^{(k)}||_2^2\right), \\ \boldsymbol{z}^{(k+1)} &:= \boldsymbol{z}^{(k)} + \boldsymbol{\beta}^{(k+1)} - \boldsymbol{\gamma}^{(k+1)}. \end{aligned}$$

Here, $\boldsymbol{z} \in \mathbb{R}^{n_{cells}}$ is called as the 'dual variable' and $\chi$ is a user-defined parameter of optimization. For EXTRACT, we define $\chi$ automatically using the eigenvalues of $\boldsymbol{X}^T\boldsymbol{X}$, the reason for which will be clear below.

The advantage of the ADMM approach can be best understood by a closer inspection of the final and second steps. The final step is a running estimate of the error in the constraints and can be calculated in a single step, whereas the second step is minimized through the projection:

$$\boldsymbol{\gamma}^{(k+1)} = \Pi_{g(\boldsymbol{\gamma})}(\boldsymbol{\beta}^{(k+1)} \; + \; \boldsymbol{z}^{(k)}),$$

where $\Pi_{g(\boldsymbol{\gamma})}(\boldsymbol{\gamma})$ performs an orthogonal projection of $\boldsymbol{\gamma}$ to the set $g(\boldsymbol{\gamma}) \neq \infty$. Fortunately, our constraints in Eq. (7) define a straightforward projection, and thus, the only step that requires a real optimization routine is the first one.

As we show below, the first optimization step of the ADMM can be solved using the fixed-point solver derived in **§2** above for the unconstrained minimization scenario. Therefore, by transforming the original problem into a simpler step-by-step approach, ADMM provides access to the speed and scalability of the fixed-point solver, which otherwise cannot minimize the constrained non-negative robust estimation problem in Eq. (7).

#### 3.2. Solving non-negative constrained robust estimation

Now, we derive the update rule of the first ADMM equation, which requires minimizing the loss:

$$\mathcal{L}_{ADMM}(\boldsymbol{\beta}) = \sum_{i=1}^{n_{samples}} \rho_{\boldsymbol{\kappa}_i}(\boldsymbol{Y}_i - \sum_{j \in cells} \boldsymbol{X}_{ij}\boldsymbol{\beta}_j) + \lambda_1||\boldsymbol{\beta}||_1 + (\lambda_2/2)||\boldsymbol{\beta}||_2^2 + (\chi/2)||\boldsymbol{\beta} - \boldsymbol{v}||_2^2 ,$$

where we define the variable $\boldsymbol{v} = \boldsymbol{\gamma}^{(k)} - \boldsymbol{z}^{(k)}$. As before, we first compute the gradient:

$$\nabla_{\boldsymbol{\beta}}\mathcal{L}_{ADMM}(\boldsymbol{\beta}) = \boldsymbol{X}^T max(\boldsymbol{U}, 0) + (\boldsymbol{X}^T\boldsymbol{X} + I[\lambda_2 + \chi])\,\boldsymbol{\beta} - \boldsymbol{X}^T\boldsymbol{Y} + \lambda_1 sgn(\boldsymbol{\beta}) - \chi\boldsymbol{v} ,$$

where the outlier detection variable, $\boldsymbol{U}$, is defined as in Eq. (5). Next, we compute the Hessian:

$$\frac{\partial^2\mathcal{L}_{ADMM}(\boldsymbol{\beta})}{\partial\boldsymbol{\beta}_k\partial\boldsymbol{\beta}_l} = \sum_{i \in samples} [\boldsymbol{X}_{ik}\boldsymbol{X}_{il} + \boldsymbol{I}_{kl}[\lambda_2 + \chi]] - \sum_{i:\, \boldsymbol{U}_i > 0} \boldsymbol{X}_{ik}\boldsymbol{X}_{il} .$$

We follow our steps from **§2**, and approximate the Hessian with $\boldsymbol{X}^T\boldsymbol{X} + \boldsymbol{I}[\lambda_2 + \chi]$, which leads to the update rule:

$$\boldsymbol{\beta}^{(t+1)} := \boldsymbol{\beta}_{rls} - (\boldsymbol{X}^T\boldsymbol{X} + \boldsymbol{I}[\lambda_2 + \chi])^{-1}[\boldsymbol{X}^T max(\boldsymbol{U}, 0) + \lambda_1 sgn(\boldsymbol{\beta}^{(t)}) + \chi(\boldsymbol{z}^{(k)} - \boldsymbol{\gamma}^{(k)}), \quad (8)$$

where we define the regularized least-squares solution as $\boldsymbol{\beta}_{rls} = (\boldsymbol{X}^T\boldsymbol{X} + \boldsymbol{I}[\lambda_2 + \chi])^{-1}\boldsymbol{X}^T\boldsymbol{Y}$. This update rule has the same form as the fixed-point solver (**Algorithm 1**) discussed in **§2**, and therefore inherits its convergence, speed, and scalability properties. Specifically, the fixed-point solver now includes $L_1$- and $L_2$-regularization terms and an additional consensus term,

$\chi(\boldsymbol{z}^{(k)} - \boldsymbol{\gamma}^{(k)})$, that enforces the convergence between the different ADMM steps. For $\chi = \lambda_1 = \lambda_2 = 0$, this solver reproduces the unconstrained and unregularized version in Eq. (4). Although we have provided a general form for the fixed-point solver, in almost all cases we found neither a need for nor a benefit from regularization of the cell extraction routines in the EXTRACT pipeline (*i.e.*, $\lambda_1 = \lambda_2 = 0$ by default). An exception was the case with highly overlapping cells, in which the L1-regularized approach during cell refinement helped prevent duplicate cell filters.

Our derivations so far have focused on the scenario where $\boldsymbol{\beta}$ is a vector, *i.e.*., solving the optimization problem for a single frame or a pixel. Fortunately, the ADMM solver above can be extended to cover the full movie by promoting $\boldsymbol{\beta}$ to a two-dimensional matrix. In other words, the optimization process does not need to be parallelized, but can be vectorized. Consequently, the ADMM approach used here is readily compatible with a GPU implementation for fast matrix multiplications.

#### 3.3. Adaptive estimation of the robustness margin

To allow practical applications of robust regression to real datasets, we need to model how $\boldsymbol{\kappa}$ depends on the local contamination in a spatiotemporal patch of the movie. This need is rooted in the reality of $Ca^{2+}$ imaging datasets, in which neuropil activity (and other contaminants) affect cell extraction in a spatiotemporally varying manner. When the ground truth $\boldsymbol{\kappa}$ varies across the movie, any arbitrary homogeneous choice for $\boldsymbol{\kappa}$ (as is done in Ref.2) can introduce an undesirable bias into the estimation procedure. To account for the spatiotemporal variations, we created an adaptive estimation procedure to jointly estimate the regression parameter $\boldsymbol{\beta}$ and $\boldsymbol{\kappa}$ from the movie, where the matrix $\boldsymbol{\kappa}^{(q)}$ contains the estimates for each cell and time point after $q$ iterations.

Our recipe is as follows: We start with a user-defined initial value of $\boldsymbol{\kappa}^{(0)}$ (by default, 0.7 s.d. of the median pixel activities across the field of view). We perform an adaptive estimation of the contamination level $\boldsymbol{\epsilon}^{(q-1)}$, which is performed on a frame-by-frame basis for each cell given the current estimates $\boldsymbol{\kappa}^{(q-1)}$ and $\boldsymbol{\beta}^{(q-1)}$. Inside the main iteration loop for robust estimation, we estimate the bias in estimated $\boldsymbol{\beta}^{(q-1)}$, which depends on the current estimate $\boldsymbol{\kappa}^{(q-1)}$. Then, we make iterative updates to the robustness parameter, $\boldsymbol{\kappa}^{(q-1)} \rightarrow \boldsymbol{\kappa}^{(q)}$, to minimize the bias in the estimation of the true parameter, $E[\boldsymbol{\beta}^{(q-1)} - \boldsymbol{\beta}^{*}]$, which is driven to zero as the number of iterations grows. The outputs consist of a $[n_{cell} \times n_t]$ matrix $\boldsymbol{\epsilon}^{(q)}$ and the updated estimates of the robustness margin $\boldsymbol{\kappa}^{(q)}$, which are linked element-wise through a functional relationship $\boldsymbol{\kappa}^{(q)} = h(\boldsymbol{\epsilon}^{(q)})$, as we show below.

For simplicity of presentation, we first discuss the estimation of $\boldsymbol{\kappa}^{(q)}$ on a single frame and a single cell, *i.e.*, a single entry of the matrix $\boldsymbol{\kappa}^{(q)}$. Therefore, we will refer to scalar values from now on, $\kappa^{(q)}$ and $\epsilon^{(q)}$, as the individual entries of interest. The estimation process, as before, can be vectorized to estimate $\boldsymbol{\kappa}^{(q)}$ across the full movie and all cells.

#### 3.3.1. Relating robustness margin to contamination level

We start by defining the function $f(x) = x\,\Phi(x) + \varphi(x)$, where $\Phi(x)$ and $\varphi(x)$ are the cumulative distribution and probability density functions of a standard normal distribution, respectively. We use this function to restate the relationship, derived in the main paper (**Proposition 1**), between $\kappa$ and $\epsilon$ as follows:

$$\Phi(\kappa) + \frac{\varphi(\kappa)}{\kappa} = \frac{1}{1-\epsilon} \Leftrightarrow 1 - \epsilon = \frac{\kappa}{f(\kappa)}\,. \tag{9}$$

Here, we can easily check the consistency of Eq. (9) for limiting cases. Specifically, as $\kappa \rightarrow \infty$,

$\frac{\kappa}{f(\kappa)} \rightarrow \frac{1}{\Phi(\kappa)} \rightarrow 1$ and thus $\epsilon \rightarrow 0$. For $\kappa = 0$, we find $\epsilon = 1$ as expected. It is worth noting that Eq. (9) provides a simple formula to compute $\epsilon$ given $\kappa$, but the inverse of the function (even though it exists, see **Figure S1F**) does not have an immediate closed form. Thus, we use a fitted function, which includes polynomials and a logarithmic function, to approximate the converse relationship, $\kappa \approx h(\epsilon)$.

#### *3.3.2. Using distributional asymmetry to link bias with positive residuals*

We now note the asymmetry in the probability distributions of the statistical model. Due to $\epsilon$ probability of having a non-Gaussian noise component, there is an asymmetric distribution in the residuals obtained as part of the regression problem. We will exploit this fact to link bias with an easily computable observable: fraction of positive residuals. We start by recalling that the statistical model of the movie (for a given time point and pixel) follows $y = \boldsymbol{x}^T \boldsymbol{\beta}^* + \sigma$, where $\sigma$ denotes the noise component, $\boldsymbol{x}$ is the cells' spatial footprint at the pixel of interest and $\boldsymbol{\beta}^{(q-1)}$ is the estimated $Ca^{2+}$ activity traces at the given time point. As we show below, a mismatch in the estimated contamination level, $\kappa^{(q)} \neq \kappa$, leads to an inevitable bias in the estimation of the $Ca^{2+}$ activity traces such that $b = \boldsymbol{x}^T [E[\boldsymbol{\beta}^{(q-1)}] - \boldsymbol{\beta}^*] \neq 0$.

Practically, it is not possible to compute $E[\boldsymbol{\beta}^{(q-1)}]$, which would entail resampling the movie with different initializations of the random noise from the statistical noise model. Instead, we only have access to $\boldsymbol{\beta}^{(q-1)}$, the estimate for a given realization of the movie, *i.e.*, the recorded movie. Here, we recall from **Supplementary Note 1** that the $Ca^{2+}$ activity traces are traditionally estimated from multiple pixels and thus have negligible variance compared to the movie noise, scaling as $O(\sigma/n_{pixels})$. Consequently, for the rest of this calculation, we approximate $E[\boldsymbol{\beta}^{(q-1)}] \approx \boldsymbol{\beta}^{(q-1)}$, *i.e.*, as a deterministic variable instead of a random variable.

As a quantity that can be easily estimated from the movie, we consider the fraction of pixels occupied by the cell of interest for a given time-point with non-negative residuals. To do so, we start by finding the distribution of the residuals under an estimated solution $E[\boldsymbol{\beta}^{(q-1)}]$ for a mismatched $\kappa^{(q-1)}$ as follows:

$$\hat{r} = y - \boldsymbol{x}^T\boldsymbol{\beta}^{(q-1)} = \sigma - \boldsymbol{x}^T(\boldsymbol{\beta}^{(q-1)} - \boldsymbol{\beta}^*) \approx \sigma - \boldsymbol{x}^T[E[\boldsymbol{\beta}^{(q-1)}] - \boldsymbol{\beta}^*] = \sigma - b.$$

Here, with the approximation $E[\boldsymbol{\beta}^{(q-1)}] \approx \boldsymbol{\beta}^{(q-1)}$, the residuals follow the same probability distribution as $\sigma$, with a non-zero mean of $b$. Then, we theoretically approximate the mean number of pixels occupied by the cell that have non-negative residuals as:

$$\begin{aligned} p &= P(\hat{r} > 0) = P(\sigma > b) = \int_b^\infty [(1-\epsilon)\varphi(\sigma) + \epsilon h_\kappa(\sigma)]d\sigma, \\ &= 1 - \int_{-\infty}^b [(1-\epsilon)\varphi(\sigma) + \epsilon h_\kappa(\sigma)]d\sigma = 1 - (1-\epsilon)\int_{-\infty}^b \varphi(\sigma)d\sigma + \epsilon \int_{-\infty}^b h_\kappa(\sigma)d\sigma \\ &\approx 1 - (1-\epsilon)\Phi(b) = 1 - \frac{\kappa}{f(\kappa)}\Phi(b) \end{aligned}$$

From the second to third row, since $b$ is small for $\kappa^{(q-1)} \approx \kappa$, we have ignored the contribution from $h_\kappa(\sigma)$, which is only nonzero for $\sigma \geq \kappa \geq 0$. However, especially in cases where there is a large mismatch in $\kappa^{(q-1)} \neq \kappa$ such that $b \gg \kappa$, this term has a finite contribution at order $O(b)$.

Fortunately, we are only interested in the first-order expansion of this quantity for analytical tractability. Thus, we account for the first-order contribution coming from the $h_\kappa(\sigma)$ term via a user-defined scaling parameter ($\alpha \leq 1$, due to the opposite signs of the linear terms in the approximation) in the Taylor approximation as:

$$p \approx 1 - \frac{\kappa^{(q)}}{f(\kappa^{(q)})}\Phi(0) - \alpha\frac{\kappa^{(q)}}{f(\kappa^{(q)})}\varphi(0)b + O(b^2). \tag{10}$$

Here, the update, $\kappa^{(q)}$, is chosen to satisfy Eq. (10). We will show below that the final solution has the right distributional properties regardless of $\alpha$, whereas $\alpha$ plays the role of a learning rate.

#### *3.3.3. Linking estimated robustness parameter to the bias*

So far we linked the bias, $b$, to the ground truth $\kappa$, yet our overall goal is to create a connection between $\kappa$ and $\kappa^{(q-1)}$, using which we can perform a more accurate estimate, $\kappa^{(q)}$. A second link naturally emerges from the observation that even though $\kappa^{(q-1)} \neq \kappa$, we still minimize a 1-sided Huber loss under $\kappa^{(q-1)}$. In fact, since $\boldsymbol{\beta}^{(q-1)}$ is obtained by minimizing the loss function $E[\rho_{\kappa^{(q-1)}}(y - \boldsymbol{x}^T\boldsymbol{\beta}^{(q-1)})]$, we can write the optimality condition as:

$$
\begin{aligned}
0 &= E[\psi_{\kappa^{(q-1)}}(y - \boldsymbol{x}^T\boldsymbol{\beta}^{(q-1)})] = (1-\epsilon)E_{\Phi}[\psi_{\kappa^{(q-1)}}(y - \boldsymbol{x}^T\boldsymbol{\beta}^{(q-1)})] + \epsilon\, E_{H}[\psi_{\kappa^{(q-1)}}(y - \boldsymbol{x}^T\boldsymbol{\beta}^{(q-1)})] \\
&\approx (1-\epsilon)E_{\Phi}[\psi_{\kappa^{(q-1)}}(\sigma - b)] + \epsilon E_{H}[\psi_{\kappa^{(q-1)}}(\sigma - b)] \\
&= (1-\epsilon)\{\int_{-\infty}^{\kappa^{(q-1)}+b} (\sigma - b)\varphi(\sigma)d\sigma + \kappa^{(q-1)} \int_{\kappa^{(q-1)}+b}^{\infty} \varphi(\sigma)d\sigma\} + \epsilon\,\kappa^{(q-1)} + O([\kappa - \kappa^{(q-1)}]^2) \\
&\approx (1-\epsilon)\{-\varphi(\kappa^{(q-1)} + b) + \kappa^{(q-1)} - (\kappa^{(q-1)} + b)\Phi(\kappa^{(q-1)} + b)\} + \epsilon\kappa^{(q-1)} \\
&= (1-\epsilon)\{-f(\kappa^{(q-1)} + b) + \kappa^{(q-1)}\} + \epsilon\kappa^{(q-1)} = 0,
\end{aligned}
$$

$$\Leftrightarrow \frac{f(\kappa^{(q-1)}+b)}{\kappa^{(q-1)}} = \frac{1}{1-\epsilon}.$$

This condition is satisfied for any $\kappa^{(q-1)}$ and corresponding bias $b$. Thus, this is also satisfied when $\kappa^{(q-1)} := \kappa$ and $b = 0$ such that we recover the original equation $\frac{f(\kappa)}{\kappa} = \frac{1}{1-\epsilon}$. Using this, we can rewrite the condition resulting from the above equation as:

$$\frac{f(\kappa)}{\kappa} = \frac{f(\kappa^{(q-1)} + b)}{\kappa^{(q-1)}}.$$

As before, to estimate $\kappa^{(q)}$, we are only interested in the Taylor expansion for small $b$:

$$\frac{f(\kappa^{(q-1)}) + \Phi(\kappa^{(q-1)})b + O(b^2)}{\kappa^{(q-1)}} = \frac{f(\kappa)}{\kappa} \Rightarrow$$

$$b \approx [\kappa^{(q-1)} \frac{f(\kappa^{(q)})}{\kappa^{(q)}} - f(\kappa^{(q-1)})]/\Phi(\kappa^{(q-1)}) = \frac{f(\kappa^{(q-1)})}{\Phi(\kappa^{(q-1)})} \frac{\epsilon^{(q)} - \epsilon^{(q-1)}}{1-\epsilon^{(q)}} \quad (11)$$

such that the updated value of the contamination, $\kappa^{(q)}$, is chosen to satisfy the relationship in Eq. (11) and we define $\epsilon^{(q)}$ as the new estimated probability of the non-Gaussian noise.

*3.3.4. Eliminating estimation bias to update the contamination estimate*

Now, we can set $b$ in Eq. (11) into Eq. (10), which eliminates the bias and defines an adaptive update procedure:

$$p \approx 1 - (1 - \epsilon^{(q)})[\Phi(0) + \alpha\varphi(0) \frac{f(\kappa^{(q-1)})}{\Phi(\kappa^{(q-1)})} \frac{\epsilon^{(q)} - \epsilon^{(q-1)}}{1-\epsilon^{(q)}}]$$

$$\Rightarrow \epsilon^{(q)} = \epsilon^{(q-1)} - \frac{1-(1-\epsilon^{(q-1)})\Phi(0)-p}{\Phi(0)-\alpha\varphi(0)f(\kappa^{(q-1)})/\Phi(\kappa^{(q-1)})}. \quad (12)$$

Once we update $\epsilon^{(q)}$, the updated $\kappa^{(q)}$ is uniquely defined through Eq. (9) and can be estimated via $\kappa^{(q)} \approx h(\epsilon^{(q)})$. The main text includes the full picture (**Algorithm 2**) and additional details, *e.g.,* regarding the estimation of the probability that residuals are non-negative ($p$).

*3.3.5. The learning rate*

Finally, we discuss the role of $\alpha$ as a learning rate. Assuming convergence ($\epsilon^{(q)} = \epsilon^{(q-1)} = \epsilon$), Eq. (12) leads to:

$$1 - (1 - \epsilon)\Phi(0) - p = 0 \quad \Rightarrow p = \frac{1}{2}(1 - \epsilon) + \epsilon.$$

Once converged, as expected, the probability of observing positive residuals is $\frac{1}{2}$ the probability of Gaussian noise plus the probability of having non-Gaussian noise, without any dependence on $\alpha$. In practice, we set $\alpha = 0.05$ by default, which empirically led to more stable convergence.

*3.3.6. Generalization to the full movie*

We estimate $\epsilon^{(q)}$ for each cell and frame, which is then broadcasted to the full field-of-view via the linear model using cells' spatial footprints, $\boldsymbol{S}$. Finally, the $\boldsymbol{\kappa}$ matrix is obtained from $\boldsymbol{\epsilon}^{(q)}$ through entry-wise application of $\boldsymbol{\kappa} \approx h(\boldsymbol{\epsilon}^{(q)})$. The adaptive $\boldsymbol{\kappa}$ estimation can be performed during both cell refinement and final robust regression steps. Overall, we find that adaptive estimation is most necessary for the final robust regression, and cell refinement, whose main purpose is to extract robust filters, can be performed with a fixed point estimate of $\boldsymbol{\kappa}$. More details on this are provided in the main text, *e.g.*, **Figure 1H–M**.

#### 3.4. Adaptive estimation for cell baselines

Revisiting Eq. (7), we had defined a non-negativity condition for the estimated variables such that $\boldsymbol{\beta} \geq 0$. However, each cell may have a slightly different baseline activity that might not exactly be located at $\boldsymbol{\beta} = 0$. To incorporate this, we relax the non-negativity condition such that $\boldsymbol{\beta}_i \leq \boldsymbol{b}_i$, where $\boldsymbol{b}_i$ is the static baseline for the cell $i$. We use $\boldsymbol{b}_i = quantile(T_i, 0.25)$ as the default value, which can be adjusted by the user if cells are expected to be active $\geq 75\%$ of the time. Here, it is worth emphasizing that we are not subtracting the dynamically varying temporal background; rather estimating the static baseline $Ca^{2+}$ activity, which is performed only as part of the final robust regression before outputting the final $Ca^{2+}$ activity traces.

### §4 Algorithmic speed-ups with custom implementation

In this section, we discuss EXTRACT's algorithmic implementation, which speeds up the estimation process without sacrificing high-quality cell extraction.

### 4.1. Spatial partitioning in EXTRACT

Here, we introduce a novel two-level approach to spatial tiling of movies: The first level considers each partition as a movie of its own and runs the same cell extraction routine on each tile, potentially in parallel. The resulting patches are post-processed to remove the duplicate cells, similar to previous pipelines[7]. The second level is a subpartitioning routine unique to EXTRACT, which leads to faster and more accurate optimization as we discuss below.

#### *4.1.1. Spatial partitioning leading to independent patches*

In the first level of tiling, the movie is divided into pre-defined spatial partitions, each analyzed independently from one another. Later, cells from all partitions are combined via merging and thresholding based on quality metrics. EXTRACT uses this type of partitioning especially for movies with a large field-of-view (FOV), similarly to previous work[7]. Unlike CAIMAN, which typically explores small patches (up to 100 pixel in width) with 5-10 neurons[9], EXTRACT can handle $\sim O(1000)$ cells per spatial patch without significant compromises to speed or additional requirements for GPU or RAM memory, as enabled by the novel implementation detailed below in this section. The ability to process large spatial patches notably increases the quality of cell extraction, even for the $L_2$ solvers (**Figs. 3E,F, S4B**). With larger patches, one can demix crosstalk between (large) components that may not be well extracted from small patches, and a larger partition size leads to minimal handling of edge cases and resulting duplicates.

#### *4.1.2. Spatial partitioning within the fixed-point solver*

While the first level of partitioning is performed at the movie level, the second level is a subpartitioning routine within the ADMM solvers. Here, the robust regression is performed in batches, yet still giving EXTRACT access to the full field of view selected during the first level partitioning. This novel second-level partitioning is unique to EXTRACT's fixed-point solver, and mainly reflects that there is no background component that needs to be explicitly modeled. This property allows the extraction of disconnected and/or large spatial profiles, such as those belonging to dendrites (**Figure S8**), by running cell extraction on large spatial patches without additional burden on the GPU and/or RAM memory.

When estimating $\hat{S}$, the subpartitioning procedure is trivial and lossless. Since each pixel is estimated independently, one can divide the FOV of the movie into smaller chunks to run the regression. For EXTRACT, during the estimation of $\hat{S}$ we pick the number of subpartitions based on the available GPU/RAM memory. Empirically, the more pixels can be uploaded to the GPU, the faster the solver runs, as vectorized matrix operations are faster than running 'for loops'. Consequently, EXTRACT benefits from increased GPU memory, but it can also adapt to the existing GPUs' specifications without outside interference.

When estimating $\hat{T}$, the subpartitioning takes place in both spatial and temporal dimensions. The case of temporal partitioning is trivial and lossless; since the robust regression is run independently for each frame. The subpartitioning in the spatial domain is not lossless but is motivated by the locality and sparsity of cells' spatial footprints. Specifically, spatial subpartitioning would only affect a small portion of the cells and still preferable to the alternative, *i.e.*, using smaller spatial patches during the first-level partitioning. Yet, the question remains, how should we combine the activities of cells from distinct spatial subpartitions?

#### *4.1.3. Merging cells within the subpartitions:*

A straightforward approach[7] is to merge the cells and re-estimate the activity, or delete one of them and keep the other. Both options are suboptimal, as they either increase runtimes significantly or lead to suboptimal estimation of $Ca^{2+}$ activity traces due to missing pixels. For EXTRACT, we take a different, efficient, method that theoretically reproduces the original cells' activity in a certain limit. The core idea is to combine the already estimated $Ca^{2+}$ activity traces from the parts of the cell, divided into two or more subpartitions, using a linear combination:

$$\hat{T} = \boldsymbol{A}_1\hat{\boldsymbol{T}}_1 + \boldsymbol{A}_2\hat{\boldsymbol{T}}_2 + ... = \sum_{i=1}^{n_{partitions}} \boldsymbol{A}_i\hat{\boldsymbol{T}}_i. \quad (13)$$

Here, $\hat{T}$ is the desired estimate of the target cell's $Ca^{2+}$ activity trace, whereas $\hat{\boldsymbol{T}}_i$ is the $Ca^{2+}$ activity trace estimate computed from the subpartition $i$. To calibrate the coefficients, $\boldsymbol{A}_i$, we consider the combination that leads to minimum estimation errors in the limit case of no cell overlaps and $\kappa \to \infty$, *i.e.*, only Gaussian noise. Otherwise, the calculations with (adaptive) robust regression are not analytically tractable.

#### *4.1.4. Calibrating the coefficients for the merging:*

For simplicity, and without loss of generality for the case with no overlap, we consider a single frame and a single cell:

$$\boldsymbol{M} = \boldsymbol{S}T + \boldsymbol{\sigma}$$

where $\boldsymbol{\sigma}$ is i.i.d. Gaussian noise with zero mean and variance $\sigma^2_{iid}$, $\boldsymbol{S}^T\boldsymbol{S} = 1$, and $T$ is now a scalar corresponding to the activity of the target cell. Once we subpartition this movie into $n_{partitions}$ spatial parts, $\boldsymbol{S} = \sum_{i=1}^{n_{partitions}} \boldsymbol{S}^{(i)}$, the statistical model becomes:

$$\boldsymbol{M} = \left(\sum_{i=1}^{n_{partitions}} \boldsymbol{S}^{(i)}\right)T + \boldsymbol{\sigma} = \sum_{i=1}^{n_{partitions}} \boldsymbol{S}^{(i)}T + \boldsymbol{\sigma},$$

where $\boldsymbol{S}^{(i)T}\boldsymbol{S}^{(j)} = \boldsymbol{\tau}_i \leq 1$ if $i = j$ and zero otherwise. Here, $\boldsymbol{\tau}_i$ quantifies the fraction of the cell's area in the partition $i$. Then, the least-squares solutions, $\hat{\boldsymbol{T}}_i$, estimated from the partitions follow the distribution:

$$\hat{\boldsymbol{T}}_i = (\boldsymbol{S}^{(i)T}\boldsymbol{S}^{(i)})^{-1}\boldsymbol{S}^{(i)T}\boldsymbol{M} = T + \boldsymbol{\tau}_i^{-1}\boldsymbol{S}^{(i)T}\boldsymbol{\sigma} = T + \tilde{\boldsymbol{\sigma}}_i,$$

where $\tilde{\boldsymbol{\sigma}}$ is a transformed noise term with zero mean. As discussed in **Supplementary Note 1**, the least-squares solution has zero bias such that $E[\hat{\boldsymbol{T}}_i] = T$. This provides the first constraint for computing the coefficients, $\boldsymbol{A}_i$, in Eq. (13):

$$E[\hat{T}] = \sum_{i=1}^{n_{partitions}} \boldsymbol{A}_i E[\hat{\boldsymbol{T}}_i] = T \quad \Rightarrow \quad \sum_{i=1}^{N} \boldsymbol{A}_i = 1.$$

To obtain a second constraint, we compute the mean-squared error for the reconstructed $\hat{T}$:

$$E[(\hat{T} - T)^2] = E[\sum_j \sum_i \boldsymbol{A}_i \boldsymbol{\tau}_i^{-1} \boldsymbol{S}^{(i)T} \boldsymbol{\sigma}\ \boldsymbol{A}_j \boldsymbol{\tau}_j^{-1} \boldsymbol{S}^{(j)T} \boldsymbol{\sigma})] = \sum_j \sum_i \boldsymbol{A}_i \boldsymbol{\tau}_i^{-1} \boldsymbol{A}_j \boldsymbol{\tau}_j^{-1} E[Tr[\boldsymbol{\sigma}^T \boldsymbol{S}^{(i)} \boldsymbol{S}^{(j)T} \boldsymbol{\sigma}])]$$

$$= \sum_i \boldsymbol{A}_i^2 \boldsymbol{\tau}_i^{-2} \sigma_{iid}^2 \boldsymbol{\tau}_i = \sigma_{iid}^2 \sum_{i=1}^{n_{partitions}} \boldsymbol{A}_i^2 \boldsymbol{\tau}_i^{-1}$$

To calibrate $\boldsymbol{A}_i$, we wish to minimize the error associated with the reconstructed signal $\widehat{T}$ subject to the zero-bias constraint. To enforce this constraint, we can simply replace $\boldsymbol{A}_N = 1 - \sum_{i=1}^{n_{partitions}-1} \boldsymbol{A}_i$. Then, taking the derivative and setting it to zero, we obtain the second condition:

$$\boldsymbol{A}_i = \boldsymbol{\tau}_i\ , \tag{14}$$

where we used the identity $\sum_{i=1}^{n_{partitions}} \boldsymbol{\tau}_i = 1$, which can be derived from the normalization $\boldsymbol{S}^T \boldsymbol{S} = \sum_{i=1}^{N} \sum_{j=1}^{N} \boldsymbol{S}^{(i)T} \boldsymbol{S}^{(j)} = \sum_{i=1}^{N} \boldsymbol{\tau}_i = 1$. Then, the error becomes: $E[(\hat{T} - T)^2] = \sigma_{iid}^2$, the same error as if we simply had not partitioned (see Eq. (8) in **Supplementary Note 1**). Thus, using the scaling factors in Eq. (14), the least-squares estimator has the same accuracy after the subpartitioning as long as the cells are not overlapping.

Having considered one theoretical limit of the robust estimator for the subpartitioning, we can now gain intuition as to how subpartitioning works. In practice, we compute the cell areas in overlapping partitions, normalize with respect to the total area, and compute the weighted $Ca^{2+}$ activity traces. The resulting final $Ca^{2+}$ activity trace provides a faster, yet still accurate, alternative to existing methods. Specifically, subpartitioning within the ADMM solver allows performing cell extraction on the lowest number of spatial patches. An alternative is to use many more spatial partitions, with ~O(10) cells each instead of ~O(1000), running cell extraction independently across all of them. This is the approach taken by existing algorithms[9], as noted above.

To validate the subpartitioning approach empirically, in all simulation benchmarks we ran EXTRACT with the subpartitioning approach, which in all cases outperformed existing cell extraction methods and least-squares estimates (**Table S1**). Our extensive benchmarking provides strong empirical evidence that the spatial subpartitioning increases the speed of the algorithm without trading off accuracy.

#### 4.2. Pixel selection

Since robust regression does not require modeling of background sources, EXTRACT can cull pixels from the robust regression that do not contain any signal for cells' $Ca^{2+}$ activity traces. This helps decrease the number of pixels used during the regression and thus also decreases runtimes and memory requirements. We refer to this implementation trick as "pixel selection."

Let us consider an example to illustrate the massive impact that pixel selection has on speed and scalability. A FOV with 1024 × 1024 has $\sim 10^6$ pixels, whereas around 1000 cells tend to occupy $\sim 10^4 - 10^5$ pixels. Instead of using the pixels across the full field-of-view,

EXTRACT can decrease the number of pixels over which to perform matrix multiplications by at least an order-of-magnitude. Moreover, in addition to boosting speed, this massive reduction in problem size comes with unique benefits, such as reductions in RAM/GPU memory requirements, faster GPI upload times, and the ability to scale up to $1000$ cells within a single spatial partition.

In general, the ability to discard a sizable portion of the movie pixels during the optimization subroutines, without trading off accuracy, rests solely on the lack of background modeling and thus that the optimization problem is the same, regardless of whether these null pixels are included or not. As shown in **Figure 5K**, EXTRACT scales linearly with the number of cells and can handle movies of ~10 TB size (**Figure 4**). For large datasets, one bottleneck seems to be movie upload times (**Figure 4E**), which takes up ~$6\%$ of the movie runtime for a movie with 10,000 frames, but ~$25\%$ of the runtime for a movie with 100,000 frames. Therefore, with emerging large-scale $Ca^{2+}$ imaging technologies that can record $10000$ or even more cells simultaneously (**Figure 4**), robust regression approaches with no background modeling are expected to scale well into the future datasets.

#### 4.3. Downsampling in space and/or time

In addition to spatial partitioning, EXTRACT benefits greatly from downsampling in space and time, regarding both speed and accuracy (**Figure S6C**). When using EXTRACT, we have different recommendations for spatial and temporal downsampling.

Spatial downsampling can increase the movie SNR and speed the cell extraction process as the total number of pixels decreases. Yet, it also decreases the number of pixels per cell, thus potentially increasing the variance of the $Ca^{2+}$ activity trace estimation problem (**Supplementary Note 1**). Depending on the brain area and imaging modality, spatial downsampling may or may not be recommended. Empirically, we found that EXTRACT runs fastest for a mean cell radius of ~ 5 pixels without sacrificing the accuracy of cell extraction.

Unlike spatial downsampling, temporal downsampling is more forgiving, especially when performed during cell finding and refinement, though excessive downsampling can lead to suboptimal results (**Figure S6C**). In EXTRACT, we recommend performing temporal downsampling to speed cell finding and cell refinement, which also increases the accuracy of cell extraction. EXTRACT uses the resulting $\boldsymbol{S}$ to perform a final robust regression on the full movie, which leads to the final $Ca^{2+}$ activity traces without compromising time resolution. Empirically, we found that downsampling in time to 2–3 Hz is desirable and enhances the speed, but more aggressive downsampling reduces the cell extraction accuracy (**Figure S6C**).

**4.4. RAM/GPU memory usage.**

An important hallmark for an algorithm aiming to scale beyond the traditional boundaries of $Ca^{2+}$ imaging is the ability to utilize RAM and GPU memory without compromising the accuracy of cell extraction or employing techniques that slow its solvers. The worst-case scenario would be if an algorithm could not process movies beyond a certain size, for instance due to insufficient RAM memory (**Figure 4D**). Fortunately, when designing EXTRACT, we tested and confirmed its usability in large-scale movies, with no indication of saturation up to a movie 10 TB in size.

The GPU memory utilization is automatically performed within the ADMM solvers and through the subpartitioning technique discussed above. The movie is divided into spatial and temporal parts according to the available GPU memory (multiplied by a slacking factor) and processed in series over the GPU. This prevents memory errors and allows additional speedups with increased GPU memory.

The RAM memory requirements depend strongly on the movie and how much time downsampling is utilized during the cell finding and refinement procedures. Yet, the spatial partitioning allows EXTRACT to be able to process movies of any practical size on personal computers, including laptops. In fact, one of the simulation experiments (**Appendix Figure 1**) is performed on a personal Apple Macbook Air with the M1 chip and 16GB RAM. Thus, practically, there is no hard requirement of high RAM memory for processing large movies with EXTRACT, though we usually recommend leaving ~4 times the size of a single spatial partition available in the RAM memory.

An important final topic, which is usually overlooked in the publications of earlier approaches[7], is the claim that parallelization across >20 CPU cores is scalable, which is not true due to RAM memory requirements. Unfortunately, when cell extraction algorithms are used without GPUs, parallelization across multiple CPU cores comes with additional constraints on RAM memory usage. Thus, whereas the CPU version of EXTRACT performs quickly and accurately with movies of lower sizes, once one aims to process large movies >1TB size, the CPU implementation would simply not be scalable. This is another motivation for the field to move towards GPU implementable algorithms, which is the main design point that motivated our development of the ADMM approach.

### §5 Conclusion: Summary of EXTRACT's design choices

In this note, we first derived the algorithm behind the fixed point solver, *i.e.*, the ADMM approach for solving the constrained robust estimation problem. Next, we discussed the advantages conveyed by the superior implementation of EXTRACT, which were mainly possible thanks to the robust regression not requiring explicit modeling of background sources. Finally, we discussed EXTRACT's RAM/GPU memory requirements and its inherent scalability to large datasets. Here, we summarize how each step contributes to the speed and scaling of EXTRACT:

#### 5.1. Fixed-point solver

One of the biggest advantages of our solver comes from the observation that the solution of the robust regression tends to be topologically close to the least-squares estimate, *i.e.*, its Hessian can be approximated via $\boldsymbol{X}^T\boldsymbol{X}$. Thus, using $\boldsymbol{X}^T\boldsymbol{X}$ as the approximate Hessian, we designed the fixed-point solver with fast convergence properties, similar to Newton's method, *and* the matrix inversion can be precomputed at the beginning.

#### 5.2. ADMM approach for enforcing constraints

There are three straightforward possibilities we considered to enforce the constraints for the robust regression: i) through interior-point methods, ii) through projected gradient descent, and iii) through the ADMM framework. The first method would require the computation of a new inverse-Hessian at any step, and is ruled out due to its slow speed. Using the ADMM framework has two main advantages over the gradient-based methods: 1) ADMM framework, but not projected descent, can directly use the fixed-point solver and therefore converges faster. 2) As we discussed above, ADMM has two copies for the variable of interest, whose agreement can be monitored for early stopping.

#### 5.3. Algorithmic speed-ups

As discussed above, the fixed-point solver is ideally set up for vectorization on (potentially multiple) GPUs, as the solver mainly performs matrix multiplication. Thus, the subpartitioning process is primarily determined by GPU and RAM memory requirements, and not speed. Moreover, since EXTRACT performs no explicit background modeling, one can reduce the number of relevant pixels, and perform downsampling in time and space for cell finding and refinement procedures. These speed-ups would not be possible without the fixed-point solver or the use of the robust regression in EXTRACT as we discussed earlier.

#### 5.4. Scalability through multiple GPUs and memory requirements

EXTRACT achieves comparable speeds when parallelized on few GPUs vs across hundreds of CPU cores, which is a necessary requirement for processing large-scale $Ca^{2+}$ imaging movies without encountering speed or RAM memory issues. Unlike previous methods that either fail to utilize GPUs[7] or can use a single GPU[10], EXTRACT achieves scaling owing to its multiple GPU support. Given that $Ca^{2+}$ imaging datasets will only get bigger, multi-GPU support will become more essential due to the scalability and memory failures of multi-threading across CPU cores. EXTRACT, with its custom written fixed-point solver, is readily suitable to scale to the datasets of the future in systems neuroscience research.

**Supplementary Note 3: Supplementary benchmarking experiments**

### §1. Overview

Throughout the main text, we performed and discussed several simulation experiments (**Table S1**) to benchmark the speed and accuracy of EXTRACT against the previous state-of-the-art cell extraction algorithms[1–3]. To achieve a comprehensive evaluation of EXTRACT, we performed several supplementary experiments, which is what we discuss in this note.

In **§2**, we validate the cell refinement module (introduced in **Figure 2**) under ideal conditions for the movie generation, *e.g.*, with only Gaussian iid noise. Next, we show that our findings presented in the main text are not specific to the size of the simulated cells (**§3**) and that motion artifacts can have deleterious effects on the cell extraction process, which may not be immediately apparent from the aggregate trace quality metrics (**§4**). Then, we show that EXTRACT can perform cell extraction under extreme conditions such as very short movie durations (**§5**) and donut shaped cell profiles (**§6**). Next, we find that EXTRACT, but not necessarily SEUDO[2], is robust to non-idealities in the estimated cell profiles and generalizes across conditions, for which it was fine-tuned (**§7**). Finally, we conclude with a summary of the insights gained from additional experiments we performed in this note (**§8**).

### §2. The use of the robust loss function is beneficial even with iid Gaussian noise

In the main text, we mainly focused on how robust loss handles outliers to improve the cell extraction process. However, what if there were no inherent outliers in the movie? Would the use of the robust loss function still be necessary to achieve high quality cell extraction if the movie contained no neuropil, no dendritic activity, and no blood vessels? In other words, what if the movie noise was completely iid Gaussian?

To address this question, we first recall our discussion from **Supplementary Note 1**, in which the $Ca^{2+}$ imaging movies were modeled with the following equation:

$$\boldsymbol{M} = \boldsymbol{S}\boldsymbol{T} + \boldsymbol{\sigma} \tag{1}$$

Here, $\boldsymbol{M}$ refers to the movie, $\boldsymbol{S}$ and $\boldsymbol{T}$ correspond to cells' ground truth spatial profiles and $Ca^{2+}$ activity traces, respectively, and $\boldsymbol{\sigma}$ corresponds to the movie noise added to each of the pixel activities. Traditionally, $\boldsymbol{\sigma}$ contains several non-Gaussian and spatiotemporally correlated contamination such as out of focus background cells, neuropil, and/or blood vessel activities. For our purposes in this section, we assume that $\boldsymbol{\sigma}$ is sampled from an iid Gaussian distribution, *i.e.*, contains only random and independent noise.

Even in this overly idealized scenario, unless the cells' spatial profiles were estimated with perfect accuracy during the cell finding module (which is neither true nor realistic), there will always be a non-Gaussian contamination in the subsequent estimation problems. Specifically, assume that $\hat{\boldsymbol{S}}$ refers to the estimated profiles such that $\hat{\boldsymbol{S}} = \boldsymbol{S} - \Delta\boldsymbol{S}$. Then, Eq. (1) follows into the following form:

$$\boldsymbol{M} = \hat{\boldsymbol{S}}\boldsymbol{T} + \Delta\boldsymbol{S}\,\boldsymbol{T} + \boldsymbol{\sigma},$$

where the term $\Delta\boldsymbol{S}\,\boldsymbol{T}$ introduces non-Gaussian contamination for the estimation of $Ca^{2+}$ activity traces, $\hat{\boldsymbol{T}}$. In words, the activity of a neighboring cell, stored in $\boldsymbol{T}$, would introduce *structured noise,* in the form of $\Delta\boldsymbol{S}\,\boldsymbol{T}$, to the pixels occupied by the target cell, whose activity we seek to reconstruct. Thus, even if no explicit background contamination exists in the $Ca^{2+}$ imaging movies, we hypothesize that robust loss function is beneficial for the cell extraction problem.

To test this hypothesis, we simulated 2p $Ca^{2+}$ imaging movies that contained varying numbers of cells and only iid Gaussian noise (**Appendix Figure 5**). We processed these movies with EXTRACT and the $L_2$ solver, the version of EXTRACT that replaces the robust loss function with the traditional $L_2$ loss function at each module. While the conventional intuition may predict that an L2 solver would be optimal, we observed that EXTRACT led to both faster and more accurate convergence during cell refinement iterations. This was true for both the cell finding and the $Ca^{2+}$ activity trace quality metrics (**Appendix Figure 5**).

It is worth noting that even though the precision and recall values for EXTRACT may sometimes seem low immediately after the cell finding, *i.e.*, for zero cell refinement iterations, this is an artifact of how we chose to pair the extracted cells to the ground truth. The cell finding in EXTRACT used stricter (adaptive) $\boldsymbol{\kappa}$, as neighboring cells that were not yet found would introduce non-Gaussian noise. Thus, the first guess had cells whose spatial profiles contained only the highest SNR pixels (in line with **Fig. 1M**). However, even after just one cell refinement iteration, the cell profiles updated to include as many pixels as beneficial to the cell extraction, matching the ground truth and outperforming the $L_2$ solver.

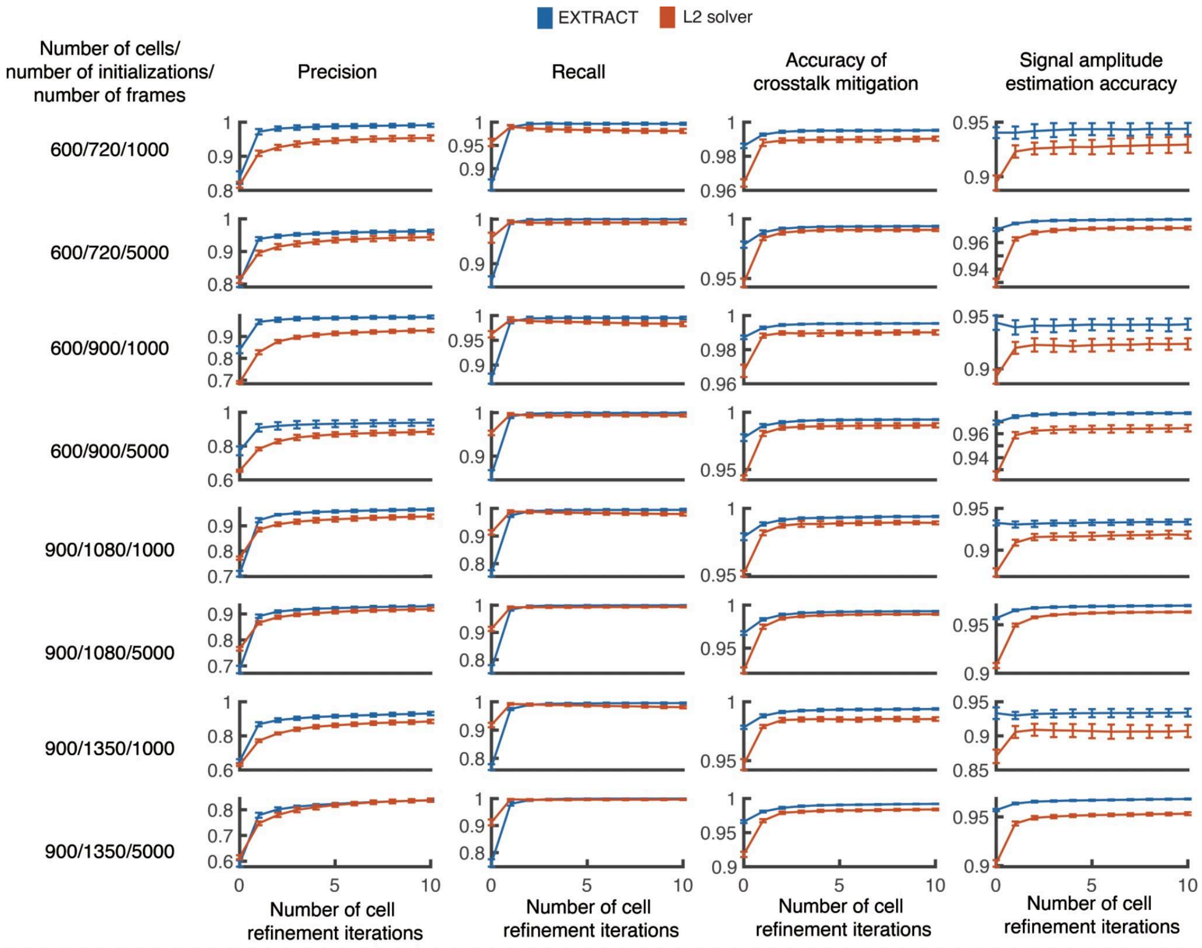

**Appendix Figure 5. Robust loss function leads to faster convergence with iid Gaussian noise.**

To test whether the use of the robust loss function is beneficial for processing $Ca^{2+}$ imaging movies with only independent Gaussian movie noise, we simulated two-photon $Ca^{2+}$ imaging movies (similar to **Figure S4B** but with no correlated noise component). Using both EXTRACT and the L2 solver, we processed movies with varying cell densities, duration, and maximum number of allowed cell candidates during the cell finding. In all cases, the use of the robust loss function led to superior cell extraction results (higher quality $Ca^{2+}$ activity traces, less duplicate and/or spurious cells) and faster convergence during the cell refinement.

### §3. EXTRACT can handle variations in imaging resolution

In the majority of the experiments, we simulated the cells using two-dimensional Gaussian functions, whose standard deviations in both x and y directions were sampled from a uniform distribution, *i.e.*, within $[3.5, 4.5]$ pixels. Since the cell extraction process makes no detailed assumption regarding the cell radius, or the spatial resolution in general, this choice should in theory have little-to-no effect on the quality of the cell extraction.

To test this implicit assumption explicitly, we simulated a set of 2p movies, which contained cells with varying levels of average radii, *i.e.*, imaging resolution (**Appendix Figure 6**). We processed these movies using EXTRACT, $L_2$ solver, CNMF[4] (MATLAB implementation of CAIMAN[1]), and ICA[3]. All algorithms, except for ICA, performed approximately the same as the imaging resolution varied. Although $L_2$ solver and CNMF had slight variations across the scales and would have likely benefited from fine-tuning of the hyperparameters, EXTRACT provided high quality cell extraction results across the scale (**Appendix Figure 6**). In contrast, ICA failed to process $Ca^{2+}$ imaging movies that contained small cells (occupying roughly 10-20 pixels), which may become a limiting factor for spatial resolution of the large scale $Ca^{2+}$ imaging movies that can be processed with ICA.

Overall, changing the cell radii measured in pixels, hence the spatial resolution, had little-to-no effect on EXTRACT’s cell extraction quality.

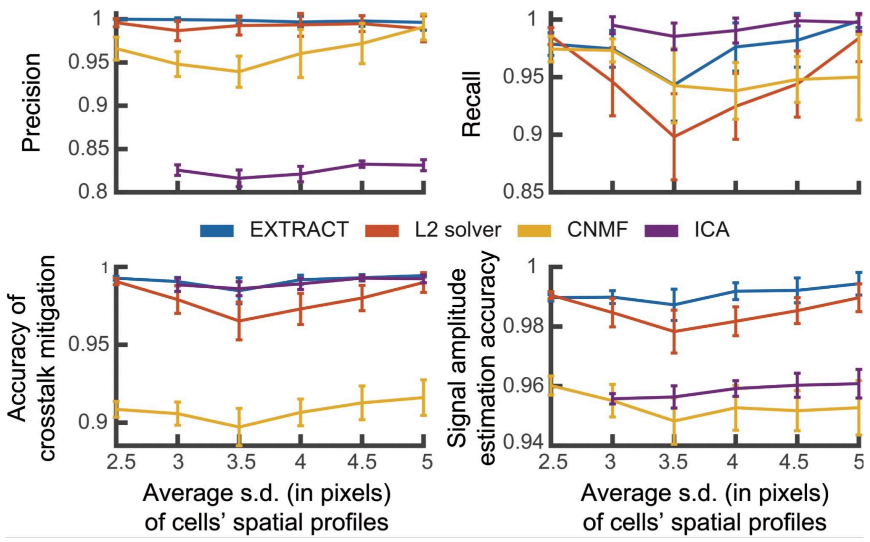

**Appendix Figure 6. EXTRACT can process movies with diverse imaging resolutions.**

To test the ability of cell extraction algorithms to perform in different imaging resolutions, we simulated various 2p movies with distinct cell radius and number that mimic zooming in on a particular region. While all algorithms performed similarly across a diverge range of imaging resolutions, ICA failed to provide sensible results for the movies that are most zoomed out.

### §4. EXTRACT requires correction of the motion artifacts

The one-sided robust loss function rejects positive outliers, *i.e.*, pixel activities that cannot be explained by the existing cells' $Ca^{2+}$ activity traces. As we have shown throughout the main text, as long as the cells are stationary, this approach improves the cell extraction results by removing crosstalk and other contaminants. However, the $Ca^{2+}$ imaging movies may contain significant brain motion, either because it may not be correctable using the existing motion correction algorithms[5,6] or the experiments may erroneously leave out motion correction as part of the preprocessing pipeline. In such cases, would the model agnostic nature of EXTRACT, specifically the lack of exponential kernels fitted to $Ca^{2+}$ activity traces[1,6], still be desirable?

To test this idea, we simulated several two-photon $Ca^{2+}$ imaging movies with varying levels of rigid motion (see **Supplementary Note 4** for details), similar to the experiments in

**Figure S6D**. We processed these movies with EXTRACT and CNMF, extracting cells’ spatial profiles and $Ca^{2+}$ activity traces. Then, we identified the $Ca^{2+}$ events by performing an exponential deconvolution using the ground truth $Ca^{2+}$ decay rate, and subsequently thresholding any event below 5% of the maximum $Ca^{2+}$ activity amplitude.

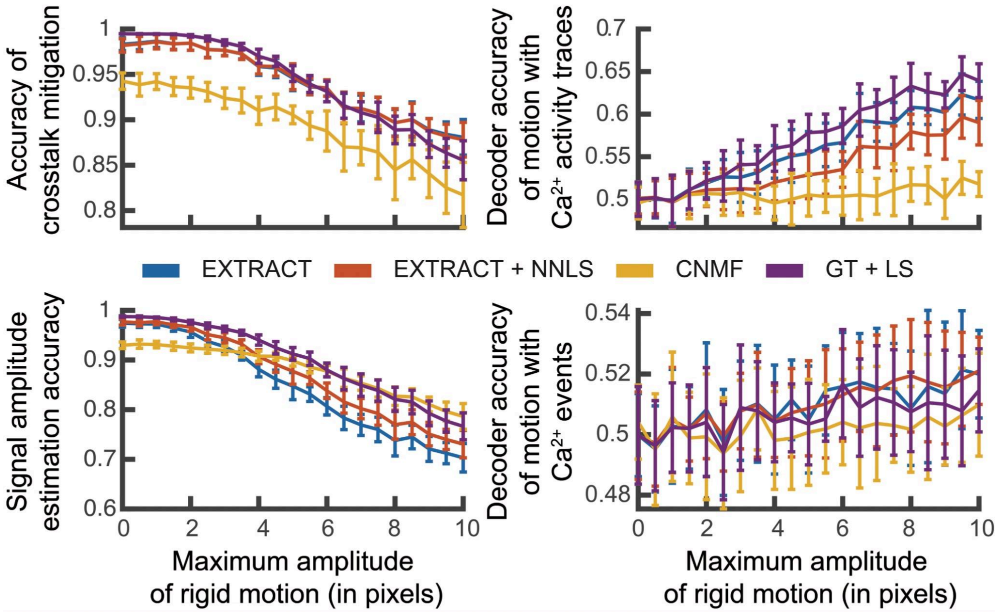

**Appendix Figure 7. EXTRACT requires motion correction, yet can handle small residuals.**

To test whether robust motion deteriorates the quality of the cell extraction outputs, we simulated two-photon $Ca^{2+}$ imaging movies with varying degrees of artificially induced (randomized) rigid motion. *Left.* All cell extraction algorithms were able to identify cells and their $Ca^{2+}$ activity traces despite residual motion, yet increasing motion artifacts deteriorated the cell extraction quality. *Right.* The motion can be decoded from the $Ca^{2+}$ activity traces, but not the $Ca^{2+}$ events, obtained via EXTRACT, whereas the exponentially decaying CNMF outputs did not encode the motion, but had low accuracy to begin with. All algorithms were virtually unaffected by rigid motions below a single pixel.

As a first test, we computed the $Ca^{2+}$ activity trace quality metrics while increasing the severity of the artificially induced brain motion (**Appendix Figure 7**, *left*). In these metrics, EXTRACT outperformed CNMF in its ability to mitigate the crosstalk, but had inaccurately estimated the amplitudes of the $Ca^{2+}$ activity events for rigid motion larger than 4 pixels. In other words, while the use of the robust regression suppressed crosstalk between neighboring cells and background contamination, part of the $Ca^{2+}$ signal amplitude was also deemed outlier due to severe motion. In fact, if the cells' spatial profiles are in motion, the neural signals could at times only coincide with a small fraction of the cells' spatial footprint and consequently be deemed outliers. On the other hand, existing pipelines, like CNMF, use exponential kernels to fit $Ca^{2+}$ activity traces, providing extra robustness to motion artifacts. Thus, the use of the robust loss function is advantageous when no brain motion is present, and suppresses the crosstalk in all cases. Yet, the assumption of a temporal kernel for neural signals, or a more agnostic version that enforces temporal consistency, may be utilized for future generations of EXTRACT as an optional parameter, which can be used to process $Ca^{2+}$ imaging movies whose motion cannot be corrected and better quality data collection is not possible.

Secondly, a hidden danger with processing movies with brain motion is the possibility that behavioral variables correlated with the brain motion may be decodable from neural activities even if there was no real coding in the neural activities. To illustrate this, we trained linear decoders, SVMs, to predict random motion times from the $Ca^{2+}$ activity traces and events (**Appendix Figure 7**, *right*). The raw $Ca^{2+}$ activity traces predicted the frames in the $Ca^{2+}$ imaging movie with motion, but the $Ca^{2+}$ events and the CNMF estimated traces did not, though the latter did not provide an accurate estimate of the $Ca^{2+}$ activity traces to begin with (**Appendix Figure 7**, *left*). This observation underscores the importance of performing event detection when analyzing $Ca^{2+}$ imaging datasets, especially those with residual motion artifacts.

To sum up, though fitting an exponential kernel to $Ca^{2+}$ activity traces may provide safeguards against the motion, a form of robust loss function should still be used in this process to allow proper estimation of $Ca^{2+}$ activity signals without crosstalk contamination. Therefore, until motion resistant robust cell extraction algorithms are developed, the process of motion correction is necessary for all $Ca^{2+}$ imaging movie preprocessing pipelines.

### §5. EXTRACT can process short $Ca^{2+}$ imaging movies

In several experimental conditions, it may be preferable, or even necessary, to record short $Ca^{2+}$ imaging movies. To provide validity evidence that EXTRACT can process very short movies, we simulated several two-photon $Ca^{2+}$ imaging movies with as little as 500 frames, corresponding to 50 seconds of imaging. As shown in **Appendix Figure 8**, EXTRACT outperformed CAIMAN in all metrics and was able to accurately perform cell extraction with simulated $Ca^{2+}$ imaging movies that have as little as one minute duration.

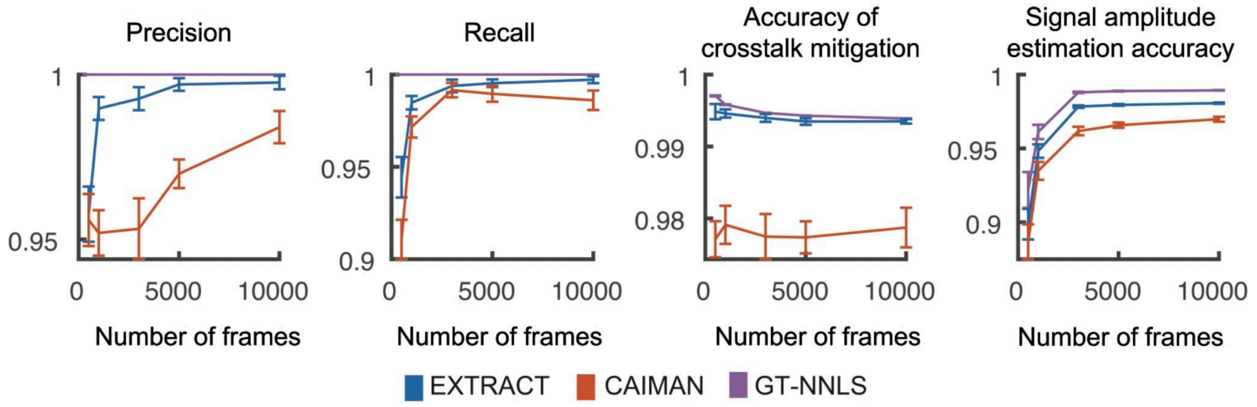

**Appendix Figure 8. Testing EXTRACT's ability to process short $Ca^{2+}$ imaging movies.**

To test whether EXTRACT can process short $Ca^{2+}$ imaging movies, we simulated 2p movies, similar to the ones in **Fig. 3A**. EXTRACT was able to process movies as short as 500 frames, roughly a minute in duration. As expected, both EXTRACT and CAIMAN benefited from the increased imaging duration.

### §6. EXTRACT can extract donut shaped cells

Though currently not in as much active use, previous work regularly benchmarked cell extraction algorithms on donut shaped neurons that arise from using $Ca^{2+}$ indicators excluding nucleus[4,7]. In line with the established literature, we performed additional experiments by simulating two-photon $Ca^{2+}$ imaging movies containing cells with donut shaped profiles (See **Methods**, following Ref.[4]). We processed these movies using EXTRACT, $L_2$ solver, CNMF[4], and ICA[3] (**Appendix Figure 9**).

EXTRACT makes no assumptions regarding cells’ spatial profiles. In line with this, EXTRACT’s cell extraction results on these movies were on par with those from the simulated $Ca^{2+}$ imaging movies with Gaussian shaped cells **(Appendix Figure 9**). However, both CNMF and ICA failed to accurately estimate cells’ spatial profiles and had multiple cell components that had only mild overlaps with the ground truth **(Appendix Figure 9**). Thus, EXTRACT’s agnostic nature was particularly beneficial for experimental conditions, for which the assumptions of previous cell extraction algorithms were not satisfied.

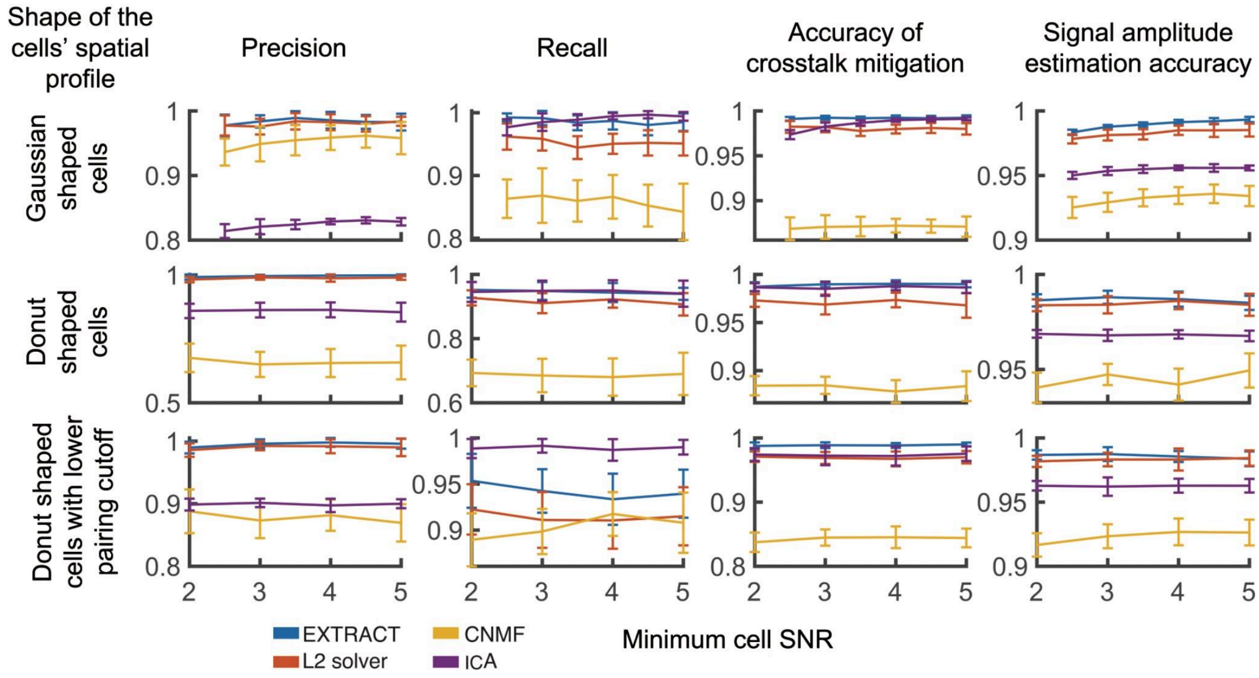

**Appendix Figure 9. Testing EXTRACT’s ability to identify donut shaped neurons.**

To illustrate that EXTRACT makes no specific assumption regarding cells’ spatial profiles, we simulated two-photon $Ca^{2+}$ imaging movies with circular (Gaussian) and donut shaped cells for varying levels of cell SNRs. While EXTRACT was able to identify both types of cells regardless of cell SNR with high correlations between ground truth and estimated spatial profiles, ICA and CNMF failed to identify donut shaped cells accurately. Even when lower pairing cutoffs were used, *i.e.*, cells’ with even mild overlaps were matched, CNMF failed to identify 10% of the donut shaped cells while returning 10% false positives.

### §7. Generalization and robustness to imperfectly estimated cell profiles

In the main text, we have observed that the quality of the final $Ca^{2+}$ activity traces is strongly dependent on the quality of the cells' estimated spatial profiles (**Fig. 3G,H**). This was true for all algorithms, *i.e.*, EXTRACT, NNLS/LS estimates of the $Ca^{2+}$ activity, and the recently developed $Ca^{2+}$ activity trace post-processing approach, SEUDO[2]. Moreover, we also observed that SEUDO required fine-tuning of its hyperparameters (**Fig. S5**), which may be an indication of generalization issues. Combined, these observations are in direct contrast to the premise of using post-processing tools to re-estimate $Ca^{2+}$ activity traces, a correction process that has been recently gaining interest[2,8]. In this section, we first quantify the generalization properties of the state-of-the-art $Ca^{2+}$ activity trace post-processing tool, SEUDO[2]. We, then, test its robustness to the variations in the quality of estimated spatial profiles, comparing and contrasting against the robust regression performed by EXTRACT.

As a first step, we tested whether hyperparameters of SEUDO can generalize out of the specific configurations for which they were optimized. We simulated 2p $Ca^{2+}$ imaging movies with 150 cells and initialized the trace estimator algorithms, both SEUDO and EXTRACT's final robust regression, with varying percentages of the ground truth cells' spatial profiles. A lower percentage of initialized cells corresponds to higher amounts of background contamination, as left-out cells are to model neuropil and introduce crosstalk. In this setting, we optimized SEUDO through a grid search for a highly contaminated movie, *i.e.* <50 cells initialized. We observed that some subsets of hyperparameters did not generalize to less contaminated movies, *i.e.* >50 cells initialized, whereas select few did (**Appendix Figure 10**, left). On the other hand, the robust regression, which we already showed to be robust to variations in hyperparameters (**Figure 1G**), generalized seamlessly across different levels of contamination, mainly thanks to the adaptive estimation of $\boldsymbol{\kappa}$ (See **Methods** and **Supplementary Note 2** for details of how the adaptive estimation is performed). In line with these results, the generalization ability of SEUDO, but not EXTRACT, was severely affected when cells' ground truth spatial profiles were replaced with some inaccurately estimated ones (**Appendix Figure 10**, middle).

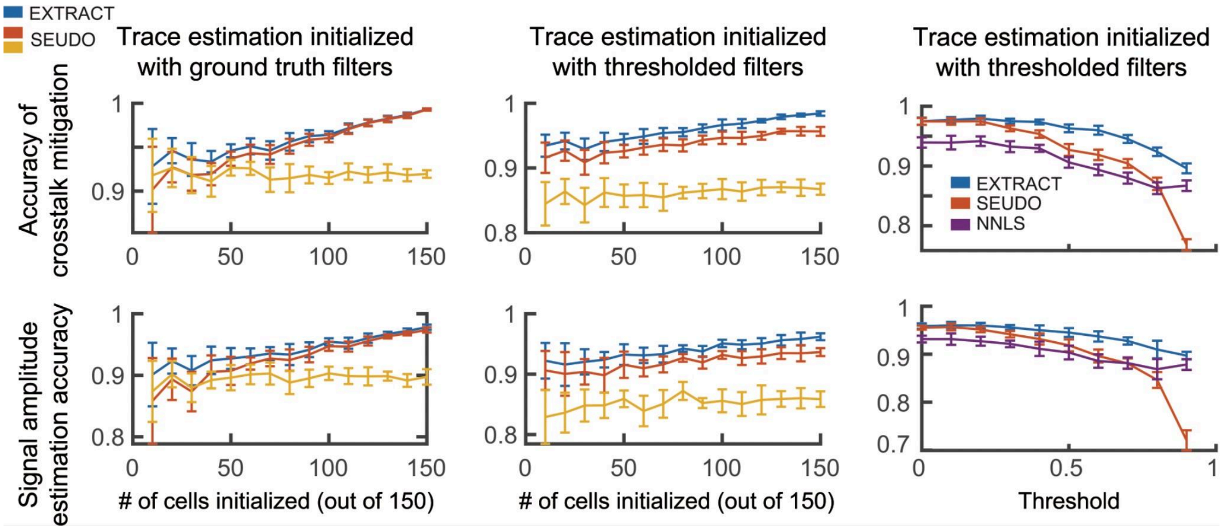

**Appendix Figure 10. Testing the generalization properties of EXTRACT and SEUDO.**

To test whether EXTRACT and/or SEUDO are robust to variations in hyperparameters that are optimized for a specific contamination level and to inaccuracies in the estimated spatial profiles, we simulated additional 2p $Ca^{2+}$ imaging movies, each containing 150 cells. *Left.* We initialized both algorithms with varying numbers of ground truth cells' spatial profiles and compared the resulting trace estimation accuracies. We picked two hyperparameter configurations from our grid search for SEUDO, which was optimal for highly contaminated movies (less than 50 cells initialized), and performed a sweep across contamination levels. SEUDO had both types of configurations, those that generalized and those that did not. The robust regression utilized by EXTRACT generalized seamlessly, thanks to the adaptive estimation of $\boldsymbol{\kappa}$. *Middle.* The same as the plot on the left, but the cells' ground truth spatial profiles were modified to introduce inaccuracies. Specifically, we thresholded them below 0.4, setting the values of "estimated" cells' profiles to be zero whenever the ground truth profile had less than 0.4 of its maximum weight. *Right.* We initialized both algorithms with 80% of the ground truth cell profiles. We varied the threshold values, *e.g.*, the one used for the experiments in the middle, with higher values corresponding to more inaccurate spatial profiles. While SEUDO performed similarly to EXTRACT when ground truth profiles were used, as the estimated filters varied from the ground truth, EXTRACT but not SEUDO remained robust to imperfect spatial profiles, with the latter underperforming compared to NNLS.

Next, we quantified the robustness of both algorithms to inaccuracies in estimated cells' profiles in more detail. To do so, we focused on the case with 80% identified cells and varied the severity of mismatch between the ground truth spatial profiles and those used during the $Ca^{2+}$ activity trace estimation (**Appendix Figure 10**, right). While EXTRACT remained robust to the inaccuracies in the estimated profiles, SEUDO did not and was eventually outperformed by even the NNLS estimates.

Combining our observations in this section with **Fig. 3G,H**, we observe that estimation of cells' spatial profiles is a crucial process that cannot be skipped or delegated to inferior cell extraction algorithms, rather should be performed accurately via robust regression. Inaccuracies in the spatial profiles have direct influence on the estimated $Ca^{2+}$ activity trace quality. Therefore, the trace post-processing is a suboptimal process by design.

### §8. Conclusion

In this note, we performed additional experiments to test aspects of EXTRACT that were not primarily discussed in the main text, but nonetheless are crucial to validate the use of the robust loss function for the cell extraction problem. Specifically, we have shown that i) the use of the robust loss function is beneficial even when the noise in the pixel activities are random and independent, ii) EXTRACT can handle variations in the imaging and experimental conditions, but still requires correction of the motion, and iii) EXTRACT, but not necessarily prior algorithms, show true signs of robustness and generalizability across datasets and non-ideal scenarios.

### Supplementary Note 4: Additional details on the large scale simulation benchmark

### §1. Overview

$Ca^{2+}$ imaging is a mature technology that allows simultaneous recordings of tens of thousands of neurons in the live animal brains[1]. To process the resulting movies in a timely manner, several automated algorithms have been developed in the literature to perform cell extraction, *i.e.*, the identification of cells' spatial profiles/locations and their $Ca^{2+}$ activity traces[2–6]. Even though some standardized benchmarks, such as Neurofinder challenge[7], aim to test the efficacy of cell extraction algorithms in identifying cells' locations correctly, a comprehensive benchmark with controlled simulated conditions and ground truth, which can facilitate accurate tests of $Ca^{2+}$ activity trace quality, has been lacking in the literature. Prior work benchmarked the cell extraction algorithms either in simple simulated conditions with a few movies[6,8], or utilized real datasets[2], in which no ground truth exists, and tends to provide anecdotal evidence for the estimated $Ca^{2+}$ activity traces or relies on human annotation[9] that may be faulty (as was found out[2] with the Neurofinder Challenge). Moreover, most results reported on $Ca^{2+}$ activity trace quality usie ill-defined metrics, such as Pearson's correlation coefficients of $Ca^{2+}$ activity traces estimated with other methods[4,10]. However, the correlation is highly susceptible to the correct or incorrect estimation of the $Ca^{2+}$ indicator decay times, rather than the true timing and amplitude of $Ca^{2+}$ events, making the results uninterpretable. Instead, in this work, we rely on interpretable and validated quality metrics and large scale simulation benchmarks with controlled conditions.

To facilitate reproducibility of our results in this work, this note discusses the methodological details of our simulation benchmark and reports some of the extra experiments we performed. Specifically, **§2** introduces the mathematical framework we used to simulate $Ca^{2+}$ imaging movies and provides the detailed lists of parameters used for each experiment in the benchmark (**Table S1**). Then, we perform additional experiments benchmarking EXTRACT's abilities under diverse experimental and imaging conditions (**§3**). Finally, we conclude with final remarks by providing a summary of our empirical test on the simulation benchmark (**§4**).

### §2. Details of the simulation benchmark

In this supplementary note, we detail the conditions of each experiment in the simulation benchmark, which consists of 33 distinct conditions to test EXTRACT's cell extraction abilities (**Table S1**). Even though we provided the simulation details of the $Ca^{2+}$ imaging movies in the **Methods** section of the main text, we start with an overview of the movie generation algorithm for completeness.

#### 2.1. An overview of the movie generation steps

To date, the benchmarking of existing cell extraction algorithms is often limited to a few simulated (idealistic) $Ca^{2+}$ imaging movies[8,11], public benchmarks that primarily focus on cell finding quality on few human annotated $Ca^{2+}$ imaging movies[7], and/or a few realistic simulations of two-photon $Ca^{2+}$ imaging movies[10], which require time and resources to generate at scale. In this work, we took an intermediary approach, generating simulated $Ca^{2+}$ imaging movies at scale using idealistic assumptions. But, we also added additional realistic components such as variations in the neural spiking amplitudes and background contamination in the form of spatiotemporally correlated noise (**Methods**), which we summarize below.

To simulate a typical 2p $Ca^{2+}$ imaging movie, we used the the following model:

$$\boldsymbol{M}_{2p} = \boldsymbol{S}_{cell}\boldsymbol{T}_{cell} + (1 - \boldsymbol{\epsilon})\boldsymbol{\sigma}_{iid} + \boldsymbol{\epsilon}\boldsymbol{\sigma}_{neuropil} + \boldsymbol{F},$$

in which $\boldsymbol{S}$ and $\boldsymbol{T}$ contain the cells' spatial footprints and $Ca^{2+}$ activity traces, $0 \leq \boldsymbol{\epsilon} \leq 1$ defines the level of spatiotemporally correlated neuropil contamination, $\boldsymbol{\sigma}_{neuropil}$, that may not necessarily be sampled from an iid Gaussian distribution, $\boldsymbol{\sigma}_{iid}$, and $\boldsymbol{F}$ is the static fluorescence background, which is set to one, without loss of generality, unless otherwise specified.

A typical 1p $Ca^{2+}$ imaging movie has the same components as the 2p $Ca^{2+}$ imaging movies, plus a set of out-of-focus (OOF) cells with varying diameters, typically modeled with $\approx [50, 180]$ μm, to model the locally changing 1p background contamination:

$$\begin{aligned} \boldsymbol{M}_{1p} &= \boldsymbol{S}_{cell}\boldsymbol{T}_{cell} + \boldsymbol{S}_{OOF}\boldsymbol{T}_{OOF} + (1-\boldsymbol{\epsilon})\boldsymbol{\sigma}_{iid} + \boldsymbol{\epsilon\sigma}_{neuropil} + \boldsymbol{F} \\ &= \boldsymbol{M}_{2p} + \boldsymbol{S}_{OOF}\boldsymbol{T}_{OOF}. \end{aligned} \tag{1}$$

This form of movie generation is consistent with the movies used to benchmark CNMF-E[6], the former state-of-the-art cell extraction algorithm for 1p $Ca^{2+}$ imaging movies.

Though earlier work also considered addition of blood vessels to these simulations[6], we chose not to add them explicitly here. Instead, our goal with the simulation benchmark has been to create conditions, matching the assumptions of former cell extraction algorithms. Specifically, EXTRACT is agnostic to the source of the background contamination, whereas other cell extraction algorithms make explicit assumptions on the background. Thus, our setting gave CNMF/-E[4,6,8] clear advantages to model the data generation process of the movie explicitly.

### 2.2. Default parameters for the simulation benchmark

Before we discuss the details of each experiment, we first note the following default values of the simulation:

***General parameters:***

- Frame rate: 10 Hz
- Total number of frames, $n_{times}$: 5000 frames
- FOV size, $ns$: $250 \times 250\, pixel^2$ corresponding to $400 \times 400\, \mu m^2$
- Total number of cells, $n_{cell}$: 600 for the cells of interest, 150 for OOF cells.
- % of correlated noise component, $\boldsymbol{\epsilon}$: 5%

***Generation of the cells' spatial footprints:***

- Size of cell filters: the s.d. of the Gaussian distribution is uniformly sampled from [3.5,4.5] pixels for cells of interest, [15,50] for OOF cells.
- Minimum distance between generated cells: 4 pixels for cells of interest, 0 for OOF cells.

***Generation of the $Ca^{2+}$ activity traces:***

- Event rate, $r_{event}$: 0.1Hz for cells of interest ($\boldsymbol{M}_{2p}$), 1Hz for OOF cells
- Decay time for the $Ca^{2+}$ indicators: 1s for cells of interest ($\boldsymbol{M}_{2p}$), 4s for OOF cells
- Refractory period: 1 frame
- Noise level, $\sigma_{pixel}$: 0.02
- Minimum cell SNR, $SNR_{min}$: four times the s.d. of the noise.
- Minimum amplitude of an OOF cell: 0.04. Note that this corresponds to an SNR of ~2.
- Spike synchronization probability, $p_{synch}$: 0 if independent, 0.5 if correlated
- Number of cells in synchronization groups, $n_{synch}$: [50,100] cells
- Variation in the spiking amplitudes, $A_{spike}$: 0.3. As discussed in the **Methods** of the main text, the $Ca^{2+}$ events are sampled following $T_{amp} = (1 + N_S) \cdot \sigma_{pixel} \cdot SNR_{min}$, with $N_s = Poisson\,(A_{spike})$.

When simulating 1p $Ca^{2+}$ imaging movies, we create two distinct movies: The first movie ($\boldsymbol{M}_{2p}$) contains the cells' contributions to the total fluorescence and the simulated noise (with $\sigma_{pixel} = 0.02$). The second movie is a background with a negligible iid noise level (with $\sigma_{pixel} = 0.002$, hence has negligible effects to the iid noise), and contains the out of focus cells. For the latter, to properly develop neuropil signals, we let the OOF cells ramp up their firing patterns for 1000 frames before starting the sampling. To obtain the final 1p $Ca^{2+}$ imaging movie ($\boldsymbol{M}_{1p}$), we combine the two simulated parts following Eq. (1).

### §3 Benchmarks testing the need for the robust estimation framework

The first six experiments in the benchmark (**Table S1**) test how the addition of the robust loss affects the individual modules of EXTRACT. Specifically:

- Cell finding: Experiments 2 and 6
- Cell refinement: Experiments 3 and 4
- Final robust regression: Experiments 1 and 5

#### Ex 1: Need for the robust estimation of $Ca^{2+}$ activity traces

This experiment isolates the estimation procedure of the $Ca^{2+}$ activity traces from the rest of the cell extraction pipeline, which is achieved by initializing the cell extraction algorithms with cells' ground truth profiles.

We simulated 20 two-photon movies with the default parameters, except, the cells had independent events and the movies had zero fluorescence background $\boldsymbol{F} = 0$. We initialized the cell extraction algorithms with 20%, 40%, 60%, 80%, and 100% ground truth cell profiles to estimate cells' $Ca^{2+}$ activity traces. For each movie, we performed the trace estimation with and without[12] adaptive estimation of $\boldsymbol{\kappa}$ while the initialization of $\boldsymbol{\kappa}$ varied across [0.1,0.3,0.5,0.7,1,2, 3,5,10,100] s.d. of the median movie pixel activity. Since $\boldsymbol{F} = 0$, the movies were already assumed to be in the $\Delta F/F$ units. This allowed applying the non-negative solver directly without movie pre-processing and/or cell-baseline correction.

The cells that were not used during the initialization are meant to mimic cells not found by the cell extraction module and/or other forms of spatiotemporally correlated activity such as blood vessels, neuropil etc. As the percentage of initialized cells decreases, more spatiotemporally correlated background activity should be mitigated in the trace estimation, *e.g.*, non-Gaussian component of the noise becomes more prominent. Since the adaptive estimation of $\boldsymbol{\kappa}$ is a non-convex problem, not all initial conditions lead to the same $\boldsymbol{\kappa}$ values. Empirically, as shown in **Fig. 1G** and **Fig. S1E,** all tested initial values, especially those close to $\sim 1$ s.d. of the median movie pixel activity, corresponded to high quality estimation of $Ca^{2+}$ activity traces.

**Ex 2: Need for the robust loss function in cell extraction**

This experiment tests whether the use of the robust loss function for extracting cells has any advantages, or whether EXTRACT's superior cell extraction compared to the existing algorithms can be explained by specific hyperparameters used and/or EXTRACT's superior software implementation.

We simulated a total of 120 one-photon and two-photon $Ca^{2+}$ imaging movies with increasing numbers of cells (300,900,1500), otherwise using the default parameters for the independently firing neurons. In the end, this corresponded to 20 simulated $Ca^{2+}$ imaging movies per tested experimental condition. As discussed in the main text, we designed an $L_2$ solver that shares the same implementation as EXTRACT, simply by using a fixed, practically 100 s.d. but effectively infinite value for $\boldsymbol{\kappa}$, which converts the robust solver into a non-negative least-squares (NNLS) solver with adaptive cell baseline estimation. Otherwise, both algorithms used the same base modules of EXTRACT (**Figure 2**). We allowed both algorithms to identify more cells during the initial cell finding stage, roughly 1.5 times the number of cells in the $Ca^{2+}$ imaging movies. This increased number stems from the fact that the cell finding module is not perfect and tends to identify several duplicates in certain cases. Moreover, in real datasets, experimenters do not know the exact number of cells in the movie, in which case overshooting first and then culling false-positives may be preferable for maximizing the recall. The duplicated cell candidates were then either to be rejected during the cell finding module or discarded during the cell refinement module.

Since the $L_2$ solver has the exact same implementation as EXTRACT, except for the utilization of the robust loss, this setting allowed us to test whether the robust loss had significant contributions to the superior cell extraction results reported throughout the paper. The theoretical prediction was as follows: As the cells became highly overlapping, the severe non-Gaussian noise contaminants, *e.g.*, crosstalk of $Ca^{2+}$ signals from neighboring sources, would significantly impede the ability of the $L_2$ solver to identify cellular profiles. On the contrary, the robust loss function would be less impacted by increased cell density and outperform the $L_2$ solver. Our empirical results, presented in **Fig. 3E**, confirmed this theoretical intuition. The use of the robust loss function led to higher quality cell identification for all cases, and the difference between the outputs of the two algorithms became more pronounced with higher cell density.

**Ex 3: Validation of cell refinement procedures with a robust loss function**

As discussed in the **Supplementary Note 1**, even though earlier approaches to cell extraction used alternating estimation of cells' spatial profiles and $Ca^{2+}$ activity traces[4], the discarding of low quality components, *i.e.*, putative cells, is traditionally performed after the fact. With EXTRACT, we introduced the cell refinement module (**Figure 2**), which performs quality checks in a closed loop in between, not after, the alternating iterations between cells' spatial and temporal components. To delineate the novel contribution of the robust loss function to this process, this experiment performs a similar test to Experiment 2, but specifically focusing on EXTRACT's cell refinement module.

As before, we simulated 20 one-photon and two-photon movies using the default parameters for independent spiking conditions. The movies were processed with both the $L_2$ solver and EXTRACT, separately, with increasing numbers of cell refinement steps. Results, shown in **Fig. S4B**, indicated that cell refinement was beneficial for both of the algorithms, whereas the latter converged to higher quality results with far less iterations (roughly 6, which is set as the default value).

**Ex 4: Faster cell refinement with robust loss function under iid Gaussian noise**

The $Ca^{2+}$ imaging movies we simulated in Experiment 3 contained small, but nonvanishing, amounts of correlated noise that aims to model spatiotemporally correlated neuropil contamination. However, would the robust loss function still be beneficial if the movies contained only iid Gaussian noise?

As before, we simulated 80 two-photon $Ca^{2+}$ imaging movies for combinations of several parameters (corresponding to 10 movies for each combination): $n_t = (1000, 5000)$, $n_{cell} = (600, 900)$, and $f_{cell} = (1.2, 1.5)$, where $f_{cell}$ is the ratio of maximum number cells that can be found during the cell finding module to the number of ground truth cells in the movie. Otherwise, we used the default parameters for independently firing neurons.

The existence of overlapping cells introduces inherent non-Gaussian and spatiotemporally correlated noise components to the cell extraction problem (discussed in theoretical details in **Supplementary Note 1**). Specifically, the signal for one of the cells may become noise for an overlapping neighboring cell. As discussed below in **Appendix Figure 5**, the use of the robust loss function was beneficial for the cell refinement process, even though the simulated $Ca^{2+}$ imaging movies contained only iid Gaussian noise components.

**Ex 5: Need for the robust estimation of cells’ spatial profiles and activity baselines**

This experiment aims to study the effects of accurately estimating the cells’ spatial profiles and baselines to the quality of final $Ca^{2+}$ activity traces, and whether robust regression plays a significant role in trace estimation once other components are estimated accurately via robust regression.

To study this question, we performed two distinct tests. In the first test, we simulated a total of 120 one-photon $Ca^{2+}$ imaging movies with the minimum cell SNR values varying across $SNR_{min} = (2.5, 3, 3.5, 4, 4.5, 5)$, 20 movies per condition. Each movie had 500 independently spiking cells in a $240 \times 240\, \mu m^2$ FOV, otherwise we used the default parameters. We processed these movies with EXTRACT using an $L_1$ regularized ($\lambda_1 = 0.1$) robust regression for estimating $Ca^{2+}$ activity traces. In the second test, we focused on $Ca^{2+}$ imaging movies with a particular cell SNR ($SNR_{min} = 4$) and performed a swept across the $L_1$ regularization parameters $\lambda_1 = (0, 10^{-3}, 10^{-2}, 10^{-1}, 1)$.

In both tests, we used EXTRACT to obtain the cells' spatial profiles, $\boldsymbol{S}$. Then, using these robustly estimated profiles, we computed additional sets of $Ca^{2+}$ activity traces via NNLS, NNLS with cell baseline adjustment, and least-squares. The results, presented in **Fig. S4C and D**, led to several important conclusions: i) a direct NNLS estimate, without adaptively estimating cells' baseline activities, was severely suboptimal as movie baselines ($\Delta \boldsymbol{F} = 0$) do not necessarily coincide with the cells' baselines ($\boldsymbol{T} = 0$). ii) After baseline adjustments, NNLS outperformed least-squares in estimating cells' $Ca^{2+}$ activity traces, in line with our discussions in **Supplementary Note 1**. iii) Robust regression outperformed all other methods, suggesting that even when everything else is accurately estimated, robust regression is needed to accurately estimate cells' $Ca^{2+}$ activity traces.

**Ex 6: Changing the robustness parameter during cell finding**

The use of the robust loss function benefits the quality of cells' estimated spatial profiles, and subsequently the final $Ca^{2+}$ activity traces. Yet, how does the specific choice of the robustness parameter, $\boldsymbol{\kappa}$, affect the quality of cell finding, especially since it is not adaptively estimated during the profile estimation procedure (**Methods, Fig. 1M**)?

To answer this question, we simulated 20 one-photon movies with low snr ($SNR_{min}$ = 2.5), otherwise using the default parameters for independently firing neurons. We processed these movies using fixed and homogenous κ values ranging across (0.05,0.1,0.2,0.3,0.4,0.5, 0.7,1,10,100) and computed the F1 score for cell finding. For these movies, we performed 10 cell refinement iterations, but did not allow the adaptive estimation of κ.

The results, presented in **Fig. S4E**, showed a complex relationship between the used fixed κ values and the resulting cell finding quality. Overall, picking $\boldsymbol{\kappa} \sim 0.2$ for the one-photon $Ca^{2+}$ imaging movies resulted in the best cell finding results. Later applications to real datasets suggested that adaptive estimation with an initial $\boldsymbol{\kappa} \sim 0.7$ provided consistently good results across diverse conditions, which is what we set as its default value.

As a result, It is worth noting that we often do not perform adaptive estimation during the cell refinement, which is why the quality of the cell finding, but not the $Ca^{2+}$ activity traces, will depend on the initial choice of $\boldsymbol{\kappa}$. Therefore, $\boldsymbol{\kappa}$ is one of the first parameters we recommend optimizing for real data applications, though, historically, little-to-no fine-tuning has been needed across mice or imaging sessions that share the same experimental and imaging conditions.

#### §4. Benchmarking EXTRACT against the state-of-the-art cell extraction algorithms

Experiments 7-21 in the benchmark (**Table S1**) test EXTRACT's cell extraction abilities against existing state-of-the-art (or still regularly used) cell extraction and trace post-processing algorithms. Specifically, we performed the following comparisons:

- Comparisons with CAIMAN[4]: Experiments 7-12
- Comparisons with averaging inside regions of interests (ROIs) [13]: Experiment 13
- Comparisons with SEUDO[9]: Experiments 14-19
- Comparisons with CNMF[8] and ICA[3]: Experiments 20-21

**Exs 7-12: Benchmarking EXTRACT against CAIMAN on diverse conditions**

As discussed in the main text, two most commonly used cell extraction algorithms in the field are CAIMAN[4] and Suite2p[2]. Both algorithms use similar $L_2$ estimation based regression methods for cell extraction, though CAIMAN is explicitly validated to process one-photon $Ca^{2+}$ imaging movies[6]. Therefore, as a representative of currently utilized cell extraction pipelines, we picked CAIMAN to benchmark EXTRACT against.

*In experiments 7-10,* we used EXTRACT and CAIMAN to process a total of 800 simulated two-photon and one-photon $Ca^{2+}$ imaging movies. Given our focus on designing an algorithm that can scale datasets of the future, *e.g.*, those with densely packed cells (See **Figure 4**), we aimed to understand how these algorithms would perform with increasing cell density under various experimental and imaging conditions. Specifically, for each experiment, we simulated 20 one-photon and two-photon $Ca^{2+}$ imaging movies with varying total number of cells, $n_{cell} = (300, 600, 900, 1200, 1500)$, in a $400 \times 400$ µm$^2$ field of view (FOV). The experiments tested the following conditions:

- Experiment 7: Standard $Ca^{2+}$ imaging movies with $SNR_{min} = 4$, independently firing neurons, and negligible neuropil $\epsilon = 0.05$.
- Experiment 8: Low SNR $Ca^{2+}$ imaging movies with $SNR_{min} = 2.5$, independently firing neurons, and negligible neuropil $\epsilon = 0.05$.
- Experiment 9: $Ca^{2+}$ imaging movies with $SNR_{min} = 4$, correlated neurons, and negligible neuropil $\epsilon = 0.05$.
- Experiment 10: $Ca^{2+}$ imaging movies with $SNR_{min} = 4$, correlated neurons, and substantial amounts of neuropil contamination $\epsilon = 0.3$.

The results, presented in **Figures 3, S2, and S3**, validated EXTRACT’s superior cell extraction and generalization abilities across diverse conditions. Moreover, though EXTRACT matched or outperformed CAIMAN’s performance, the difference was more dramatic with the increased cell densities, in line with the results of Experiment 2 above.

*Experiment 11:* Next, to test whether EXTRACT can process very short $Ca^{2+}$ imaging movies, we simulated additional two-photon movies with increasing durations (from roughly 1 to 17 minutes), but otherwise using with the default parameters, *e.g.*, $n_{cell} = 600$ cells firing independently etc. The results, shown in **Appendix Figure 8**, have confirmed EXTRACT's ability to extract cells from very short $Ca^{2+}$ imaging movies, as little as c.a. one minute duration. In contrast, CAIMAN had decreased precision and recall values even in simulated $Ca^{2+}$ imaging movies with several minutes of equivalent imaging time.

*Experiment 12:* In our experiments so far, we used a particular configuration of CAIMAN and EXTRACT, locally fine-tuned to the condition of interest from a set of suggested hyperparameters (**Methods**). But, how generalizable are these results, *e.g.*, when both EXTRACT and CAIMAN are tuned to the optimal parameters? To answer this question, we sought to obtain a precision-recall (PR) curve for both algorithms by performing a grid search over their hyperparameters. The PR curve outputs the precision and recall values of cell finding in a continuum, tracing out an upper-bound curve.

To obtain the PR curve, we simulated a single two-photon $Ca^{2+}$ imaging movie using the default parameters and with neurons firing in a correlated manner. For CAIMAN, we performed a grid search across the following parameters:

- Number of cells per partition, K: (30,50,100,150,200)
- Half-size of partitions in pixels, rf: (20,40,60,80)
- Radius of average neurons in pixels, gSig: (3,4,5) and accordingly gSiz = 2*gSig+1
- Trace SNR threshold, min_SNR: (2.5,3,3.5)
- Space correlation threshold, rval_thr: (0.4,0.7,0.9)

- Rank of the background component, nb: (1,2,3)

By default, we picked p=1 (autoregressive model), frame_rate = 10Hz, decay_time = 1s to match the ground truth data. For EXTRACT, we performed a grid search across the following parameters:

- Number of maximum cells that can be initialized: (500,600,700,800)
- Initial values of $\boldsymbol{\kappa}$ (used without adaptive updates when estimating $\boldsymbol{S}$): (0.1,0.3,0.5,0.7,1)
- Minimum cell SNR: (3.5,4,4.5)
- Spatial corruption values: (1,2,3,4)
- Number of cell refinement iterations: (2,4,6,8,10)

In all cases, we used a single partition for EXTRACT, which is by default for small FOVs (See Supplementary Note 2 for algorithmic details on why this is feasible with EXTRACT). In contrast, when we tried to use a single partition for CAIMAN, we surprisingly received error messages for some combinations of the hyperparameters and were forced to use multiple spatial partitions.

The results, shown in **Figures 3B and S4A**, illustrate EXTRACT's superiority in cell finding, tracing out the upper left portion of the PR curve (with the upper left most point corresponding to a perfect precision and recall). The cell extraction outputs by CAIMAN not only had higher variance, but also were inferior. Consequently, the results of our benchmarking experiments cannot be explained by specific choices of hyperparameters, rather lie in the inherent differences between EXTRACT and CAIMAN.

**Ex 13: Suboptimality of averaging pixel activities inside ROIs**

One of the earlier approaches to cell extraction, especially when $Ca^{2+}$ imaging movies contained only few cells, involved hand-drawing circles around cells' spatial location, called regions of interests, and computing the $Ca^{2+}$ activity traces by averaging pixel activities therein (See for example[14]). Such methods have clear disadvantages compared to a simple least-squares estimates, which can be implemented with similar computational complexity, with an extra matrix inversion that can be pre-computed (See **Supplementary Note 1**). Yet, averaging pixel activities inside ROIs is still being implemented in the published algorithms today[13,15].

This experiment focuses on the superiority of computing continuous valued spatial profiles for cells over using hand-drawn binary locations, and thereby illustrates the short-comings of ROI averaging. To illustrate this concept, we simulated 20 two-photon $Ca^{2+}$ imaging movies with 1000 frames duration, $SNR_{min} = 2.5$, neurons firing in a correlated manner, $F = 0$, and otherwise using the default values. We used EXTRACT as a post-processing tool to perform robust regression and NNLS. To obtain the $Ca^{2+}$ activity traces, we used augmented ground truth spatial profiles such that $\hat{\boldsymbol{S}} = \boldsymbol{S}(\boldsymbol{S} > 0.4)$. We initialized both trace estimation algorithms with varying numbers of ground truth cells, *i.e.*, (50,100,150,200,250,300,350,400, 450,500,550,600) out of 600. As before, any cell that was not known to the trace estimation algorithm introduced non-Gaussian noise contaminants. To compute the binarized regions of interests, we considered the binarized versions of $\hat{\boldsymbol{S}}$, *i.e.*, $\hat{\boldsymbol{S}}_{binarized} = (\hat{\boldsymbol{S}} > 0)$.

As discussed in **Appendix Figure 2**, averaging pixel activities inside the regions of interest had the same $Ca^{2+}$ activity trace quality regardless of the contamination level, which corresponded to the lowest accuracy achieved with NNLS estimates under the highest level of contamination. On the other hand, once the contamination level decreased, *i.e.*, more neighboring and overlapping cells were identified by the cell extraction algorithm, NNLS estimates, but not those averaged within ROIs, asymptotically reached EXTRACT's accuracy. Therefore, even for real-time algorithms for which computation speed is particularly important, a least-squares based estimation procedure may provide higher accuracy $Ca^{2+}$ activity traces while having similar complexity to the methods computing activity averages within ROIs[13,15].

**Exs 14-19: Comparison of EXTRACT vs SEUDO as post-processing tools**

The suboptimality of estimated $Ca^{2+}$ activity traces by $L_2$ approaches has been known and studied in the literature, often leading to the development of trace post-processing algorithms[9,16,17]. The recently developed SEUDO[9] is the current state of the art approach, which is what we benchmarked EXTRACT against.

*Experiments 14-16:* SEUDO is designed to remove neuropil and other types of contaminants that cannot be explained by the extracted cells' activities. Previous work provided validity evidence in the form of human annotators[9], which may be biased and make mistakes in their annotations as previously shown[2] for the NeuroFinder benchmark[7]. In this work, we first set out to replicate these claims with controlled simulations.

As a first test, we simulated 10 two-photon $Ca^{2+}$ imaging movies, which contained an additional neuropil contamination similar to the one-photon $Ca^{2+}$ imaging movies, but milder. Specifically, for each movie, we first simulated the standard two-photon component with 150 cells, varying spiking amplitudes ($A_{var} = 1$), $n_t = 1000$ frames, high neuropil $\epsilon = 0.3$, correlated spiking with $p_{synch} = 0.5$ and $n_{synch} = [20, 40]$, otherwise using default parameters in a $160 \times 160\, \mu m^2$ FOV. Then, we simulated a secondary background movie with 40 cells, cell diameters ranging $[55, 90]\, \mu m$, no minimum distance requirement between cells, 0.5 Hz event rate, 3s decay rate, varying spiking amplitudes ($A_{var} = 1$) and minimum spike amplitude of 0.02, roughly matching the random noise in the original movie. Before adding to the original movie, we performed a band pass spatial filtering on the background movie using the default Butterworth filter described in the methods section (average cell radius 6, high/low pass cutoffs = 3). With the bandpass filtering and the spiking levels matching the noise s.d. of the original movie, the background movie resembled neuropil contamination in noisy two-photon $Ca^{2+}$ imaging movies that SEUDO was developed to process. These movies had higher local neuropil contamination compared to our standard two-photon $Ca^{2+}$ imaging movies, but had little-to-no global contamination as was the case for the simulated one-photon $Ca^{2+}$ imaging movies.

With these movies, we first performed a grid search for SEUDO across its hyperparameters, *i.e.*, $\sigma = (200, 20, 2, 0.2, 0.02, 0.002)$, $\lambda = (10^{-4}, 10^{-3}, 10^{-2}, 10^{-1})$, and $a = (1, 2, 3, 4, 5, 6, 7, 8)$. We also tried padSpace between 5 and 20, eventually settling to 5 pixels. Throughout all experiments in this work, we wrote our own, significantly faster, wrapper for SEUDO, which allowed us to parallelize it across multiple CPU cores. Our analysis revealed that SEUDO was able to properly reject the neuropil contamination when we allowed it access to full imaging duration, which is how we set up SEUDO's implementation. The alternative, *i.e.*, using the auto classification option of transient times and performing trace estimation only during those times[9], increased the processing speed but significantly decreased the accuracy.

Experiments 14-16 used $Ca^{2+}$ imaging movies simulated using the above formula and SEUDO optimized through the grid search. For experiment 14, we initialized the two-photon $Ca^{2+}$ imaging movies with varying percentages of cells' ground truth spatial profiles, (10, 20, 30, 40, 50, 60, 70, 80, 90, 100, 110, 120, 130, 140, 150) out of 150 cells. We post-processed these movies with SEUDO and EXTRACT to obtain the $Ca^{2+}$ activity traces. For experiment 15, we followed the same procedure as in experiment 14, but thresholded the spatial profiles (by 0.4) similar to Experiment 13. For experiment 16, we fixed the number of cells to be initialized to 120 (out of 150) and varied the threshold values across [0.1-0.9], emulating imperfect cell finding processes. Specifically, a higher threshold corresponds to higher deviation of cells' spatial profiles from the ground truth. The results, presented in **Appendix Figure 10**, validated SEUDO as a valid post-processing tool when cell finding is properly performed, which was nonetheless non-robust to variations in the estimated spatial profiles.

*Experiments 17-19:* Next, we tested SEUDO as a post-processing algorithm by utilizing the cells' spatial profiles obtained from existing cell extraction algorithms. To do so, we simulated two-photon $Ca^{2+}$ imaging movies using the same procedure (except for Experiment 19, see below) as in Experiments 14-16. Specifically, for each movie, we first simulated the standard two-photon component with varying spiking amplitudes ($A_{var} = 3$), $n_t = 3000$ frames, low neuropil $\epsilon = 0.05$, cells spiking independently, otherwise using default parameters in a $160 \times 160\,\mu m^2$ FOV. For experiment 17 (testing variations in SNR), we simulated the movies with 100 cells and varied the minimum SNR values of the cells across (2.5, 5). For experiment 18 (testing variations in cell density), we fixed the minimum SNR values of the cells to 4, but varied the numbers of cells across (40, 80, 120, 160, 200). Similar to before, for both of these experiments, we simulated a secondary background movie with 40 cells, cell diameters ranging $[28, 38]\ \mu m$, no minimum distance requirement between cells, 0.5 Hz event rate, 3s decay rate, varying spiking amplitudes ($A_{var} = 1$), and minimum spike amplitude of $0.005 \times SNR_{min}$. Before adding to the original movie, we once again performed a band pass spatial filtering on the background movie. For experiment 19 (testing no neuropil condition), we did not use a background movie, set the neuropil contamination to zero, $\epsilon = 0$, fixed the minimum SNR values of the cells to 4 and varied the numbers of cells across (40, 80, 120, 160, 200). For each condition, we simulated 20 distinct $Ca^{2+}$ imaging movies. The $Ca^{2+}$ imaging movie we used for optimizing SEUDO was taken from experiment 17, with a minimum cell SNR value of 5. Specifically, we performed a grid search for SEUDO across its hyperparameters, *i.e.*, $\sigma = (200, 20, 2, 0.2, 0.02)$, $\lambda = (10^{-4}, 10^{-3}, 10^{-2}, 10^{-1})$, and $a = (1, 2, 3, 4, 5, 6, 7, 8)$.

We processed these simulated movies with CNMF and EXTRACT to obtain cells' spatial profiles. These extracted filters, not the ground truth, are used to extract the $Ca^{2+}$ activity traces with SEUDO, NNLS, and robust regression. To allow fair comparisons, we used the median centered $\Delta F/F$ movie for all post-processing approaches and turned off the high-pass filtering for EXTRACT. Results, presented in **Figures 3G, H and S5 A, B, C, E, F**, validated SEUDO as a better post-processing tool compared to CNMF, in line with the findings by prior work[9], but also has shown SEUDO's inability to provide higher quality $Ca^{2+}$ activity traces once the spatial profiles were badly estimated. Moreover, SEUDO was not necessary, and suboptimal to robust regression, once cells' spatial profiles were estimated using the robust regression. Hence, these experiments confirmed that SEUDO was applicable for a narrow range of experimental non-idealities encountered with CNMF[8] or Suite2p[2], which are now mitigated with EXTRACT.

**Exs 20-21: Benchmarking EXTRACT against MATLAB based cell extraction algorithms**

To benchmark EXTRACT against the commonly used MATLAB based cell extraction algorithms, specifically ICA[3] and CNMF[8], we simulated another set of two-photon $Ca^{2+}$ imaging movies, 20 per condition, with following parameters:

- *Experiment 20:* Number of cells - $n_{cell} = 100$, spiking amplitude variation - $A_{var} = 1$, total number of frames - $n_t = 3000$, neuropil contamination - $\epsilon = 0.1$, FOV - $160 \times 160\ \mu m^2$, correlated spiking with $p_{synch} = 0.5$ and $n_{synch} = [5, 10]$, minimum cell SNR - ranging across (2.5, 3, 3.5, 4, 4.5, 5), otherwise default values.
- *Experiment 21:* Number of cells - $n_{cell} = 100$, spiking amplitude variation - $A_{var} = 1$, total number of frames - $n_t = 5000$, neuropil contamination - $\epsilon = 0.05$, FOV - $160 \times 160\ \mu m^2$, correlated spiking with $p_{synch} = 0.5$ and $n_{synch} = [5, 10]$, minimum cell SNR - (2, 3, 4, 5), with donut shaped cells, and otherwise default values.

As before, to allow fair comparisons, we used the median centered $\Delta F/F$ movie for ICA and turn off high-pass filtering, allowing only $\Delta F$ (median) subtraction, for EXTRACT. The results, presented in **Appendix Figure 9**, indicated that both CNMF and ICA led to inferior cell extraction results, whereas ICA surprisingly suppressed the CNMF despite being developed several years earlier. Yet, we observed that ICA was rather susceptible to the details of the simulated configurations, and failed to process some movies altogether (specifically, see Experiment 26 below). EXTRACT and ICA, but not CNMF, were able to identify high quality spatial profiles corresponding to donut shaped cells. Overall, though CNMF was easier to use, ICA provided higher quality cell extraction results; whereas EXTRACT often outperformed the two algorithms in this benchmark.

### §5. Validation experiments for EXTRACT's constraints and algorithmic choices

In **Supplementary Note 2**, we introduced several unique algorithmic choices and domain-knowledge inspired constraints for EXTRACT. Yet, such choices should always be validated. To achieve this, we designed the experiments 22-25, which tested the non-negativity constraint (*Experiments 22-23*), the use of spatial high-pass filtering (*Experiment 24*), and temporal downsampling of $Ca^{2+}$ imaging movies for faster and more accurate processing (*Experiment 25*). The details are provided below.

#### Ex 22: Empirical validation of the theoretical $Ca^{2+}$ activity trace estimation errors

In **Supplementary Note 1**, we provided a theoretical account of $Ca^{2+}$ activity trace estimation problem. As part of this analysis, we arrived at a simple expression for the theoretical trace estimation error that would be achieved by the least-squares estimate, given as:

$$E[||\boldsymbol{T} - \hat{\boldsymbol{T}}_{ls}||_2^2] = Tr[(\boldsymbol{S}^T\boldsymbol{S})^{-1}]\sigma_{iid}^2,$$

where $\sigma_{iid}$ is the standard deviation of the (zero mean) normally distributed noise in the pixel activities, $\boldsymbol{T}$ and $\boldsymbol{S}$ stand for cells' $Ca^{2+}$ activity traces and spatial profiles, respectively.

To empirically test this prediction, we simulated two-photon $Ca^{2+}$ imaging movies, 20 per condition, with 100 cells, $n_t = 1000$ frames in a $160 \times 160\ \mu m^2$ FOV, a minimum cell SNR of 5, spiking amplitude variation - $A_{var} = 1$, independent spiking of neurons, no neuropil $\epsilon = 0$, no global baseline $F = 0$, and with varying average cell radii, (2, 2.5, 3, 3.5, 4, 4.5, 5) s.d. in pixels. Using EXTRACT as a post-processing tool on these movies and using cells' ground truth spatial profiles, we computed the empirical and theoretical trace estimation errors for least-squares estimates, as well as the empirical trace estimation errors for NNLS and non-negative robust regression. The relationship between trace estimation errors, normalized by $\sigma_{iid}^2$, are presented in **Appendix Figure 1**. As expected, we observed a near perfect match between the predicted and empirical errors for the least-squares estimates, whereas both NNLS and non-negative robust regression had far lower errors compared to the least-squares estimate.

**Ex 23: Empirical validation of the non-negativity constraint**

EXTRACT, similar to prior cell extraction algorithms[4], utilizes a non-negativity constraint when estimating the cells' spatial profiles and $Ca^{2+}$ activity traces. As discussed in **Supplementary Note 1**, the use of non-negativity constraint can significantly decrease the variance in the least-squares estimates.

To validate the non-negativity constraint, we simulated several two-photon $Ca^{2+}$ imaging movies, 20 per condition. Specifically, we varied the minimum cell SNR values within (2, 2.5, 3, 3.5, 4), otherwise used the default parameters for independent spiking, and set $F = 0$. Initializing with 480 out of 600 cells' ground truth spatial profiles, we used EXTRACT as a post-processing tool to obtain $Ca^{2+}$ activity traces estimated by least-squares and NNLS. The results, shown in **Appendix Figure 3**, provided additional empirical evidence that non-negativity constraint may increase the quality of the estimated $Ca^{2+}$ activity traces, though our discussions in Experiment 5 suggest that non-negativity constraint need to be supplemented with an adaptive estimation of cells’ baseline activities.

**Ex 24: Empirical validation of the spatial high-pass filtering**

When performing cell extraction, several past cell extraction algorithms perform different forms of spatial filtering, either directly on the $Ca^{2+}$ imaging movie or the extracted $Ca^{2+}$ activity traces. Perhaps the most common version is the neuropil subtraction[2,17], in which a fraction of the neighboring pixel activities are subtracted from the pixel activities encapsulated by cells’ spatial profiles. Another commonly used approach is the ring model by CNMF-E[6], where the pixel activities on a large ring centered at the cells’ origin are subtracted from the final signal, similar to the neuropil subtraction but at a larger scale. However, neither of these approaches are interpretable and/or guaranteed to not explain away signals from the estimated $Ca^{2+}$ activity traces. In EXTRACT, we use a more principled approach of spatial high-pass filtering, in which the filter cutoff can be designed to prevent removing any pixel activity that is not correlated at least several times the typical cell radius (**Methods**).

To validate the use of spatial high-pass filtering for cell extraction, we simulated 20 two-photon and one-photon $Ca^{2+}$ imaging movies with high neuropil contamination $\epsilon = 0.3$, otherwise using the default parameters for correlated spiking between cells. We initialized EXTRACT with 480 (out of 600) cells’ ground truth spatial profiles and performed two cell refinement iterations to adjust the spatial profiles to the filtered $Ca^{2+}$ imaging movie (See **Supplementary Note 1** for a discussion on how spatial high-pass filtering affects the cells’ spatial profiles). With this procedure, we obtained the $Ca^{2+}$ activity traces as we varied the high-pass cutoff values between (2, 3, 4, 5, 6, 7, 8, 9, 10), where 10 corresponds to mildest filtering of contaminants that are correlated in space roughly 10 times larger than a typical cells’ size. The results, presented in **Appendix Figure 4**, have validated the need for mild high-pass filtering, without which the neuropil contamination would deteriorate the estimated $Ca^{2+}$ activity traces.

**Ex 25: Empirical validation of the temporal downsampling during cell finding and cell refinement**

Several prior algorithms perform temporal downsampling to speed up the cell extraction process[2,4], a design that we also implemented within EXTRACT. In our case, EXTRACT may utilize a temporally downsampled $Ca^{2+}$ imaging movie for the cell finding and cell refinement modules, but the final $Ca^{2+}$ activity traces are computed from the original movie, not leading to any decreased temporal resolution in the final results.

To test the effects of temporal downsampling, we simulated two-photon $Ca^{2+}$ imaging movies, 10 per condition. Specifically, we simulated movies with 30 Hz frame rate, $n_t = 72000$ number of frames, no spiking amplitude variation - $A_{var} = 0$, low neuropil $\epsilon = 0.05$, varying levels of minimum cell SNRs (1.5, 2.5, 4), independent spiking cells, otherwise using the default conditions. We processed these movies with temporal downsampling factors varying within (1, 2, 3, 4, 5, 6, 8, 10, 15, 30), *i.e.*, from 30Hz to 1Hz. The results, shown in **Fig. S6C**, indicated that EXTRACT benefits from temporal downsampling in terms of both speed and accuracy and is able to perform cell extraction in $Ca^{2+}$ imaging movies sampled down to 2 Hz without significant deterioration of the accuracy. Moreover, the cell extraction results with 1 Hz movies were still accurate, comparable to the stereotypical accuracies achieved by CAIMAN, though we recommend downsampling down to 2 Hz as the optimal value.

#### §6. Benchmarks testing the limits of EXTRACT

Experiments 26-30 test the behavior of EXTRACT in limiting cases including tiny and very large cells (*Experiment 26*), increased spatiotemporal correlations in cell activities (*Experiments 27*), cells encapsulated by other cells (*Experiment 28*), and under severe motion artifacts (*Experiments 29 and 30*). The details are provided below.

**Ex 26: Comparisons with varying cell radii**

In our benchmark, almost all experiments were performed using the same spatial resolution, *i.e.*, single cell spatial profile is a Gaussian function with a standard deviation roughly 4 pixels. But, not all experiments use the same spatial resolution, some large scale $Ca^{2+}$ imaging movies may have only a few pixels for each cell. Therefore, to test whether EXTRACT can identify cells in extreme (very high to low) resolutions, we simulated several 20 two-photon $Ca^{2+}$ imaging movies with following specifications: Average radius of cells - varying (2.5, 3, 3.5, 4, 4.5, 5) s.d. in pixels with appropriately adjusted number of cells to keep the total cell area approximately the same - $n_{cell} = (160, 112, 82, 63, 50, 40)$, spiking amplitude variation - $A_{var} = 1$, total number of frames - $n_t = 3000$, low neuropil - $\epsilon = 0.1$, FOV - $100x100$ pixel$^2$, cells spiking in a correlated manner with $p_{synch} = 0.5$ and $n_{synch} = [5, 10]$, otherwise using the default values. The results, presented in **Appendix Figure 6**, showed negligible variations in EXTRACT's outputs across diverse resolutions, validating EXTRACT's use for when cells occupy as little as tens of pixels.

**Ex 27: Robustness to increased spatiotemporal correlations**

Another extreme scenario, which is encountered in practice particularly with imaging of Purkinje cells, is when cells are firing in a highly spatiotemporally correlated manner. In such cases, cell extraction algorithms may have problems identifying individual cells within the global activations.

To test whether EXTRACT can perform cell extraction despite highly correlated neural activities, we simulated $Ca^{2+}$ imaging movies with increasing spatiotemporal correlations. Specifically, we simulated two sets of two-photon $Ca^{2+}$ imaging movies, 20 per condition, with long ($n_t = 5000$ frames) and short ($n_t = 1000$ frames) imaging durations. The movies included neurons firing in a correlated manner, with the correlation probabilities varying $p_{synch} = (0,\ 0.1,\ 0.2,\ 0.3, 0.4,\ 0.5,\ 0.6,\ 0.7,\ 0.8,\ 0.9)$ and $n_{synch} = [200, 600]$, and otherwise using default conditions. Given that large numbers of neurons were able to fire jointly, spatiotemporal correlations in activity was controlled by the joint spiking probability $p_{synch}$.

We used EXTRACT and CAIMAN to process these simulated $Ca^{2+}$ imaging movies. Owing to high spatiotemporal correlations, we used high-pass filtering to identify the cells' spatial profiles with EXTRACT, but turned off the high-pass filtering when obtaining the final $Ca^{2+}$ activity traces. This is because high-pass filtering could explain away correlated firings of neurons, which is one of the few cases where spatial filtering of any kind, including the neuropil subtraction, can hurt significantly.

The results, presented in **Fig. S6A,B**, are in line with our empirical studies on real datasets with Purkinje cells (**Fig. S8**), and validated EXTRACT's use for cases when cells are firing in an extremely correlated manner. Even when cells were firing almost always jointly (90% of the time), EXTRACT was able to identify the individual cells as long as the simulated $Ca^{2+}$ imaging movies were long enough such that individual cells fired a few spikes independently.

**Ex 28: Demixing of activities in the extreme limit - cells encapsulated by other cells**

To date, existing cell algorithms have shown surprising capabilities of demixing $Ca^{2+}$ activity traces from cells that are largely overlapping[8]. Yet, in this experiment, we tested a new limit: a smaller cell is encapsulated by another out-of-focus bigger one. Specifically, we simulated two separate movies in a $30 \times 30$ pixels$^2$ FOV, each with one cell. The smaller cell had a diameter of 6 pixels, the larger cell was as large as the full FOV. Both movies had a minimum cell SNR of 5, otherwise were simulated using default values. The final movie was obtained by summing two movies and adding the global background, $F = 1$.

We processed this specially designed movie with EXTRACT. Despite allowing EXTRACT to find up to 20 cells, EXTRACT accurately identified *only* two cells, discarding any spurious duplicates and setting a new record for the cell extraction process. The results are presented in **Fig. 3F** and may be crucial for one-photon $Ca^{2+}$ imaging experiments.

**Exs 29-30: Testing EXTRACT's cell extraction quality under rigid residual motion**

Like the current state of the art cell extraction algorithms[2,4,8], EXTRACT also requires correcting the brain's motion before extracting cells. This requirement is rooted in EXTRACT's assumption that the movie consists of stationary cells' activities. But, these algorithms are not perfect and sometimes can lead to few to sub-pixel errors. What happens then?

*Experiment 29:* To answer this question and test the ability of EXTRACT to handle rigid motion, we simulated two-photon $Ca^{2+}$ imaging movies, 20 per condition, with low snr (minimum cell SNR = 2.5), correlated spiking, $n_t = 1000$ frames, with $F = 0$, and otherwise default parameters. To apply rigid shifts, which may be at the level of subpixels, we sampled a random number for each frame. If the random number was below $0.05$ and no shift had been applied within a 5 frame radius, we induced a shift that exponentially decayed over the course of 10 time frames, *i.e.*, 1s. The shift was randomly chosen for both $x$ and $y$ dimensions using a uniform random distribution between [-max shift, max shift], where the maximum shift is varied between 0 and 10 pixels. The shifts were applied using the shift_reconstruct function from the NormCorre pipeline[18]. Then, we initialized the movies with 480 (out of 600) cells' ground truth spatial profiles and extracted the $Ca^{2+}$ activity traces via the robust regression, NNLS, and the least-squares. For this experiment, EXTRACT was not allowed to use high-pass filtering. The results are presented in **Fig. S6D**, and indicated that EXTRACT was able to handle rigid shifts up to few pixels, though we strongly recommend correction of the brain's motion with an appropriate motion correction algorithm[18].

*Experiment 30:* Similar to before, we simulated several two-photon $Ca^{2+}$ imaging movies, 20 per condition, with 100 cells, $n_t = 3000$ frames, on a $160 \times 160\ \mu m^2$ FOV, with independent spikes, $F = 0$, otherwise using default parameters. Unlike Experiment 29, in this experiment, the cell extraction algorithms were required to identify the cells' spatial profiles as well. EXTRACT, once again, was not allowed to use the high-pass filtering to prevent unfair advantage. Using the extracted $Ca^{2+}$ activity traces, we used exponential deconvolution, with 5% amplitude threshold, to obtain the event times; and employed support-vector machines to decode the frames with motion from the $Ca^{2+}$ activity traces and the extracted events.

The results are presented in **Appendix Figure 7** and are in line with the results from Experiment 29. Specifically, motion deteriorated the quality of cell extraction results for all algorithms, yet subpixel motion was tolerable. However, the random motion could be decoded from the $Ca^{2+}$ activity traces obtained by EXTRACT and NNLS, but not from the $Ca^{2+}$ events. In contrast, despite having low quality $Ca^{2+}$ activity traces to begin with, CNMF did not encode the random motion in the estimated $Ca^{2+}$ activity traces, likely due to the simultaneous convolution and deconvolution performed with the exponential kernel. This observation provides an intuition for future work. Specifically, algorithms developed to process $Ca^{2+}$ imaging movies with substantial motion may consider incorporating time-consistency components that can lead to ignoring instantaneous drops due to motion.

Overall, Experiments 29 and 30 showed that cell extraction requires correction of the brain's motion; yet remaining subpixel motion can often be tolerated.

#### §7. Benchmarks testing the fast solver of EXTRACT

Experiments 31-33 test the speed and scaling of EXTRACT with increasing number of cells (*Experiment 31*) and increasing FOV (*Experiment 32 and 33*). The details are provided below.

**Exs 31-32: Benchmarking EXTRACT's ADMM based solver**

One of our novel contributions with EXTRACT is its fast solver, which minimizes a constrained optimization problem using a second-order optimization routine that has the complexity equivalent to gradient descent. We provided theoretical details in **Supplementary Note 2** and here discuss the experiments we conducted to validate these claims.

*Experiment 31:* To test how fast EXTRACT is with respect to Matlab's non-negative least-squares solver, *i.e.*, the lsqlin function, we simulated two-photon $Ca^{2+}$ imaging movies, 20 per condition. Specifically, we simulated movies with a FOV of $100 \times 100$ pixels$^2$, varying number of cells$n_{cell} = (50, 100, 150, 200, 250, 300)$, $n_t = 18000$ frames, correlated spiking with $p_{synch} = 0.5$ and $n_{synch} = [10, 20]$, minimum cell distance of 1 pixel, no neuropil $\epsilon = 0$, otherwise using default parameters with $F = 0$. We used EXTRACT as a NNLS solver, and compared it to lsqlin, which was parallelized across individual frames. To test specifically the ADMM based solver and the subpartitioning routines, we performed the pixel deletion (See **Supplementary Note 2**) for the Matlab's NNLS solver as well. The latter was allowed to utilize 10 CPU cores, whereas EXTRACT used a single GPU. The results, shown in **Fig. S6E**, indicated several factors of speed improvement with EXTRACT's novel solver compared to the MATLAB's native solver.

*Experiment 32:* Similar to Experiment 31, we simulated two-photon $Ca^{2+}$ imaging movies, 20 per condition, but this time aimed to understand the scaling of both solvers with increasing FOV size. We simulated movies with varying number of cells $n_{cell} = (25, 100, 225, 400, 625)$, correspondingly increasing FOV size $(50^2, 100^2, 150^2, 200^2, 250^2)$ pixels$^2$, $n_t = 18000$ frames, correlated spiking, minimum cell distance of 1 pixel, no neuropil $\epsilon = 0$, otherwise using default parameters with $F = 0$.

To process these movies, we once again used EXTRACT’s ADMM based solver to perform NNLS, and compared the results to lsqlin, which was parallelized across individual frames using 10 CPU cores. The results, shown in **Fig. S6E**, indicated up to two orders of magnitude speed up for larger FOVs with EXTRACT’s ADMM based solver, providing additional evidence for EXTRACT’s scalibility.

**Ex 33: Benchmarking EXTRACT’s speed against SEUDO as a post-processing tool**

To test the speed and the scaling ability of SEUDO as a post-processing tool, we performed two sub-experiments. In the first experiment, we simulated 20 two-photon movies with following properties: Number of cells - $n_{cell} = 150$, spiking amplitude variation - $A_{var} = 1$, total number of frames - $n_t = 1000$, high neuropil - $\epsilon = 0.30$, FOV - $160 \times 160\, \mu m^2$, correlated spiking with $p_{synch} = 0.5$ and $n_c = [20, 40]$, $F = 1$, otherwise using default values. We also simulated the background following the recipe in Experiments 14. We parallelized SEUDO across 18 CPU cores and obtained the $Ca^{2+}$ activity traces of varying numbers of cells (from 10 to 150). In contrast, for EXTRACT, we used only a single GPU. In the second experiment, we simulated $Ca^{2+}$ imaging movies with increasing numbers of frames, *i.e.*, $n_t = (100, 500, 1000, 3000, 5000)$, otherwise same as above. We initialized both EXTRACT and SEUDO with the 50 (out of 150) cells’ spatial profiles and estimated their $Ca^{2+}$ activity traces.

The results, presented in **Fig. S5D**, illustrated EXTRACT’s scaling properties in comparison to SEUDO. Specifically, with the increasing number of frames and/or cells, EXTRACT became two orders of magnitude faster than SEUDO as a post-processing tool.

**§8. Computer specifications for the benchmark**

To process the simulation movies in this massive benchmark in a timely manner, which nonetheless would take a few months even in the most parallelized format, we used a combination of computers. Some experiments were parallelized across computers for faster processing. The computers had the following specifications:

- A computer with two NVIDIA GeForce RTX 3090 GPUs and an Intel Core(TM) i9-10980XE processor with 18 CPU cores (*Speed benchmarking experiment 33*)
- A computer with two NVIDIA GeForce RTX 3080 Ti GPUs and an Intel Core(TM) i9-10980XE processor with 18 CPU cores
- A computer with an Nvidia Geforce RTX 3090 Ti GPU and Intel Core i9-9900X Skylake X 10-Core processor (*Speed benchmarking experiments 31 and 32*)
- A computer with Nvidia Geforce RTX 2070 GPU and Intel Core i9-9900X Skylake X 10-Core processor
- Apple Macbook Air with M1 chip
- Stanford High Performance Computing Cluster

### §9. Conclusion

In this note, we provided the details on the extensive simulation benchmark we utilized to test EXTRACT's capabilities and benchmark it against the existing cell extraction algorithms. The code for the simulation movies is provided as part of our Github code, and interested users can perform additional experiments to test EXTRACT's and other cell extraction algorithms' capabilities on further experimental and imaging conditions.

### Supplementary Note 5: "User manual for EXTRACT"

### §1. Overview

This user manual aims to provide the basic principles for *how to efficiently and effectively use* EXTRACT, our tractable and robust automated cell extraction tool for $Ca^{2+}$ imaging presented in the main text. For the purpose of this manual, we assume no detailed understanding of robust regression or other technical details presented in the main text. Instead, we will consider EXTRACT as a black box algorithm, controlled by a set of parameters, which takes as an input the $Ca^{2+}$ imaging movie and outputs the $Ca^{2+}$ activity traces of the identified cells in the movie and their spatial profiles. The inner workings of EXTRACT are mainly relevant for those who wish to develop similar algorithms; whereas a phenomenological understanding of EXTRACT's modules has been enough for many users to date to effectively and efficiently utilize EXTRACT. Below is a summary of the content for each section.

**§2** introduces an example code (**Tutorial 1**) to quickly start using EXTRACT. Here, on a simulated $Ca^{2+}$ imaging movie, we showcase some of the most helpful features of EXTRACT, *e.g.*, how to watch the movie before processing, how to set some crucial hyperparameters, and how to perform post-extraction quality controls. **§3** provides an overview of EXTRACT's modules in a user-friendly way and a full list of key hyperparameters that users need to be familiar with. Moreover, this section also contains a short tutorial regarding parallel computing and graphics processing unit utilization (**Tutorial 2**). **§4-6** introduce new tutorials (**Tutorials 3-6**) to discuss the effect of key hyperparameters of each module in detail. Specifically, we propose a 2-step optimization procedure for EXTRACT's hyperparameters, in which the modular structure is exploited to optimize the hyperparameters in a modular manner. The tutorials illustrate how to systematically and easily perform this procedure to quickly fine-tune the hyperparameters and include additional tips and tricks for processing challenging, particularly low SNR, movies with EXTRACT. **§7** concludes with final remarks. While working on the tutorials, we recommend keeping **Figure S9** nearby, which provides a visual summary of this user manual.

### §2. Code for a quick start

Throughout this user manual, whenever we refer to a variable or a code piece in MATLAB, we denote them with a distinct font. For example, the variable representing the input movie is referred to as `M`. With that, in essence, EXTRACT can be compactly run with a single line of code:

```
output = extractor(M,config);
```

In this section, we provide the basic information needed to quickly start using EXTRACT. Here, we first introduce and discuss EXTRACT's inputs and outputs, and then provide a code walkthrough on a simulated $Ca^{2+}$ imaging movie (Tutorial 1).

### 2.1. EXTRACT inputs

The EXTRACT algorithm has two inputs. The first input, `M`, corresponds to the movie being processed. It can either be a three-dimensional movie matrix, with the time component being the third dimension, or a command (see below) that points to the location of the movie. The second input is a structure array that contains the configuration parameters for EXTRACT to use. This structure array can be passed as an empty array, in which case EXTRACT uses the default parameters.

#### Inputting the $Ca^{2+}$ imaging movie into EXTRACT

As noted above, the input movie, `M`, can be provided as a three-dimensional matrix. For the second option of inputting the movie as a command, we will discuss two options. For these discussions, we will use an example h5 dataset, whose file name is `example.h5` and the movie is contained inside the dataset `/mov`.

The first option is to input the movie in a single string format. If it is inputted as a string, the string must be in the format `filepath:dataset`, where the dataset name should not contain a colon (:), though having it in the filepath is fine. For the example movie, the string would be:

```
M = 'example.h5:/mov';
```

As an alternative option, especially if the dataset name contains a colon (:), then `M` can be inputted as a cell such that the first component would include the path, whereas the second component would include the dataset name. In our example, this would be:

```
M = {}; M{1} = 'example.h5'; M{2} = '/mov';
```

If the movie is not within the direct path, but is rather inside a folder, say `C:\movie_folder`, then the file path in both cases would be updated to `'C:\movie_folder\example.h5'` for both options, *e.g.*, `M = 'C:\movie_folder\example.h5:/mov';`

**Structure array containing EXTRACT's configurations**

The configurations are inputted into the EXTRACT in the form of a structure array in MATLAB. We often refer to this array as `config`. We discuss the most important configuration parameters below, which are also illustrated in **Figure S9** in the main text.

- `avg_cell_radius`: An order of magnitude estimate for an average cell's radius in the movie. This parameter sets the scale on which EXTRACT will operate. For example, this parameter is used for normalization during the application of the high-pass filtering or for several quality metrics (See **§3** for details). Therefore, though setting it to a significantly larger or lower value will impair performance, EXTRACT is fairly robust to the variations in this parameter. A simple and effective way to estimate this parameter would be to plot the maximum projections of the $Ca^{2+}$ imaging movie (**§4**). Default: 6 pixels.
- `num_partitions_x` and `num_partitions_y`: User specified number of spatial partitions in x and y dimensions of the movie, often selected for large-scale $Ca^{2+}$ imaging movies to minimize the RAM usage. If the user does not decide on particular values, EXTRACT performs partitioning automatically (once any side length of the movie exceeds 512 pixels). Practically, we recommend picking the partition sizes such that each spatial chunk contains around thousands of cells, or each partition's file size is around a quarter of the full RAM memory, whichever is smaller. EXTRACT performs the stitching of the results automatically once all partitions are processed. Default: none.
- `cellfind_max_steps`: The maximum number of cells that can be initialized during the cell finding of EXTRACT in each spatial partition. The default is 1000, which may need to be adjusted depending on the particular dataset. Specifically, it can be increased so that EXTRACT finds more cells per partition or decreased to prevent increased false positives.
- `max_iter`: The number of cell refinement iterations (See **Figure 2**). Often, 5-10 iterations are enough to cull out false-positives, whereas too many iterations can lead to unnecessarily increased runtimes (**§5** and **§6**). Default: 6 iterations.
- `cellfind_min_snr` and `thresholds.T_min_snr`: Two different definitions of signal-to-noise ratio for cells' estimated $Ca^{2+}$ activity traces. The former is used only during cell finding, whereas the letter is used for both cell finding and cell refinement. A cell's estimated signal quality should be higher than these values. For `cellfind_min_snr`, a typical value of 1 is reasonable for most datasets. Increase this value to decrease false positives during cell finding with no effect on the cell refinement procedure. For

`thresholds.T_min_snr`, a default value is 7 (and lowest acceptable value is 3). This parameter is often very effective to cull away false-positives during cell refinement (**§6**).

- `trace_output_option`: This parameter decides which type of solver EXTRACT will use to produce the final estimated $Ca^{2+}$ activity traces (**§7**). Viable options include: `'nonneg'`, `'baseline_adjusted'`, `'no_constraint'`, `'nonnegative_least_squares'`, `'least_squares'`.

These parameters are perhaps the most immediately relevant ones for an end user to quickly start using EXTRACT, which is what Tutorial 1 focuses on. There are several other parameters relevant to the users, including the robustness parameter $\boldsymbol{\kappa}$, which is adaptively optimized and thus not necessarily needs to be tuned by a first-time user. We will discuss these in the following sections of this user manual with additional tutorials.

#### 2.2. EXTRACT outputs

EXTRACT outputs are compactly saved inside a structure array, which has four major fields: `'config'`, `'info'`, `'spatial_weights'`, and `'temporal_weights'`. `Config` contains the configuration parameters used by EXTRACT, whereas `info` includes some cell extraction statistics that are not directly relevant for a first time user. The remaining fields are three and two-dimensional matrices, respectively:

- `spatial_weights` contains the spatial profiles and locations of extracted cells and has the shape [movie_height x movie_width x number_of_cells_found].
- `temporal_weights` contains the inferred $Ca^{2+}$ activity traces for each cell and has the shape [number_of_movie_frames x number_of_cells_found].

In the main text and other supplementary notes, (transformed versions of) these matrices have often been referred to with the notations $\boldsymbol{S}$ and $\boldsymbol{T}$, respectively.

#### 2.3. Tutorial 1: A quick start to EXTRACT

In this first tutorial, we will work on a simulated $Ca^{2+}$ imaging movie. Our main goal is to get the first time users started with EXTRACT, though future tutorials will focus on further important hyperparameters. The walkthrough below is provided in the Github as a live script, which also contains the necessary data files for running the tutorial.

**Running EXTRACT**

Let us start with a simple cell extraction procedure. The code below uses the most important parameters we discussed above and performs a basic cell extraction:

```
load('example.mat');
config=[];
config = get_defaults(config);
config.avg_cell_radius=7;
config.trace_output_option='no_constraint';
config.num_partitions_x=1;
config.num_partitions_y=1;
config.use_gpu=0;
config.max_iter = 10;
config.cellfind_min_snr=0;
config.thresholds.T_min_snr=10;
output=extractor(M,config);
```

Fortunately, since this is a simulation movie, we can quantify the quality of cell extraction. Within the ‘example.mat’ file provided as part of the tutorial, we have cells' ground truth $Ca^{2+}$ activities, T_ground, and the spatial profiles, S_ground. To compare the cell extraction results, we first need to compute the same quantities for the extracted cells:

```
T_ex = output.temporal_weights';
S_ex = full(output.spatial_weights);
[h,w,k]=size(S_ex);
S_ex=reshape(S_ex,h*w,k);
```

Note that there are 20 ground truth cells in the provided simulated $Ca^{2+}$ imaging movie. The code above also outputs 20 cells. But, do they match? Did we find all the cells, or did we find some duplicates or spurious ones? To check this, we can use a spatial correlation based function (See **Methods**) that matches the extracted cells to the ground truth:

```
idx_match = match_sets(S_ex, S_ground,0.8);
```

As can be seen from the accompanying tutorial code, EXTRACT indeed finds all the cells.

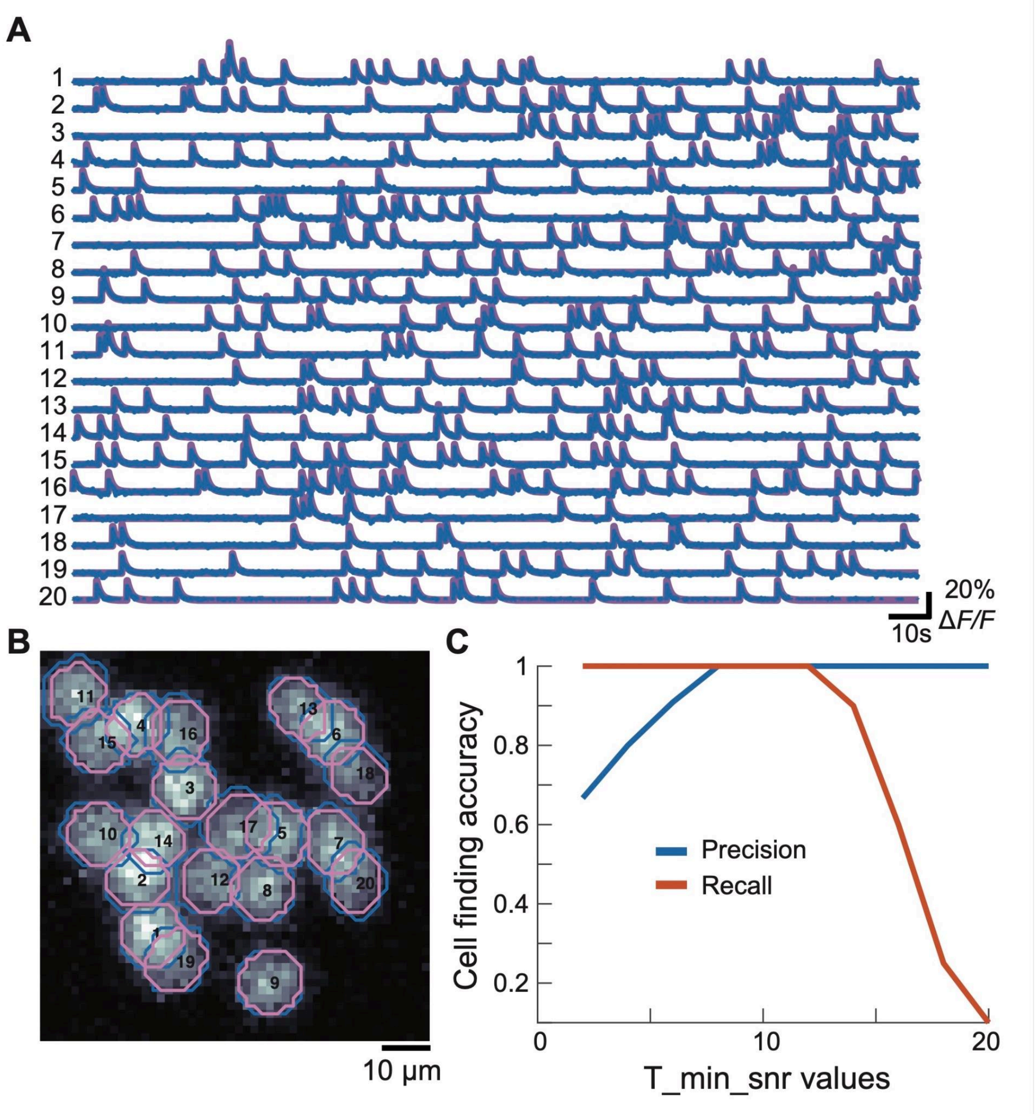

**Appendix Figure 11. Cell extraction results for a simulated Ca2+ imaging movie.**
The extracted cells’ $Ca^{2+}$ activity traces (**A**) and spatial profiles (**B**) superposed on top of the ground truth cells. **C** A simple parameter sweep experiment reveals the importance of correctly, but not necessarily finely, tuning the expected SNR value for the cells.

#### Visualizing cell extraction outputs

Using the following code, we can visualize the cell extraction outputs, which is shown in **Appendix Figure 11 A, B**:

```
color_extract = [0 0.4470 0.7410];
color_gt      = [144 103 167]./255;
color_l2 = [1,0.5,1];
```

```
plot_stacked_traces_double(T_ground(idx_match(2,:),:),...
T_ex(idx_match(1,:),:),1,{color_gt,color_extract},[],[],{5,3});
ims_ex = reshape(S_ex,h,w,[]);
ims_g = reshape(S_ground,h,w,[]);
max_im = max(M,[],3);
plot_simulated_cellmap(ims_g,...
max_im,ims_ex(:,:,idx_match(1,:)),color_extract,color_l2)
```

**Importance of SNR parameter**

Having shown the results of an example, well optimized, cell extraction, we next show the importance of correctly, not necessarily finely, choosing one of the most important parameters: `thresholds.T_min_snr`. This parameter needs to be adjusted to allow low SNR cells to be picked up or extremely low SNR garbage to be discarded. Using the code below, we can test how very low or very high values of this parameter affect the cell extraction quality.

```
thr_all = linspace(2,20,10);
precision = zeros(1,size(thr_all,2));
recall    = zeros(1,size(thr_all,2));
for i = 1:size(thr_all,2)
   config.thresholds.T_min_snr=thr_all(i);
   config.verbose = 0;
   output=extractor(M,config);
   S_ex=reshape(full(output.spatial_weights),h*w,[]);
   idx_match = match_sets(S_ex, S_ground,0.8);
   precision(i) = size(idx_match,2)/size(S_ex,2);
   recall(i) = size(idx_match,2)/size(S_ground,2);
   fprintf('%d finished.\n',i);
end
```

The results are shown in **Appendix Figure 11C.** As can be seen from the figure, choosing a very low `T_min_snr` leads to high recall, but can lead to diminished precision. This is because garbage/duplicate cells cannot be discarded despite having very low SNR values. On the other side, increasing `T_min_snr` too much can lead to several actual cells being discarded, leading to decreased recall values. Fortunately, the exact ground truth value does not need to be matched, as here flexible choices, *i.e.*, those between 8-12, all provided perfect cell extraction results. Empirically, we find that `T_min_snr` may need to be adjusted all the way down to ~3 for low SNR movies, whereas a value of 7-10 would be preferable to more effectively discard garbage for moderate to high SNR movies.

### §3. Introduction to EXTRACT: Understanding the modules and hyperparameters

In this section, we provide a comprehensive list of the hyperparameters that an EXTRACT end user should be broadly familiar with. While the list is arguably long, most hyperparameters are self explanatory (*e.g.*, `skip_dff` parameter that controls whether the $\Delta F/F$ transform should be applied during the preprocessing module). In Figure **2** of the main text, we provided a broad overview of EXTRACT's internal modules: preprocessing, cell finding, cell refinement, and final robust regression. Fortunately, we designed almost all of the hyperparameters by keeping this modularity in mind, which is why we will introduce them below in batches. We note that some parameters were already listed in **§2**, but for completeness we list them here again.

The default values for all hyperparameters can be found inside the function `get_defaults.m`. Similar to this user manual, the parameters there are also listed in batches depending on the module they control (also see **Figure S9**). Whenever in doubt, we recommend checking the exact spelling of the hyperparameters with this script. We advise that users do not change this script, the defaults are picked to be most appropriate for a general set of movies. Finally, please note that this script will not overwrite user provided configurations.

#### 3.1. General control hyperparameters

There are 16 general control parameters immediately relevant for EXTRACT users, which we list below:

- `avg_cell_radius`: An order of magnitude estimate for an average cell's radius in the movie. This parameter sets the scale on which EXTRACT will operate. For example, this parameter is used for normalization during the application of the high-pass filtering or for several quality metrics (See, *e.g.*, the preprocessing module below). Therefore, though setting it to a significantly larger or lower value will impair performance, EXTRACT is fairly robust to the variations in this parameter. A simple and effective way to estimate this parameter would be to plot the maximum projections of the $Ca^{2+}$ imaging movie (**§4**). Default: 6 pixels.

- `downsample_time_by`: For cell finding and refinement purposes only, EXTRACT can use a temporally downsampled version of the movie. This speeds up the cell extraction process and tends to increase the cell finding quality. In general, we recommend setting this parameter such that the downsampled movie has approximately few Hz sampling rate. For example, if the original movie is 20Hz, a reasonable value would be 5. Default: 1.
- `dendrite_aware`: If the movie includes dendrites and/or other non-conventional regions of interest, set this to 1. When this parameter is on, EXTRACT skips several checks regarding the shape and the size of the estimated cell profiles (See Section 3.4 below). Default: 0.
- `Remove_duplicate_cells`, `T_dub_thresh` and `S_corr_thresh`: If the movie is partitioned, it is possible that the same cells are found in two separate slightly overlapping partitions. These parameters control the removal of duplicate cells in these overlap regions. `Remove_duplicate_cells` is a control flag, 'true' by default. The other two parameters set the correlation thresholds for the overlapping cells' activities and spatial profiles. When both are surpassed between two cells, these cells are considered duplicates of each other and the cell with the higher area is retained.
- `use_gpu`: If you have a graphics processing unit (GPU), set this parameter to 1.
- `parallel_cpu`: Boolean flag for parallel processing movie partitions over the CPU cores. EXTRACT either parallelizes across the CPU cores or uses the GPU. Often, GPUs are faster. Default: 0.
- `multi_gpu`: Same as before, but for multiple GPUs. Default: 0.
- `num_workers`: This parameter is relevant when either `parallel_cpu` or `multi_gpu` is on. When using parallel processing, this parameter can be used to set the desired number of CPU cores or GPUs. Default: # of available cores/GPUs to Matlab - 1.
- `use_sparse_arrays`: If this is set to 1, the cells' spatial profiles stored inside the structure array `output.spatial_weights` will be stored as sparse matrices, instead of single format. Turn this flag on when processing large movies. Default: 0.

- `compact_output`: If set to `true`, then the output will not include secondary, but sometimes useful, cell extraction information. Turning this flag on often reduces the size of the output file substantially. Default: 0.
- `hyperparameter_tuning_flag`: If this flag is on, the cell extraction process stops after cell finding and a single cell refinement step. See **§6** and **Tutorial 5** for additional details for how to use this. Default: 0.
- `verbose`: Controls the logs that are outputted to the console. 0: No output at all. 1: EXTRACT emits basic details during the signal extraction process within each spatial patch. 2: Similar to 1, but EXTRACT provides a rather detailed summary (default when serial processing). 3: Only the most basic general information and no particular information from within the spatial patches (default when parallel processing).
- `num_partitions_x` and `num_partitions_y`: User specified number of spatial partitions in x and y dimensions of the movie, often selected for large-scale $Ca^{2+}$ imaging movies to minimize the RAM usage. If the user does not decide on particular values, EXTRACT performs partitioning automatically (once any side length of the movie exceeds 512 pixels). Practically, we recommend picking the partition sizes such that each spatial chunk contains around thousands of cells, or each partition's file size is around a quarter of the full RAM memory, whichever is smaller. EXTRACT performs the stitching of the results automatically once all partitions are processed. Default: none.

#### 3.2. Preprocessing module

Controlling the preprocessing module is rather straightforward, which involves four relevant parameters listed below:

- `preprocess`: Turns the full preprocessing module on or off. Default: 1.
- `skip_dff`: To perform the $\Delta F/F$ transformation of the $Ca^{2+}$ imaging movies in a numerically stable manner, EXTRACT subtracts the static baseline of the movie but does not perform the division explicitly. Instead, we store the static baseline values in a separate array, which is used to normalize the final $Ca^{2+}$ activity traces. Setting this flag to 0 skips the $\Delta F$ subtraction and assigns $F = 1$ to all pixels. Default: 0.
- `F_per_pixel`: When the preprocessing module is skipped, this parameter (a matrix) defines the static baseline values for each pixel. Default: none.

- spatial_highpass_cutoff: This parameter defines the cutoff values of the spatial high-pass filter applied to the $Ca^{2+}$ imaging movie. The exact cutoff value is proportional to avg_cell_radius/spatial_highpass_cutoff, normalized with respect to the average cell radius in the movie. A larger value of this parameter leads to mild filtering, with no filtering being performed when it is set to infinity. Default: 5.

#### 3.3. Cell finding module

The following are the parameters associated with the cell finding module:

- spatial_lowpass_cutoff: This parameter defines the cutoff values of the spatial low-pass filter applied to the $Ca^{2+}$ imaging movie for the cell finding purposes only. The exact cutoff value is proportional to avg_cell_radius*spatial_lowpass_cutoff, normalized with respect to the average cell radius in the movie. A larger value of this parameter leads to mild filtering, with no filtering being performed when it is set to infinity. Default: 2.
- cellfind_min_snr and thresholds.T_min_snr: Two different definitions of signal-to-noise ratio for cells' estimated $Ca^{2+}$ activity traces. The former is used only during cell finding, whereas the letter is used for both cell finding and cell refinement. A cell's estimated signal quality should be higher than these values. For cellfind_min_snr, a typical value of 1 is reasonable for most datasets. Increase this value to decrease false positives during cell finding with no effect on the cell refinement procedure. For thresholds.T_min_snr, a default value is 7 (and lowest acceptable value is 3). This parameter is often very effective to cull away false-positives during cell refinement (**§6**).
- cellfind_max_steps: The maximum number of cells that can be initialized during the cell finding of EXTRACT in each spatial partition. The default is 1000, which may need to be adjusted depending on the particular dataset. Specifically, it can be increased so that EXTRACT finds more cells per partition or decreased to prevent increased false positives.
- cellfind_kappa_std_ratio and cellfind_adaptive_kappa: The robustness parameter, $\kappa$, utilized during the cell finding module is initially set to the estimated noise in the movie times cellfind_kappa_std_ratio. Large values of $\kappa$ correspond to assuming low contamination by non-Gaussian noise, setting $\kappa = \infty$ leads to least squares estimates. When cellfind_adaptive_kappa is set to one, this parameter is adaptively estimated from the $Ca^{2+}$ imaging movie. Note that these parameters affect only

the cell finding module, there are parameters for the cell refinement and final robust regression modules. Default: $\kappa = 0.7$, the adaptive estimation is off by default.

- `init_with_gaussian`: If true, then during cell finding, each cell is initialized with a gaussian shape prior to robust estimation of their spatial profiles. If false, then initialization is done by computing a correlation image. If cells are of different shape and size, keep this off. Default: 0.
- `avg_yield_threshold`: During cell finding, If the yield in the last few components falls below this threshold, the cell finding module is terminated. Default: 1/10.
- `visualize_cellfinding`: To visualize the cell finding process, turn this flag on. This parameter is particularly useful for optimizing hyperparameters (**§5**), but increases the runtimes significantly. Default: 0.
- `S_init`: Optionally, one can provide cells' profiles with this parameter and EXTRACT will use these as the initial set of cells, skipping the cell finding module. Default: none.

#### 3.4. Cell refinement module

The main purpose of the cell refinement module is to correctly estimate the true cells' spatial profiles and delete the spurious and/or duplicate cells. Here, we first provide the list of general hyperparameters of the cell refinement module:

- `kappa_std_ratio`: The robustness parameter, $\kappa$, utilized during the cell refinement module is initially set to the estimated noise in the movie times `kappa_std_ratio`. Large values of $\kappa$ correspond to assuming low contamination by non-Gaussian noise, setting $\kappa = \infty$ leads to least squares estimates. Default: 0.7.
- `adaptive_kappa`: This parameter can take three distinct values. 0: No adaptive estimation of the robustness parameter during cell refinement or final robust regression. 1: Adaptive estimation is performed during the final robust regression only. 2: Adaptive estimation is performed during both cell refinement and final robust regression. Default: 1.
- `max_iter`: The number of cell refinement iterations. Default : 6.
- `l1_penalty_factor`: A scalar that determines the strength of $L_1$ regularization penalty to be applied when estimating the temporal components. The penalty is applied only to cells that overlap in space and whose temporal components are correlated. Use larger values if spurious cells are observed in the vicinity of high SNR cells, particularly helpful for culling out duplicate cells. Default: 0.

Inside the cell refinement module, EXTRACT computes several quality metrics and discards cell candidates that fail to score sufficiently. The thresholds on these quality metrics are determined by the outside user inside the structure array: `thresholds`. The following parameters controlling these quality metrics are most relevant for EXTRACT users:

- `T_min_snr`: Cells with lower SNR value than this parameter will be eliminated. This parameter is shared with the cell finding module, see above. Default: 7.
- `size_lower_limit` and `size_upper_limit`: These factors are multiplied with the average cell area (determined from `avg_cell_radius`) and any cell with an area outside of these will be eliminated during the cell refinement. Defaults: 0.1 and 10.
- `spatial_corrupt_thresh`: Spatial corruption indices, quantifying the non-uniformity in cells' spatial profiles, are calculated at each step of the alternating minimization routine. Images that have an index higher than these are eliminated. Default: 1.5.
- `eccent_thresh`: Cells with eccentricity higher than this will be eliminated. One can intuitively think of high eccentricity as cells becoming more elliptic. This parameter eliminates blood vessels in most one-photon $Ca^{2+}$ imaging movies. Default: 6.
- `T_dup_corr_thresh` and `S_dup_corr_thresh`: Through alternating estimation, cells that have higher trace correlation than `T_dup_corr_thresh` and higher image correlation than `S_dup_corr_thresh` are eliminated. Defaults: 0.95 and 0.8.
- `low_ST_index_thresh`: EXTRACT requires cells' $Ca^{2+}$ activity traces to explain a portion of the fluorescence signal in the pixels encapsulated by the cells. This parameter culls out the identified components, in which the inferred $Ca^{2+}$ activity traces do not explain the pixel activities well. In practice, this parameter may be set to '0.01' and is very helpful for removing garbage at the expense of culling out low SNR cells. In most movies, this parameter is not needed. See **Spatiotemporal match metrics** Section in the **Methods** for additional details. Default: -1 (no checks).

#### 3.5. Final robust regression module

The final robust regression module performs, essentially, one final robust regression to estimate cells' $Ca^{2+}$ activity traces. Below are the relevant parameters:

- `Regression_only`: When this flag is on, EXTRACT becomes a $Ca^{2+}$ activity trace estimator. Depending on the `trace_output_option`, EXTRACT can be used to perform robust regression, least-squares regression, and/or non-negative least-squares regression. Cell finding and refinement modules will be skipped. `S_init` should be initialized with the cells' spatial profiles, over which the regression will be performed. Default: 0.

- `trace_output_option`: This parameter decides which type of solver EXTRACT will use to produce the final estimated $Ca^{2+}$ activity traces (**§7**). Viable options include: `'nonneg'`, `'baseline_adjusted'`, `'no_constraint'`, `'nonnegative_least_squares'`, `'least_squares'`. Default: `'baseline_adjusted'`.
- `trace_quantile`: When `'baseline_adjusted'` solver is used, EXTRACT uses an adaptive process to compute the cells' activity baselines. This parameter provides the percentile, which will be used as the cells' activity baselines. Default: 0.25.

#### 3.6. Parameters for speed optimization

EXTRACT default parameters are optimized for accuracy. In many cases, additional speed improvements can be achieved by simple tricks that do not affect the cell extraction quality. Below are the relevant parameters:

- `cellfind_max_iter`: Number of alternating estimations during the one-by-one cell finding process. Default is 10, but decreasing down to 3 often leads to little to no performance decreases.
- `max_iter_S` and `max_iter_T`: Maximum number of iterations for S and T estimation steps during the cell finding. Both are 100 by default, but decreasing down to 20 is usually perfectly acceptable, since the robust regression converges fast.
- `max_iter_T_final`: This parameter is used to set the maximum number of ADMM iterations (See **Methods**) at the final regression. Most cases do not require any adjustments, be careful when decreasing to gain speed as final activity traces are obtained through this step. If this number is too small, the cells' $Ca^{2+}$ activity traces may appear distorted at their maximum values due to insufficient estimation iterations. Default: 100.

#### 3.7. Tutorial 2: Parallelizing cell extraction across multiple CPU cores

In this brief tutorial, we show the importance of running EXTRACT on multiple GPUs and/or CPU cores. The computer we ran this tutorial on, whose results will be shown as part of the live script in our Github[1], has two NVIDIA Geforce RTX 3090s and an Intel i9-10980XE CPU with 18 physical cores. To start with, we first simulate an example two-photon $Ca^{2+}$ imaging movie using the code:

```
if ~isfile('Example_2p_movie.h5')
    [opts_2p] = get_2p_defaults();
    opts_2p.ns = 500;
    rng(1)
    create_2p_movie(opts_2p,'Example_2p_movie');
End
```

This code simulates a two-photon $Ca^{2+}$ imaging movie with a $500 \times 500$ pixels$^2$ field-of-view. In the upcoming tutorials, we will show how to optimize EXTRACT to achieve optimal cell extraction. For this tutorial, our goal is to showcase the speed of EXTRACT under various conditions. Thus, we will use the already optimized parameters:

```
M = 'Example_2p_movie.h5:/mov';
config = get_defaults([]);
config.adaptive_kappa = 2;
config.spatial_highpass_cutoff = inf;
config.downsample_time_by = 4;
config.num_partitions_x = 4;
config.num_partitions_y = 4;
config.thresholds.T_dup_corr_thresh = 0.99;
config.thresholds.spatial_corrupt_thresh = 0.1;
config.thresholds.eccent_thresh = 3;
config.max_iter = 10;
config.thresholds.size_upper_limit = 3;
config.cellfind_min_snr = 0;
```

```
config.verbose = 0;
config.trace_output_option = 'no_constraint';
```

To start with, we run this movie on a single CPU core using the code:

```
config.use_gpu = 0;
output = extractor(M,config);
```

This code, on our computer, took around 15 minutes to complete. In a similar manner, by using the parameters `use_gpu`, `parallel_cpu`, `multi_gpu`, and `num_workers`, we ran the same movie on 16 CPU cores (7 mins), a single GPU (5 mins), and two GPUs (3.8 mins).

Overall, GPU performance was best, which was followed by parallelization across CPU cores. Please note that the runtimes are mainly due to some fixed costs (also shown in **Figure 5K**). These extra costs are negligible for large scale movies (as in **Figure 4**), but prevent major speedups with multiple GPUs for short movies like the one we considered here.

#### §4. Prerequisites: Pre-processing of movies

In this section, we will mainly be discussing the **Tutorial 3**, which provides a brief introduction to EXTRACT's preprocessing module. Here, we discuss loading (and watching) $Ca^{2+}$ imaging movies from the h5 files, fast spatial high-pass filtering using EXTRACT's GPU accelerated bandpass filter, and other supporting functions for the preprocessing pipeline.

As before, this tutorial is provided in our Github repository[1]. We start by simulating a one-photon $Ca^{2+}$ imaging movie using the following code:

```
if ~isfile('Example_1p_movie.h5')
   [opts_2p,opts_back] = get_1p_defaults();
   opts_2p.ns = 100;
   opts_2p.n_cell = 100;
   opts_back.ns = 100;
   opts_back.n_cell = 20;
```

```
    opts_back.cell_radius = [20,40];
    rng(1)
    create_1p_movie(opts_2p,opts_back,'Example_1p_movie');
end
```

Throughout this tutorial, we use this simulated movie to introduce the most important aspects of the preprocessing module.

#### 4.1. Reading and watching the $Ca^{2+}$ imaging movies

For EXTRACT, we recommend to use the '.h5' format for storing and reading $Ca^{2+}$ imaging movies. Each '.h5' file is accompanied by at least one dataset, which are conventionally named with a backslash, *e.g.*, '`/example_dataset`'. In our example, the dataset that contains the movie is called '`/mov`'. If unsure, one can always check the dataset name and size using the following command:

```
info = h5info('Example_1p_movie.h5');
```

The movie can be read using MATLAB's native `h5read` function. The `info` array above tells us that the movie matrix has the shape $100 \times 100 \times 5000$. We can read the first 1000 frames using the following code:

```
M = h5read('Example_1p_movie.h5','/mov',[1,1,1],[100,100,1000]);
```

Here, the first entry is the name of the h5 file, the second entry dataset, the third entry the start coordinates, and the final entry is the how much to include in that coordinate, reading in rectangular chunks. If the third and forth entries are not given, h5read reads the full movie into RAM memory.

Once loaded on the RAM memory, the $Ca^{2+}$ imaging movie can be watched using the native EXTRACT function:

```
view_movie(M)
```

It is often good practice to watch the raw $Ca^{2+}$ imaging movies for potential motion and/or experimental artifacts before starting the cell extraction process.

#### 4.2. Spatial high-pass filtering in EXTRACT

Prior work utilizes distinct strategies to either model the background contamination[2] or explicitly remove neuropil[3]. With EXTRACT, we use a principled approach that can remove spatially correlated neuropil activities while retaining the cells' $Ca^{2+}$ activity signals, *i.e.*, utilize spatial high-pass filtering. Within EXTRACT's preprocessing module (`preprocess_movie.m`), we have coded a GPU accelerated band-pass filter, which can be applied as a spatial high-pass filter with the following code:

```
avg_cell_radius = 6;
spatial_highpass_cutoff = 5;
spatial_lowpass_cutoff = inf;
use_gpu = 0;
M_proc = spatial_bandpass(M, avg_cell_radius, ...
 spatial_highpass_cutoff, spatial_lowpass_cutoff, ...
 use_gpu);
view_movie(M_proc)
```

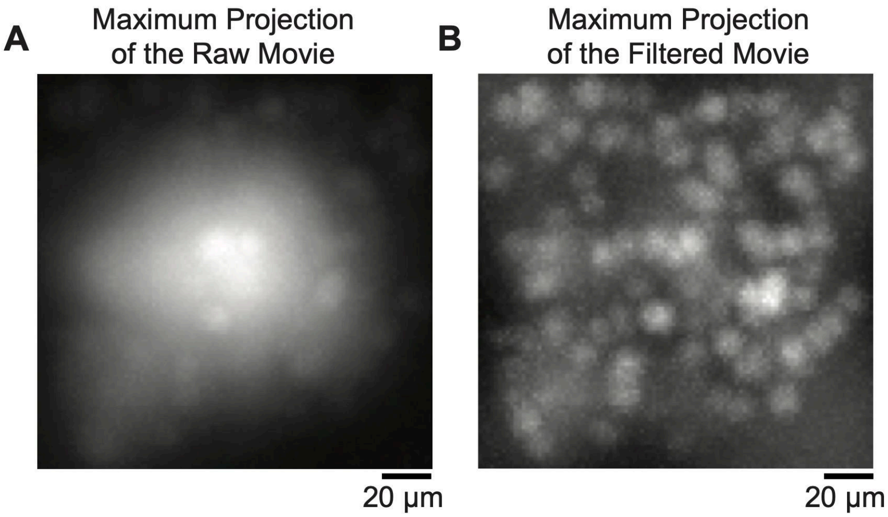

**Appendix Figure 12. Spatial high-pass filtering removes global neuropil contamination.**
**A** The maximum projection of the raw simulated one-photon $Ca^{2+}$ imaging movie shows signs of global contamination. **B** Performing a spatial high-pass filtering on the full movie removes the global background and allows visual identification of the cells' spatial profiles.

As shown in **Appendix Figure 12**, the spatial filtering is particularly useful for removing global background. As we show next, this process does not remove cells' $Ca^{2+}$ events (**Appendix Figure 13**), which is theoretically consistent with the choice of the cutoff threshold.

#### 4.3. Preprocessing and saving movies in batches

When large $Ca^{2+}$ imaging movies need to be processed in batches, the preprocessing module of EXTRACT can be run in an offline manner, and by looping through the movie frames to prevent memory errors. This can be achieved with a few lines of code:

```
config = get_defaults([]);
config.partition_size_time = 500;
preprocess_save('Example_1p_movie.h5:/mov',config)
```

We conclude the **Tutorial 3** with an example cell extraction with this preprocessed movie, whose results are shown in the accompanying live script (See our Github repository) and **Appendix Figure 13**.

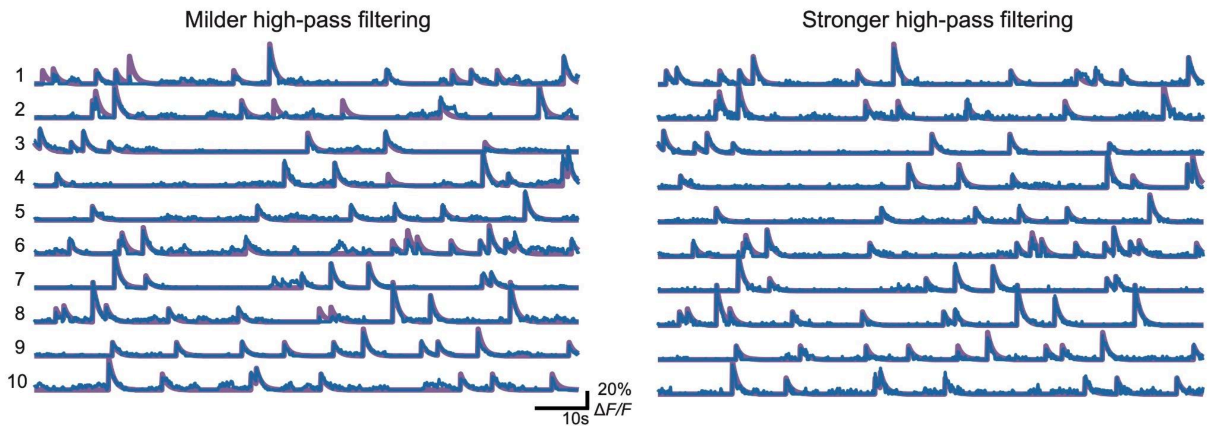

**Appendix Figure 13. Spatial high-pass filtering does not remove cells' $Ca^{2+}$ events.**
Since the cutoff value for the spatial high-pass filtering is chosen far away from the spatial frequencies containing cells, the spiking events are not removed from the $Ca^{2+}$ activity traces. In fact, in certain cases, stronger high-pass filtering may allow higher quality cell extraction, and therefore better recovery of the spiking events in noisy $Ca^{2+}$ imaging movies. Here, we are using a simulated movie with extensive global neuropil contamination.

### §5.Two-step hyperparameter optimization of the cell finding and refinement modules

EXTRACT's cell finding and refinement modules can be optimized in a two-step optimization This section primarily discusses **Tutorials 4** and **5**, where we discuss the optimization of hyperparameters for EXTRACT's cell finding and refinement modules, respectively. For these tutorials, we use a low SNR two-photon $Ca^{2+}$ imaging movie, courtesy of Parker Lab[4].

#### 5.1. Step one: Optimizing the cell finding module

Given the modularity of EXTRACT's cell finding module, we can optimize its hyperparameters in a modular and decoupled manner in **Tutorial 4**. We start by preprocessing the raw $Ca^{2+}$ imaging movie and watching the resulting filtered version for potential artifacts via the code:

```
M = h5read('jones.h5','/data');
config = get_defaults([]);
M_proc = preprocess_movie(M,config);
view_movie(M_proc(:,:,1:100))
```

This particular movie has rather low signal and high levels of noise, making it challenging to process with the default values. To test this, we can turn off the cell refinement process and watch the cell extraction real-time as it is happening with following the code:

```
config.max_iter = 0;
config.visualize_cellfinding = 1;
output = extractor(M,config);
```

As shown in the accompanying live script[1], EXTRACT's cell finding module fails to identify several cell candidates that are visible to the naked eye.

To optimize the cell finding module, we first note that the earlier run had shown an early cutoff for the trace snr values, indicating that more cells might have been found if `thresholds.T_min_snr` had a lower SNR threshold. Second, the `cellfind_min_snr` value was centered around 5 (also see **Appendix Figure 14** below), which is often at the order of tens to hundreds. Thus, as a first step, we pick very low values for both parameters. Next, since our main concern is being able to identify low SNR cells, we also decrease `spatial_lowpass_cutoff` value, which performs a low-pass filtering and smooths the movie for cell finding purposes only:

```
M = 'jones.h5:/data';
config = get_defaults([]);
config.downsample_time_by = 4;
config.spatial_lowpass_cutoff = 1;
config.use_gpu = 0;
config.max_iter = 0;
config.visualize_cellfinding = 1;
config.cellfind_min_snr = 0;
config.thresholds.T_min_snr = 3.5;
output = extractor(M,config);
```

The results, shown in **Appendix Figure 14,** contain several duplicate and/or spurious cell candidates, which need to be removed by the cell refinement module that we discuss next.

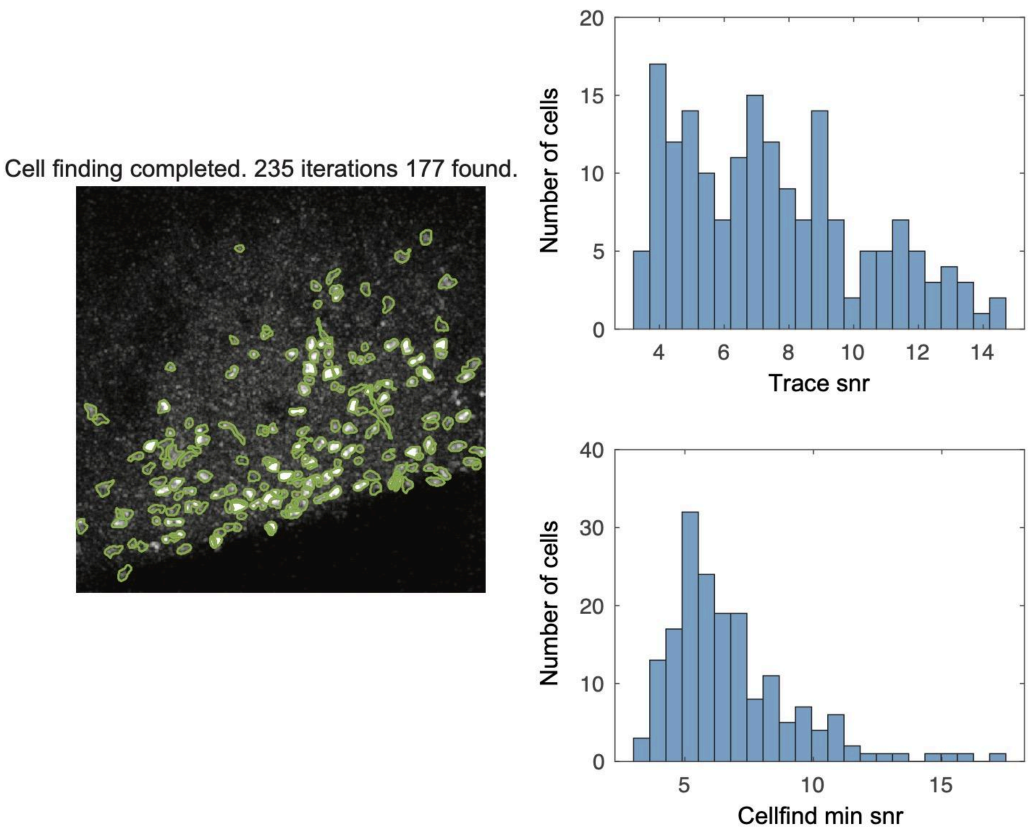

**Appendix Figure 14. The dynamical visualization of the cell finding module.**

If `visualize_cellfinding = 1`, EXTRACT provides a dynamical visualization of the cell finding process. Here, the cells are identified on a projection map on the left and the cell finding metrics are presented real-time on the right. In this tutorial, multiple low SNR cells (with SNR < 7) were identified that would have been rejected by the default values.

#### 5.2. Step two: Optimizing the cell refinement module

Having discussed the first step of the optimization routine, with **Tutorial 5**, we now focus on the cell refinement module. Within EXTRACT, we have a flag for optimizing the cell refinement module (`hyperparameter_tuning_flag`). When this parameter is true, EXTRACT performs a single cell refinement step and saves the quality metrics. Then, the function, `plot_hyperparameter_curves.m`, can be used to plot the quality metrics:

```
M = 'jones.h5:/data';
config = get_defaults([]);
config.downsample_time_by = 4;
config.spatial_lowpass_cutoff = 1;
```

```
config.use_gpu = 0;
config.hyperparameter_tuning_flag = 1;
config.cellfind_min_snr = 0;
config.thresholds.T_min_snr = 3.5;
config.cellfind_min_snr = 0;
config.adaptive_kappa = 2;
output = extractor(M,config);
plot_hyperparameter_curves(output)
```

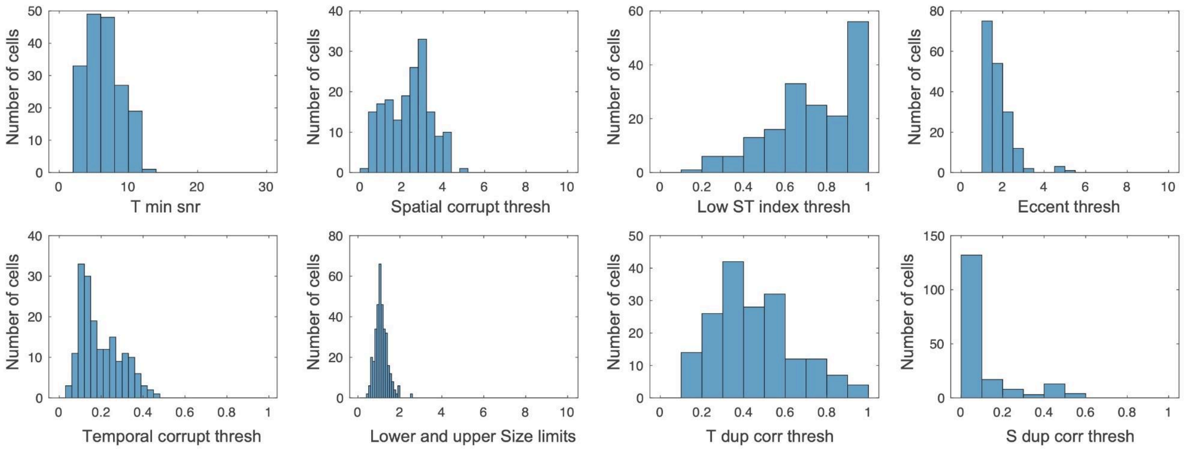

**Appendix Figure 15. A summary of the trace quality metrics for hyperparameter tuning.**
When `hyperparameter_tuning_flag = 1`, EXTRACT performs only a single cell refinement step and saves the computed trace quality metrics. Using the helper function `plot_hyperparameter_curves.m`, the user can visualize these metrics and broadly decide on the first guess of the thresholding parameters.

The resulting plots (shown in **Appendix Figure 15**) constitute the starting points for the tuning of the quality metric thresholds:

- Firstly, many data points in the `T_min_snr` curve seem to be right at the threshold. We might wish to lower it further (here down to 3.2) and check if it prevents discarding true positive cells.
- Next, the spatial corruption values in these plots are at ~ 4-5, much higher than the default value of 1.5. This is expected for very noisy movies. We need to increase this threshold value if we wish to keep the true cells (here up to 5).
- Next, the extracted cell areas seem to be rather small, usually a sign that either our average cell radius is quite off or that `kappa_std_ratio` is too strict. We changed the latter to 1 in this example, which helped recover some incorrectly discarded cells.

- We picked the number of iterations to be 10, which was empirically when the cell profiles stabilized (you can check this by setting `visualize_cellfinding = 1`).
- Moreover, we kept the adaptive estimation for cell refinement (`adaptive_kappa = 2`), which was particularly helpful for this low SNR movie.
- Finally, we observe a long tail in the `T_dup_corr_threshold` curves, which is why we set the correlation threshold to 0.8. We finetuned this value while checking the final outputs for a few runs and ensuring that duplicate cells were properly discarded.

With that, we were able to optimize the cell refinement module with a few runs, each took around 2-3 minutes. For larger movies, the same procedure can be performed on a smaller chunk of the movie for faster optimization.

#### 5.3. The final cell extraction result

Bringing all together, we ran EXTRACT on the full movie after a quick optimization, which gave rise to the following code:

```
M = 'jones.h5:/data';
config = get_defaults([]);
config.downsample_time_by = 4;
config.spatial_lowpass_cutoff = 1;
config.use_gpu = 0;
config.max_iter = 10;
config.cellfind_min_snr = 0;
config.thresholds.T_min_snr = 3.2;
config.thresholds.spatial_corrupt_thresh = 5;
config.thresholds.T_dup_corr_thresh = 0.8;
config.adaptive_kappa = 2;
config.kappa_std_ratio = 1;
output = extractor(M,config);
figure
plot_output_cellmap(output,[],[],'clim_scale',[0.2, 0.999])
figure
T_ex = output.temporal_weights';
```

```
plot_stacked_traces_double(T_ex(1:31,:),[],0)
```

The cell extraction results are shown in **Appendix Figure 16**. In our experience, these optimized parameters are robust and often perform well across days and mice, as long as the imaging and experimental conditions are not significantly different.

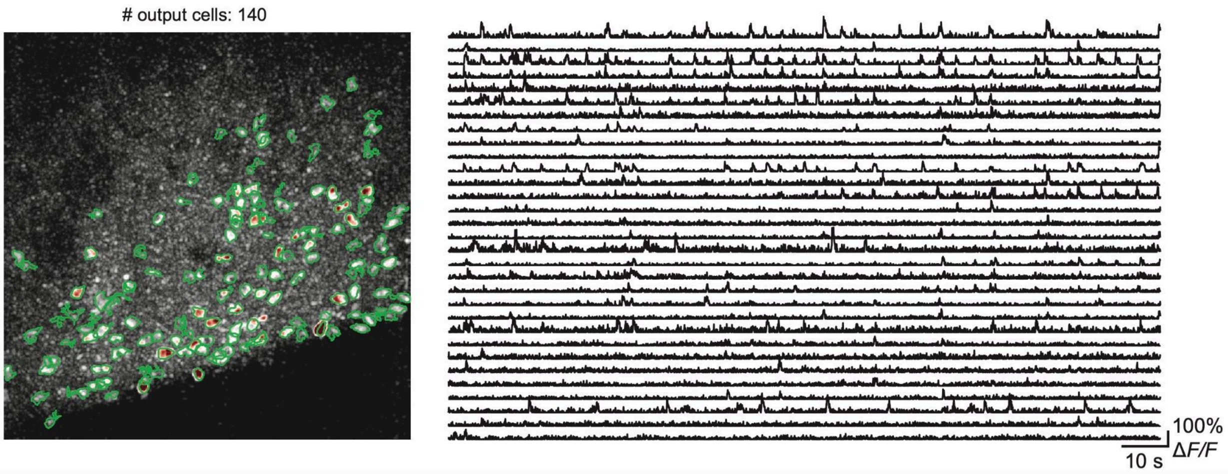

**Appendix Figure 16. Visualization of the final cell extraction results with EXTRACT.**
After optimization, EXTRACT provided high quality cell extraction results with little to no spurious or duplicate cells (left) and the $Ca^{2+}$ activity traces from the first 30 extracted cells (right).

#### §6. How to use the final robust regression module as a stand-alone tool

Apart from being utilized as a cell extraction tool, EXTRACT can also be used as a post-processing tool to obtain $Ca^{2+}$ activity traces via robust regression given the cells' spatial profiles. In **Tutorial 6**, we quickly show how to configure EXTRACT to do this by using the movie from the first tutorial:

```
load('example.mat');
M = M-1; % ground truth movie with F = 0;
```

We initialize EXTRACT with the first 15 cells, whereas the other 5 are undetected, simulating the existence of neuropil and other contamination sources. Then, we can compute both the robust and the least-squares estimates of $Ca^{2+}$ activity traces with the following code:

```
config=[];
```

```
config = get_defaults(config);
config.avg_cell_radius=7;
S = S_ground(:,1:15);
config.S_init = S;
config.preprocess = 0; %using a preprocessed movie
config.F_per_pixel = ones(50,50); % F values are all one
config.trace_output_option='no_constraint';
config.use_gpu=0;
config.regression_only = 1;
output_rb=extractor(M,config);
T_ex = output_rb.temporal_weights';
config.trace_output_option='least_squares';
output_ls=extractor(M,config);
T_ls = output_ls.temporal_weights';
```

Here, once `regression_only=1`, EXTRACT skips the cell finding and refinement modules. In this scenario, `S_init` should contain the spatial profiles (2D or 3D, both acceptable). The results are shown in **Appendix Figure 17**. Note that both estimates are imperfect, though robust regression preferentially suppresses the contamination from unidentified cells.

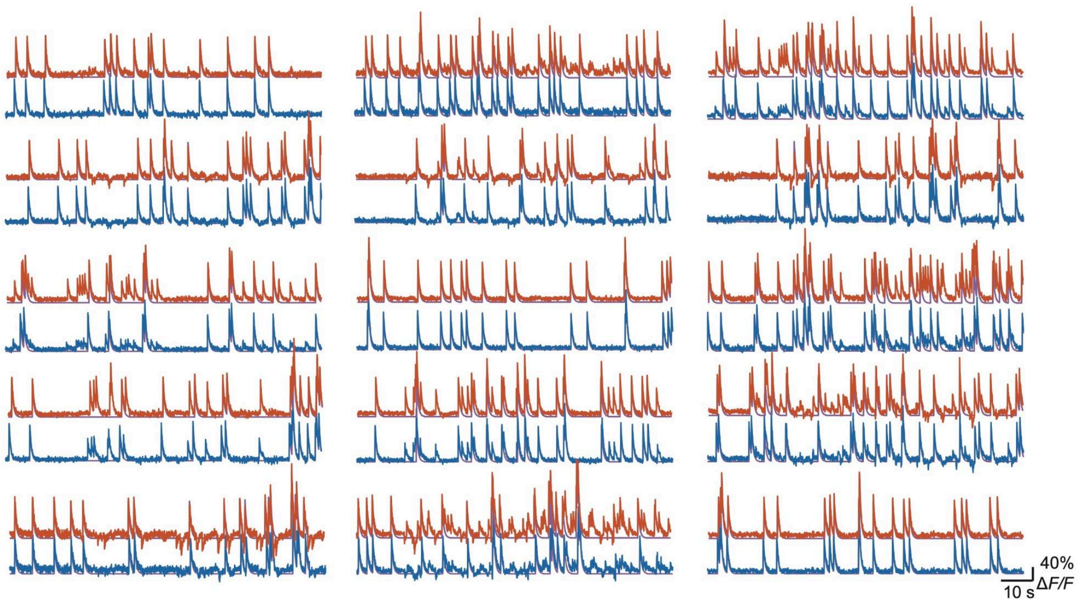

**Appendix Figure 17. EXTRACT as a post-processing tool for estimating $Ca^{2+}$ activity traces.**
Using the movie from **Tutorial 1**, we estimated cells’ $Ca^{2+}$ activity traces by using EXTRACT as a post-processing tool. We initialized EXTRACT with 15 out of 20 cells and estimated the $Ca^{2+}$ activity traces with robust (blue) and least-squares (red) regression. Though both algorithms showed occasional crosstalk, robust regression led to preferential suppression of contamination and consequently to more accurate $Ca^{2+}$ activity traces.

### §7. Conclusion

In this document, we provided a quick start guide for EXTRACT and introduced the two-step optimization routine that we regularly use to quickly optimize its hyperparameters. We have shared six tutorials in our accompanying repository and a descriptive figure (**Figure S9**), which should hopefully help first-time users get started with EXTRACT. For any additional questions or comments, please open an issue in the Github repository or contact via. We also occasionally hold tutorial sessions to help new labs get started, which you can request via the same email.
